# *Pseudomonas* can prevent the parasitic fungus, while keeping the crop fungus unaffected, in the gardens of *Odontotermes obesus*

**DOI:** 10.1101/2021.01.29.428757

**Authors:** Renuka Agarwal, Manisha Gupta, Abin Antony, Ruchira Sen, Rhitoban Raychoudhury

## Abstract

Insects that farm monocultures of fungi are canonical examples of nutritional symbiosis as well as independent evolution of agriculture in non-human animals. But just like in human agriculture, these fungal crops face constant threat of invasion by weeds which, if unchecked, takes over the crop fungus. In fungus-growing termites, the crop fungus (*Termitomyces*) faces such challenges from the parasitic fungus *Pseudoxylaria*. The mechanism by which *Pseudoxylaria* is suppressed is not known. However, evidence suggests that some bacterial secondary symbionts can serve as defensive mutualists by preventing the growth of *Pseudoxylaria*. However, such secondary symbionts must possess the dual, yet contrasting, capabilities of suppressing the weedy fungus while keeping the growth of the crop fungus unaffected. This study describes the isolation, identification and culture-dependent estimation of the roles of several such putative defensive mutualists from the colonies of the wide-spread fungus-growing termite from India, *Odontotermes obesus*. From the 38 bacterial cultures tested, a strain of *Pseudomonas* showed significantly greater suppression of the weedy fungus than the crop fungus. Moreover, a 16S rRNA pan-microbiome survey, using the Nanopore platform, revealed *Pseudomonas* to be a part of the core microbiota of *Odontotermes obesus*. A meta-analysis of microbiota composition across different species of *Odontotermes* also confirms the wide-spread prevalence of *Pseudomonas* within this termite. These evidence indicate that *Pseudomonas* could be playing the role of defensive mutualist within *Odontotermes*.

## Introduction

Domestication of specific fungal cultivars by insects, for food, has evolved independently over the past 50 million years in three major insect lineages of ants, termites, and beetles [1-3]. Curiously, fungus-growing ants and termites show parallel symbiotic evolution in two distinct geographical locations. The former is found in the new world, whereas, the latter (Subfamily: Macrotermitinae) are found in tropical Africa and Asia [4]. However, both systems face the invasion of parasitic fungi taking over their crop. In fungus-growing termites, the most prominent of these parasites is *Pseudoxylaria* (Phylum: Ascomycota) which invade the monoculture of the crop fungus, *Termitomyces* (Phylum: Basidiomycota) [2, 5-7]. Termites have been reported to regulate this invasion by actively grooming out any unwanted spores [8], selectively burying the parasitic fungi [9] and applying chemical secretions with specific anti-fungal properties [10]. However, the effectiveness of these measures in controlling *Pseudoxylaria* remains unknown [4, 11].

The presence of fungicide-producing mutualistic microbes in fungus-growing ants, which arguably suppresses the growth of parasitic fungi in their colonies [12-15], points to a similar role of secondary bacterial symbionts in fungus-growing termites as well [16, 17]. This conjecture is strengthened by the fact that even after cultivating very divergent crop fungi, the microbiota across all insect-fungal systems shows a remarkable phylogenetic and compositional similarity [18, 19]. This multi-level convergent evolution suggests the importance of such secondary microbes in maintaining successful nutritional symbiosis across all fungus-growing insects [18, 19]. The evolutionary logic for the presence of such mutualistic microbes within these symbioses is inescapable. Since, the termites cannot subsist on any fungi other than *Termitomyces* [20, 21] and *Pseudoxylaria* has been found in almost all fungus-growing termites [22], colonies with such mutualistic microbes would have a strong selective advantage. Consequently, many strains of *Bacillus* and *Streptomyces* have been found from termite colonies which can act against *Pseudoxylaria* [11, 23-26]. But, most of these strains also inhibit the growth of the cultivar fungus [23, 24].

This indicates that such a secondary symbiont cannot be a generic anti-fungal agent as it will also adversely affect the crop fungus on which the termites subsist. Thus, a potential mutualist must have the dual, but contrasting, capability of acting as an antagonist towards *Pseudoxylaria*, but, as a mutualist towards *Termitomyces*. Unfortunately, there is no clear and unambiguous evidence of such a microbe from fungus-growing termites. Moreover, the presence of such microbes has not been demonstrated across the core microbiota of individual termite populations as these surveys are either from the gut or from the comb [11, 27]. But if these mutualists are indeed essential then they must be present within the different castes as well as the fungus comb of the termite. This is especially true for winged reproductives (alates) which are the mobile castes of termites that move out of the natal colony to find new ones. Alates are presumed to be equipped with the full complement of the microbiota required for the successful establishment of new colonies and would be selected against if they lack such critical secondary mutualists [28, 29]. Thus, the presence of any such a secondary symbiont should permeate across the different castes as well as the comb. However, since no such analysis of caste-specific microbiota has been reported from any fungus-growing termites, it is not possible to ascertain whether the previously reported microbes are an essential part of the symbiosis.

In this study, we present evidence of the presence of several potential mutualistic bacteria within the core microbial community of the fungus-growing termite *Odontotermes obesus*. We isolated 38 different bacterial cultures from different castes and the comb and tested their potential to be an antagonist towards *Pseudoxylaria*. Then we tested some of these antagonists to ascertain whether they also impede the growth of the crop fungus. A particular strain of *Pseudomonas* appears from these two contrasting screens to be an ideal mutualist as it strongly impedes the growth of *Pseudoxylaria* but show almost negligible inhibition of *Termitomyces*. To test whether *Pseudomonas* is a part of the symbiotic community, we also enumerated the pan-microbiome of *O. obesus* by obtaining Nanopore reads of 16S rRNA gene fragments (V3-V4 regions) from six different samples (comb, both male and female alates, both major and minor workers and nymphal castes). From this pan-microbiome, we then established the presence of *Pseudomonas* within the core microbiota of *O. obesus*. Finally, we performed a meta-analysis of microbiota across many different *Odontotermes* sp. and ascertain the ubiquity *Pseudomonas* in this termite genus.

## Materials and Methods

### Source of termites, combs and their DNA extraction

Different castes, nymphs and fungus combs of *O. obesus* were collected from two different mounds of the IISER Mohali campus (Fig. S1-S3). The combs were collected in sterile plastic bags, termites were removed and then crushed within them. These were divided into 0.5 g portions in sterile glass containers. To minimize environmental contamination and degradation, DNA was extracted from some of these portions within 3 hours of the collection while others were used for microbial isolation. Major workers, minor workers and nymphs collected from these combs were also kept in separate sterile vials. Male and female alates were collected during swarming, which began after the first rains during June 2018. DNA was extracted from individual termite using the CTAB (Cetyl Trimethyl Ammonium Bromide) method [30] (supplementary methods S1).

DNA extractions from fungus combs often have extensive humic acid contamination which can hinder downstream PCR reactions. To remove this humic acid, the precipitation step with Phenol: Chloroform: Isoamyl alcohol were repeated until the final pellet obtained was white in color [11].

### Isolation and identification of bacterial strains

Workers, nymphs, and alates were washed in the sterilized water to clean them of soil particles and then homogenized in 500 µl of sterile 1X PBS buffer with sterile plastic pestles. 0.5 gm portions of comb were also similarly homogenized. A dilution series (1x -10^−6^x) of these homogenates were then plated on Luria Bertani Agar (LBA) and Tryptic Soya Agar (TSA) for any bacterial growth at 30°C for 24 hrs. For strain identification, DNA from individual bacterial colonies was extracted with CTAB buffer and amplified with the primer set 341F*/*806R [31] for a portion of the 16S rRNA gene (supplementary methods S1). The sequences thus obtained were used to identify the bacterial strains by a BLAST search in NCBI. These were submitted to GenBank (accession number MN908295-MN908332). Since, multiple strains within each bacterial genus were obtained, the nomenclature of these strains were modified to read as ‘Genera-NCBI Accession Number’ (Table S1).

### Bacterial-fungal interactions

Bacterial growth curves for all the 38 isolated strains were determined at 30°C using Freedom EVO (Tecan Life Sciences) to ensure that interaction with the fungi was done when they were in their respective log phases. To identify whether these bacteria prevent the growth of the parasitic fungus, paper disc diffusion assay was used in 90 mm diameter Petri dishes containing Potato Dextrose Agar media (PDA) [23]. Paper discs soaked with 10 µl of bacterial cultures were inoculated at the center of the Petri dish along with actively growing *Pseudoxylaria* in two plugs. Control growth plates were set up for both the bacteria and fungus and incubated in dark at 30°C in a Heratherm™ Compact Microbiological Incubator (Thermo Fisher Scientific). The magnitude of inhibition by bacteria was evaluated by using the following formula from Royse and Ries [32]:

Magnitude of Inhibition = (Total Area of fungal growth in the control in mm^2^) – (Total Area of fungal growth in the interaction in mm^2^) / Total Area of fungal growth in the control in mm^2^

This assay was slightly modified for interaction against *Termitomyces*, as it grows much more slowly than *Pseudoxylaria*. While *Pseudoxylaria* plugs can fill up a control Petri plate within 5 days, *Termitomyces* plugs can take over 30 days to do so (Fig. S4). To account for this disparity, a liquid suspension of *Termitomyces* nodules and mycelia was prepared by homogenizing a 4 cm^2^ block of actively growing culture in sterile 1X PBS and spread on plates with Potato-Dextrose with Yeast Malt (PYME) Agar [33]. These plates were first incubated for 36 hours at 30°C, before adding bacterial discs [24]. Another modification was the use of four bacterial discs against *Termitomyces* compared to a single bacterial disc against *Pseudoxylaria*. This enabled the establishment of contact between *Termitomyces* and the bacterial strains much faster than would have been possible with the use of single discs. These two modifications standardized the growing conditions of the two fungi so that final data could be taken on the 7^th^ day after inoculation. All the interaction plates were evaluated for any type of inhibition every 24 hours for 7 days post-inoculation by taking photographs using the Panasonic Lumix G2 camera (Fig. S5). Photographs from the 7^th^ day were counted as the final data and were analyzed in Adobe Photoshop CS6 by measuring the area of fungal growth in pixels and converting it into millimeters (100 pixels = 1 mm^2^). All interaction assays were repeated with sample sizes ranging from 3-5 plates. Representative figures for all the bacterial-fungal interaction assays are given in supplementary figures (Fig. S6-S8).

### Sample preparation, running the Nanopore platform and obtaining sequences

To identify the microbiota present within termite castes and fungus combs, the V3-V4 region of the 16S rRNA gene was amplified using primers 341F/806R [31]. Six different DNA samples (major worker, minor worker, nymph, male alate, female alate and comb) were prepared for pan-microbiome analysis on the Nanopore platform (supplementary methods S1).

Samples were sequenced on a MinION platform (MinION Mk1B) on FLO-MIN106 flowcells for 48 hours using MinKNOW software with the protocol *NC_48Hr_sequencing_FLO-MIN106_SQK-LSK108_plus_Basecaller*. After the completion of the run, sequences were separated and trimmed according to their barcodes and quality (supplementary methods S1). Cleaned sequences were deposited to NCBI Sequence Read Archive (SRA) under the BioProject PRJNA608773 (BioSample accessions: SAMN14208512 - SAMN14208517).

### Taxonomic identification of the pan-microbiome

A customized microbial repository was made by combining all the available 16S rRNA gene fragments from NCBI FTP site and the DictDb v 3.0 database [34]. LAST v 973 [35], was used to identify the sequences with the following parameters: match score of 1, gap opening penalty of 1, and gap extension penalty of 1 [36]. To estimate the sequencing depth rarefaction curves were generated with the Vegan package v 2.5-4 in R v 3.6.2. The estimated community richness (Chao1) and diversity indices (Evenness, Shannon, Simpson, and Inv Simpson) [37-39] were also calculated in R.

We compared the gut and comb microbiota from the other known *Odontotermes* sp. [31, 40, 41] with the results obtained from this study using weighted principle coordinate analysis (PCoA) in phyloseq package of R. Nineteen different microbiota were used for the similarity analysis as these also amplified the same 16S rRNA gene region and used the same reference database.

## Results

### Eight different bacterial strains prevent the growth of the *Pseudoxylaria*

38 different bacterial cultures were obtained from the termites and the comb. These were then tested for their ability to prevent the growth of *Pseudoxylaria*. Four different kinds of interactions were found: clear zone of inhibition, reduced growth of fungus near the bacteria, contact inhibition, and negligible inhibition [13]. As fig. 1 (left panel) indicates, eight different bacteria showed significant capability to prevent the growth of *Pseudoxylaria*.

**Fig. 1.**
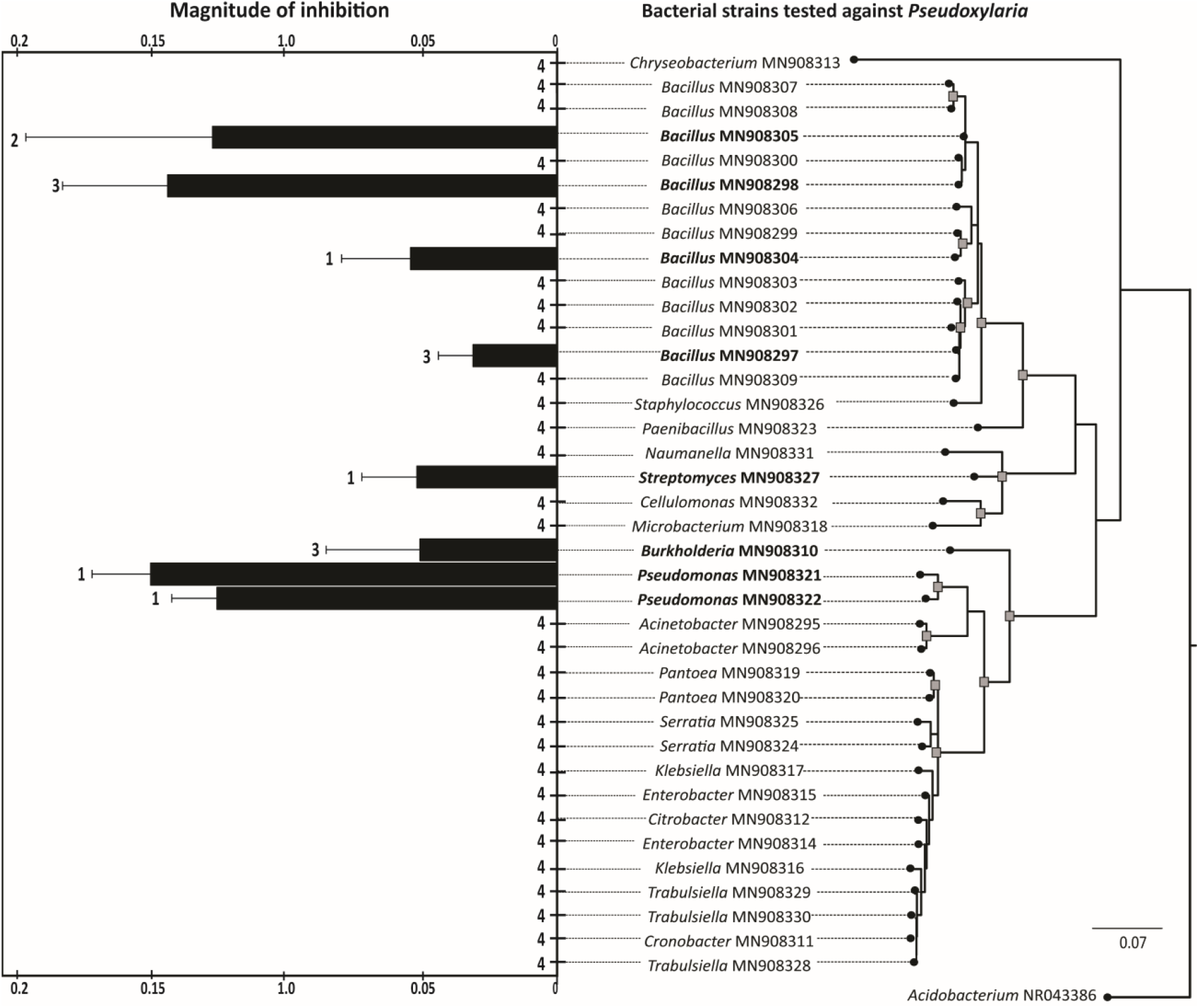
Magnitude of inhibition (left panel) against the parasitic fungus, *Pseudoxylaria*, of the 38 bacterial cultures (right panel) obtained from *O. obesus* colonies. Whisker over horizontal bars represent standard deviation and number indicates the type of inhibition shown by the bacterial strains (1= clear zone of inhibition, 2= reduced growth near bacteria, 3= contact inhibition, 4= negligible inhibition). The phylogenetic analysis was run on MEGAX with the K2+g substitution model using *Acidobacterium* as the outgroup. 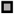 represents > 75 bootstrap value. Bacterial names in bold indicate significant inhibition of *Pseudoxylaria*. Numbers following the genera names indicate NCBI accession number.

A clear zone of inhibition was seen in the interaction between *Pseudoxylaria* and both the strains of *Pseudomonas*, as well as *Bacillus*-MN908304 and *Streptomyces*-MN908327 (Fig. 2, Fig. S6). Whereas, against *Bacillus*-MN908305, *Pseudoxylaria* showed reduced growth near the bacteria (Fig. 2 and Fig. S6). Against three other bacteria (*Bacillus*-MN908297, *Bacillus*-MN908298 and *Burkholderia*-MN908310) *Pseudoxylaria* showed contact inhibition (Fig. 2, Fig. S6). For all the rest of the 30 bacterial cultures, no discernible reduction of fungus growth was observed as it grew over the bacterial colonies (Fig. S7).

**Fig. 2.**
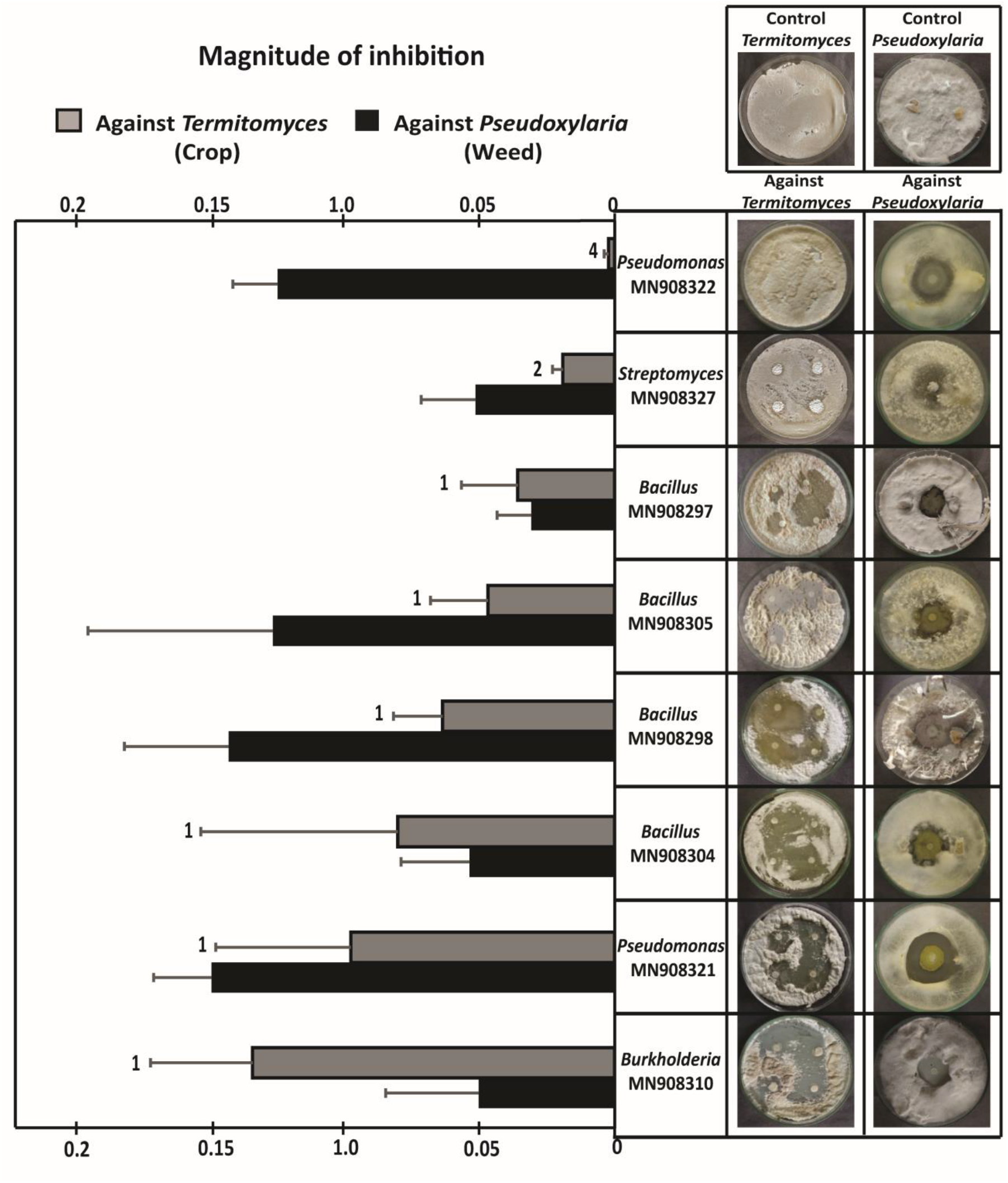
A comparison of the magnitude of inhibition against the crop fungus (*Termitomyces*) and the parasitic fungus (*Pseudoxylaria*) for the eight selected bacterial strains. The data for *Pseudoxylaria* is same as in fig. 1 and is represented here for comparison. The right panel shows representative pictures of *Termitomyces* and *Pseudoxylaria* interactions with the candidate bacterial cultures with the top panel showing growths in control plates. The bacterial strains are arranged on the basis of the magnitude of inhibition against the crop fungus with the top most strain showing the least inhibition. Numbers on *Termitomyces* bars represent the type of inhibition shown by the bacterial strains (1= clear zone of inhibition, 2= reduced growth near bacteria and 4= negligible inhibition).

Thus, the eight strains of bacteria were found to be capable of preventing the growth of the parasitic fungus in culture. As fig. 1 (right panel) indicates there is hardly any phylogenetic similarity across these bacterial strains with this ability, indicating, that this capability might have evolved independently or different modes of prevention of fungal growth are employed by these bacteria.

### Five bacterial cultures show greater inhibition of the parasitic fungus than the crop fungus

These eight strains were then tested against the crop fungus, to test whether these show generic anti-fungal properties and also impedes the growth *Termitomyces* (Fig. S8). As fig. 2 indicates these bacteria show variable degrees of inhibition of *Termitomyces*. However, when compared with the magnitude of inhibition against both the crop and parasitic fungi, it was clear that *Bacillus*-MN908297, *Bacillus*-MN908304 and *Burkholderia*-MN908310 behaved as a generalist fungal growth inhibitor (Fig. 2) and affected the crop fungus to a greater degree than the parasitic fungus. The remaining five bacterial strains showed variable levels of inhibition against the crop fungus but the magnitude of the inhibition was lower as compared to *Pseudoxylari*a. Of particular interest were the two *Pseudomonas* strains obtained as *Pseudomonas*-MN908321 showed the highest magnitude of inhibition against *Pseudoxylaria* but also showed inhibition, albeit to a lesser degree, against *Termitomyces*. But *Pseudomonas*-MN908322 showed high levels of inhibition of *Pseudoxylaria* as well as almost negligible levels of inhibition against *Termitomyces*. Thus, *Pseudomonas*-MN908322 emerged as an excellent candidate for the role of a mutualist. The inhibition by bacterial strains was not due to lack of nutrition and this was confirmed by re-growing *Pseudoxylaria* and *Termitomyces* on used media (supplementary methods S1).

Thus, these two rounds of selection proved that the microbial communities of *O. obesus* can harbor multiple bacteria which have the potential to be a defensive mutualist and *Pseudomonas*-MN908322 emerged as the best candidate.

### *Pseudomonas* is a part of the core microbiota of *O. obesus*

The pan-microbiome of *O. obesus* was established from two types of workers (major and minor), nymphal castes, alates (male and female) and the comb. Over 2.6 × 10^5^ high-quality Nanopore reads were obtained from the six different samples (Table 1) with the highest number of reads from male alates (∼1.1 × 10^5^ reads) and the lowest from minor workers (∼0.03 ×10^5^ reads). The rarefaction curves (Fig. 3) showed exhaustive sampling of three different samples (comb, major worker and male alate) indicating sufficient coverage. However, the number of reads obtained for minor workers and nymph samples were low (Fig. 3). The number of bacterial OTUs identified ranged between 975 (in comb) to 212 (minor worker). The entire pan-microbiome of *O. obesus* had 1045 unique genera (Fig. 4, Table S2).

**Table 1.**
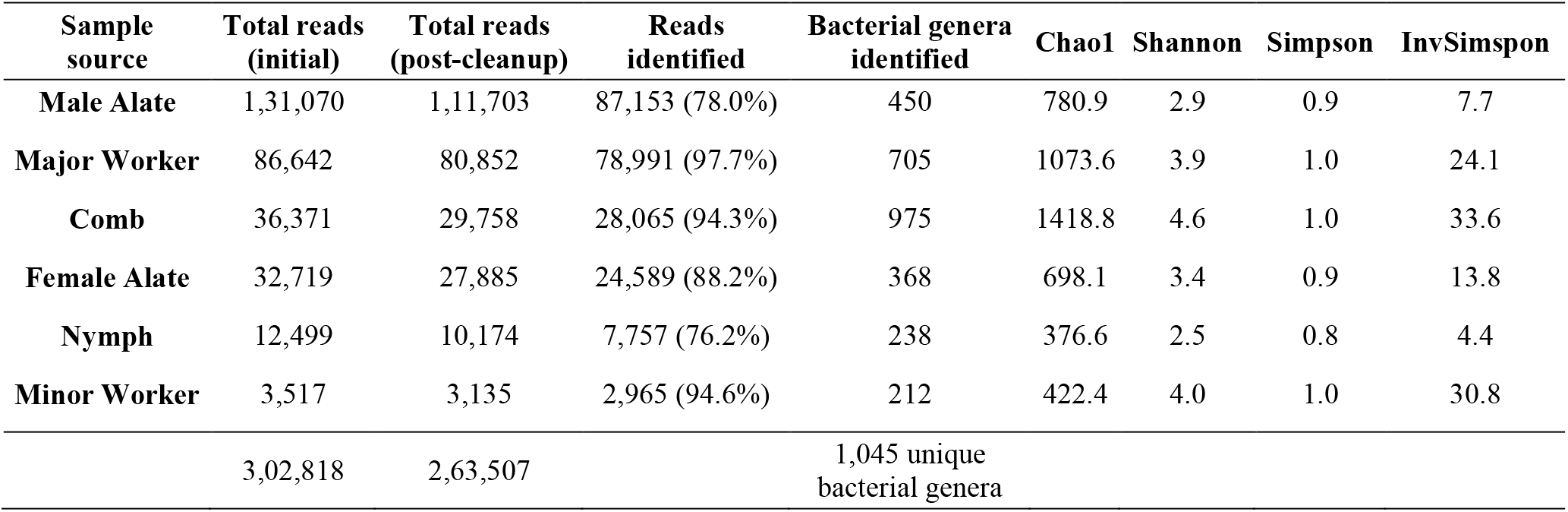
Number of reads obtained to identify the pan-microbiome of *O. obesus*. The table also shows different diversity indices for all the samples. The details of the taxonomic distribution, as well as abundance, for all the identified reads are given in supplementary material, table S2.

**Fig. 3.**
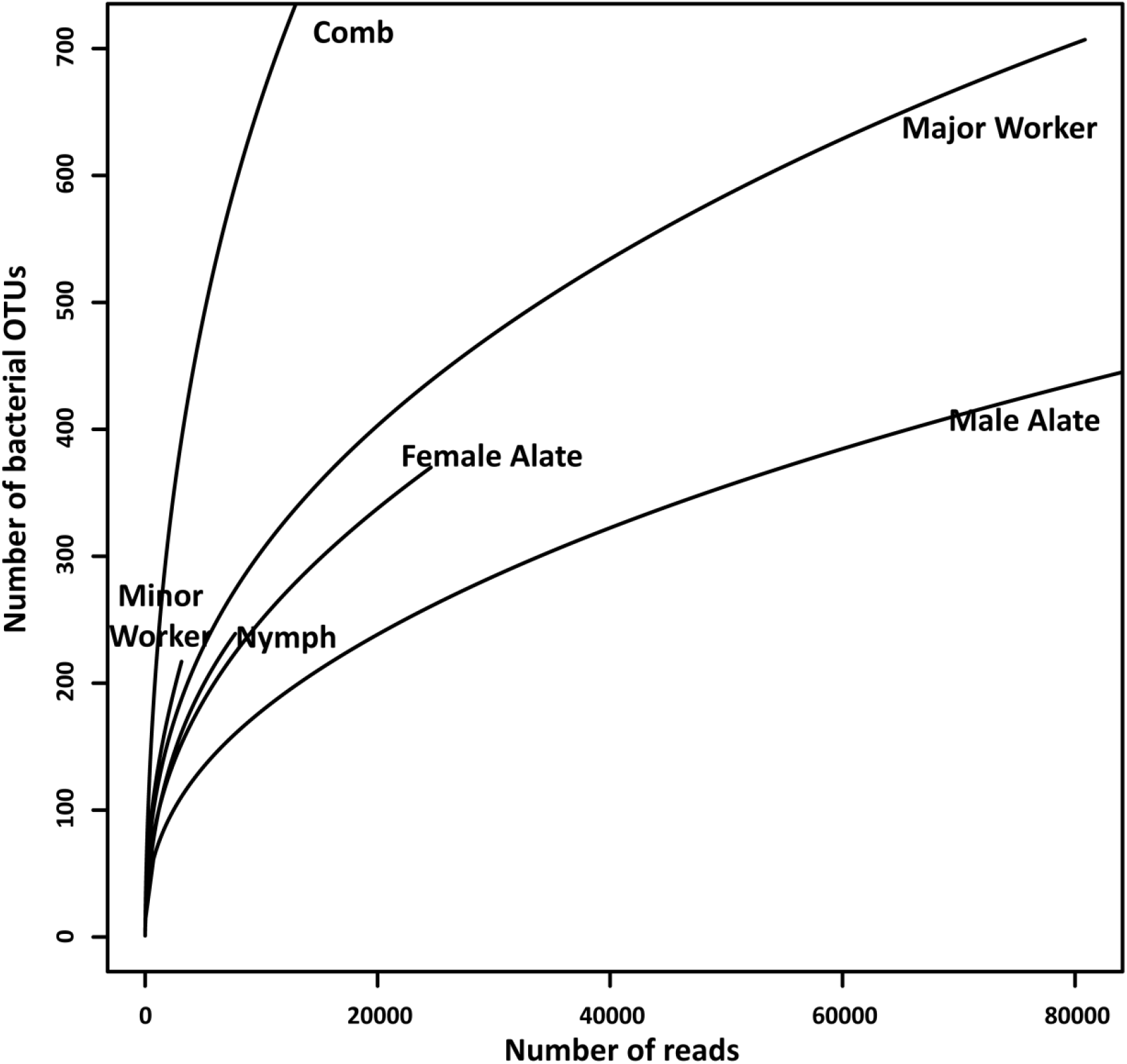
Rarefaction analysis of bacterial 16S rRNA OTUs from six different sources from the colony of *O. obesus*.

**Fig 4.**
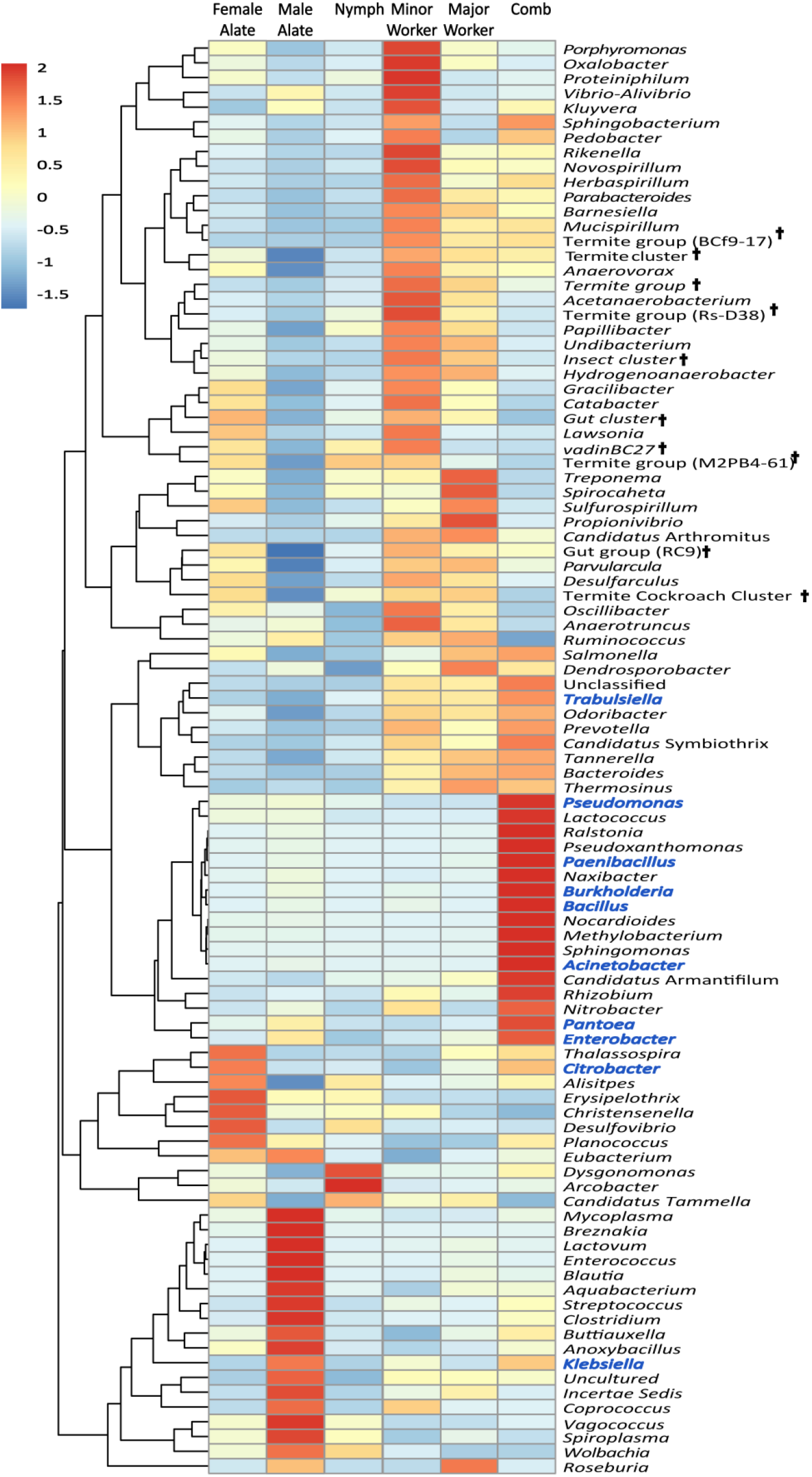
Core-microbiota of *O. obesus*. Bacterial strains that were present in at least five samples of this study are shown in the heatmap. The genera mentioned in blue were also obtained in culture. **†** are the genera classified according to DictDB database.

To identify the core microbiota, the list of all the identified bacterial genera were parsed to find how many of them occur in at least five out of the six different samples. We used this criterion as little is known about caste-specific microbial variation in fungus-growing termites, especially, whether difference in caste roles are also reflected in their microbial composition. As fig. 4 indicates the core microbiota consists of 95 different bacterial genera. *Pseudomonas* was found across five of the six samples and was absent only from minor workers. As minor workers and nymph samples also had the least number of reads (0.03×10^5^ and 0.1×10^5^ reads, respectively), the absence of *Pseudomonas* in the former can probably be explained by the low depth of sequence coverage (Fig. 3).

## Discussion

The major goal of this study was to find whether secondary symbionts with mutualistic capabilities are present within the bacterial community of *O. obesus* which can help in preventing parasitic fungal invasion. We hypothesized that as the crop fungus was the only source of nutrition for these termites, any mutualist present in the community would therefore be selected to be present in any successful colonies of *O. obesus*. However, the critical requisite for any such putative bacterial mutualist must be their dual, but contrasting, capability of preventing the parasitic fungus but not the crop fungus. The 38 different bacterial cultures obtained were therefore subjected to these two rounds of selection against the two fungi. *Pseudomonas*-MN908322 emerged as the best candidate as it strongly inhibits *Pseudoxylaria* but not *Termitomyces*. We further show that *Pseudomonas* is also widely present in the caste members of *O. obesus* by constructing the core microbiota indicating that these are indeed key microbes in this symbiosis.

This is the first study reporting *Pseudomonas* as a potential mutualist in a fungus-growing termite. This is surprising, since, *Pseudomonas* is widely known to prevent many different kinds of fungi and is even used as a commercial antifungal agent [42]. Moreover, they are often one of the dominant microbial genera within fungus-growing insects [19]. This study has uncovered two strains of *Pseudomonas* which have strong inhibitory effects on *Pseudoxylaria*. However, *Pseudomonas*-MN908321 also shows some inhibition of *Termitomyces*, but still less than against *Pseudoxylaria*. But *Pseudomonas*-MN908322 shows negligible levels of inhibition of *Termitomyces* and remains the best candidate for a mutualist. As the phylogenetic analysis shows (Fig. 5), *Pseudomonas*-MN908321 is a close relative of *P. aeruginosa* which has known constitutive antifungal activities [43]. However, *Pseudomonas*-MN908322 clusters with *P. plecoglossicida, P. fluorescens* and *P. putida*. Curiously, *P. putida* can act as a positive enhancer of fungal growth, where it can help augment the mycelial growth of many different fungi within Basidiomycota [43]. It is significant to note that the crop fungus, *Termitomyces*, also belongs to the phylum Basidiomycota.

**Fig 5.**
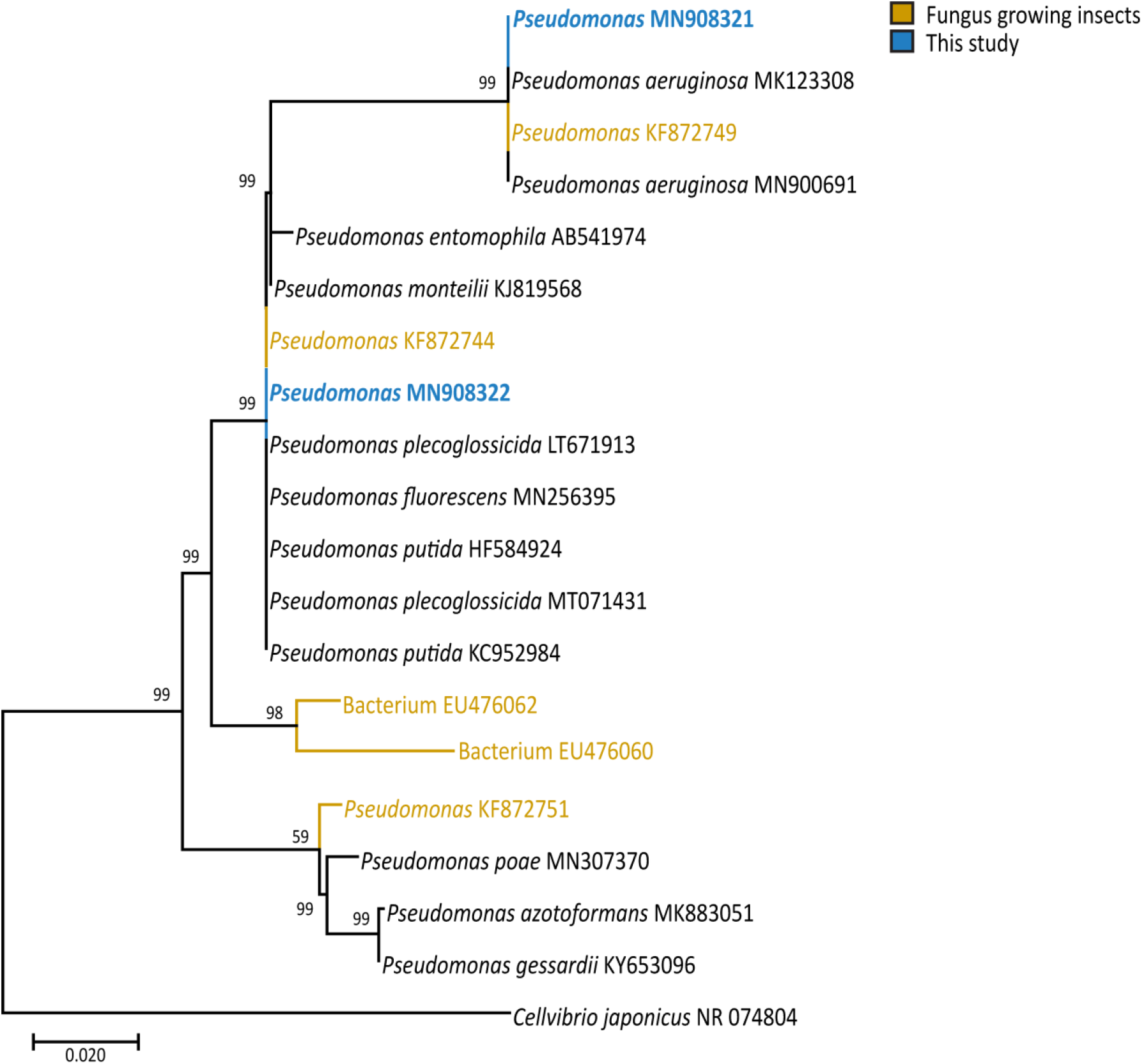
A maximum likelihood phylogenetic tree of *Pseudomonas* obtained in this study (blue) with other related strains. Taxa in yellow are found in fungus-growing insects. The phylogenetic tree was constructed using partial 16S rRNA gene sequences in MEGAX with 1000 bootstrap replicates using K2+g model. The sequences from other fungus growing insects were obtained from Cardoza et al. (2009) [48], Muwawa et al. (2016) [49]. *Cellvibrio japonicus* was used as the outgroup.

Our study also confirms the importance of *Bacillus* and *Streptomyces* as potential defensive mutualists in fungus-growing termites. The role of *Bacillus* is well documented in fungus-growing termites [11, 23] where some bacillaene-producing strains can act as a selective inhibitor of *Pseudoxylaria* [23]. Four strains of *Bacillus* (MN908297, MN908298, MN908304 and MN908305) obtained in this study show substantial growth limiting capabilities against *Pseudoxylaria* but what is also significant is the other nine different *Bacillus* strains do not. Thus, even within a specific genus of bacteria this capability to prevent the parasitic fungus shows phenotypic variability. This finding agrees with previous studies [11, 23] where different *Bacillus* strains showed a wide variance in their ability to prevent the parasitic fungus.

*Streptomyces* can also provide chemical defense by producing specific inducible secondary metabolites in fungus-growing insects [44, 45]. A wide range of chemicals produced by *Streptomyces* can not only act as antifungal agents but can also show antibacterial properties in some insects [46].

The genus *Odontotermes* is one of the most speciose and wide-spread of all fungus-growing termites with around 390 species spread across Africa and Tropical Asia [47]. Therefore, identifying the roles of the different secondary symbionts within *Odontotermes* colonies can be a key to understand the evolutionary origins of this symbiosis. This study presents an extensive analysis of the different bacterial genera found within O. *obesus* colonies. To test how representative these communities are across different species of *Odontotermes*, a comparison was done with the data obtained in this study with previously published data from nineteen different species of *Odontotermes* [31, 40, 41]. The microbial abundance datasets obtained from this study were combined from all the six different samples and a consolidated list of bacterial genera was made for comparison. As fig. 6 shows, *O. obesus* microbial composition differs substantially from other *Odontotermes* species. A weighted PCoA analysis of the Unifrac distances among these samples also shows a limited clustering (Fig. S10). However, the presence of *Pseudomonas* could be detected in fourteen of the nineteen samples (Fig. 6). This presence of *Pseudomonas* in *Odontotermes* colonies, across their range, further underscores their importance to this symbiosis.

**Fig 6.**
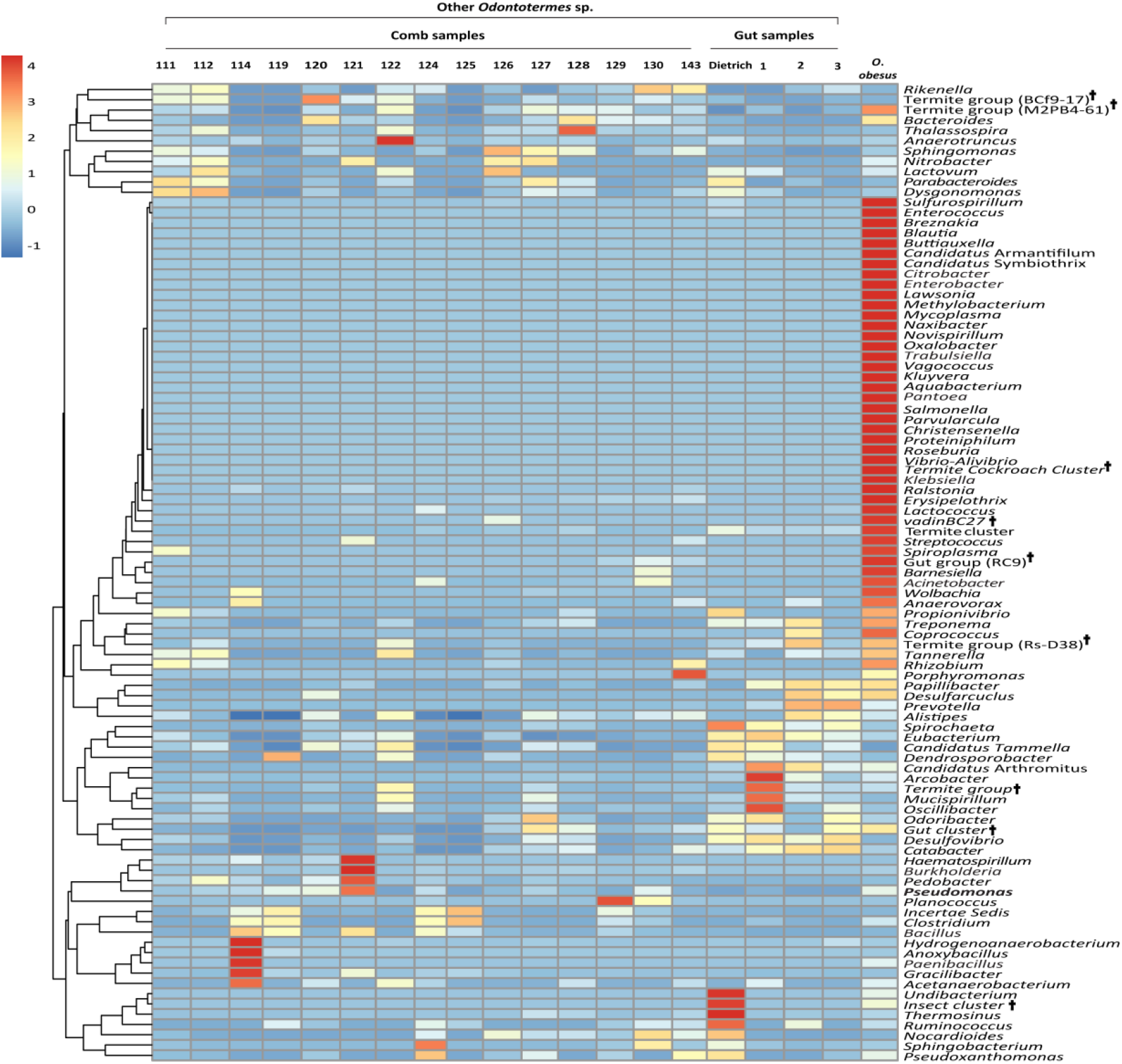
Comparison of *O. obesus* core-microbiota with the microbiota of 19 other *Odontotermes* species to evaluate the presence of *Pseudomonas. Pseudomonas* was found in 15 samples. The data from 19 different *Odontotermes* sp. were compiled from Dietrich et al., 2014 [40]; Otani et al., 2014 [31] and Otani et al., 2016 [41] (numbering of the *Odontotermes* sp. reflects their original nomenclature from the studies cited). **†** are the genera classified according to DictDB database.

The results from this study also raise several important questions about microbial community ecology and termite biology. First, how many different mutualistic bacterial strains are there in *O. obesus* colonies and what are their specific roles? We obtained 38 different cultures but high-throughput methods employed (Nanopore platform) identified at least 1045 different bacterial genera. Thus, the existence of other bacterial mutualists cannot be ruled out. Another confounding factor is the difficulty in identifying the number of strains present from any one particular bacterial genus. This question is important as our results indicate that there are at least two different cultures of *Pseudomonas*, showing varying degrees of mutualistic capabilities. Until and unless unambiguous enumeration of the number of strains is done, it is difficult to identify specific functions with specific secondary symbionts.

The other major question that emerges from this study is, what roles, if any, are the termites playing in preventing the parasitic fungus. Specifically, are termites using any of these identified mutualists in preventing *Pseudoxylaria*, or are the mutualists doing the prevention independent of the termites? The answer must await future experiments which include both termites as well as the identified candidate mutualists.

## Acknowledgements

We thank Dr. Nivedita Saha, Zoological Survey of India, for identifying different termite species and Nimisha E.S. for help with the fieldwork. We also thank Paras Verma for assistance with the bioinformatics analysis. We acknowledge all EVOGEN lab members for their valuable comments on manuscript.

## Declarations

### Availability of data and material

The microbial dataset generated is available as supplementary material to this article. 16S rRNA fragments obtained have been submitted to GenBank (accession number MN908295–MN908332) and Nanopore sequencing generated have been submitted to NCBI Sequence Read Archive (SRA) under the BioProject PRJNA608773 (BioSample accessions: SAMN14208512-SAMN14208517).

### Code availability

Not applicable

### Authors’ Contributions

RA and RR conceived and planned the study. RA performed field collections, sequencing, microbial culturing, data collection and analysis. AA helped in fieldwork and pilot experiments. Nanopore Sequencing and its analysis were done by RA and MG. The statistical analysis of interaction experiments was done by RS and RA. RR and RA wrote the paper.

