## Supplemental File for "*Pseudomonas* can prevent the parasitic fungus, while keeping the crop fungus unaffected, in the gardens of *Odontotermes obesus*"

**Supplementary methods S1**

**DNA extraction of termites and fungus comb**

Individuals were put in 1 ml of CTAB buffer containing 100 mM Tris (pH 8), 2% CTAB, 1.4 M NaCl, 20 mM EDTA, 1% β-Mercaptoethanol and 0.3 g $mL^{-1}$ proteinase K. These were crushed and then incubated at 60˚C for 30 minutes. These were then subjected to Phenol: Chloroform: Isoamyl alcohol (PCI: 25:24:1) precipitation and the purified DNA pellet obtained was dissolved in 1X TE (pH 8) buffer.

DNA extractions from fungus combs often have extensive humic acid contamination which can hinder downstream PCR reactions. To remove this humic acid, the precipitation step with PCI were repeated until the final pellet obtained was white in color [1, 2]. DNA samples were used for PCR only if the 260/280 nm ratio were within the range of 1.3-1.9, which were obtained through a NanoDrop Spectrophotometer 2000 (Thermo-Fisher Scientific).

**PCR amplification, sequencing and phylogenetic analysis of bacterial isolates**

Bacterial colonies were identified by amplifying a portion of the 16S rRNA gene using primer set 341F/806R [3]. The amplification reaction was prepared to a final volume of 20 µl containing: 14.5 µl sterile distilled water, 0.4 µl dNTPs (10 µM), 2 µl 17.5 mM buffer, 0.5 µl of each primer (10 µM), 2 µl template and 0.1 µl *Taq* Polymerase (Himedia). PCR was performed under following conditions: an initial denaturation step at 95°C for 3 minutes, followed by 33 cycles of denaturation (95°C, 45 seconds), annealing (56°C, 45 seconds), extension (72°C, 1 minute) with a final extension at 72°C for 10 minutes. The PCR products were cleaned with Exonuclease I and Shrimp alkaline phosphatase and then sequenced with BigDye® Terminator v.3.1 cycle sequencing kit for both strands. The chromatograms obtained were cleaned with Sequencher v.5.2.4 (GeneCodes Corp.) and manually edited with Bioedit v.7.0.5.3 [4]. Taxonomic identification of these sequences was achieved through BLAST with the parameters of >90% sequence identity and/or the first hit. The sequences were aligned with other similar homologues, obtained from NCBI, using the ClustalW in Bioedit. The Maximum likelihood phylogenetic tree of aligned sequences was constructed using MEGAX v.10.1.7 [5] with 1000 bootstrap replicates. Suitable substitution models for various phylogenetic analyses were also obtained through MEGAX. We further selected the isolated bacteria based on their sequence similarity of the 16S rRNA gene so that all the selected strains had a unique sequence profile.

**Reusing of media from bacterial- fungal interactions**

One confounding factor of taking the absence of fungal growth as a measure of inhibition by the bacteria is fungal growth can stop because the bacteria has exhausted the nutrition across its area of growth. This can result in erroneous estimation of inhibition, especially with contact inhibition, where one microbe cannot grow over the other due to unavailability of further nutrition from the PDA. Although, this seems unlikely, yet, to rule out this possibility the media from the area of bacterial growth was reused for fungal growth. For some interaction plates, the 7 days old media, on which bacteria was growing, was filtered, autoclaved, and re-plated to grow and analyze the growth of both *Pseudoxylaria* and *Termitomyces*. Both these fungi showed equal rates of growth on these reused as well as fresh PDA media (Fig. S9).

**Sample preparation and obtaining sequences from Nanopore Sequencing**

The amplification reaction was prepared to a final volume of 20 µl containing: 14.5 µl sterile distilled water, 0.4 µl dNTPs (10 µM), 2 µl 17.5 mM buffer, 0.5 µl of each primer (10 µM), 2 µl template and 0.1 µl *Taq* Polymerase (Himedia). PCR was performed under following conditions: an initial denaturation step at 95°C for 3 minutes, followed by 33 cycles of denaturation (95°C, 45 seconds), annealing (56°C, 45 seconds), extension (72°C, 1 minute) with a final extension at 72°C for 10 minutes. PCR products were cleaned using Wizard® SV Gel and PCR clean-Up System (Promega). Purified PCR products were quantified using NanoDrop 2000 (Thermo Scientific) and Qubit 3.0 Fluorometer (Life Technologies). To obtain a comprehensive assessment of microbial diversity, three individual DNA extractions from two mounds were pooled in equimolar concentrations. This was done for comb DNA and all the termite castes, except alates, where three individual DNA extractions from males and females were pooled.

An initial library preparation was done using the SQK-LSK 108 Ligation Sequencing Kit 1D-vR9 (Oxford Nanopore Technologies) where poly- 'A' tails were added to the purified PCR product by Ultra II End-prep enzyme mix. The resultant mixture was purified using 1X AMPure XP beads and eluted in nuclease-free water. These were individually ligated with barcode adaptors and purified using 1X AMPure XP beads and eluted in nuclease-free water. These six different samples were then individually barcoded with unique barcodes using the EXP-PBC001 PCR barcoding kit I (https://community.nanoporetech.com/protocols).

Barcoded samples were then purified using Wizard® SV Gel and the PCR clean-Up System (Promega) and quantified using Qubit 3.0 Fluorometer (Life Technologies). All the six barcoded samples were pooled in equal amount (20-25 ng $\mu l^{-1}$) before being loaded on the Nanopore flowcell. 5 µl of λ phage DNA was added to the library to serve as a control.

After the completion of the run, sequences were first separated according to their barcodes by using the program ont-albacore (Oxford Nanopore Technologies Ltd.) These were then converted to FASTA and FASTQ files using *Poretools* (v 0.5.1) for downstream analysis [6]. High-quality reads (average read quality score ≥ 10) were selected using the program *Nanofilt* [7]. The obtained high-quality sequences were then further processed where the barcodes and adaptors were removed using Porechop (https://github.com/rrwick/Porechop). The sequences exhibited the average accuracy of about 89% which was estimated by aligning control λ phage DNA with existing NCBI λ sequences (NC_001416.1) using LAST v. 973.

### Supplementary figures

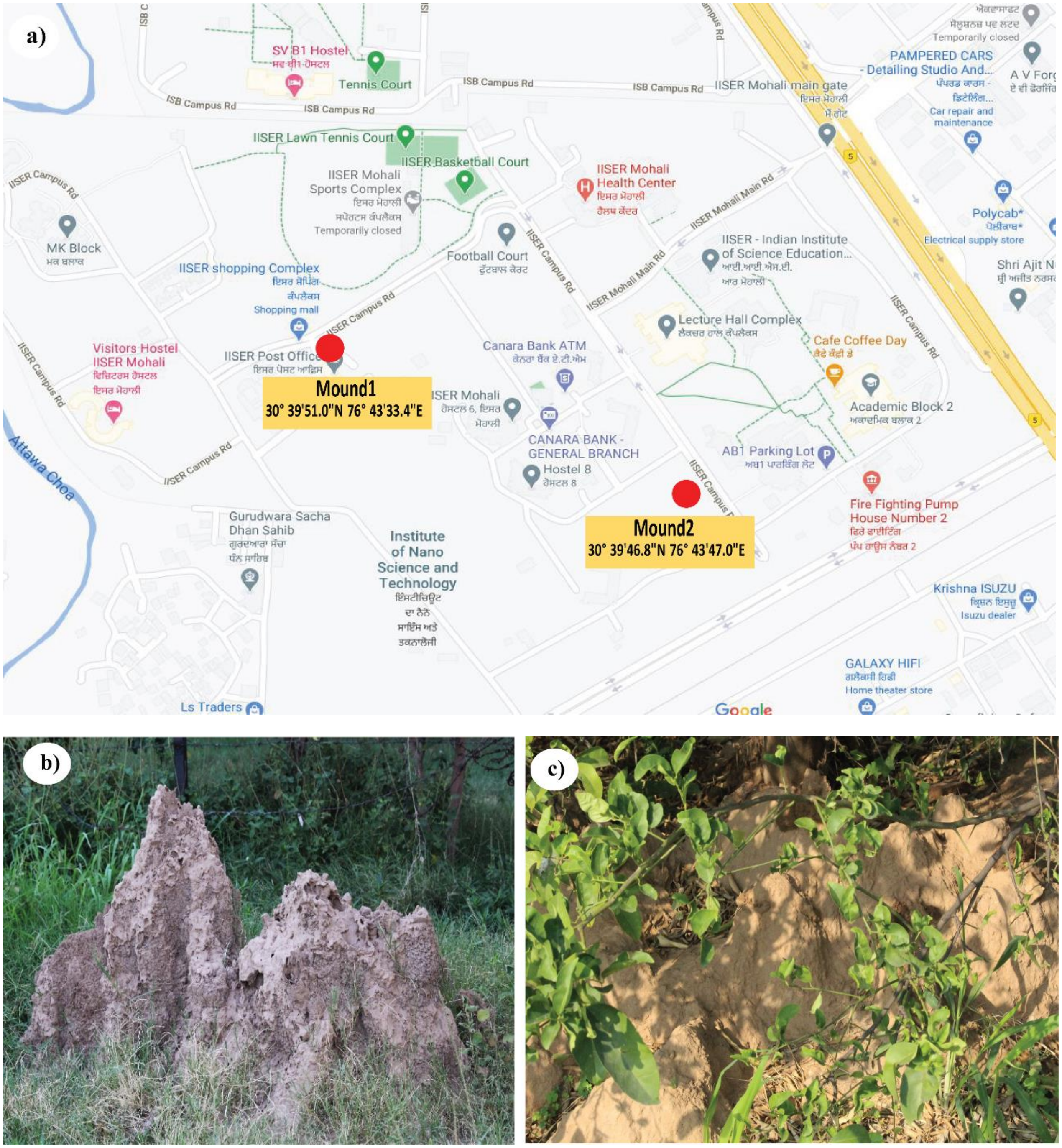

Fig. S1: a) Geographical location of *Odontotermes obesus* colonies used in the study. b) & c) Close-up of the two mounds of *O. obesus* used in the study.

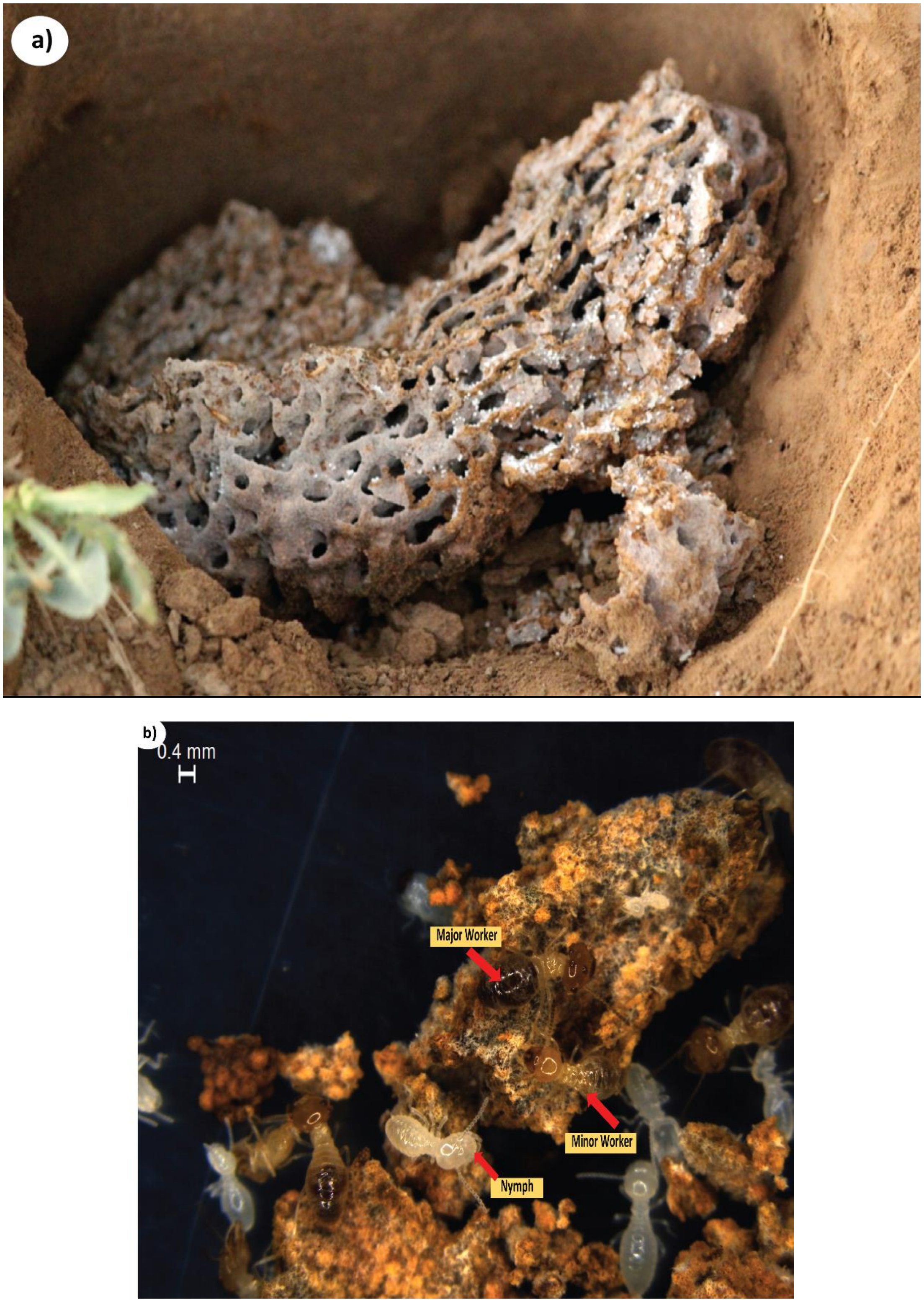

Fig. S2: a) Detailed view of fungus comb b) Different castes of termites on the fungus comb.

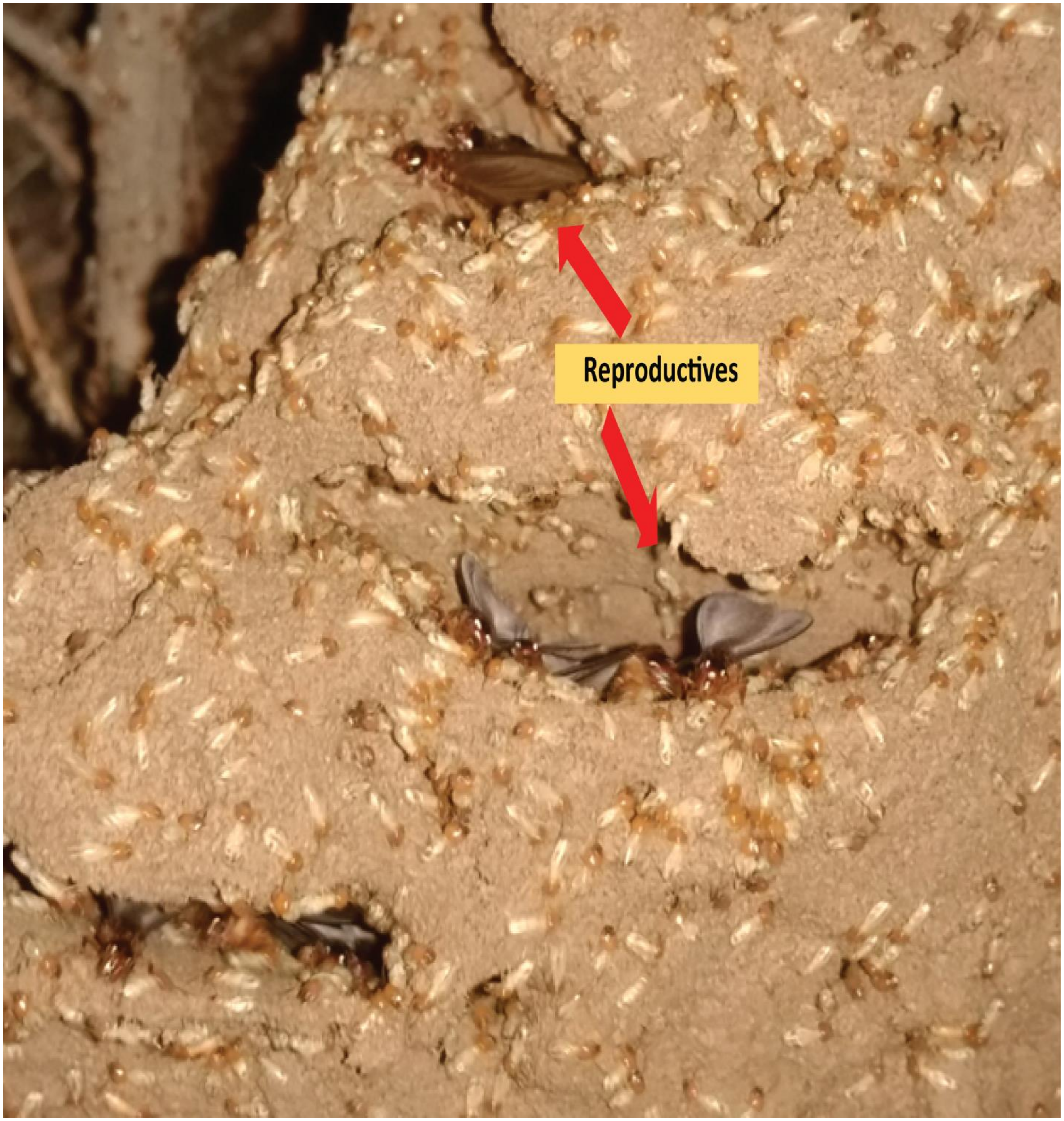

Fig. S3: Reproductives emerging out from the gaps in mound during monsoon season.

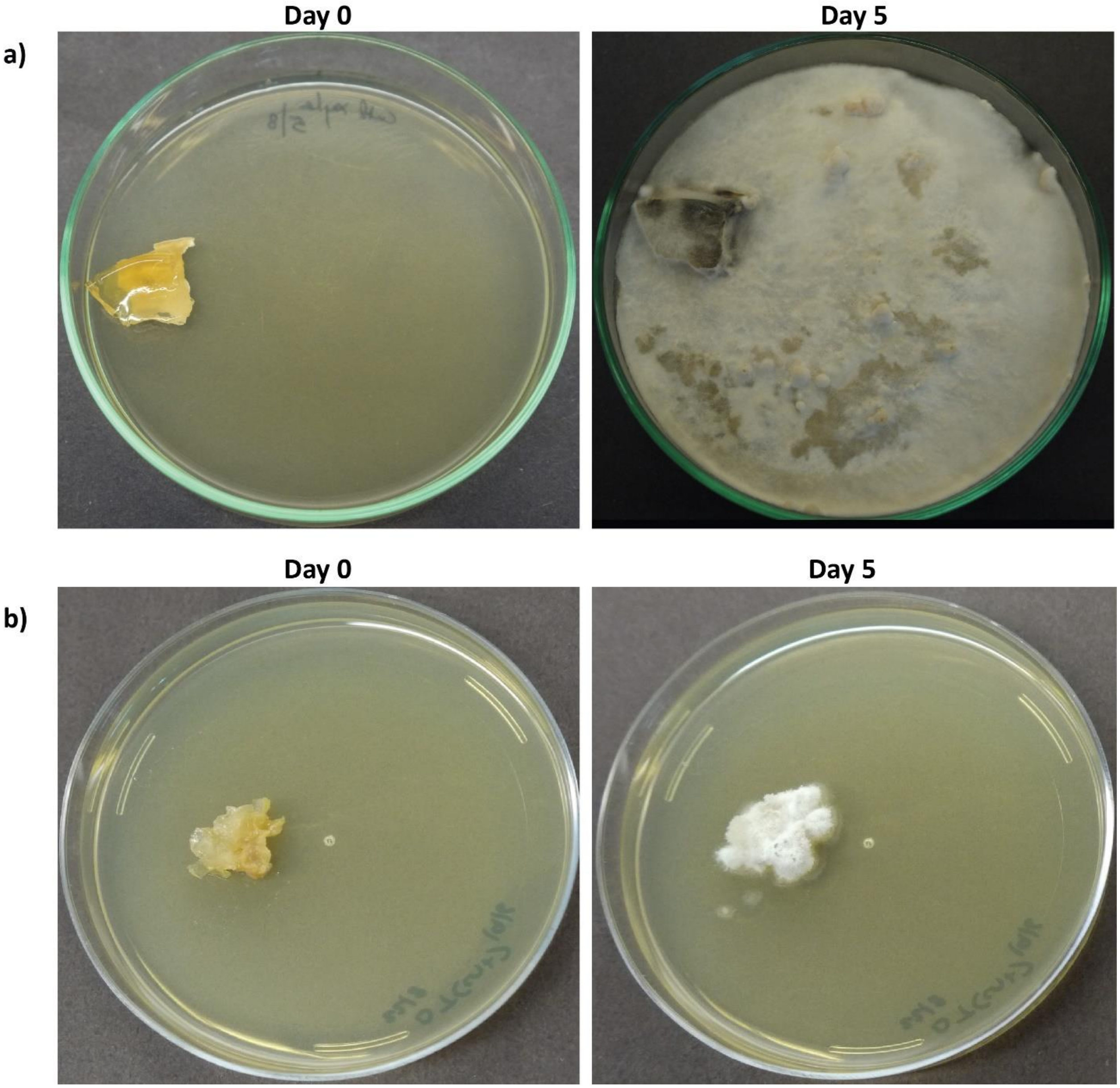

Fig. S4: Differences in the growth rates of a) *Pseudoxylaria* and b) *Termitomyces* are shown by comparing their pictures from different days.

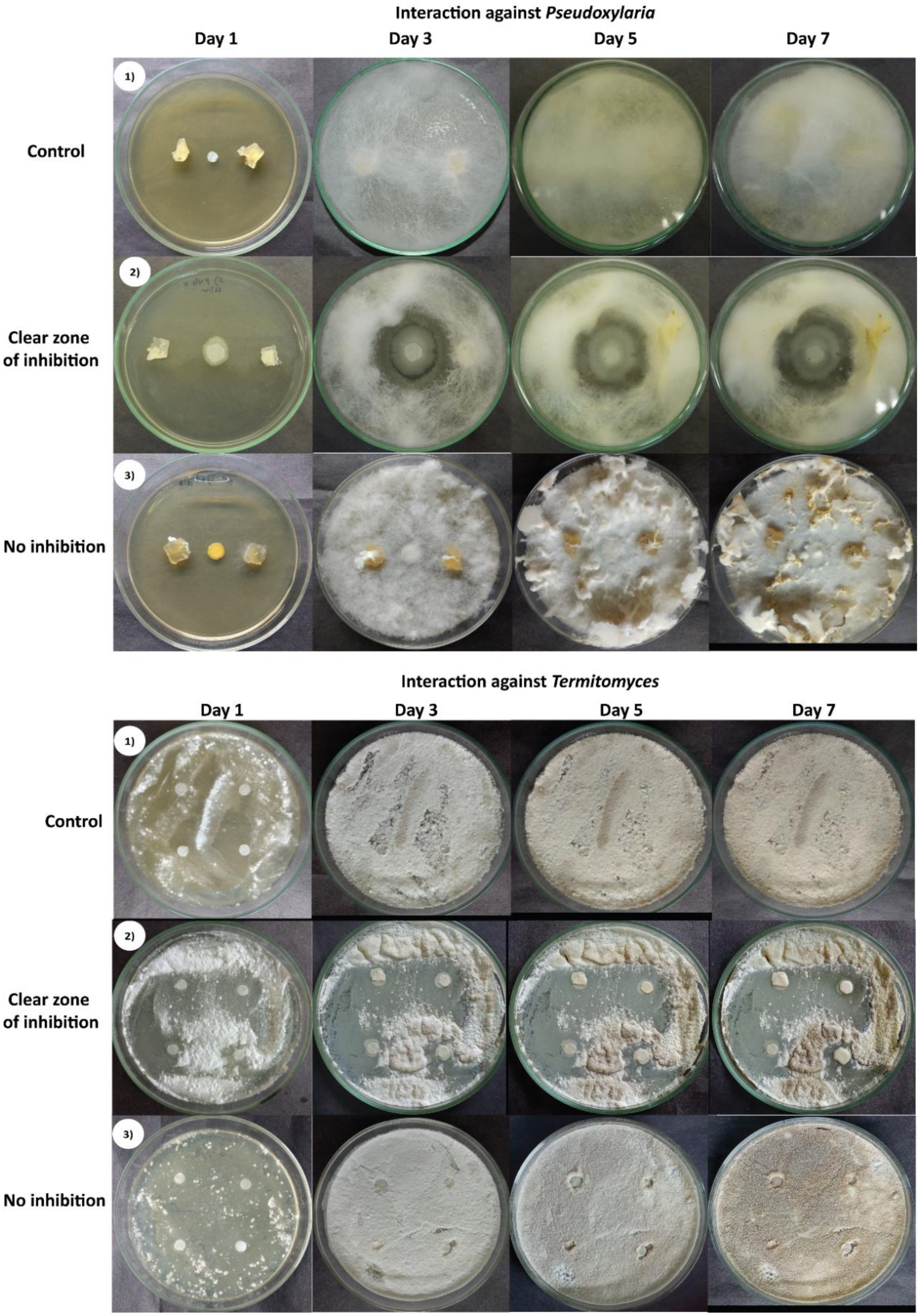

Fig. S5: Representative pictures of bacterial and fungal interaction.

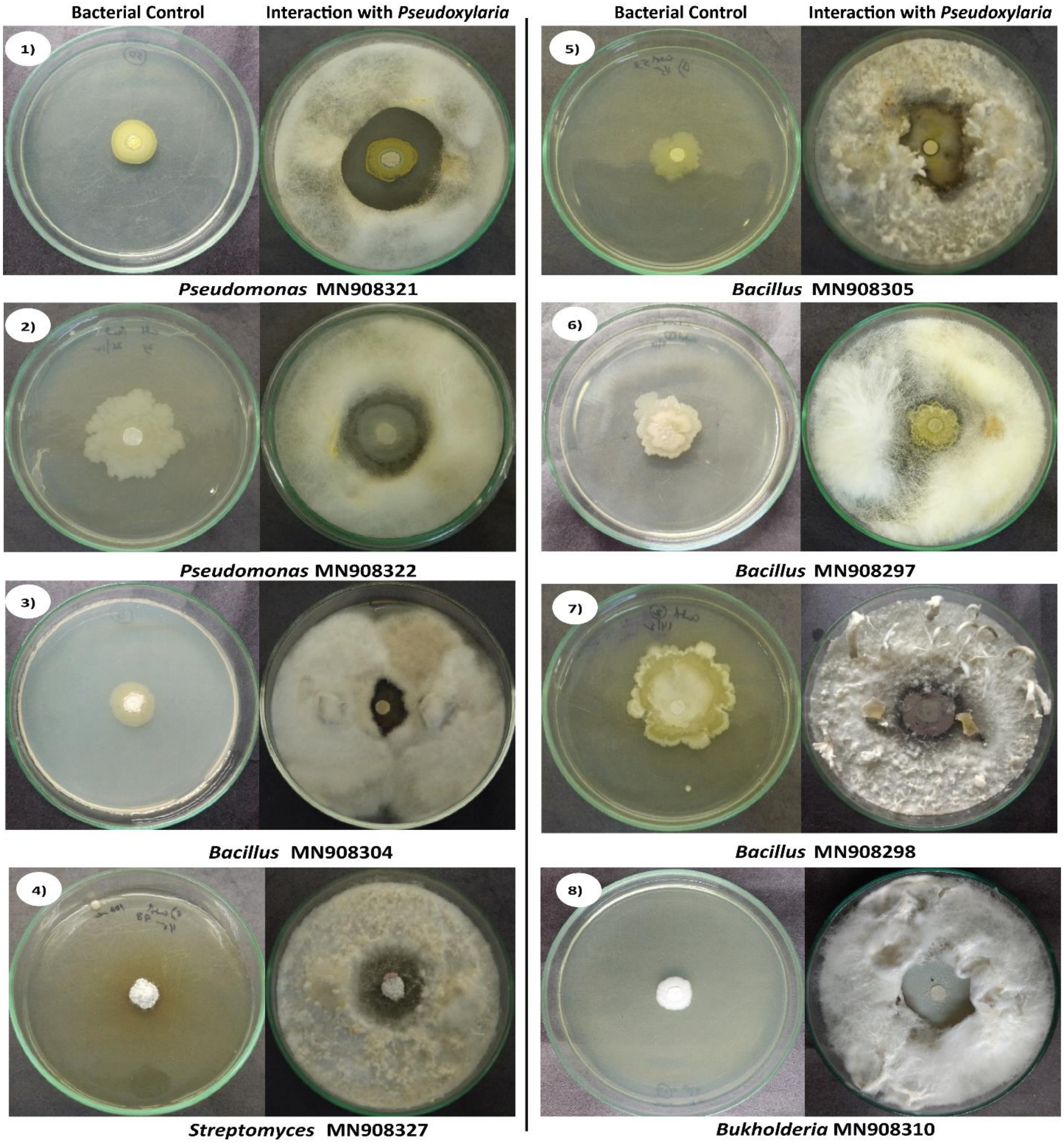

Fig. S6: Assay for the prevention of the growth of *Pseudoxylaria*. All the interactions are categorized into four types: **1)-4) clear zone of inhibition**, **5) reduced growth near bacteria** and **6)-8) contact inhibition.** Each interaction is represented with its bacterial control and one representative test plate. All the photos are from day 7 of the beginning of the experiment.

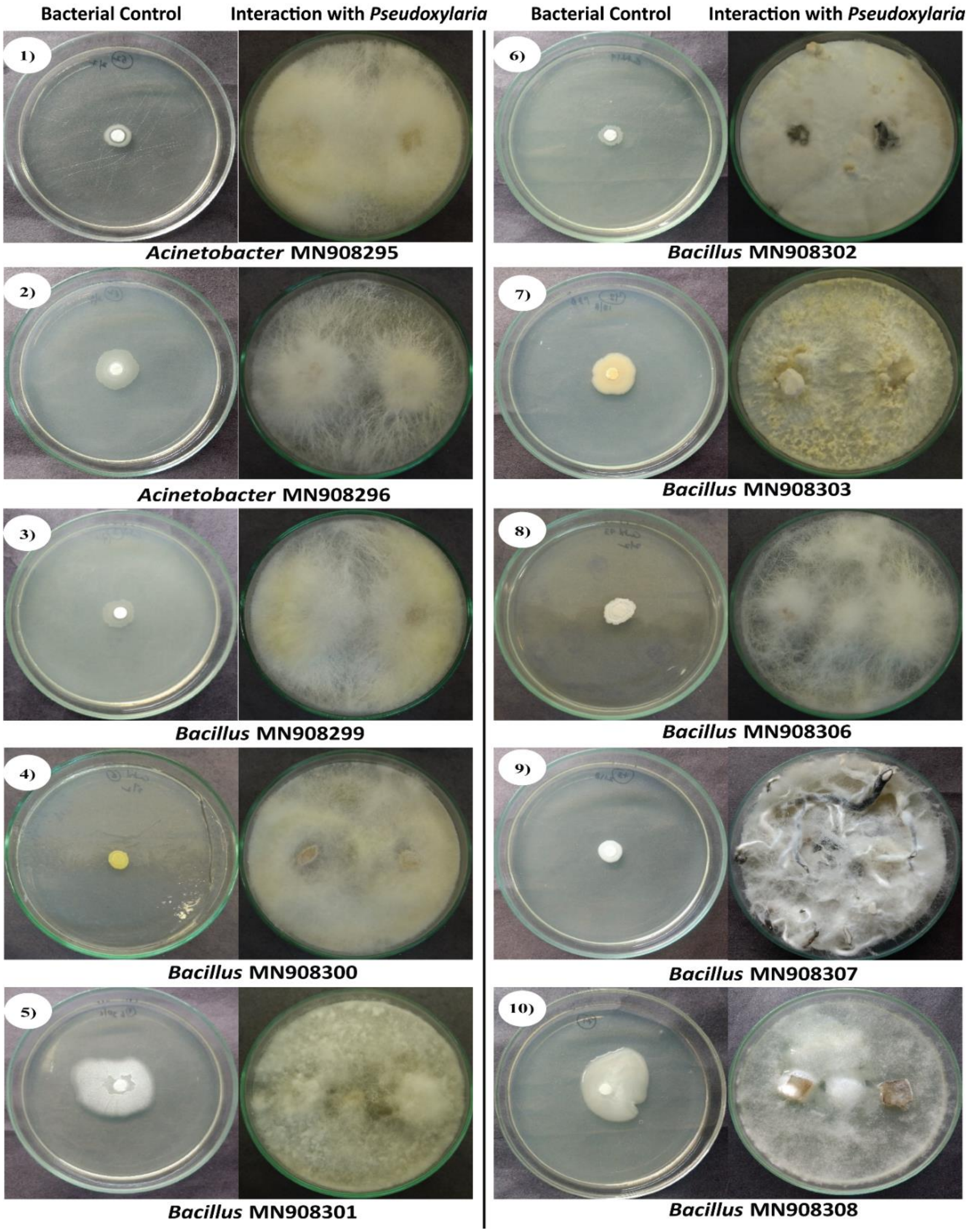

Fig. S7: The interaction assays between *Pseudoxylaria* and the 30 bacterial strains (1-30) which showed **negligible inhibition** of *Pseudoxylaria*. Each interaction is represented with its bacterial control and one representative test plate. All the photos are from day 7 of the beginning the experiment.

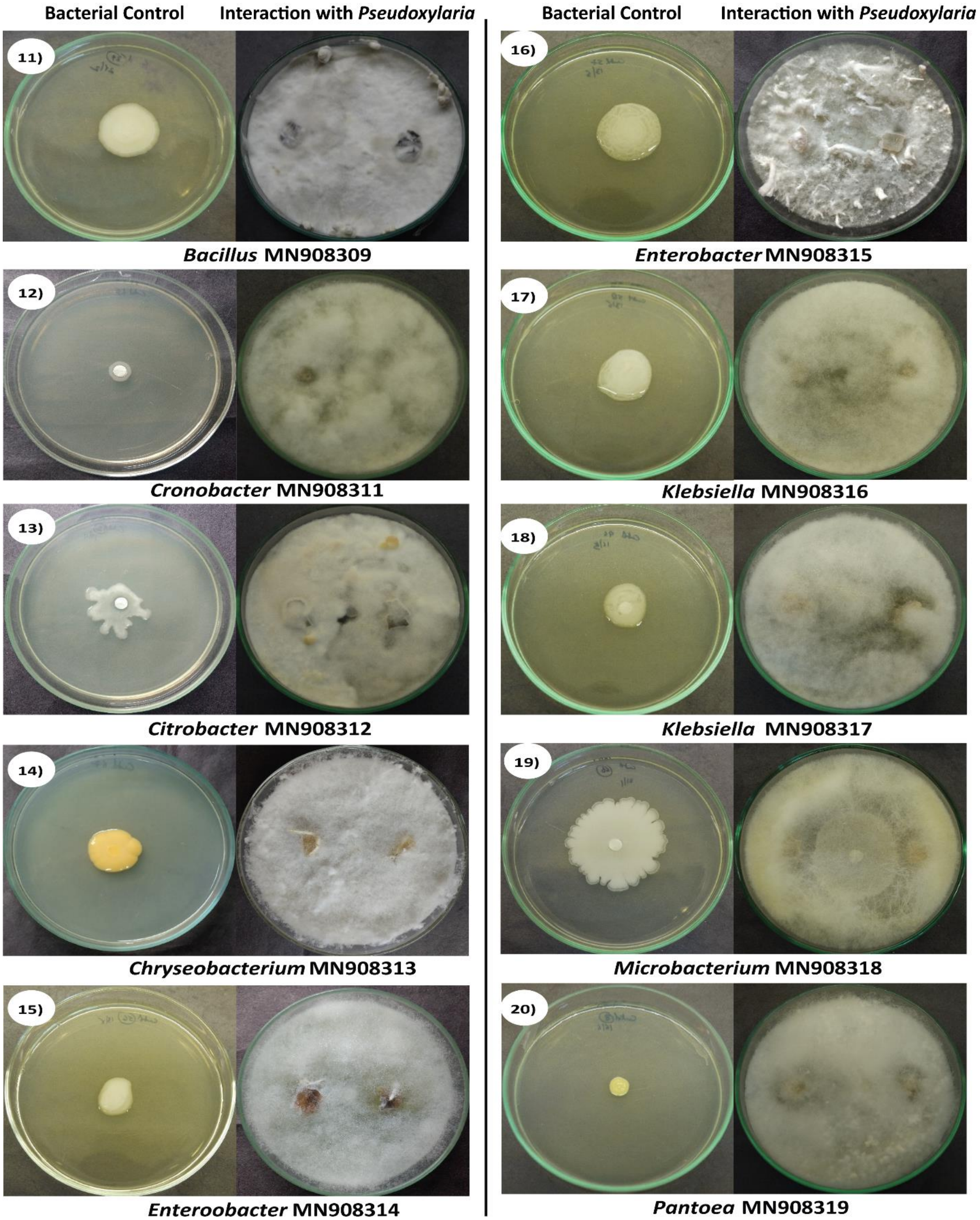

Fig. S7 continued.

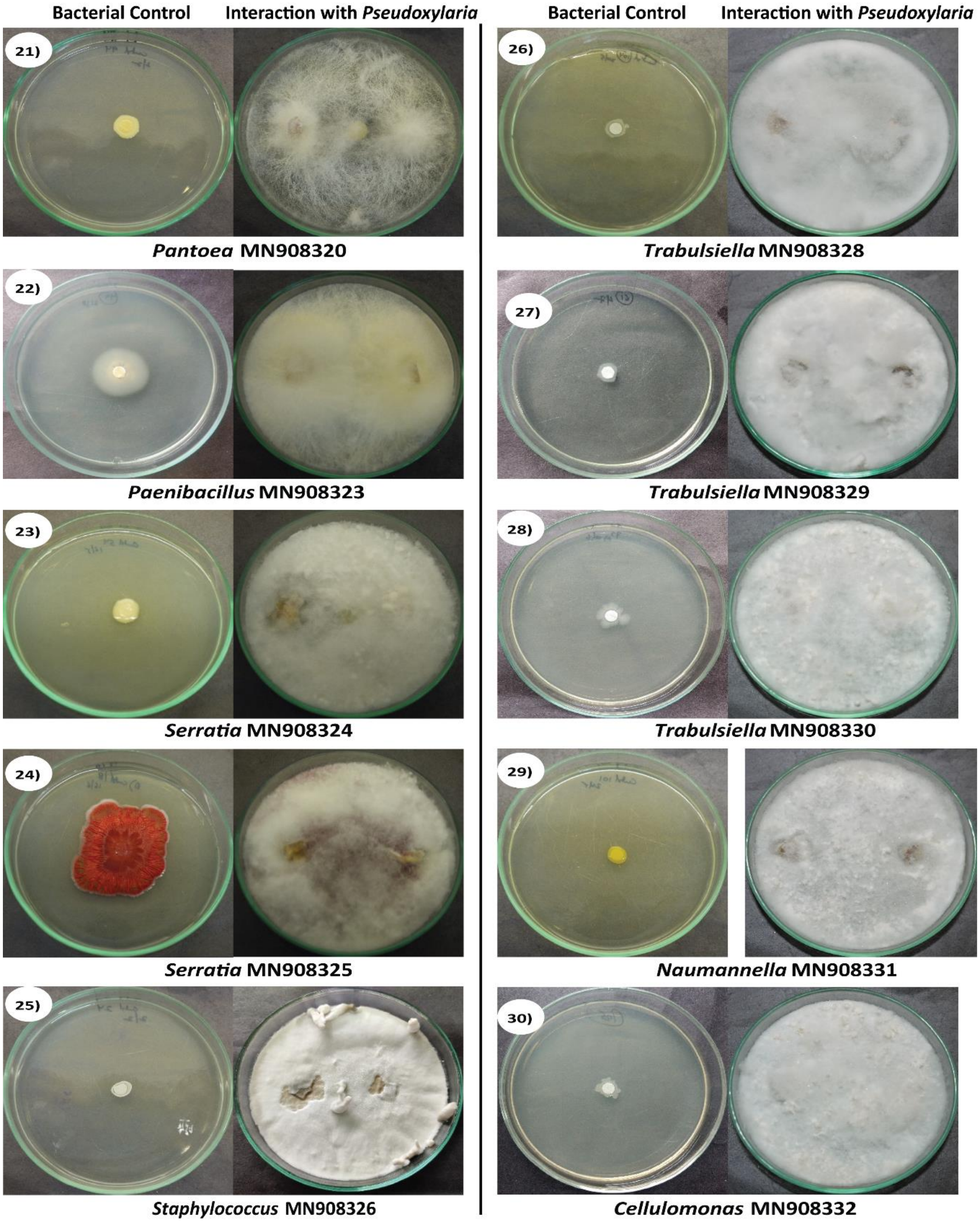

Fig. S7 continued.

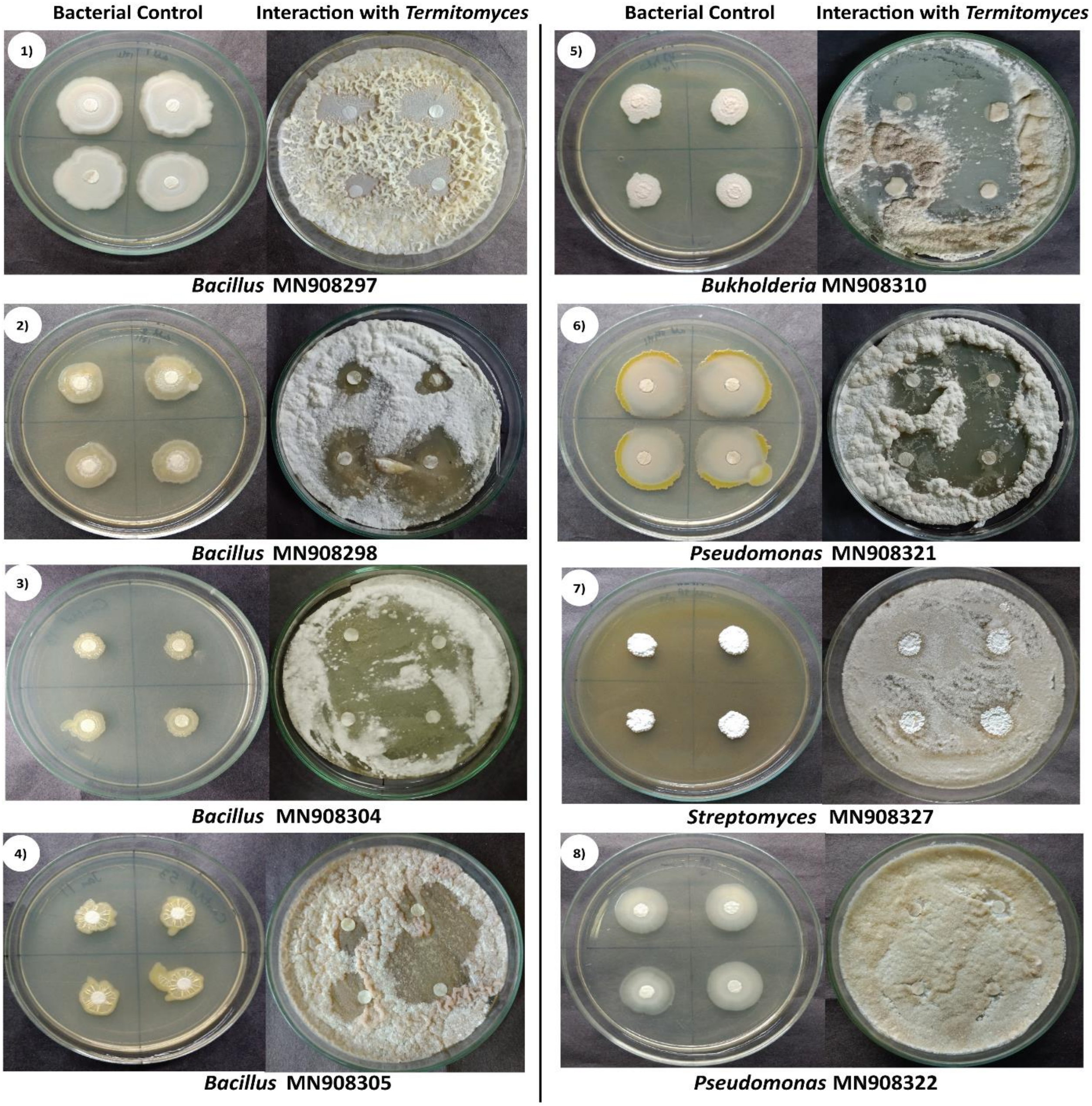

Fig. S8: Interaction assay of selected 8 selected bacterial strains against *Termitomyces*. All the interactions are categorized into four types: 1)-6) **clear zone of inhibition**, 7) **contact inhibition** and 8) **negligible inhibition** of the *Termitomyces*. Each interaction is represented with its bacterial control and a representative test plate. All the photos are from day 7 of the beginning of the experiment.

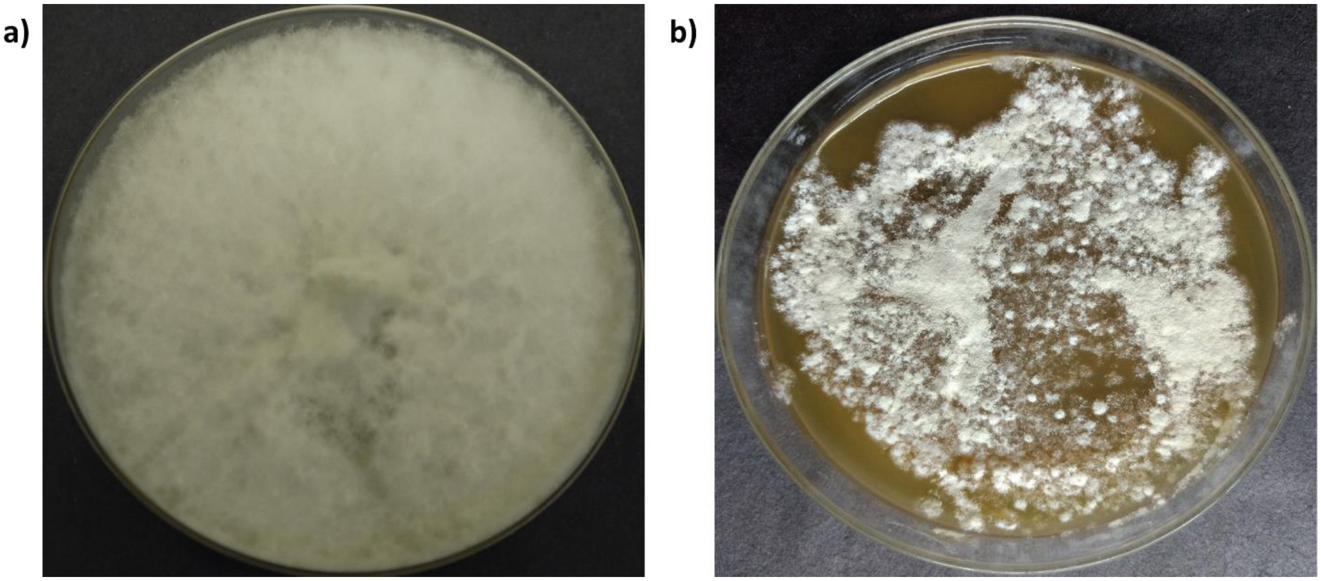

Fig. S9: Day 5 pictures of a) *Pseudoxylaria* and b) *Termitomyces* grown on previously used media from the interaction assay that was autoclaved and re-plated.

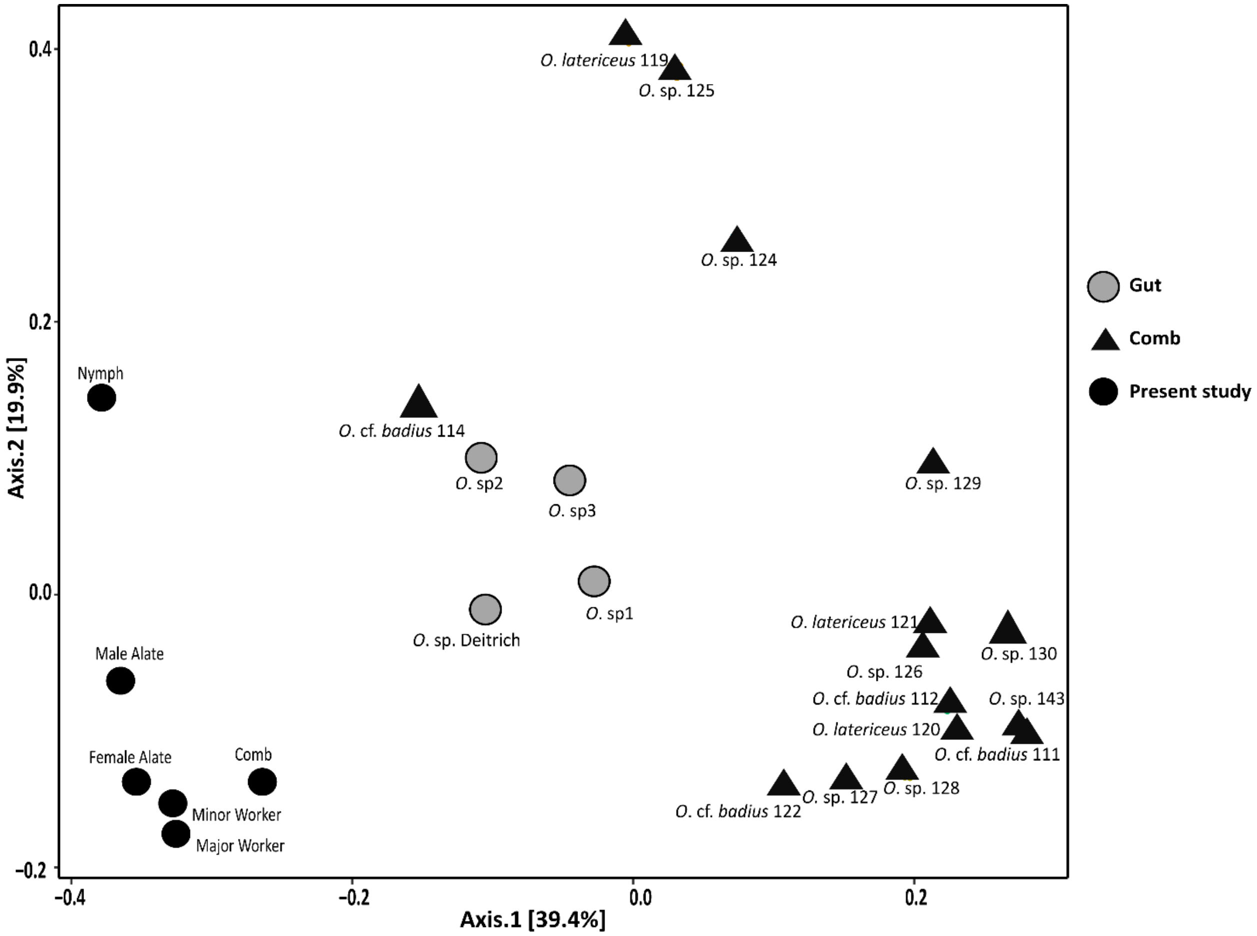

Fig. S10: PCoA similarity analysis of comb and worker gut samples from different *Odontotermes* sp. together with the samples of this study using weighted Unifrac distance.

**Table S1: Details of the strains isolated for the culture-dependent assay.**

| Isolated cultures (Genus-Accession no.) | Nearest neighbor in GenBank | % identity |
|---|---|---|
| *Acinetobacter* sp. **(MN908295)** | MT076253 | 100 |
| *Acinetobacter* sp. **(MN908296)** | CP045428 | 100 |
| *Bacillus* sp. **(MN908297)** | MN866795 | 99.86 |
| *Bacillus* sp. **(MN908298)** | MT081483 | 100 |
| *Bacillus* sp. **(MN908299)** | CP047059 | 100 |
| *Bacillus* sp. **(MN908300)** | MT072181 | 100 |
| *Bacillus* sp. **(MN908301)** | MT091981 | 100 |
| *Bacillus* sp. **(MN908302)** | MT081301 | 100 |
| *Bacillus* sp. **(MN908303)** | MG893109 | 99.91 |
| *Bacillus* sp. **(MN908304)** | MT075838 | 100 |
| *Bacillus* sp. **(MN908305)** | MT089708 | 99.71 |
| *Bacillus* sp. **(MN908306)** | MT086207 | 100 |
| *Bacillus* sp. **(MN908307)** | MN988922 | 99.50 |
| *Bacillus* sp. **(MN908308)** | CP048876 | 99.89 |
| *Bacillus* sp. **(MN908309)** | MT072200 | 100 |
| *Burkholderia* sp. **(MN908310)** | MK048540 | 100 |
| *Cellulomonas* sp. **(MN908332)** | MK106264 | 100 |
| *Chryseobacterium* sp.**(MN908313)** | MK116543 | 99.90 |
| *Citrobacter* sp. **(MN908312)** | MN400096 | 100 |
| *Cronobacter* sp. **(MN908311)** | JN222370 | 99.91 |
| *Enterobacter* sp. **(MN908314)** | MT085978 | 99.62 |
| *Enterobacter* sp. **(MN908315)** | MK318776 | 99.90 |
| *Klebsiella* sp. **(MN908316)** | CP047675 | 99.91 |
| *Klebsiella* sp. **(MN908317)** | LR134333 | 100 |
| *Microbacterium* sp. **(MN908318)** | KR906327 | 99.90 |
| *Naumannella* sp. **(MN908331)** | KP326341 | 98.43 |
| *Pantoea* sp. **(MN908319)** | MF462970 | 100 |
| *Pantoea* sp. **(MN908320)** | MN932340 | 100 |
| *Pseudomonas* sp. **(MN908321)** | MT072143 | 100 |
| *Pseudomonas* sp. **(MN908322)** | MN889010 | 100 |
| *Paenibacillus* sp. **(MN908323)** | AM162327 | 99.63 |
| *Serratia* sp. **(MN908324)** | MF280132 | 99.89 |
| *Serratia* sp. **(MN908325)** | MG719570 | 100 |
| *Staphylococcus* sp. **(MN908326)** | MT071657 | 100 |
| *Streptomyces* sp. **(MN908327)** | KM522701 | 99.89 |
| *Trabulsiella* sp. **(MN908328)** | KR902558 | 99.81 |
| *Trabulsiella* sp. **(MN908329)** | NR_043860 | 99.51 |
| *Trabulsiella* sp. **(MN908330)** | NR_114235 | 100 |

**Table S2: Relative abundances of the bacterial communities obtained from Nanopore Sequencing.**

| Phylum | Class | Order | Family | Genus | Major Worker | Minor worker | Nymph | Comb | Female Alate | Male Alate |
|---|---|---|---|---|---|---|---|---|---|---|
| Acidobacteria | Acidobacteria | Acidobacteriales | Acidobacteriaceae | *Acidobacterium* | 0.001 | 0.000 | 0.000 | 0.007 | 0.000 | 0.000 |
| Acidobacteria | Acidobacteria | Acidobacteriales | Acidobacteriaceae | AMD cluster | 0.000 | 0.000 | 0.000 | 0.004 | 0.000 | 0.000 |
| Acidobacteria | Acidobacteria | Acidobacteriales | Acidobacteriaceae | marine benthic group 1 | 0.000 | 0.000 | 0.000 | 0.004 | 0.000 | 0.000 |
| Acidobacteria | Acidobacteria | Acidobacteriales | Acidobacteriaceae | marine benthic group 2 | 0.000 | 0.000 | 0.000 | 0.004 | 0.000 | 0.000 |
| Acidobacteria | Acidobacteria | Acidobacteriales | Acidobacteriaceae | Uncultured 1 | 0.000 | 0.000 | 0.000 | 0.057 | 0.000 | 0.000 |
| Acidobacteria | Acidobacteria | Acidobacteriales | Acidobacteriaceae | Uncultured 15 | 0.000 | 0.000 | 0.000 | 0.004 | 0.000 | 0.000 |
| Acidobacteria | Acidobacteria | Acidobacteriales | Acidobacteriaceae | Uncultured 18 | 0.000 | 0.000 | 0.000 | 0.018 | 0.000 | 0.000 |
| Acidobacteria | Acidobacteria | Acidobacteriales | Acidobacteriaceae | Uncultured 2 | 0.000 | 0.000 | 0.000 | 0.011 | 0.000 | 0.000 |
| Acidobacteria | Acidobacteria | Acidobacteriales | Acidobacteriaceae | Uncultured 26 | 0.000 | 0.000 | 0.000 | 0.093 | 0.000 | 0.000 |
| Acidobacteria | Acidobacteria | Acidobacteriales | Acidobacteriaceae | Uncultured 27 | 0.000 | 0.000 | 0.000 | 0.007 | 0.000 | 0.000 |
| Acidobacteria | Acidobacteria | Acidobacteriales | Acidobacteriaceae | Uncultured 28 | 0.000 | 0.000 | 0.000 | 0.004 | 0.000 | 0.000 |
| Acidobacteria | Acidobacteria | Acidobacteriales | Acidobacteriaceae | Uncultured 29 | 0.000 | 0.000 | 0.000 | 0.014 | 0.000 | 0.001 |
| Acidobacteria | Acidobacteriia | Acidobacteriales | Acidobacteriaceae | *Candidatus* Chloroacidobacterium | 0.005 | 0.000 | 0.000 | 0.096 | 0.000 | 0.000 |
| Acidobacteria | Acidobacteriia | Acidobacteriales | Acidobacteriaceae | *Candidatus* Solibacter | 0.001 | 0.000 | 0.000 | 0.011 | 0.004 | 0.001 |
| Acidobacteria | Acidobacteriia | Acidobacteriales | Acidobacteriaceae | *Edaphobacter* | 0.001 | 0.000 | 0.013 | 0.000 | 0.000 | 0.000 |
| Acidobacteria | Acidobacteriia | Acidobacteriales | Acidobacteriaceae | Mixed environment cluster | 0.002 | 0.000 | 0.000 | 0.011 | 0.000 | 0.000 |
| Acidobacteria | Acidobacteriia | Acidobacteriales | Acidobacteriaceae | *Terriglobus* | 0.001 | 0.000 | 0.000 | 0.000 | 0.000 | 0.000 |
| Acidobacteria | Acidobacteriia | Acidobacteriales | Acidobacteriaceae | Uncultured 22 | 0.002 | 0.000 | 0.000 | 0.043 | 0.000 | 0.000 |
| Acidobacteria | Acidobacteriia | Acidobacteriales | Acidobacteriaceae | Uncultured 23 | 0.061 | 0.032 | 0.013 | 0.018 | 0.004 | 0.001 |
| Acidobacteria | Acidobacteriia | Acidobacteriales | Acidobacteriaceae | Uncultured 25 | 0.001 | 0.000 | 0.000 | 0.032 | 0.000 | 0.000 |
| Acidobacteria | Acidobacteriia | Acidobacteriales | Acidobacteriaceae | Uncultured 31 | 0.005 | 0.032 | 0.000 | 0.363 | 0.012 | 0.001 |
| Acidobacteria | Acidobacteriia | Acidobacteriales | Acidobacteriaceae | Uncultured 7 | 0.001 | 0.000 | 0.000 | 0.000 | 0.000 | 0.000 |
| Acidobacteria | Acidobacteriia | Acidobacteriales | Acidobacteriaceae | Uncultured 9 | 0.004 | 0.000 | 0.000 | 0.031 | 0.000 | 0.000 |
| Acidobacteria | Blastocatellia | Blastocatellales | Blastocatellaceae | *Aridibacter* | 0.001 | 0.000 | 0.000 | 0.046 | 0.000 | 0.000 |
| Acidobacteria | Holophagae | CA002 | Unclassified | Unclassified | 0.000 | 0.000 | 0.000 | 0.000 | 0.000 | 0.001 |

**Table S2 continued.**

| | | | | | | | | | | |
|---|---|---|---|---|---|---|---|---|---|---|
| Acidobacteria | Holophagae | Cluster 32-20 | Unclassified | Unclassified | 0.000 | 0.000 | 0.000 | 0.004 | 0.000 | 0.000 |
| Acidobacteria | Holophagae | F-1404R | Unclassified | Unclassified | 0.000 | 0.032 | 0.000 | 0.018 | 0.000 | 0.000 |
| Acidobacteria | Holophagae | Holophagales | Holophagaceae | marine group | 0.000 | 0.000 | 0.000 | 0.004 | 0.000 | 0.000 |
| Acidobacteria | Holophagae | iii1 8 | Unclassified | Unclassified | 0.000 | 0.000 | 0.000 | 0.025 | 0.004 | 0.001 |
| Acidobacteria | Holophagae | NKB17 | Unclassified | Unclassified | 0.000 | 0.000 | 0.000 | 0.004 | 0.000 | 0.000 |
| Acidobacteria | Holophagae | NS72 | Unclassified | Unclassified | 0.000 | 0.000 | 0.000 | 0.007 | 0.004 | 0.000 |
| Acidobacteria | Holophagae | SJA-36 | Unclassified | Unclassified | 0.000 | 0.000 | 0.000 | 0.000 | 0.004 | 0.000 |
| Acidobacteria | Holophagae | Sva0725 | Unclassified | Unclassified | 0.000 | 0.000 | 0.000 | 0.011 | 0.000 | 0.000 |
| Acidobacteria | Holophagae | Sva0725 | Unclassified | Unclassified | 0.001 | 0.000 | 0.000 | 0.000 | 0.000 | 0.000 |
| Acidobacteria | Holophagae | TK85 | Unclassified | Unclassified | 0.001 | 0.000 | 0.000 | 0.000 | 0.000 | 0.000 |
| Acidobacteria | Holophagae | Unclassified | Unclassified | Unclassified | 0.001 | 0.000 | 0.000 | 0.000 | 0.000 | 0.000 |
| Acidobacteria | RB25 | Unclassified | Unclassified | Unclassified | 0.000 | 0.032 | 0.000 | 0.025 | 0.000 | 0.000 |
| Acidobacteria | RB25 | Unclassified | Unclassified | Unclassified | 0.002 | 0.000 | 0.000 | 0.000 | 0.000 | 0.000 |
| Actinobacteria | Acidimicrobiia | Acidimicrobiales | Acidimicrobiaceae | *Aciditerrimonas* | 0.000 | 0.000 | 0.000 | 0.004 | 0.000 | 0.000 |
| Actinobacteria | Actinobacteria | Acidimicrobiales | Acidimicrobiaceae 1 | *Ilumatobacter* | 0.000 | 0.000 | 0.000 | 0.014 | 0.000 | 0.000 |
| Actinobacteria | Actinobacteria | Acidimicrobiales | Acidimicrobiaceae 1 | marine group | 0.000 | 0.000 | 0.000 | 0.004 | 0.000 | 0.001 |
| Actinobacteria | Actinobacteria | Acidimicrobiales | Acidimicrobiaceae 1 | Uncultured 3 | 0.000 | 0.000 | 0.000 | 0.011 | 0.000 | 0.000 |
| Actinobacteria | Actinobacteria | Acidimicrobiales | Acidimicrobiaceae 1 | Uncultured 5 | 0.000 | 0.000 | 0.000 | 0.011 | 0.004 | 0.000 |
| Actinobacteria | Actinobacteria | Acidimicrobiales | Acidimicrobiaceae 2 | Unclassified | 0.000 | 0.000 | 0.000 | 0.004 | 0.000 | 0.000 |
| Actinobacteria | Actinobacteria | Acidimicrobiales | Candidatus Microthrix | *Candidatus* Microthrix | 0.000 | 0.000 | 0.000 | 0.007 | 0.000 | 0.000 |
| Actinobacteria | Actinobacteria | Acidimicrobiales | Iamiaceae | *Iamia* | 0.000 | 0.000 | 0.000 | 0.029 | 0.000 | 0.000 |
| Actinobacteria | Actinobacteria | Acidimicrobiales | marine group 1 | Unclassified | 0.000 | 0.000 | 0.000 | 0.007 | 0.000 | 0.000 |
| Actinobacteria | Actinobacteria | Acidimicrobiales | marine group 2 | Unclassified | 0.000 | 0.000 | 0.000 | 0.011 | 0.000 | 0.000 |
| Actinobacteria | Actinobacteria | Acidimicrobiales | Unclassified | Unclassified | 0.000 | 0.000 | 0.000 | 0.014 | 0.004 | 0.000 |
| Actinobacteria | Actinobacteria | Acidimicrobiales | Uncultured 2 | Unclassified | 0.000 | 0.000 | 0.000 | 0.011 | 0.000 | 0.000 |
| Actinobacteria | Actinobacteria | Acidimicrobiales | Uncultured 3 | Unclassified | 0.000 | 0.000 | 0.000 | 0.004 | 0.000 | 0.000 |
| Actinobacteria | Actinobacteria | Acidimicrobiales | Uncultured 4 | Unclassified | 0.001 | 0.000 | 0.000 | 0.011 | 0.000 | 0.000 |

**Table S2 continued.**

| | | | | | | | | | | |
|---|---|---|---|---|---|---|---|---|---|---|
| Actinobacteria | Actinobacteria | Acidimicrobiales | Uncultured 6 | Unclassified | 0.002 | 0.000 | 0.000 | 0.018 | 0.000 | 0.000 |
| Actinobacteria | Actinobacteria | Actinomycetales | Actinomycetaceae | *Actinomyces* | 0.001 | 0.000 | 0.000 | 0.000 | 0.000 | 0.000 |
| Actinobacteria | Actinobacteria | Actinomycetales | Kineosporiaceae | *Kineosporia* | 0.001 | 0.000 | 0.000 | 0.004 | 0.004 | 0.001 |
| Actinobacteria | Actinobacteria | Actinomycetales | Termite cluster 1 | Subcluster a | 0.001 | 0.000 | 0.000 | 0.000 | 0.000 | 0.000 |
| Actinobacteria | Actinobacteria | Actinomycetales | Termite cluster 2 | Unclassified | 0.001 | 0.000 | 0.000 | 0.000 | 0.000 | 0.000 |
| Actinobacteria | Actinobacteria | Actinomycetales 1 | Streptomycetaceae | *Streptomyces* 1 | 0.001 | 0.000 | 0.000 | 0.057 | 0.000 | 0.000 |
| Actinobacteria | Actinobacteria | Actinomycetales 1 | Streptomycetaceae | *Streptomyces* 3 | 0.000 | 0.000 | 0.000 | 0.000 | 0.000 | 0.001 |
| Actinobacteria | Actinobacteria | Actinomycetales 1 | Streptomycetaceae | *Streptomyces* 5 | 0.000 | 0.000 | 0.000 | 0.004 | 0.000 | 0.000 |
| Actinobacteria | Actinobacteria | Actinopolysporales | Actinopolysporaceae | *Actinopolyspora* | 0.000 | 0.000 | 0.000 | 0.004 | 0.000 | 0.000 |
| Actinobacteria | Actinobacteria | AKIW543 | Unclassified | Unclassified | 0.000 | 0.000 | 0.000 | 0.078 | 0.000 | 0.000 |
| Actinobacteria | Actinobacteria | AKIW543 | Unclassified | Unclassified | 0.002 | 0.000 | 0.000 | 0.000 | 0.000 | 0.000 |
| Actinobacteria | Actinobacteria | Bifidobacteriales | Bifidobacteriaceae | *Bifidibacterium* | 0.000 | 0.000 | 0.000 | 0.007 | 0.000 | 0.000 |
| Actinobacteria | Actinobacteria | Catenulisporales | Catenulisporaceae | *Catenulispora* | 0.001 | 0.000 | 0.000 | 0.000 | 0.000 | 0.000 |
| Actinobacteria | Actinobacteria | Coriobacteriales | Coriobacteriaceae | *Collinsella* | 0.001 | 0.000 | 0.000 | 0.004 | 0.000 | 0.000 |
| Actinobacteria | Actinobacteria | Coriobacteriales | Coriobacteriaceae | *Enterorhabdus* | 0.001 | 0.000 | 0.026 | 0.000 | 0.004 | 0.005 |
| Actinobacteria | Actinobacteria | Coriobacteriales | Coriobacteriaceae | marine group | 0.007 | 0.000 | 0.013 | 0.018 | 0.000 | 0.000 |
| Actinobacteria | Actinobacteria | Coriobacteriales | Coriobacteriaceae | Unclassified | 0.000 | 0.000 | 0.000 | 0.000 | 0.000 | 0.001 |
| Actinobacteria | Actinobacteria | Coriobacteriales | Coriobacteriaceae | Uncultured 10 | 1.169 | 1.454 | 1.843 | 0.314 | 0.809 | 0.438 |
| Actinobacteria | Actinobacteria | Coriobacteriales | Coriobacteriaceae | Uncultured 11 | 0.004 | 0.000 | 0.000 | 0.000 | 0.004 | 0.001 |
| Actinobacteria | Actinobacteria | Coriobacteriales | Coriobacteriaceae | Uncultured 4 | 0.000 | 0.000 | 0.013 | 0.000 | 0.000 | 0.000 |
| Actinobacteria | Actinobacteria | Coriobacteriales | Coriobacteriaceae | Uncultured 8 | 0.005 | 0.000 | 0.026 | 0.011 | 0.004 | 0.000 |
| Actinobacteria | Actinobacteria | Coriobacteriales | Coriobacteriaceae | Uncultured 8 | 0.000 | 0.000 | 0.000 | 0.000 | 0.000 | 0.006 |
| Actinobacteria | Actinobacteria | Coriobacteriales | Coriobacteriaceae | Unclassified | 0.002 | 0.000 | 0.000 | 0.000 | 0.000 | 0.000 |
| Actinobacteria | Actinobacteria | Corynebacteriales | Corynebacteriaceae | *Corynebacterium* | 0.000 | 0.000 | 0.000 | 0.004 | 0.008 | 0.000 |
| Actinobacteria | Actinobacteria | Corynebacteriales | Corynebacteriaceae | *Corynebacterium* 1 | 0.000 | 0.000 | 0.013 | 0.014 | 0.012 | 0.000 |
| Actinobacteria | Actinobacteria | Corynebacteriales | Corynebacteriaceae | *Corynebacterium* 4 | 0.000 | 0.000 | 0.000 | 0.004 | 0.000 | 0.000 |
| Actinobacteria | Actinobacteria | Corynebacteriales | Corynebacteriaceae | Unclassified | 0.000 | 0.000 | 0.000 | 0.000 | 0.000 | 0.001 |

**Table S2 continued.**

| | | | | | | | | | | |
|---|---|---|---|---|---|---|---|---|---|---|
| Actinobacteria | Actinobacteria | Corynebacteriales | Dietziaceae | *Dietzia* | 0.000 | 0.000 | 0.000 | 0.004 | 0.000 | 0.000 |
| Actinobacteria | Actinobacteria | Corynebacteriales | Mycobacteriaceae | *Mycobacterium* | 0.001 | 0.000 | 0.000 | 0.456 | 0.000 | 0.002 |
| Actinobacteria | Actinobacteria | Corynebacteriales | Nocardiaceae | *Nocardia* | 0.001 | 0.000 | 0.000 | 0.007 | 0.000 | 0.000 |
| Actinobacteria | Actinobacteria | Corynebacteriales | Nocardiaceae | *Rhodococcus* | 0.000 | 0.000 | 0.000 | 0.018 | 0.004 | 0.000 |
| Actinobacteria | Actinobacteria | Corynebacteriales | Nocardiaceae | *Rhodococcus* 2 | 0.000 | 0.000 | 0.000 | 0.004 | 0.000 | 0.000 |
| Actinobacteria | Actinobacteria | Corynebacteriales | Nocardiaceae | *Smaragdicoccus* | 0.000 | 0.000 | 0.000 | 0.000 | 0.000 | 0.002 |
| Actinobacteria | Actinobacteria | Corynebacteriales | Tsukamurellaceae | *Tsukamurella* | 0.000 | 0.000 | 0.000 | 0.004 | 0.000 | 0.000 |
| Actinobacteria | Actinobacteria | Corynebacteriales | Unclassified | Unclassified | 0.000 | 0.000 | 0.000 | 0.004 | 0.000 | 0.000 |
| Actinobacteria | Actinobacteria | Frankiales | Nakamurellaceae | Unclassified | 0.000 | 0.000 | 0.000 | 0.007 | 0.000 | 0.000 |
| Actinobacteria | Actinobacteria | Geodermatophilales | Geodermatophilaceae | *Geodermatophilus* | 0.000 | 0.000 | 0.000 | 0.011 | 0.000 | 0.002 |
| Actinobacteria | Actinobacteria | Glycomycetales | Glycomycetaceae | *Glycomyces* | 0.000 | 0.000 | 0.000 | 0.004 | 0.000 | 0.000 |
| Actinobacteria | Actinobacteria | Kineosporiales | Kineosporiaceae | *Kineococcus* | 0.000 | 0.000 | 0.000 | 0.004 | 0.000 | 0.000 |
| Actinobacteria | Actinobacteria | Kineosporiales | Kineosporiaceae | *Quadrisphaera* | 0.000 | 0.000 | 0.000 | 0.000 | 0.000 | 0.001 |
| Actinobacteria | Actinobacteria | Kineosporiales | Kineosporiaceae | *Thalassiella* | 0.000 | 0.000 | 0.000 | 0.004 | 0.000 | 0.000 |
| Actinobacteria | Actinobacteria | MB-A2-108 | Unclassified | Unclassified | 0.000 | 0.000 | 0.000 | 0.036 | 0.000 | 0.000 |
| Actinobacteria | Actinobacteria | Micrococcales | Beutenbergiaceae | *Beutenbergia* | 0.000 | 0.000 | 0.000 | 0.004 | 0.000 | 0.000 |
| Actinobacteria | Actinobacteria | Micrococcales | Beutenbergiaceae | *Miniimonas* | 0.000 | 0.000 | 0.000 | 0.004 | 0.000 | 0.000 |
| Actinobacteria | Actinobacteria | Micrococcales | Beutenbergiaceae | *Serinibacter* | 0.001 | 0.000 | 0.000 | 0.000 | 0.004 | 0.000 |
| Actinobacteria | Actinobacteria | Micrococcales | Brevibacteriaceae | *Brevibacterium* | 0.000 | 0.000 | 0.000 | 0.011 | 0.000 | 0.000 |
| Actinobacteria | Actinobacteria | Micrococcales | Cellulomonadaceae | *Cellulomonas* | 0.001 | 0.000 | 0.000 | 0.093 | 0.000 | 0.000 |
| Actinobacteria | Actinobacteria | Micrococcales | Dermabacteraceae | *Brachybacterium* | 0.002 | 0.000 | 0.000 | 0.014 | 0.000 | 0.001 |
| Actinobacteria | Actinobacteria | Micrococcales | Dermacoccaceae | *Kytococcus* | 0.000 | 0.000 | 0.000 | 0.004 | 0.000 | 0.000 |
| Actinobacteria | Actinobacteria | Micrococcales | Dermatophilaceae | *Dermatophilus* | 0.001 | 0.000 | 0.000 | 0.000 | 0.000 | 0.001 |
| Actinobacteria | Actinobacteria | Micrococcales | Intrasporangiaceae | *Janibacter* | 0.000 | 0.000 | 0.000 | 0.014 | 0.000 | 0.000 |
| Actinobacteria | Actinobacteria | Micrococcales | Intrasporangiaceae | *Knoellia* | 0.000 | 0.000 | 0.000 | 0.004 | 0.000 | 0.000 |
| Actinobacteria | Actinobacteria | Micrococcales | Intrasporangiaceae | *Kribbia* | 0.000 | 0.000 | 0.000 | 0.004 | 0.000 | 0.000 |
| Actinobacteria | Actinobacteria | Micrococcales | Intrasporangiaceae | *Phycicoccus* | 0.000 | 0.000 | 0.000 | 0.004 | 0.000 | 0.000 |

**Table S2 continued.**

| | | | | | | | | | | |
|---|---|---|---|---|---|---|---|---|---|---|
| Actinobacteria | Actinobacteria | Micrococcales | Intrasporangiaceae | *Serinicoccus* | 0.000 | 0.000 | 0.000 | 0.004 | 0.000 | 0.000 |
| Actinobacteria | Actinobacteria | Micrococcales | Microbacteriaceae | *Agromyces* | 0.000 | 0.000 | 0.000 | 0.025 | 0.000 | 0.000 |
| Actinobacteria | Actinobacteria | Micrococcales | Microbacteriaceae | *Allohumibacter* | 0.000 | 0.000 | 0.000 | 0.004 | 0.000 | 0.000 |
| Actinobacteria | Actinobacteria | Micrococcales | Microbacteriaceae | *Clavibacter* | 0.001 | 0.000 | 0.000 | 0.014 | 0.000 | 0.000 |
| Actinobacteria | Actinobacteria | Micrococcales | Microbacteriaceae | *Curtobacterium* | 0.000 | 0.000 | 0.000 | 0.018 | 0.000 | 0.000 |
| Actinobacteria | Actinobacteria | Micrococcales | Microbacteriaceae | *Galbitalea* | 0.000 | 0.000 | 0.000 | 0.004 | 0.000 | 0.000 |
| Actinobacteria | Actinobacteria | Micrococcales | Microbacteriaceae | *Leifsonia* | 0.001 | 0.000 | 0.000 | 0.007 | 0.000 | 0.000 |
| Actinobacteria | Actinobacteria | Micrococcales | Microbacteriaceae | *Leucobacter* | 0.001 | 0.000 | 0.000 | 0.004 | 0.000 | 0.000 |
| Actinobacteria | Actinobacteria | Micrococcales | Microbacteriaceae | *Microbacterium* | 0.002 | 0.000 | 0.000 | 0.029 | 0.004 | 0.003 |
| Actinobacteria | Actinobacteria | Micrococcales | Microbacteriaceae | *Microcella* | 0.000 | 0.000 | 0.000 | 0.004 | 0.000 | 0.000 |
| Actinobacteria | Actinobacteria | Micrococcales | Microbacteriaceae | *Pseudoclavibacter* | 0.000 | 0.000 | 0.000 | 0.007 | 0.000 | 0.000 |
| Actinobacteria | Actinobacteria | Micrococcales | Microbacteriaceae | *Rudaibacter* | 0.000 | 0.000 | 0.000 | 0.004 | 0.000 | 0.000 |
| Actinobacteria | Actinobacteria | Micrococcales | Micrococcaceae | *Arthrobacter* | 0.001 | 0.000 | 0.000 | 0.078 | 0.000 | 0.001 |
| Actinobacteria | Actinobacteria | Micrococcales | Micrococcaceae | *Kocuria* | 0.001 | 0.000 | 0.000 | 0.007 | 0.000 | 0.000 |
| Actinobacteria | Actinobacteria | Micrococcales | Micrococcaceae | *Micrococcus* | 0.000 | 0.000 | 0.000 | 0.021 | 0.000 | 0.002 |
| Actinobacteria | Actinobacteria | Micrococcales | Micrococcaceae | *Nesterenkonia* | 0.000 | 0.000 | 0.000 | 0.000 | 0.004 | 0.001 |
| Actinobacteria | Actinobacteria | Micrococcales | Micrococcaceae | *Rothia* | 0.000 | 0.000 | 0.000 | 0.014 | 0.000 | 0.000 |
| Actinobacteria | Actinobacteria | Micrococcales | Promicromonosporaceae | *Cellulosimicrobium* | 0.000 | 0.000 | 0.000 | 0.021 | 0.000 | 0.000 |
| Actinobacteria | Actinobacteria | Micrococcales | Promicromonosporaceae | *Isoptericola* | 0.000 | 0.000 | 0.000 | 0.007 | 0.000 | 0.000 |
| Actinobacteria | Actinobacteria | Micrococcales | Promicromonosporaceae | *Promicromonospora* | 0.000 | 0.000 | 0.000 | 0.007 | 0.000 | 0.000 |
| Actinobacteria | Actinobacteria | Micrococcales | Unclassified | *Timonella* | 0.000 | 0.000 | 0.000 | 0.004 | 0.000 | 0.000 |
| Actinobacteria | Actinobacteria | Micrococcales 1 | Micrococcaceae | *Arthrobacter* 10 | 0.000 | 0.000 | 0.000 | 0.018 | 0.000 | 0.000 |
| Actinobacteria | Actinobacteria | Micrococcales 1 | Micrococcaceae | *Arthrobacter* 12 | 0.000 | 0.000 | 0.000 | 0.004 | 0.000 | 0.000 |
| Actinobacteria | Actinobacteria | Micrococcales 1 | Micrococcaceae | *Arthrobacter* 13 | 0.000 | 0.000 | 0.000 | 0.007 | 0.004 | 0.000 |
| Actinobacteria | Actinobacteria | Micrococcales 1 | Micrococcaceae | *Arthrobacter* 14 | 0.000 | 0.000 | 0.000 | 0.004 | 0.000 | 0.000 |
| Actinobacteria | Actinobacteria | Micrococcales 1 | Micrococcaceae | *Arthrobacter* 17 | 0.000 | 0.000 | 0.000 | 0.071 | 0.000 | 0.000 |
| Actinobacteria | Actinobacteria | Micrococcales 1 | Micrococcaceae | *Arthrobacter* 21 | 0.000 | 0.000 | 0.000 | 0.007 | 0.000 | 0.001 |

**Table S2 continued.**

| | | | | | | | | | | |
|---|---|---|---|---|---|---|---|---|---|---|
| Actinobacteria | Actinobacteria | Micrococcales 1 | Micrococcaceae | *Arthrobacter* 6 | 0.000 | 0.000 | 0.000 | 0.021 | 0.000 | 0.000 |
| Actinobacteria | Actinobacteria | Micrococcales 1 | Micrococcaceae | *Arthrobacter* 8 | 0.000 | 0.000 | 0.000 | 0.004 | 0.000 | 0.000 |
| Actinobacteria | Actinobacteria | Micrococcales 1 | Micrococcaceae | *Citricoccus* | 0.000 | 0.000 | 0.000 | 0.004 | 0.000 | 0.000 |
| Actinobacteria | Actinobacteria | Micrococcales 1 | Micrococcaceae | *Kocuria* 2 | 0.002 | 0.000 | 0.000 | 0.000 | 0.000 | 0.000 |
| Actinobacteria | Actinobacteria | Micrococcales 1 | Micrococcaceae | *Micrococcus* 1 | 0.000 | 0.000 | 0.000 | 0.004 | 0.000 | 0.000 |
| Actinobacteria | Actinobacteria | Micrococcales 1 | Micrococcaceae | *Micrococcus* 3 | 0.000 | 0.000 | 0.000 | 0.004 | 0.000 | 0.000 |
| Actinobacteria | Actinobacteria | Micrococcales 1 | Micrococcaceae | Unclassified | 0.000 | 0.000 | 0.000 | 0.011 | 0.000 | 0.000 |
| Actinobacteria | Actinobacteria | Micrococcales 1 | Sanguibacteraceae | *Sanguibacter* | 0.001 | 0.032 | 0.000 | 0.000 | 0.000 | 0.000 |
| Actinobacteria | Actinobacteria | Micrococcales 3 | Cellulomonadaceae | *Cellulomonas* 1 | 0.000 | 0.000 | 0.000 | 0.029 | 0.000 | 0.001 |
| Actinobacteria | Actinobacteria | Micrococcales 3 | Cellulomonadaceae | *Cellulomonas* 2 | 0.000 | 0.000 | 0.000 | 0.014 | 0.000 | 0.000 |
| Actinobacteria | Actinobacteria | Micrococcales 3 | Microbacteriaceae | *Agreia* | 0.000 | 0.000 | 0.000 | 0.004 | 0.000 | 0.000 |
| Actinobacteria | Actinobacteria | Micrococcales 3 | Microbacteriaceae | *Frigoribacterium* 1 | 0.000 | 0.000 | 0.000 | 0.021 | 0.000 | 0.000 |
| Actinobacteria | Actinobacteria | Micrococcales 3 | Microbacteriaceae | *Schumannella* | 0.000 | 0.000 | 0.000 | 0.004 | 0.000 | 0.000 |
| Actinobacteria | Actinobacteria | Micrococcales 3 | Microbacteriaceae | Unclassified | 0.000 | 0.000 | 0.000 | 0.007 | 0.000 | 0.000 |
| Actinobacteria | Actinobacteria | Micrococcales 3 | Microbacteriaceae | Uncultured 1 | 0.001 | 0.000 | 0.000 | 0.007 | 0.000 | 0.000 |
| Actinobacteria | Actinobacteria | Micrococcales 3 | Microbacteriaceae | Uncultured 2 | 0.000 | 0.000 | 0.000 | 0.021 | 0.000 | 0.000 |
| Actinobacteria | Actinobacteria | Micrococcales 3 | Microbacteriaceae | Unclassified | 0.001 | 0.000 | 0.000 | 0.000 | 0.000 | 0.000 |
| Actinobacteria | Actinobacteria | Micrococcales 3 | Microbacteriaceae | *Zimmermanella* 2 | 0.001 | 0.000 | 0.000 | 0.000 | 0.000 | 0.000 |
| Actinobacteria | Actinobacteria | Micrococcales 3 | Microbacteriaceae | *Zimmermanella* 3 | 0.000 | 0.000 | 0.000 | 0.004 | 0.000 | 0.000 |
| Actinobacteria | Actinobacteria | Micrococcales 4 | Intrasporangiaceae 1 | *Lapillicoccus* | 0.000 | 0.000 | 0.000 | 0.004 | 0.000 | 0.000 |
| Actinobacteria | Actinobacteria | Micrococcales 4 | Intrasporangiaceae 1 | *Oryzihumus* | 0.000 | 0.000 | 0.000 | 0.004 | 0.000 | 0.000 |
| Actinobacteria | Actinobacteria | Micrococcales 4 | Intrasporangiaceae 1 | *Terracoccus* sp a | 0.000 | 0.000 | 0.000 | 0.004 | 0.000 | 0.000 |
| Actinobacteria | Actinobacteria | Micrococcales 4 | Intrasporangiaceae 1 | *Tetrasphaera* | 0.000 | 0.000 | 0.000 | 0.007 | 0.000 | 0.000 |
| Actinobacteria | Actinobacteria | Micrococcales 4 | Intrasporangiaceae 1 | Unclassified | 0.000 | 0.000 | 0.000 | 0.004 | 0.000 | 0.000 |
| Actinobacteria | Actinobacteria | Micromonosporales | Micromonosporaceae | *Actinoplanes* | 0.000 | 0.000 | 0.000 | 0.021 | 0.000 | 0.000 |
| Actinobacteria | Actinobacteria | Micromonosporales | Micromonosporaceae | *Actinoplanes* 5 | 0.000 | 0.000 | 0.000 | 0.004 | 0.000 | 0.000 |
| Actinobacteria | Actinobacteria | Micromonosporales | Micromonosporaceae | *Catenuloplanes* | 0.000 | 0.000 | 0.000 | 0.007 | 0.000 | 0.000 |

**Table S2 continued.**

| | | | | | | | | | | |
|---|---|---|---|---|---|---|---|---|---|---|
| Actinobacteria | Actinobacteria | Micromonosporales | Micromonosporaceae | *Krasilnikovia* | 0.000 | 0.000 | 0.000 | 0.004 | 0.000 | 0.000 |
| Actinobacteria | Actinobacteria | Micromonosporales | Micromonosporaceae | *Micromonospora* | 0.000 | 0.000 | 0.000 | 0.029 | 0.000 | 0.000 |
| Actinobacteria | Actinobacteria | Micromonosporales | Micromonosporaceae | *Micromonospora* 1 | 0.000 | 0.000 | 0.000 | 0.004 | 0.000 | 0.000 |
| Actinobacteria | Actinobacteria | Micromonosporales | Micromonosporaceae | *Pseudosporangium* | 0.000 | 0.000 | 0.000 | 0.004 | 0.000 | 0.000 |
| Actinobacteria | Actinobacteria | Nakamurellales | Nakmurellaceae | *Nakamurella* | 0.000 | 0.000 | 0.000 | 0.004 | 0.000 | 0.000 |
| Actinobacteria | Actinobacteria | PeM15 | Unclassified | Unclassified | 0.000 | 0.000 | 0.000 | 0.004 | 0.000 | 0.001 |
| Actinobacteria | Actinobacteria | Propionibacteriales | Nocardioidaceae | *Actinopolymorpha* | 0.001 | 0.000 | 0.000 | 0.004 | 0.000 | 0.000 |
| Actinobacteria | Actinobacteria | Propionibacteriales | Nocardioidaceae | *Aeromicrobium* | 0.001 | 0.000 | 0.000 | 0.021 | 0.000 | 0.005 |
| Actinobacteria | Actinobacteria | Propionibacteriales | Nocardioidaceae | *Kribbella* | 0.000 | 0.000 | 0.000 | 0.004 | 0.000 | 0.000 |
| Actinobacteria | Actinobacteria | Propionibacteriales | Nocardioidaceae | *Marmoricola* | 0.000 | 0.000 | 0.000 | 0.021 | 0.004 | 0.002 |
| Actinobacteria | Actinobacteria | Propionibacteriales | Nocardioidaceae | *Nocardioides* | 0.002 | 0.000 | 0.013 | 0.253 | 0.008 | 0.005 |
| Actinobacteria | Actinobacteria | Propionibacteriales | Nocardioidaceae | *Pimelobacter* | 0.000 | 0.000 | 0.000 | 0.004 | 0.000 | 0.000 |
| Actinobacteria | Actinobacteria | Propionibacteriales | Nocardioidaceae | *Propionicimonas* | 0.000 | 0.000 | 0.000 | 0.004 | 0.000 | 0.001 |
| Actinobacteria | Actinobacteria | Propionibacteriales | Propionibacteriaceae | *Aestuariimicrobium* | 0.098 | 0.000 | 0.000 | 0.029 | 0.020 | 0.006 |
| Actinobacteria | Actinobacteria | Propionibacteriales | Propionibacteriaceae | *Brooklawnia* | 0.002 | 0.000 | 0.000 | 0.000 | 0.000 | 0.000 |
| Actinobacteria | Actinobacteria | Propionibacteriales | Propionibacteriaceae | *Friedmanniella* | 0.001 | 0.000 | 0.000 | 0.000 | 0.004 | 0.000 |
| Actinobacteria | Actinobacteria | Propionibacteriales | Propionibacteriaceae | *Luteococcus* | 0.004 | 0.000 | 0.000 | 0.004 | 0.000 | 0.000 |
| Actinobacteria | Actinobacteria | Propionibacteriales | Propionibacteriaceae | *Propionibacterium* | 0.001 | 0.000 | 0.000 | 0.004 | 0.000 | 0.000 |
| Actinobacteria | Actinobacteria | Propionibacteriales | Propionibacteriaceae | *Tessaracoccus* | 0.014 | 0.000 | 0.000 | 0.000 | 0.000 | 0.000 |
| Actinobacteria | Actinobacteria | Propionibacteriales | Propionibacteriaceae | Uncultured 1 | 0.002 | 0.000 | 0.000 | 0.000 | 0.000 | 0.000 |
| Actinobacteria | Actinobacteria | Propionibacteriales | Propionibacteriaceae | Uncultured 1 | 0.000 | 0.000 | 0.000 | 0.000 | 0.000 | 0.001 |
| Actinobacteria | Actinobacteria | Propionibacteriales | Propionibacteriaceae | Uncultured 2 | 0.001 | 0.000 | 0.000 | 0.000 | 0.000 | 0.000 |
| Actinobacteria | Actinobacteria | Propionibacteriales | Propionibacteriaceae | Unclassified | 0.005 | 0.000 | 0.000 | 0.000 | 0.000 | 0.000 |
| Actinobacteria | Actinobacteria | Pseudonocardiales | Pseudonocardiaceae | *Actinomycetospora* | 0.000 | 0.000 | 0.000 | 0.011 | 0.000 | 0.000 |
| Actinobacteria | Actinobacteria | Pseudonocardiales | Pseudonocardiaceae | *Actinophytocola* | 0.000 | 0.000 | 0.000 | 0.004 | 0.000 | 0.000 |
| Actinobacteria | Actinobacteria | Pseudonocardiales | Pseudonocardiaceae | *Kibdelosporangium* | 0.000 | 0.000 | 0.000 | 0.004 | 0.000 | 0.000 |
| Actinobacteria | Actinobacteria | Pseudonocardiales | Pseudonocardiaceae | *Lentzea* | 0.000 | 0.000 | 0.000 | 0.004 | 0.000 | 0.000 |

**Table S2 continued.**

| | | | | | | | | | | |
|---|---|---|---|---|---|---|---|---|---|---|
| Actinobacteria | Actinobacteria | Pseudonocardiales | Pseudonocardiaceae | *Pseudonocardia* | 0.000 | 0.000 | 0.000 | 0.025 | 0.000 | 0.000 |
| Actinobacteria | Actinobacteria | Pseudonocardiales | Pseudonocardiaceae | *Saccharopolyspora* | 0.000 | 0.000 | 0.000 | 0.014 | 0.000 | 0.000 |
| Actinobacteria | Actinobacteria | Pseudonocardiales 6 | Actinosynnemataceae | *Saccharothrix* | 0.000 | 0.000 | 0.000 | 0.004 | 0.000 | 0.000 |
| Actinobacteria | Actinobacteria | RL185-aaj71c12 | Unclassified | Unclassified | 0.000 | 0.000 | 0.000 | 0.004 | 0.000 | 0.000 |
| Actinobacteria | Actinobacteria | Solirubrobacterales | Cluster 288-2 | Unclassified | 0.000 | 0.000 | 0.000 | 0.011 | 0.000 | 0.000 |
| Actinobacteria | Actinobacteria | Solirubrobacterales | Cluster 319-6M6 | Unclassified | 0.000 | 0.000 | 0.000 | 0.004 | 0.000 | 0.000 |
| Actinobacteria | Actinobacteria | Solirubrobacterales | Cluster 480-2 | Unclassified | 0.000 | 0.000 | 0.000 | 0.068 | 0.000 | 0.000 |
| Actinobacteria | Actinobacteria | Solirubrobacterales | Elev-16S-1332 | Unclassified | 0.000 | 0.000 | 0.000 | 0.014 | 0.000 | 0.000 |
| Actinobacteria | Actinobacteria | Solirubrobacterales | Patulibacteraceae | *Patulibacter* | 0.000 | 0.000 | 0.000 | 0.061 | 0.000 | 0.000 |
| Actinobacteria | Actinobacteria | Solirubrobacterales | Solirubrobacteriaceae | *Solirubrobacter* | 0.002 | 0.000 | 0.000 | 0.093 | 0.004 | 0.000 |
| Actinobacteria | Actinobacteria | Solirubrobacterales | Unclassified | Unclassified | 0.000 | 0.000 | 0.000 | 0.018 | 0.000 | 0.000 |
| Actinobacteria | Actinobacteria | Solirubrobacterales | YNPFFP1 | Unclassified | 0.000 | 0.000 | 0.000 | 0.014 | 0.000 | 0.000 |
| Actinobacteria | Actinobacteria | Sporichthyales | Sporichthyaceae | *Sporichthya* | 0.000 | 0.000 | 0.000 | 0.007 | 0.000 | 0.000 |
| Actinobacteria | Actinobacteria | Streptomycetales | Streptomycetaceae | *Kitasatospora* | 0.001 | 0.000 | 0.000 | 0.004 | 0.000 | 0.000 |
| Actinobacteria | Actinobacteria | Streptomycetales | Streptomycetaceae | *Streptomyces* | 0.001 | 0.000 | 0.000 | 0.157 | 0.004 | 0.000 |
| Actinobacteria | Actinobacteria | Streptosporangiales | Nocardiopsaceae | *Nocardiopsis* | 0.000 | 0.000 | 0.000 | 0.004 | 0.000 | 0.000 |
| Actinobacteria | Actinobacteria | Streptosporangiales | Streptosporangiaceae | *Nonomuraea* | 0.000 | 0.000 | 0.000 | 0.007 | 0.000 | 0.000 |
| Actinobacteria | Actinobacteria | Streptosporangiales | Streptosporangiaceae | *Planomonospora* | 0.000 | 0.000 | 0.000 | 0.004 | 0.000 | 0.000 |
| Actinobacteria | Actinobacteria | Streptosporangiales | Streptosporangiaceae | *Sphaerisporangium* | 0.000 | 0.000 | 0.000 | 0.007 | 0.000 | 0.000 |
| Actinobacteria | Actinobacteria | Streptosporangiales | Streptosporangiaceae | Unclassified | 0.000 | 0.000 | 0.000 | 0.000 | 0.000 | 0.001 |
| Actinobacteria | Actinobacteria | Streptosporangiales | Thermomonosporaceae | *Actinocorallia* | 0.000 | 0.000 | 0.000 | 0.004 | 0.000 | 0.001 |
| Actinobacteria | Actinobacteria | Streptosporangiales | Thermomonosporaceae | *Actinomadura* | 0.000 | 0.000 | 0.000 | 0.004 | 0.000 | 0.000 |
| Actinobacteria | Actinobacteria | Streptosporangiales | Unclassified | *Thermomonospora* | 0.000 | 0.000 | 0.000 | 0.004 | 0.000 | 0.000 |
| Actinobacteria | Actinobacteria | TakashiAC-B11 | Unclassified | Unclassified | 0.000 | 0.000 | 0.000 | 0.007 | 0.000 | 0.000 |
| Actinobacteria | Coriobacteriia | Coriobacteriales | Atopobiaceae | *Atopobium* | 0.002 | 0.000 | 0.000 | 0.000 | 0.000 | 0.000 |
| Actinobacteria | Coriobacteriia | Eggerthellales | Eggerthellaceae | *Eggerthella* | 0.001 | 0.000 | 0.013 | 0.000 | 0.012 | 0.001 |
| Actinobacteria | Coriobacteriia | Eggerthellales | Eggerthellaceae | *Gordonibacter* | 0.005 | 0.000 | 0.000 | 0.000 | 0.008 | 0.002 |

**Table S2 continued.**

| | | | | | | | | | | |
|---|---|---|---|---|---|---|---|---|---|---|
| Actinobacteria | Coriobacteriia | Eggerthellales | Eggerthellaceae | *Raoultibacter* | 0.007 | 0.000 | 0.000 | 0.007 | 0.000 | 0.007 |
| Actinobacteria | Coriobacteriia | Eggerthellales | Eggerthellaceae | *Slackia* | 0.000 | 0.000 | 0.013 | 0.000 | 0.004 | 0.000 |
| Actinobacteria | Thermoleophilia | Solirubrobacterales | Conexibacteraceae | *Conexibacter* | 0.000 | 0.000 | 0.000 | 0.004 | 0.000 | 0.000 |
| Actinobacteria | Thermoleophilia | Solirubrobacterales | Parviterribacteraceae | *Parviterribacter* | 0.000 | 0.000 | 0.000 | 0.018 | 0.000 | 0.000 |
| Armatimonadetes | Unclassified | Unclassified | Unclassified | Unclassified | 0.000 | 0.000 | 0.000 | 0.018 | 0.004 | 0.000 |
| Armatimonadetes | Unclassified | Unclassified | Unclassified | Unclassified | 0.004 | 0.000 | 0.000 | 0.000 | 0.000 | 0.000 |
| Bacteroidetes | Bacteroidia | Bacteroidales | Bacteroidaceae | *Anaerorhabdus* | 0.000 | 0.000 | 0.000 | 0.000 | 0.004 | 0.000 |
| Bacteroidetes | Bacteroidia | Bacteroidales | Bacteroidaceae | *Bacteroides* | 4.256 | 2.903 | 0.103 | 4.536 | 0.622 | 0.310 |
| Bacteroidetes | Bacteroidia | Bacteroidales | Bacteroidaceae | *Mediterranea* | 0.001 | 0.000 | 0.000 | 0.000 | 0.000 | 0.000 |
| Bacteroidetes | Bacteroidia | Bacteroidales | Dysgomonadaceae | *Proteiniphilum* | 0.001 | 0.032 | 0.000 | 0.004 | 0.008 | 0.006 |
| Bacteroidetes | Bacteroidia | Bacteroidales | Dysgonamonadaceae | *Dysgonomonas* | 0.842 | 0.861 | 0.000 | 1.582 | 1.066 | 3.093 |
| Bacteroidetes | Bacteroidia | Bacteroidales | gir-aah93h0 | Unclassified | 0.000 | 0.000 | 0.000 | 0.000 | 0.004 | 0.003 |
| Bacteroidetes | Bacteroidia | Bacteroidales | Lentimicrobiaceae | *Lentimicrobium* | 0.000 | 0.000 | 0.000 | 0.000 | 0.008 | 0.000 |
| Bacteroidetes | Bacteroidia | Bacteroidales | Marinilabiaceae | *Alkaliflexus* | 0.011 | 0.000 | 0.000 | 0.014 | 0.008 | 0.002 |
| Bacteroidetes | Bacteroidia | Bacteroidales | Marinilabiaceae | Unclassified | 0.000 | 0.000 | 0.000 | 0.000 | 0.000 | 0.001 |
| Bacteroidetes | Bacteroidia | Bacteroidales | Marinilabiaceae | Uncultured 1 | 0.291 | 0.159 | 0.000 | 0.442 | 0.460 | 0.000 |
| Bacteroidetes | Bacteroidia | Bacteroidales | Marinilabiaceae | Uncultured 1 | 0.000 | 0.000 | 0.000 | 0.000 | 0.000 | 0.542 |
| Bacteroidetes | Bacteroidia | Bacteroidales | Marinilabiaceae | Uncultured 2 | 0.001 | 0.032 | 0.000 | 0.000 | 0.004 | 0.002 |
| Bacteroidetes | Bacteroidia | Bacteroidales | Odoribacteraceae | *Butyricimonas* | 0.002 | 0.000 | 0.000 | 0.004 | 0.004 | 0.000 |
| Bacteroidetes | Bacteroidia | Bacteroidales | Odoribacteraceae | *Culturomica* | 0.012 | 0.000 | 0.000 | 0.039 | 0.000 | 0.005 |
| Bacteroidetes | Bacteroidia | Bacteroidales | Odoribacteraceae | *Odoribacter* | 0.114 | 0.128 | 0.000 | 0.143 | 0.057 | 0.034 |
| Bacteroidetes | Bacteroidia | Bacteroidales | Paludibacteraceae | *Paludibacter* | 0.020 | 0.032 | 0.000 | 0.039 | 0.000 | 0.013 |
| Bacteroidetes | Bacteroidia | Bacteroidales | Porphyromonadaceae | *Microbacter* | 0.001 | 0.000 | 0.000 | 0.000 | 0.000 | 0.000 |
| Bacteroidetes | Bacteroidia | Bacteroidales | Porphyromonadaceae | *Porphyromonas* | 0.011 | 0.032 | 0.000 | 0.007 | 0.012 | 0.005 |
| Bacteroidetes | Bacteroidia | Bacteroidales | Porphyromonadaceae 2 | Termite cluster 1 | 0.004 | 0.000 | 0.000 | 0.007 | 0.004 | 0.007 |
| Bacteroidetes | Bacteroidia | Bacteroidales | Porphyromonadaceae 2 | Termite cluster 2 | 0.181 | 0.351 | 0.000 | 0.246 | 0.094 | 0.000 |
| Bacteroidetes | Bacteroidia | Bacteroidales | Porphyromonadaceae 2 | Termite cluster 2 | 0.000 | 0.000 | 0.000 | 0.000 | 0.000 | 0.142 |

**Table S2 continued.**

| | | | | | | | | | | |
|---|---|---|---|---|---|---|---|---|---|---|
| Bacteroidetes | Bacteroidia | Bacteroidales | Porphyromonadaceae 2 | Termite cluster 3 | 1.657 | 1.308 | 0.000 | 0.759 | 0.667 | 0.418 |
| Bacteroidetes | Bacteroidia | Bacteroidales | Porphyromonadaceae 2 | Unclassified | 0.000 | 0.000 | 0.013 | 0.000 | 0.000 | 0.002 |
| Bacteroidetes | Bacteroidia | Bacteroidales | Porphyromonadaceae 3 | Cluster IV | 0.070 | 0.064 | 0.000 | 0.114 | 0.207 | 0.163 |
| Bacteroidetes | Bacteroidia | Bacteroidales | Porphyromonadaceae 4 | *Petrimonas* | 0.001 | 0.000 | 0.000 | 0.000 | 0.004 | 0.000 |
| Bacteroidetes | Bacteroidia | Bacteroidales | Porphyromonadaceae 5 | Termite Cluster | 0.001 | 0.000 | 0.000 | 0.000 | 0.000 | 0.000 |
| Bacteroidetes | Bacteroidia | Bacteroidales | Porphyromonadaceae 7 | *Barnesiella* | 0.051 | 0.064 | 0.000 | 0.032 | 0.012 | 0.006 |
| Bacteroidetes | Bacteroidia | Bacteroidales | Porphyromonadaceae Cluster V | *Candidatus* Armantifilum | 0.052 | 0.032 | 0.000 | 0.171 | 0.012 | 0.028 |
| Bacteroidetes | Bacteroidia | Bacteroidales | Porphyromonadaceae Cluster V | *Candidatus* Symbiothrix | 0.038 | 0.064 | 0.000 | 0.086 | 0.004 | 0.014 |
| Bacteroidetes | Bacteroidia | Bacteroidales | Porphyromonadaceae Cluster V | Cockroach cluster | 0.004 | 0.128 | 0.000 | 0.011 | 0.012 | 0.001 |
| Bacteroidetes | Bacteroidia | Bacteroidales | Porphyromonadaceae Cluster V | Mixed gut cluster | 0.000 | 0.000 | 0.000 | 0.000 | 0.138 | 0.116 |
| Bacteroidetes | Bacteroidia | Bacteroidales | Porphyromonadaceae Cluster V | Termite Cockroach cluster | 0.012 | 0.000 | 0.000 | 0.014 | 0.012 | 0.001 |
| Bacteroidetes | Bacteroidia | Bacteroidales | Porphyromonadaceae Gut group | Environmental cluster | 0.000 | 0.000 | 0.000 | 0.004 | 0.000 | 0.000 |
| Bacteroidetes | Bacteroidia | Bacteroidales | Porphyromonadaceae Gut group | Mammalian cluster | 0.014 | 0.000 | 0.000 | 0.014 | 0.016 | 0.006 |
| Bacteroidetes | Bacteroidia | Bacteroidales | Porphyromonadaceae Gut group | Mixed gut cluster | 0.000 | 0.000 | 0.000 | 0.000 | 0.008 | 0.000 |
| Bacteroidetes | Bacteroidia | Bacteroidales | Porphyromonadaceae Gut group | Termite cluster I | 0.192 | 0.542 | 0.000 | 0.445 | 0.106 | 0.052 |
| Bacteroidetes | Bacteroidia | Bacteroidales | Porphyromonadaceae Gut group | Termite cluster II | 0.719 | 0.510 | 0.013 | 1.229 | 1.452 | 0.392 |
| Bacteroidetes | Bacteroidia | Bacteroidales | Porphyromonadaceae Gut group | Unclassified | 0.000 | 0.000 | 0.000 | 0.000 | 0.000 | 0.002 |
| Bacteroidetes | Bacteroidia | Bacteroidales | Prevotellaceae | *Paraprevotella* | 0.001 | 0.000 | 0.000 | 0.004 | 0.000 | 0.000 |
| Bacteroidetes | Bacteroidia | Bacteroidales | Prevotellaceae | *Prevotella* 1 | 0.068 | 0.128 | 0.000 | 0.139 | 0.024 | 0.006 |
| Bacteroidetes | Bacteroidia | Bacteroidales | Prevotellaceae | *Prevotella* 2 | 0.001 | 0.000 | 0.000 | 0.000 | 0.000 | 0.000 |
| Bacteroidetes | Bacteroidia | Bacteroidales | Prevotellaceae | Uncultured 10 | 0.006 | 0.000 | 0.000 | 0.025 | 0.000 | 0.000 |
| Bacteroidetes | Bacteroidia | Bacteroidales | Prevotellaceae | Uncultured 7 | 0.004 | 0.000 | 0.000 | 0.004 | 0.000 | 0.000 |
| Bacteroidetes | Bacteroidia | Bacteroidales | Prevotellaceae | Uncultured 8 | 0.005 | 0.035 | 0.000 | 0.000 | 0.000 | 0.000 |

**Table S2 continued.**

| | | | | | | | | | | |
|---|---|---|---|---|---|---|---|---|---|---|
| Bacteroidetes | Bacteroidia | Bacteroidales | RF16 | Unclassified | 0.002 | 0.032 | 0.000 | 0.007 | 0.004 | 0.002 |
| Bacteroidetes | Bacteroidia | Bacteroidales | Rikenellaceae | *Alisitpes* IV | 4.318 | 2.520 | 0.039 | 6.809 | 2.501 | 1.383 |
| Bacteroidetes | Bacteroidia | Bacteroidales | Rikenellaceae | *Alistipes* | 0.014 | 0.000 | 0.000 | 0.032 | 0.077 | 0.033 |
| Bacteroidetes | Bacteroidia | Bacteroidales | Rikenellaceae | *Alistipes* I | 0.067 | 0.096 | 0.000 | 0.175 | 0.268 | 0.126 |
| Bacteroidetes | Bacteroidia | Bacteroidales | Rikenellaceae | *Alistipes* II | 6.981 | 6.571 | 0.064 | 9.514 | 7.211 | 5.654 |
| Bacteroidetes | Bacteroidia | Bacteroidales | Rikenellaceae | *Alistipes* III | 0.023 | 0.510 | 0.026 | 0.242 | 16.788 | 11.507 |
| Bacteroidetes | Bacteroidia | Bacteroidales | Rikenellaceae | BCf9-17 termite group | 0.720 | 1.116 | 0.013 | 0.777 | 0.061 | 0.014 |
| Bacteroidetes | Bacteroidia | Bacteroidales | Rikenellaceae | dgA-11 gut group | 0.009 | 0.000 | 0.000 | 0.000 | 0.004 | 0.002 |
| Bacteroidetes | Bacteroidia | Bacteroidales | Rikenellaceae | gut cluster c | 0.088 | 0.096 | 0.000 | 0.207 | 0.085 | 0.028 |
| Bacteroidetes | Bacteroidia | Bacteroidales | Rikenellaceae | hoa5-07d05 gut group | 0.000 | 0.000 | 0.000 | 0.000 | 0.004 | 0.000 |
| Bacteroidetes | Bacteroidia | Bacteroidales | Rikenellaceae | M2PB4-61 termite group | 1.626 | 3.668 | 0.026 | 0.894 | 3.400 | 3.708 |
| Bacteroidetes | Bacteroidia | Bacteroidales | Rikenellaceae | *Millionella* | 0.000 | 0.000 | 0.000 | 0.000 | 0.000 | 0.001 |
| Bacteroidetes | Bacteroidia | Bacteroidales | Rikenellaceae | RC9 gut group | 0.171 | 0.223 | 0.013 | 0.146 | 0.187 | 0.110 |
| Bacteroidetes | Bacteroidia | Bacteroidales | Rikenellaceae | *Rikenella* | 0.019 | 0.064 | 0.000 | 0.021 | 0.004 | 0.000 |
| Bacteroidetes | Bacteroidia | Bacteroidales | Rikenellaceae | *Rikenella* 1 | 0.001 | 0.000 | 0.000 | 0.004 | 0.000 | 0.000 |
| Bacteroidetes | Bacteroidia | Bacteroidales | Rikenellaceae | *Rikenella* 3 | 0.000 | 0.000 | 0.000 | 0.000 | 0.004 | 0.001 |
| Bacteroidetes | Bacteroidia | Bacteroidales | Rikenellaceae | Rs-D38 termite group | 0.553 | 1.148 | 0.000 | 0.143 | 0.171 | 0.303 |
| Bacteroidetes | Bacteroidia | Bacteroidales | Rikenellaceae | SP3-e08 | 0.000 | 0.000 | 0.000 | 0.004 | 0.000 | 0.000 |
| Bacteroidetes | Bacteroidia | Bacteroidales | Rikenellaceae | *Tidjanibacter* | 0.004 | 0.000 | 0.000 | 0.007 | 0.045 | 0.025 |
| Bacteroidetes | Bacteroidia | Bacteroidales | Rikenellaceae | Unclassified | 0.000 | 0.000 | 0.000 | 0.007 | 0.012 | 0.000 |
| Bacteroidetes | Bacteroidia | Bacteroidales | Rikenellaceae | Unclassified | 0.004 | 0.000 | 0.000 | 0.000 | 0.000 | 0.000 |
| Bacteroidetes | Bacteroidia | Bacteroidales | Rikenellaceae | vadinBC27 wastewater-sludge group | 0.012 | 0.066 | 0.000 | 0.011 | 0.045 | 0.039 |
| Bacteroidetes | Bacteroidia | Bacteroidales | S24 7 | Cluster I | 0.070 | 0.096 | 0.077 | 0.061 | 0.028 | 0.017 |
| Bacteroidetes | Bacteroidia | Bacteroidales | S24 7 | Cluster II | 0.046 | 0.096 | 0.000 | 0.043 | 0.000 | 0.001 |
| Bacteroidetes | Bacteroidia | Bacteroidales | Tannerellaceae | *Parabacteroides* | 0.189 | 0.319 | 0.000 | 0.160 | 0.041 | 0.038 |
| Bacteroidetes | Bacteroidia | Bacteroidales | Tannerellaceae | *Tannerella* | 3.169 | 2.488 | 0.013 | 3.485 | 0.789 | 0.971 |
| Bacteroidetes | Bacteroidia | Bacteroidales | Unclassified | Unclassified | 0.000 | 0.000 | 0.000 | 0.000 | 0.000 | 0.001 |

**Table S2 continued.**

| | | | | | | | | | | |
|---|---|---|---|---|---|---|---|---|---|---|
| Bacteroidetes | Bacteroidia | Marinilabiliales | Marinilabiliaceae | *Alkalitalea* | 0.010 | 0.000 | 0.000 | 0.004 | 0.000 | 0.002 |
| Bacteroidetes | Bacteroidia | Marinilabiliales | Marinilabiliaceae | *Carboxylicivirga* | 0.001 | 0.000 | 0.000 | 0.000 | 0.000 | 0.000 |
| Bacteroidetes | Bacteroidia | Marinilabiliales | Marinilabiliaceae | *Mangroviflexus* | 0.000 | 0.000 | 0.000 | 0.004 | 0.008 | 0.001 |
| Bacteroidetes | Bacteroidia | Marinilabiliales | Marinilabiliaceae | *Marinilabilia* | 0.001 | 0.000 | 0.000 | 0.000 | 0.000 | 0.000 |
| Bacteroidetes | Bacteroidia | Marinilabiliales | Marinilabiliaceae | *Natronoflexus* | 0.021 | 0.000 | 0.000 | 0.011 | 0.004 | 0.003 |
| Bacteroidetes | Bacteroidia | Marinilabiliales | Marinilabiliaceae | *Saccharicrinis* | 0.000 | 0.000 | 0.000 | 0.000 | 0.004 | 0.000 |
| Bacteroidetes | Bacteroidia | Marinilabiliales | Prolixibacteraceae | *Draconibacterium* | 0.000 | 0.000 | 0.000 | 0.004 | 0.004 | 0.000 |
| Bacteroidetes | Bacteroidia | Marinilabiliales | Prolixibacteraceae | *Mangrovibacterium* | 0.005 | 0.000 | 0.000 | 0.000 | 0.000 | 0.000 |
| Bacteroidetes | Bacteroidia | Marinilabiliales | Prolixibacteraceae | *Mariniphaga* | 0.017 | 0.000 | 0.000 | 0.039 | 0.004 | 0.000 |
| Bacteroidetes | Bacteroidia | Marinilabiliales | Prolixibacteraceae | *Meniscus* | 0.004 | 0.000 | 0.000 | 0.007 | 0.000 | 0.000 |
| Bacteroidetes | Bacteroidia | Marinilabiliales | Prolixibacteraceae | *Prolixibacter* | 0.005 | 0.000 | 0.000 | 0.000 | 0.004 | 0.000 |
| Bacteroidetes | Bacteroidia | Marinilabiliales | Prolixibacteraceae | *Tangfeifania* | 0.000 | 0.000 | 0.000 | 0.004 | 0.000 | 0.001 |
| Bacteroidetes | Chitinophagia | Chitinophagales | Chitinophagaceae | *Asinibacterium* | 0.000 | 0.000 | 0.013 | 0.043 | 0.000 | 0.000 |
| Bacteroidetes | Chitinophagia | Chitinophagales | Chitinophagaceae | *Chitinophaga* | 0.002 | 0.000 | 0.000 | 0.118 | 0.000 | 0.000 |
| Bacteroidetes | Chitinophagia | Chitinophagales | Chitinophagaceae | *Ferruginibacter* | 0.001 | 0.000 | 0.000 | 0.000 | 0.000 | 0.000 |
| Bacteroidetes | Chitinophagia | Chitinophagales | Chitinophagaceae | *Flavihumibacter* | 0.000 | 0.000 | 0.000 | 0.011 | 0.000 | 0.000 |
| Bacteroidetes | Chitinophagia | Chitinophagales | Chitinophagaceae | *Flavitalea* | 0.000 | 0.000 | 0.000 | 0.021 | 0.000 | 0.001 |
| Bacteroidetes | Chitinophagia | Chitinophagales | Chitinophagaceae | *Hydrobacter* | 0.000 | 0.000 | 0.000 | 0.007 | 0.004 | 0.000 |
| Bacteroidetes | Chitinophagia | Chitinophagales | Chitinophagaceae | *Hydrotalea* | 0.001 | 0.000 | 0.000 | 0.271 | 0.000 | 0.000 |
| Bacteroidetes | Chitinophagia | Chitinophagales | Chitinophagaceae | *Lacibacter* | 0.000 | 0.000 | 0.000 | 0.007 | 0.000 | 0.000 |
| Bacteroidetes | Chitinophagia | Chitinophagales | Chitinophagaceae | *Niabella* | 0.000 | 0.000 | 0.000 | 0.036 | 0.000 | 0.000 |
| Bacteroidetes | Chitinophagia | Chitinophagales | Chitinophagaceae | *Niastella* | 0.000 | 0.000 | 0.000 | 0.011 | 0.000 | 0.000 |
| Bacteroidetes | Chitinophagia | Chitinophagales | Chitinophagaceae | *Panacibacter* | 0.000 | 0.000 | 0.000 | 0.004 | 0.000 | 0.000 |
| Bacteroidetes | Chitinophagia | Chitinophagales | Chitinophagaceae | *Parafilimonas* | 0.000 | 0.000 | 0.000 | 0.004 | 0.000 | 0.000 |
| Bacteroidetes | Chitinophagia | Chitinophagales | Chitinophagaceae | *Parasegetibacter* | 0.001 | 0.000 | 0.000 | 0.000 | 0.000 | 0.000 |
| Bacteroidetes | Chitinophagia | Chitinophagales | Chitinophagaceae | *Pseudoflavitalea* | 0.000 | 0.000 | 0.000 | 0.004 | 0.000 | 0.000 |
| Bacteroidetes | Chitinophagia | Chitinophagales | Chitinophagaceae | *Sediminibacterium* | 0.002 | 0.000 | 0.039 | 0.061 | 0.000 | 0.002 |

**Table S2 continued.**

| | | | | | | | | | | |
|---|---|---|---|---|---|---|---|---|---|---|
| Bacteroidetes | Chitinophagia | Chitinophagales | Chitinophagaceae | *Taibaiella* | 0.001 | 0.000 | 0.000 | 0.011 | 0.000 | 0.000 |
| Bacteroidetes | Chitinophagia | Chitinophagales | Chitinophagaceae | *Terrimonas* | 0.000 | 0.000 | 0.000 | 0.007 | 0.000 | 0.000 |
| Bacteroidetes | Chitinophagia | Chitinophagales | Chitinophagaceae | *Vibrionimonas* | 0.002 | 0.000 | 0.013 | 0.056 | 0.000 | 0.000 |
| Bacteroidetes | Cytophagia | Cytophagales | Cyclobacteriaceae | *Algoriphagus* | 0.000 | 0.000 | 0.000 | 0.018 | 0.000 | 0.000 |
| Bacteroidetes | Cytophagia | Cytophagales | Cyclobacteriaceae | *Cyclobacterium* | 0.001 | 0.000 | 0.000 | 0.000 | 0.004 | 0.000 |
| Bacteroidetes | Cytophagia | Cytophagales | Cytophagaceae | *Cytophaga* | 0.004 | 0.000 | 0.000 | 0.000 | 0.000 | 0.000 |
| Bacteroidetes | Cytophagia | Cytophagales | Cytophagaceae | *Dyadobacter* | 0.000 | 0.000 | 0.000 | 0.135 | 0.000 | 0.000 |
| Bacteroidetes | Cytophagia | Cytophagales | Cytophagaceae | *Larkinella* | 0.000 | 0.000 | 0.000 | 0.018 | 0.000 | 0.000 |
| Bacteroidetes | Cytophagia | Cytophagales | Cytophagaceae | *Ohtaekwangia* | 0.001 | 0.000 | 0.000 | 0.064 | 0.000 | 0.000 |
| Bacteroidetes | Cytophagia | Cytophagales | Cytophagaceae | *Siphonobacter* | 0.000 | 0.000 | 0.000 | 0.004 | 0.000 | 0.000 |
| Bacteroidetes | Cytophagia | Cytophagales | Cytophagaceae | *Spirosoma* | 0.002 | 0.000 | 0.000 | 0.018 | 0.000 | 0.000 |
| Bacteroidetes | Cytophagia | Cytophagales | Hymenobacteraceae | *Adhaeribacter* | 0.001 | 0.000 | 0.000 | 0.061 | 0.000 | 0.000 |
| Bacteroidetes | Cytophagia | Cytophagales | Hymenobacteraceae | *Hymenobacter* | 0.001 | 0.000 | 0.000 | 0.004 | 0.004 | 0.002 |
| Bacteroidetes | Cytophagia | Cytophagales | Hymenobacteraceae | *Pontibacter* | 0.004 | 0.000 | 0.000 | 0.032 | 0.000 | 0.000 |
| Bacteroidetes | Cytophagia | Cytophagales | Hymenobacteraceae | *Rufibacter* | 0.000 | 0.000 | 0.000 | 0.004 | 0.000 | 0.000 |
| Bacteroidetes | Cytophagia | Cytophagales | Persicobacteraceae | *Fulvitalea* | 0.009 | 0.000 | 0.000 | 0.014 | 0.000 | 0.000 |
| Bacteroidetes | Cytophagia | Cytophagales | Unclassified | *Chryseolinea* | 0.000 | 0.000 | 0.000 | 0.004 | 0.000 | 0.000 |
| Bacteroidetes | Flavobacteria | Flavobacteriales | Blattabacteriaceae | Armored scale insect endosymbionts | 0.001 | 0.000 | 0.000 | 0.000 | 0.000 | 0.001 |
| Bacteroidetes | Flavobacteria | Flavobacteriales | Blattabacteriaceae | *Blattabacterium* | 0.000 | 0.000 | 0.000 | 0.004 | 0.004 | 0.000 |
| Bacteroidetes | Flavobacteria | Flavobacteriales | Blattabacteriaceae | *Candidatus Sulcia* | 0.000 | 0.000 | 0.000 | 0.004 | 0.000 | 0.000 |
| Bacteroidetes | Flavobacteria | Flavobacteriales | Cryomorphaceae | *Owenweeksia* | 0.000 | 0.000 | 0.000 | 0.004 | 0.000 | 0.000 |
| Bacteroidetes | Flavobacteria | Flavobacteriales | Cryomorphaceae 2 | *Crocinitomix* | 0.000 | 0.000 | 0.000 | 0.004 | 0.000 | 0.000 |
| Bacteroidetes | Flavobacteria | Flavobacteriales | Cryomorphaceae 2 | Unclassified | 0.000 | 0.000 | 0.000 | 0.000 | 0.004 | 0.000 |
| Bacteroidetes | Flavobacteria | Flavobacteriales | Flavobacteriaceae | *Capnocytophaga* | 0.007 | 0.000 | 0.000 | 0.004 | 0.012 | 0.002 |
| Bacteroidetes | Flavobacteria | Flavobacteriales | Flavobacteriaceae 1 | *Costertonia* | 0.000 | 0.000 | 0.000 | 0.000 | 0.004 | 0.000 |
| Bacteroidetes | Flavobacteria | Flavobacteriales | Flavobacteriaceae 1 | *Croceibacter* | 0.000 | 0.000 | 0.000 | 0.000 | 0.000 | 0.002 |
| Bacteroidetes | Flavobacteria | Flavobacteriales | Flavobacteriaceae 1 | *Flavobacterium* 1 | 0.002 | 0.000 | 0.000 | 0.004 | 0.020 | 0.001 |

**Table S2 continued.**

| | | | | | | | | | | |
|---|---|---|---|---|---|---|---|---|---|---|
| Bacteroidetes | Flavobacteria | Flavobacteriales | Flavobacteriaceae 1 | *Flavobacterium* 2 | 0.001 | 0.000 | 0.000 | 0.000 | 0.000 | 0.001 |
| Bacteroidetes | Flavobacteria | Flavobacteriales | Flavobacteriaceae 1 | *Mesonia* | 0.000 | 0.000 | 0.000 | 0.000 | 0.004 | 0.000 |
| Bacteroidetes | Flavobacteria | Flavobacteriales | Flavobacteriaceae 1 | *Pibocella* | 0.000 | 0.000 | 0.000 | 0.000 | 0.004 | 0.001 |
| Bacteroidetes | Flavobacteria | Flavobacteriales | Flavobacteriaceae 1 | *Salinimicrobium* | 0.001 | 0.000 | 0.000 | 0.000 | 0.000 | 0.000 |
| Bacteroidetes | Flavobacteria | Flavobacteriales | Flavobacteriaceae 1 | *Tenacibaculum* 2 | 0.001 | 0.000 | 0.000 | 0.000 | 0.000 | 0.000 |
| Bacteroidetes | Flavobacteria | Flavobacteriales | Flavobacteriaceae 1 | *Tenacibaculum* 3 | 0.001 | 0.000 | 0.000 | 0.000 | 0.000 | 0.000 |
| Bacteroidetes | Flavobacteria | Flavobacteriales | Flavobacteriaceae 1 | Unclassified | 0.000 | 0.000 | 0.000 | 0.000 | 0.000 | 0.002 |
| Bacteroidetes | Flavobacteria | Flavobacteriales | Flavobacteriaceae 1 | Uncultured 3 | 0.000 | 0.000 | 0.000 | 0.000 | 0.004 | 0.000 |
| Bacteroidetes | Flavobacteria | Flavobacteriales | Flavobacteriaceae 2 | Unclassified | 0.000 | 0.000 | 0.000 | 0.004 | 0.000 | 0.000 |
| Bacteroidetes | Flavobacteria | Flavobacteriales | Flavobacteriaceae 2 | Uncultured a | 0.001 | 0.000 | 0.000 | 0.000 | 0.000 | 0.000 |
| Bacteroidetes | Flavobacteria | Flavobacteriales | Flavobacteriaceae 2 | Unclassified | 0.002 | 0.000 | 0.000 | 0.000 | 0.000 | 0.000 |
| Bacteroidetes | Flavobacteria | Flavobacteriales | NS9 marine group | Unclassified | 0.001 | 0.000 | 0.000 | 0.000 | 0.000 | 0.000 |
| Bacteroidetes | Flavobacteria | Flavobacteriales | Unclassified | Unclassified | 0.000 | 0.000 | 0.000 | 0.004 | 0.000 | 0.000 |
| Bacteroidetes | Flavobacteriia | Flavobacteriales | Crocinitomicaceae | *Fluviicola* | 0.000 | 0.000 | 0.007 | 0.007 | 0.008 | 0.000 |
| Bacteroidetes | Flavobacteriia | Flavobacteriales | Flavabacteriaceae | *Spongiimicrobium* | 0.000 | 0.000 | 0.000 | 0.000 | 0.004 | 0.000 |
| Bacteroidetes | Flavobacteriia | Flavobacteriales | Flavabacteriaceae | *Zobellia* | 0.000 | 0.000 | 0.000 | 0.004 | 0.008 | 0.000 |
| Bacteroidetes | Flavobacteriia | Flavobacteriales | Flavobacteriaceae | *Aquimarina* | 0.000 | 0.000 | 0.000 | 0.000 | 0.000 | 0.001 |
| Bacteroidetes | Flavobacteriia | Flavobacteriales | Flavobacteriaceae | *Arenibacter* | 0.000 | 0.000 | 0.000 | 0.000 | 0.000 | 0.001 |
| Bacteroidetes | Flavobacteriia | Flavobacteriales | Flavobacteriaceae | *Cellulophaga* | 0.000 | 0.000 | 0.000 | 0.000 | 0.004 | 0.002 |
| Bacteroidetes | Flavobacteriia | Flavobacteriales | Flavobacteriaceae | *Cloacibacterium* | 0.007 | 0.000 | 0.000 | 0.082 | 0.000 | 0.001 |
| Bacteroidetes | Flavobacteriia | Flavobacteriales | Flavobacteriaceae | *Elizabethkingia* | 0.001 | 0.000 | 0.000 | 0.000 | 0.000 | 0.000 |
| Bacteroidetes | Flavobacteriia | Flavobacteriales | Flavobacteriaceae | *Eudoraea* | 0.000 | 0.000 | 0.000 | 0.000 | 0.004 | 0.000 |
| Bacteroidetes | Flavobacteriia | Flavobacteriales | Flavobacteriaceae | *Flavobacterium* | 0.001 | 0.000 | 0.000 | 0.011 | 0.004 | 0.001 |
| Bacteroidetes | Flavobacteriia | Flavobacteriales | Flavobacteriaceae | *Kordia* | 0.001 | 0.000 | 0.000 | 0.000 | 0.000 | 0.000 |
| Bacteroidetes | Flavobacteriia | Flavobacteriales | Flavobacteriaceae | *Leptobacterium* | 0.000 | 0.000 | 0.000 | 0.000 | 0.004 | 0.000 |
| Bacteroidetes | Flavobacteriia | Flavobacteriales | Flavobacteriaceae | *Lutaonella* | 0.000 | 0.000 | 0.000 | 0.004 | 0.000 | 0.001 |
| Bacteroidetes | Flavobacteriia | Flavobacteriales | Flavobacteriaceae | *Moheibacter* | 0.000 | 0.000 | 0.000 | 0.007 | 0.000 | 0.000 |

**Table S2 continued.**

| | | | | | | | | | | |
|---|---|---|---|---|---|---|---|---|---|---|
| Bacteroidetes | Flavobacteriia | Flavobacteriales | Flavobacteriaceae | *Muricauda* | 0.000 | 0.000 | 0.000 | 0.000 | 0.000 | 0.002 |
| Bacteroidetes | Flavobacteriia | Flavobacteriales | Flavobacteriaceae | *Myroides* | 0.000 | 0.000 | 0.013 | 0.000 | 0.000 | 0.002 |
| Bacteroidetes | Flavobacteriia | Flavobacteriales | Flavobacteriaceae | *Nonlabens* | 0.001 | 0.000 | 0.000 | 0.000 | 0.000 | 0.000 |
| Bacteroidetes | Flavobacteriia | Flavobacteriales | Flavobacteriaceae | *Psychroserpens* | 0.000 | 0.000 | 0.000 | 0.000 | 0.004 | 0.000 |
| Bacteroidetes | Flavobacteriia | Flavobacteriales | Flavobacteriaceae | *Robiginitalea* | 0.001 | 0.000 | 0.000 | 0.000 | 0.000 | 0.000 |
| Bacteroidetes | Flavobacteriia | Flavobacteriales | Flavobacteriaceae | *Salegentibacter* | 0.002 | 0.000 | 0.000 | 0.000 | 0.004 | 0.001 |
| Bacteroidetes | Flavobacteriia | Flavobacteriales | Flavobacteriaceae | *Tenacibaculum* | 0.001 | 0.000 | 0.000 | 0.000 | 0.000 | 0.000 |
| Bacteroidetes | Flavobacteriia | Flavobacteriales | Flavobacteriaceae | *Weeksella* | 0.001 | 0.000 | 0.000 | 0.000 | 0.000 | 0.000 |
| Bacteroidetes | SM1A07 | Unclassified | Unclassified | Unclassified | 0.000 | 0.000 | 0.000 | 0.000 | 0.000 | 0.001 |
| Bacteroidetes | Sphingobacteria | Sphingobacteriales 1 | BD2 2 | Unclassified | 0.006 | 0.032 | 0.000 | 0.000 | 0.004 | 0.001 |
| Bacteroidetes | Sphingobacteria | Sphingobacteriales 1 | BSV13 | Unclassified | 0.002 | 0.000 | 0.000 | 0.004 | 0.000 | 0.000 |
| Bacteroidetes | Sphingobacteria | Sphingobacteriales 1 | env OPS 17 | Unclassified | 0.000 | 0.000 | 0.000 | 0.021 | 0.000 | 0.001 |
| Bacteroidetes | Sphingobacteria | Sphingobacteriales 1 | KD1-131 | Unclassified | 0.000 | 0.000 | 0.000 | 0.000 | 0.000 | 0.001 |
| Bacteroidetes | Sphingobacteria | Sphingobacteriales 1 | NS11-12 marine group | Unclassified | 0.000 | 0.000 | 0.000 | 0.011 | 0.000 | 0.000 |
| Bacteroidetes | Sphingobacteria | Sphingobacteriales 1 | PHOS-HE51 | Unclassified | 0.002 | 0.000 | 0.000 | 0.007 | 0.000 | 0.000 |
| Bacteroidetes | Sphingobacteria | Sphingobacteriales 1 | SB 1 | Termite cluster | 1.719 | 1.563 | 0.026 | 2.405 | 0.484 | 0.250 |
| Bacteroidetes | Sphingobacteria | Sphingobacteriales 1 | SB 1 | Unclassified | 0.000 | 0.577 | 0.000 | 0.453 | 0.004 | 0.000 |
| Bacteroidetes | Sphingobacteria | Sphingobacteriales 1 | SB 1 | Unclassified | 0.503 | 0.000 | 0.000 | 0.000 | 0.000 | 0.000 |
| Bacteroidetes | Sphingobacteria | Sphingobacteriales 1 | SB 5 | Unclassified | 0.000 | 0.000 | 0.000 | 0.004 | 0.000 | 0.000 |
| Bacteroidetes | Sphingobacteria | Sphingobacteriales 1 | Sphingobacteriaceae | *Mucilaginibacter* 1 | 0.000 | 0.000 | 0.000 | 0.007 | 0.012 | 0.000 |
| Bacteroidetes | Sphingobacteria | Sphingobacteriales 1 | Sphingobacteriaceae | *Mucilaginibacter* 2 | 0.001 | 0.000 | 0.000 | 0.000 | 0.041 | 0.008 |
| Bacteroidetes | Sphingobacteria | Sphingobacteriales 1 | Sphingobacteriaceae | *Pedobacter* 1 | 0.000 | 0.000 | 0.000 | 0.000 | 0.004 | 0.000 |
| Bacteroidetes | Sphingobacteria | Sphingobacteriales 1 | Sphingobacteriaceae | *Pedobacter* 2 | 0.000 | 0.032 | 0.000 | 0.007 | 0.000 | 0.000 |
| Bacteroidetes | Sphingobacteria | Sphingobacteriales 1 | Sphingobacteriaceae | *Pedobacter* 5 | 0.000 | 0.000 | 0.000 | 0.004 | 0.000 | 0.001 |
| Bacteroidetes | Sphingobacteria | Sphingobacteriales 1 | Sphingobacteriaceae | *Sphingobacterium* 1 | 0.000 | 0.000 | 0.000 | 0.004 | 0.000 | 0.000 |
| Bacteroidetes | Sphingobacteria | Sphingobacteriales 1 | Sphingobacteriaceae | *Sphingobacterium* 2 | 0.000 | 0.000 | 0.000 | 0.004 | 0.000 | 0.001 |
| Bacteroidetes | Sphingobacteria | Sphingobacteriales 1 | Sphingobacteriaceae | *Sphingobacterium* 3 | 0.000 | 0.000 | 0.000 | 0.007 | 0.004 | 0.000 |

**Table S2 continued.**

| | | | | | | | | | | |
|---|---|---|---|---|---|---|---|---|---|---|
| Bacteroidetes | Sphingobacteria | Sphingobacteriales 1 | Sphingobacteriaceae | *Sphingobacterium* 4 | 0.000 | 0.000 | 0.000 | 0.000 | 0.004 | 0.000 |
| Bacteroidetes | Sphingobacteria | Sphingobacteriales 1 | Sphingobacteriaceae | Unclassified | 0.000 | 0.000 | 0.000 | 0.021 | 0.004 | 0.002 |
| Bacteroidetes | Sphingobacteria | Sphingobacteriales 1 | ST-12K33 | Unclassified | 0.001 | 0.000 | 0.000 | 0.004 | 0.000 | 0.000 |
| Bacteroidetes | Sphingobacteria | Sphingobacteriales 1 | Unclassified | Unclassified | 0.000 | 0.000 | 0.000 | 0.014 | 0.000 | 0.002 |
| Bacteroidetes | Sphingobacteria | Sphingobacteriales 1 | vadinHA17 | Unclassified | 0.037 | 0.032 | 0.000 | 0.043 | 0.037 | 0.018 |
| Bacteroidetes | Sphingobacteria | Sphingobacteriales 1 | WCHB1-32 | Unclassified | 0.000 | 0.032 | 0.000 | 0.000 | 0.004 | 0.000 |
| Bacteroidetes | Sphingobacteria | Sphingobacteriales 1 | WCHB1-69 | Unclassified | 0.000 | 0.000 | 0.000 | 0.000 | 0.004 | 0.003 |
| Bacteroidetes | Sphingobacteria | Sphingobacteriales 2 | Chitinophagaceae | Unclassified | 0.000 | 0.000 | 0.000 | 0.029 | 0.000 | 0.005 |
| Bacteroidetes | Sphingobacteria | Sphingobacteriales 2 | Chitinophagaceae | Uncultured 1 | 0.000 | 0.000 | 0.000 | 0.011 | 0.000 | 0.000 |
| Bacteroidetes | Sphingobacteria | Sphingobacteriales 2 | Chitinophagaceae | Uncultured 10 | 0.002 | 0.000 | 0.000 | 0.029 | 0.000 | 0.000 |
| Bacteroidetes | Sphingobacteria | Sphingobacteriales 2 | Chitinophagaceae | Uncultured 11 | 0.002 | 0.000 | 0.000 | 0.039 | 0.000 | 0.000 |
| Bacteroidetes | Sphingobacteria | Sphingobacteriales 2 | Chitinophagaceae | Uncultured 12 | 0.000 | 0.000 | 0.000 | 0.007 | 0.000 | 0.000 |
| Bacteroidetes | Sphingobacteria | Sphingobacteriales 2 | Chitinophagaceae | Uncultured 2 | 0.002 | 0.000 | 0.000 | 0.043 | 0.000 | 0.000 |
| Bacteroidetes | Sphingobacteria | Sphingobacteriales 2 | Chitinophagaceae | Uncultured 4 | 0.000 | 0.000 | 0.000 | 0.007 | 0.000 | 0.000 |
| Bacteroidetes | Sphingobacteria | Sphingobacteriales 2 | Chitinophagaceae | Uncultured 5 | 0.002 | 0.000 | 0.000 | 0.021 | 0.000 | 0.011 |
| Bacteroidetes | Sphingobacteria | Sphingobacteriales 2 | Chitinophagaceae | Uncultured 6 | 0.000 | 0.000 | 0.000 | 0.007 | 0.000 | 0.000 |
| Bacteroidetes | Sphingobacteria | Sphingobacteriales 2 | Chitinophagaceae | Uncultured 9 | 0.000 | 0.000 | 0.000 | 0.004 | 0.000 | 0.000 |
| Bacteroidetes | Sphingobacteria | Sphingobacteriales 2 | Saprospiraceae | *Lewinella* | 0.000 | 0.000 | 0.000 | 0.000 | 0.000 | 0.001 |
| Bacteroidetes | Sphingobacteria | Sphingobacteriales 2 | Saprospiraceae | Unclassified | 0.000 | 0.000 | 0.000 | 0.004 | 0.000 | 0.000 |
| Bacteroidetes | Sphingobacteria | Sphingobacteriales 2 | Saprospiraceae | Uncultured 1 | 0.000 | 0.000 | 0.000 | 0.000 | 0.000 | 0.001 |
| Bacteroidetes | Sphingobacteria | Sphingobacteriales 2 | Saprospiraceae | Uncultured 2 | 0.001 | 0.000 | 0.000 | 0.011 | 0.000 | 0.000 |
| Bacteroidetes | Sphingobacteria | Sphingobacteriales 3 | Cyclobacteriaceae | Uncultured 1 | 0.000 | 0.000 | 0.000 | 0.004 | 0.000 | 0.000 |
| Bacteroidetes | Sphingobacteria | Sphingobacteriales 3 | Cytophagaceae 1 | *Flexibacter* 1 | 0.001 | 0.000 | 0.000 | 0.175 | 0.000 | 0.000 |
| Bacteroidetes | Sphingobacteria | Sphingobacteriales 3 | Cytophagaceae 3 | *Pontibacter* 1 | 0.000 | 0.000 | 0.000 | 0.000 | 0.000 | 0.001 |
| Bacteroidetes | Sphingobacteria | Sphingobacteriales 3 | Flammeovirgaceae 1 | Unclassified | 0.002 | 0.000 | 0.000 | 0.000 | 0.000 | 0.000 |
| Bacteroidetes | Sphingobacteria | Sphingobacteriales 3 | Flammeovirgaceae 2 | *Candidatus* Cardinium | 0.001 | 0.000 | 0.000 | 0.004 | 0.004 | 0.003 |
| Bacteroidetes | Sphingobacteria | Sphingobacteriales 4 | Rhodothermaceae | Unclassified | 0.000 | 0.000 | 0.000 | 0.004 | 0.000 | 0.000 |

**Table S2 continued.**

| | | | | | | | | | | |
|---|---|---|---|---|---|---|---|---|---|---|
| Bacteroidetes | Sphingobacteria | Sphingobacteriales 4 | Rhodothermaceae | Uncultured 2 | 0.000 | 0.000 | 0.000 | 0.004 | 0.000 | 0.000 |
| Bacteroidetes | Sphingobacteria | Sphingobacteriales 4 | Rhodothermaceae | Uncultured 3 | 0.000 | 0.000 | 0.000 | 0.007 | 0.000 | 0.000 |
| Bacteroidetes | Sphingobacteriia | Sphingobacteriales | Sphingobacteriaceae | *Arcticibacter* | 0.001 | 0.000 | 0.000 | 0.000 | 0.000 | 0.000 |
| Bacteroidetes | Sphingobacteriia | Sphingobacteriales | Sphingobacteriaceae | *Mucilaginibacter* | 0.005 | 0.000 | 0.000 | 0.029 | 0.041 | 0.014 |
| Bacteroidetes | Sphingobacteriia | Sphingobacteriales | Sphingobacteriaceae | *Parapedobacter* | 0.000 | 0.000 | 0.000 | 0.000 | 0.004 | 0.008 |
| Bacteroidetes | Sphingobacteriia | Sphingobacteriales | Sphingobacteriaceae | *Pedobacter* | 0.001 | 0.000 | 0.000 | 0.014 | 0.004 | 0.005 |
| Bacteroidetes | Sphingobacteriia | Sphingobacteriales | Sphingobacteriaceae | *Solitalea* | 0.000 | 0.000 | 0.000 | 0.011 | 0.016 | 0.002 |
| Bacteroidetes | Sphingobacteriia | Sphingobacteriales | Sphingobacteriaceae | *Sphingobacterium* | 0.004 | 0.064 | 0.000 | 0.050 | 0.004 | 0.005 |
| Bacteroidetes | Sphingobacteriia | Sphingobacteriales | Sphingomonadaceae | *Nubsella* | 0.000 | 0.000 | 0.000 | 0.004 | 0.000 | 0.000 |
| Bacteroidetes | VC2 1 Bac22 | Unclassified | Unclassified | Unclassified | 0.010 | 0.000 | 0.000 | 0.004 | 0.024 | 0.008 |
| Balneolaeota | Balneolia | Balneolales | Balneolaceae | *Aliifodinibius* | 0.000 | 0.000 | 0.000 | 0.004 | 0.000 | 0.000 |
| BHI80-139 | Unclassified | Unclassified | Unclassified | Unclassified | 0.000 | 0.000 | 0.000 | 0.000 | 0.004 | 0.000 |
| Caldiserica | Caldisericia | Caldisericales | TTA-B1 | Unclassified | 0.000 | 0.000 | 0.000 | 0.004 | 0.000 | 0.000 |
| Candidate phylum BD1 5 | Unclassified | Unclassified | Unclassified | Unclassified | 0.025 | 0.064 | 0.000 | 0.004 | 0.004 | 0.000 |
| Candidate phylum BRC1 | Unclassified | Unclassified | Unclassified | Unclassified | 0.001 | 0.000 | 0.013 | 0.000 | 0.000 | 0.001 |
| Candidate phylum OD1 | Unclassified | Unclassified | Unclassified | Unclassified | 0.000 | 0.000 | 0.000 | 0.004 | 0.000 | 0.000 |
| Candidate phylum OP11 | Unclassified | Unclassified | Unclassified | Unclassified | 0.012 | 0.000 | 0.000 | 0.007 | 0.008 | 0.000 |
| Candidate phylum OP3 | Unclassified | Unclassified | Unclassified | Unclassified | 0.001 | 0.000 | 0.000 | 0.007 | 0.000 | 0.000 |
| Candidate phylum OP8 | Unclassified | Unclassified | Unclassified | Unclassified | 0.030 | 0.000 | 0.000 | 0.000 | 0.004 | 0.002 |
| Candidate phylum TG3 | Subphylum 1 | incertae sedis | Termite cluster III | Subcluster IIIa | 0.001 | 0.000 | 0.000 | 0.000 | 0.000 | 0.000 |
| Candidate phylum TG3 | Subphylum 2 | incertae sedis | Termite cluster IV | Subcluster IVa | 0.000 | 1.978 | 0.000 | 0.121 | 0.130 | 0.098 |
| Candidate phylum TM6 | Unclassified | Unclassified | Unclassified | Unclassified | 0.000 | 0.000 | 0.000 | 0.584 | 0.004 | 0.000 |
| Candidate phylum TM6 | Unclassified | Unclassified | Unclassified | Unclassified | 0.000 | 0.000 | 0.000 | 0.000 | 0.000 | 0.001 |

**Table S2 continued.**

| | | | | | | | | | | |
|---|---|---|---|---|---|---|---|---|---|---|
| Candidate phylum TM6 | Unclassified | Unclassified | Unclassified | Unclassified | 0.001 | 0.000 | 0.000 | 0.000 | 0.000 | 0.000 |
| Candidate phylum TM7 | Termite cluster 1 | Unclassified | Unclassified | Unclassified | 0.001 | 0.000 | 0.000 | 0.000 | 0.000 | 0.000 |
| Candidate phylum TM7 | Termite cluster 2 | Unclassified | Unclassified | Unclassified | 0.002 | 0.000 | 0.000 | 0.029 | 0.000 | 0.000 |
| Candidate phylum TM7 | Termite cockroach cluster | Unclassified | Unclassified | Unclassified | 0.160 | 0.287 | 0.013 | 0.403 | 0.085 | 0.041 |
| Candidate phylum TM7 | Unclassified | Unclassified | Unclassified | Unclassified | 0.066 | 0.000 | 0.013 | 0.392 | 0.024 | 0.039 |
| Candidate phylum WS3 | Unclassified | Unclassified | Unclassified | Unclassified | 0.002 | 0.000 | 0.000 | 0.032 | 0.000 | 0.000 |
| Candidate phylum WS6 | Unclassified | Unclassified | Unclassified | Unclassified | 0.000 | 0.000 | 0.013 | 0.004 | 0.000 | 0.000 |
| Candidatus Melainabacteria | Unclassified | Vampirovibrionales | Unclassified | *Vampirovibrio* | 0.000 | 0.000 | 0.000 | 0.004 | 0.000 | 0.000 |
| Chlamydiae | Chlamydiae | Chlamydiales | Criblamydia Criblamydia | Unclassified | 0.000 | 0.000 | 0.000 | 0.004 | 0.000 | 0.000 |
| Chlamydiae | Chlamydiae | Chlamydiales | cvE6 | Unclassified | 0.000 | 0.000 | 0.000 | 0.089 | 0.000 | 0.000 |
| Chlamydiae | Chlamydiae | Chlamydiales | Parachlamydiaceae | *Candidatus* Protochlamydia | 0.000 | 0.000 | 0.000 | 0.036 | 0.000 | 0.000 |
| Chlamydiae | Chlamydiae | Chlamydiales | Parachlamydiaceae | Unclassified | 0.000 | 0.000 | 0.000 | 0.004 | 0.000 | 0.000 |
| Chlamydiae | Chlamydiae | Chlamydiales | Simkaniaceae | *Candidatus* Rhabdochlamydia | 0.000 | 0.000 | 0.000 | 0.032 | 0.004 | 0.000 |
| Chlamydiae | Chlamydiae | Chlamydiales | Simkaniaceae | Unclassified | 0.000 | 0.000 | 0.000 | 0.014 | 0.000 | 0.000 |
| Chlamydiae | Chlamydiia | Parachlamydiales | Parachlamydiaceae | *Neochlamydia* | 0.000 | 0.000 | 0.000 | 0.036 | 0.000 | 0.000 |
| Chlamydiae | Chlamydiia | Parachlamydiales | Parachlamydiaceae | *Parachlamydia* | 0.000 | 0.000 | 0.000 | 0.011 | 0.000 | 0.000 |
| Chlorobi | Chlorobia | Chlorobiales | BSV26 | Unclassified | 0.001 | 0.000 | 0.000 | 0.004 | 0.000 | 0.000 |
| Chlorobi | Chlorobia | Chlorobiales | Chlorobiaceae | Uncultured 5 | 0.001 | 0.000 | 0.000 | 0.000 | 0.000 | 0.000 |
| Chlorobi | Chlorobia | Chlorobiales | OPB56 | Termite cluster | 0.458 | 0.670 | 0.000 | 0.321 | 0.761 | 0.314 |
| Chlorobi | Chlorobia | Chlorobiales | OPB56 | Unclassified | 0.000 | 0.000 | 0.000 | 0.004 | 0.004 | 0.001 |
| Chlorobi | Chlorobia | Chlorobiales | SJA-28 | Unclassified | 0.000 | 0.000 | 0.000 | 0.004 | 0.000 | 0.000 |
| Chloroflexi | Anaerolineae | Anaerolineales | Anaerolineaceae | Uncultured 11 | 0.000 | 0.000 | 0.000 | 0.025 | 0.000 | 0.002 |
| Chloroflexi | Anaerolineae | Anaerolineales | Anaerolineaceae | Uncultured 3 | 0.005 | 0.000 | 0.000 | 0.007 | 0.004 | 0.000 |
| Chloroflexi | Anaerolineae | Anaerolineales | Anaerolineaceae | Uncultured 7 | 0.000 | 0.000 | 0.000 | 0.004 | 0.000 | 0.000 |

**Table S2 continued.**

| | | | | | | | | | | |
|---|---|---|---|---|---|---|---|---|---|---|
| Chloroflexi | Caldilineae | Caldilineales | Caldilineaceae | *Caldilinea* | 0.002 | 0.000 | 0.000 | 0.007 | 0.008 | 0.001 |
| Chloroflexi | Caldilineae | Caldilineales | Caldilineaceae | Uncultured 1 | 0.001 | 0.000 | 0.000 | 0.011 | 0.000 | 0.000 |
| Chloroflexi | Caldilineae | Caldilineales | Caldilineaceae | Uncultured 5 | 0.000 | 0.000 | 0.000 | 0.000 | 0.004 | 0.000 |
| Chloroflexi | Chloroflexi | Chloroflexales | Chloroflexaceae | *Chloroflexus* | 0.000 | 0.000 | 0.000 | 0.007 | 0.000 | 0.000 |
| Chloroflexi | Chloroflexi | Chloroflexales | Chloroflexaceae | *Chloronema* | 0.000 | 0.000 | 0.000 | 0.004 | 0.000 | 0.000 |
| Chloroflexi | Chloroflexi | Chloroflexales | Chloroflexaceae | *Roseiflexus* | 0.000 | 0.000 | 0.000 | 0.014 | 0.000 | 0.000 |
| Chloroflexi | Chloroflexi | Chloroflexales | FFCH7168 | Unclassified | 0.000 | 0.000 | 0.000 | 0.004 | 0.000 | 0.000 |
| Chloroflexi | Chloroflexi | Chloroflexales | Oscillochloridaceae | *Oscillochloris* | 0.000 | 0.000 | 0.000 | 0.007 | 0.000 | 0.000 |
| Chloroflexi | Chloroflexi | Herpetosiphonales | Herpetosiphonaceae | *Herpetosiphon* | 0.001 | 0.000 | 0.000 | 0.007 | 0.000 | 0.000 |
| Chloroflexi | GIF9 | Unclassified | Unclassified | Unclassified | 0.001 | 0.000 | 0.000 | 0.004 | 0.000 | 0.000 |
| Chloroflexi | Gitt-GS-136 | Unclassified | Unclassified | Unclassified | 0.000 | 0.000 | 0.000 | 0.007 | 0.000 | 0.000 |
| Chloroflexi | JG30-KF-CM66 | Unclassified | Unclassified | Unclassified | 0.000 | 0.000 | 0.000 | 0.011 | 0.000 | 0.000 |
| Chloroflexi | JG37-AG 4 | Unclassified | Unclassified | Unclassified | 0.000 | 0.000 | 0.000 | 0.004 | 0.000 | 0.000 |
| Chloroflexi | KD4-96 | Unclassified | Unclassified | Unclassified | 0.000 | 0.000 | 0.000 | 0.039 | 0.000 | 0.000 |
| Chloroflexi | Ktedonobacteria | Ktedonobacteriales | Ktedonobacteriaceae | *Ktedonobacter* | 0.001 | 0.000 | 0.000 | 0.000 | 0.000 | 0.000 |
| Chloroflexi | S085 | Unclassified | Unclassified | Unclassified | 0.000 | 0.000 | 0.000 | 0.021 | 0.000 | 0.000 |
| Chloroflexi | SAR202 clade | Unclassified | Unclassified | Unclassified | 0.004 | 0.000 | 0.000 | 0.000 | 0.000 | 0.001 |
| Chloroflexi | Thermomicrobia | AKYG1722 | Unclassified | Unclassified | 0.000 | 0.000 | 0.000 | 0.004 | 0.000 | 0.000 |
| Chloroflexi | Thermomicrobia | JG30-KF-CM45 | Unclassified | Unclassified | 0.000 | 0.000 | 0.000 | 0.014 | 0.000 | 0.000 |
| Chloroflexi | TK10 | Unclassified | Unclassified | Unclassified | 0.000 | 0.000 | 0.000 | 0.018 | 0.004 | 0.000 |
| Cyanobacteria | Cyanobacteria | Cyanobacteriales | C0d-2 | Unclassified | 0.007 | 0.000 | 0.000 | 0.011 | 0.000 | 0.002 |
| Cyanobacteria | Cyanobacteria | Cyanobacteriales | Chloroplast | Chloroplast | 0.011 | 0.000 | 0.000 | 0.107 | 0.000 | 0.001 |
| Cyanobacteria | Cyanobacteria | Cyanobacteriales | ML635J-21 | Cluster TG2 | 0.203 | 0.191 | 0.000 | 0.086 | 0.138 | 0.063 |
| Cyanobacteria | Cyanobacteria | Cyanobacteriales | ML635J-21 | Unclassified | 0.000 | 0.000 | 0.000 | 0.007 | 0.000 | 0.000 |
| Cyanobacteria | Cyanobacteria | Cyanobacteriales | MLE1-12 | Unclassified | 0.001 | 0.000 | 0.000 | 0.032 | 0.000 | 0.000 |
| Cyanobacteria | Cyanobacteria | Cyanobacteriales | SHA-109 | Uncultured | 0.001 | 0.000 | 0.000 | 0.000 | 0.000 | 0.000 |
| Cyanobacteria | Cyanobacteria | Cyanobacteriales | SubsectionI | Uncultured | 0.000 | 0.033 | 0.000 | 0.000 | 0.000 | 0.000 |

**Table S2 continued.**

| | | | | | | | | | | |
|---|---|---|---|---|---|---|---|---|---|---|
| Cyanobacteria | Cyanobacteria | Cyanobacteriales | SubsectionII | SubgroupII | 0.000 | 0.000 | 0.000 | 0.011 | 0.000 | 0.000 |
| Cyanobacteria | Cyanobacteria | Cyanobacteriales | SubsectionIII | *Leptolyngbya* 3 | 0.001 | 0.000 | 0.000 | 0.053 | 0.000 | 0.000 |
| Cyanobacteria | Cyanobacteria | Cyanobacteriales | SubsectionIII | *Phormidium* | 0.001 | 0.000 | 0.000 | 0.000 | 0.000 | 0.000 |
| Cyanobacteria | Cyanobacteria | Cyanobacteriales | SubsectionIII | *Symploca* | 0.000 | 0.000 | 0.000 | 0.004 | 0.000 | 0.000 |
| Cyanobacteria | Cyanobacteria | Cyanobacteriales | SubsectionIII | Unclassified | 0.000 | 0.000 | 0.000 | 0.007 | 0.000 | 0.000 |
| Cyanobacteria | Cyanobacteria | Cyanobacteriales | SubsectionIII | Uncultured | 0.000 | 0.000 | 0.000 | 0.011 | 0.000 | 0.000 |
| Cyanobacteria | Cyanobacteria | Cyanobacteriales | Uncultured 7 | Unclassified | 0.000 | 0.000 | 0.000 | 0.004 | 0.000 | 0.000 |
| Cyanobacteria | Cyanobacteria | Cyanobacteriales | WD272 | Unclassified | 0.000 | 0.000 | 0.000 | 0.000 | 0.004 | 0.000 |
| Cyanobacteria | Unclassified | Synechococcales | Acaryochloridaceae | *Acaryochloris* | 0.000 | 0.000 | 0.000 | 0.004 | 0.000 | 0.000 |
| Cyanobacteria | Unclassified | Synechococcales | Trichocoleusaceae | *Trichocoleus* | 0.000 | 0.000 | 0.000 | 0.004 | 0.000 | 0.000 |
| Deferribacteres | Deferribacteres | Deferribacterales | Deferribacteraceae | *Calditerrivibrio* | 0.000 | 0.000 | 0.000 | 0.000 | 0.000 | 0.001 |
| Deferribacteres | Deferribacteres | Deferribacterales | Deferribacteraceae | *Denitrovibrio* | 0.001 | 0.000 | 0.000 | 0.000 | 0.000 | 0.000 |
| Deferribacteres | Deferribacteres | Deferribacterales | Deferribacteraceae | *Mucispirillum* | 0.482 | 0.797 | 0.000 | 0.531 | 0.110 | 0.022 |
| Deferribacteres | Deferribacteres | Deferribacterales | Deferribacteraceae | Uncultured | 0.000 | 0.000 | 0.000 | 0.004 | 0.000 | 0.000 |
| Deferribacteres | Deferribacteres | Unclassified Deferribacterales | Caldithrix Caldithrix | Unclassified | 0.000 | 0.000 | 0.000 | 0.004 | 0.000 | 0.001 |
| Deferribacteres | Deferribacteres | Unclassified Deferribacterales | LCP-89 | Unclassified | 0.000 | 0.000 | 0.000 | 0.004 | 0.000 | 0.000 |
| Deferribacteres | Deferribacteres | Unclassified Deferribacterales | SAR406 clade(Marine group A) | Unclassified | 0.000 | 0.000 | 0.000 | 0.004 | 0.000 | 0.000 |
| Deinococcus-Thermus | Deinococci | Deinococcales | Deinococcaceae | *Deinococcus* | 0.001 | 0.000 | 0.000 | 0.000 | 0.000 | 0.001 |
| Deinococcus-Thermus | Deinococci | Thermales | Thermaceae | *Meiothermus* | 0.000 | 0.000 | 0.000 | 0.004 | 0.000 | 0.000 |
| Deinococcus-Thermus | Deinococcus-Thermus | KD3-62 | Uncultured | Uncultured | 0.001 | 0.000 | 0.000 | 0.000 | 0.000 | 0.000 |
| Elusimicrobia | Elusimicrobia | Elusimicrobiales | Lineage IId | Unclassified | 0.001 | 0.000 | 0.000 | 0.000 | 0.000 | 0.000 |
| Elusimicrobia | Elusimicrobia | Elusimicrobiales | Lineage IV | Unclassified | 0.001 | 0.000 | 0.000 | 0.007 | 0.008 | 0.000 |
| Elusimicrobia | Elusimicrobia | Endomicrobiales | Endomicrobiaceae | *Endomicrobium* | 0.016 | 0.064 | 0.000 | 0.025 | 0.000 | 0.000 |
| Elusimicrobia | Elusimicrobia | Endomicrobiales | Endomicrobiaceae | Environmental cluster I | 0.001 | 0.000 | 0.000 | 0.000 | 0.000 | 0.000 |
| Fibrobacteres | Subphylum 2 | Insect cluster | Termite cluster I | Subcluster Ia | 0.000 | 0.000 | 0.000 | 0.075 | 0.000 | 0.000 |

**Table S2 continued.**

| | | | | | | | | | | |
|---|---|---|---|---|---|---|---|---|---|---|
| Fibrobacteres | Subphylum 2 | Insect cluster | Termite cluster I | Subcluster Ib | 0.000 | 0.000 | 0.000 | 0.110 | 0.000 | 0.001 |
| Firmicutes | Bacilli | Bacillales | Alicyclobacillaceae | *Alicyclobacillus* | 0.001 | 0.000 | 0.000 | 0.000 | 0.000 | 0.000 |
| Firmicutes | Bacilli | Bacillales | Bacillaceae | *Amphibacillus* | 0.000 | 0.000 | 0.000 | 0.007 | 0.000 | 0.000 |
| Firmicutes | Bacilli | Bacillales | Bacillaceae | *Anoxybacillus* | 0.001 | 0.000 | 0.013 | 0.004 | 0.004 | 0.001 |
| Firmicutes | Bacilli | Bacillales | Bacillaceae | *Bacillus* | 0.005 | 0.096 | 0.077 | 1.154 | 0.004 | 0.002 |
| Firmicutes | Bacilli | Bacillales | Bacillaceae | *Bacillus* 1 | 0.000 | 0.000 | 0.000 | 0.043 | 0.004 | 0.000 |
| Firmicutes | Bacilli | Bacillales | Bacillaceae | *Bacillus* 10 | 0.000 | 0.000 | 0.000 | 0.014 | 0.000 | 0.000 |
| Firmicutes | Bacilli | Bacillales | Bacillaceae | *Bacillus* 11 | 0.001 | 0.000 | 0.039 | 0.050 | 0.004 | 0.000 |
| Firmicutes | Bacilli | Bacillales | Bacillaceae | *Bacillus* 12 | 0.000 | 0.000 | 0.000 | 0.004 | 0.000 | 0.000 |
| Firmicutes | Bacilli | Bacillales | Bacillaceae | *Bacillus* 14 | 0.001 | 0.000 | 0.039 | 0.310 | 0.000 | 0.001 |
| Firmicutes | Bacilli | Bacillales | Bacillaceae | *Bacillus* 2 | 0.000 | 0.032 | 0.000 | 0.021 | 0.000 | 0.000 |
| Firmicutes | Bacilli | Bacillales | Bacillaceae | *Bacillus* 3 | 0.000 | 0.000 | 0.000 | 0.004 | 0.000 | 0.000 |
| Firmicutes | Bacilli | Bacillales | Bacillaceae | *Bacillus* 4 | 0.002 | 0.000 | 0.013 | 0.057 | 0.000 | 0.002 |
| Firmicutes | Bacilli | Bacillales | Bacillaceae | *Bacillus* 5 | 0.000 | 0.000 | 0.000 | 0.007 | 0.000 | 0.000 |
| Firmicutes | Bacilli | Bacillales | Bacillaceae | *Bacillus* 6 | 0.000 | 0.000 | 0.000 | 0.007 | 0.000 | 0.000 |
| Firmicutes | Bacilli | Bacillales | Bacillaceae | *Bacillus* 7 | 0.001 | 0.000 | 0.000 | 0.591 | 0.000 | 0.000 |
| Firmicutes | Bacilli | Bacillales | Bacillaceae | *Bacillus* 9 | 0.000 | 0.000 | 0.000 | 0.007 | 0.000 | 0.000 |
| Firmicutes | Bacilli | Bacillales | Bacillaceae | *Fictibacillus* | 0.000 | 0.000 | 0.013 | 0.004 | 0.000 | 0.000 |
| Firmicutes | Bacilli | Bacillales | Bacillaceae | *Gracilibacillus* | 0.000 | 0.000 | 0.000 | 0.004 | 0.000 | 0.000 |
| Firmicutes | Bacilli | Bacillales | Bacillaceae | *Lysinibacillus* | 0.000 | 0.000 | 0.000 | 0.007 | 0.004 | 0.001 |
| Firmicutes | Bacilli | Bacillales | Bacillaceae | *Oceanobacillus* | 0.000 | 0.000 | 0.000 | 0.025 | 0.000 | 0.000 |
| Firmicutes | Bacilli | Bacillales | Bacillaceae | *Pontibacillus* | 0.000 | 0.000 | 0.000 | 0.004 | 0.000 | 0.000 |
| Firmicutes | Bacilli | Bacillales | Bacillaceae | *Psychrobacillus* | 0.000 | 0.000 | 0.000 | 0.004 | 0.000 | 0.000 |
| Firmicutes | Bacilli | Bacillales | Bacillaceae | *Salisediminibacterium* | 0.000 | 0.000 | 0.000 | 0.000 | 0.004 | 0.000 |
| Firmicutes | Bacilli | Bacillales | Bacillaceae | *Terribacillus* | 0.000 | 0.000 | 0.000 | 0.025 | 0.000 | 0.001 |
| Firmicutes | Bacilli | Bacillales | Bacillaceae | *Virgibacillus* | 0.000 | 0.000 | 0.000 | 0.007 | 0.000 | 0.000 |
| Firmicutes | Bacilli | Bacillales | Bacillaceae 5 | Unclassified | 0.000 | 0.000 | 0.000 | 0.004 | 0.000 | 0.000 |

**Table S2 continued.**

| | | | | | | | | | | |
|---|---|---|---|---|---|---|---|---|---|---|
| Firmicutes | Bacilli | Bacillales | Bacillaceae 6 | Unclassified | 0.000 | 0.000 | 0.000 | 0.000 | 0.004 | 0.000 |
| Firmicutes | Bacilli | Bacillales | Family XI Incertae Sedis | *Gemella* | 0.000 | 0.000 | 0.000 | 0.007 | 0.000 | 0.001 |
| Firmicutes | Bacilli | Bacillales | Family XII Incertae Sedis | *Exiguobacterium* | 0.000 | 0.000 | 0.000 | 0.011 | 0.000 | 0.001 |
| Firmicutes | Bacilli | Bacillales | Family XII Incertae Sedis | Unclassified | 0.000 | 0.000 | 0.000 | 0.004 | 0.000 | 0.000 |
| Firmicutes | Bacilli | Bacillales | Listeriaceae | *Brochothrix* | 0.000 | 0.000 | 0.026 | 0.000 | 0.000 | 0.000 |
| Firmicutes | Bacilli | Bacillales | Listeriaceae | *Listeria* | 0.002 | 0.000 | 0.000 | 0.000 | 0.000 | 0.000 |
| Firmicutes | Bacilli | Bacillales | Paenibacillaceae | *Ammoniphilus* | 0.000 | 0.000 | 0.000 | 0.007 | 0.000 | 0.000 |
| Firmicutes | Bacilli | Bacillales | Paenibacillaceae | *Aneurinibacillus* | 0.000 | 0.000 | 0.000 | 0.018 | 0.000 | 0.000 |
| Firmicutes | Bacilli | Bacillales | Paenibacillaceae | *Brevibacillus* | 0.001 | 0.000 | 0.000 | 0.014 | 0.000 | 0.000 |
| Firmicutes | Bacilli | Bacillales | Paenibacillaceae | *Cohnella* | 0.001 | 0.000 | 0.000 | 0.021 | 0.000 | 0.000 |
| Firmicutes | Bacilli | Bacillales | Paenibacillaceae | *Gorillibacterium* | 0.000 | 0.000 | 0.000 | 0.004 | 0.000 | 0.000 |
| Firmicutes | Bacilli | Bacillales | Paenibacillaceae | *Oxalophagus* | 0.000 | 0.000 | 0.000 | 0.004 | 0.000 | 0.000 |
| Firmicutes | Bacilli | bacillales | Paenibacillaceae | *Paenibacillus* | 0.002 | 0.000 | 0.000 | 0.125 | 0.000 | 0.001 |
| Firmicutes | Bacilli | Bacillales | Paenibacillaceae 1 | Unclassified | 0.000 | 0.000 | 0.000 | 0.011 | 0.000 | 0.000 |
| Firmicutes | Bacilli | Bacillales | Paenibacillaceae 2 | *Paenibacillus* 1 | 0.002 | 0.000 | 0.000 | 0.000 | 0.000 | 0.000 |
| Firmicutes | Bacilli | Bacillales | Paenibacillaceae 2 | *Paenibacillus* 12 | 0.000 | 0.000 | 0.000 | 0.014 | 0.000 | 0.000 |
| Firmicutes | Bacilli | Bacillales | Paenibacillaceae 2 | *Paenibacillus* 13 | 0.002 | 0.000 | 0.000 | 0.014 | 0.004 | 0.000 |
| Firmicutes | Bacilli | Bacillales | Paenibacillaceae 2 | *Paenibacillus* 17 | 0.000 | 0.000 | 0.013 | 0.011 | 0.000 | 0.000 |
| Firmicutes | Bacilli | Bacillales | Paenibacillaceae 2 | *Paenibacillus* 6 | 0.000 | 0.000 | 0.000 | 0.004 | 0.000 | 0.000 |
| Firmicutes | Bacilli | Bacillales | Paenibacillaceae 2 | *Paenibacillus* 8 | 0.000 | 0.000 | 0.000 | 0.014 | 0.000 | 0.000 |
| Firmicutes | Bacilli | Bacillales | Paenibacillaceae 2 | Unclassified | 0.000 | 0.000 | 0.013 | 0.000 | 0.000 | 0.000 |
| Firmicutes | Bacilli | Bacillales | Planococcaceae | *Bhargavaea* | 0.000 | 0.000 | 0.000 | 0.004 | 0.000 | 0.000 |
| Firmicutes | Bacilli | Bacillales | Planococcaceae | *Chryseomicrobium* | 0.000 | 0.000 | 0.000 | 0.004 | 0.000 | 0.000 |
| Firmicutes | Bacilli | Bacillales | Planococcaceae | *Kurthia* | 0.000 | 0.000 | 0.000 | 0.004 | 0.004 | 0.000 |
| Firmicutes | Bacilli | Bacillales | Planococcaceae | *Planococcus* | 0.001 | 0.000 | 0.000 | 0.007 | 0.024 | 0.006 |
| Firmicutes | Bacilli | Bacillales | Planococcaceae | *Planomicrobium* | 0.000 | 0.000 | 0.000 | 0.000 | 0.008 | 0.000 |
| Firmicutes | Bacilli | Bacillales | Planococcaceae | *Savagea* | 0.000 | 0.000 | 0.000 | 0.004 | 0.000 | 0.000 |

**Table S2 continued.**

| | | | | | | | | | | |
|---|---|---|---|---|---|---|---|---|---|---|
| Firmicutes | Bacilli | Bacillales | Planococcaceae | *Sporosarcina* | 0.000 | 0.000 | 0.000 | 0.004 | 0.000 | 0.000 |
| Firmicutes | Bacilli | Bacillales | Planococcaceae | *Ureibacillus* | 0.000 | 0.000 | 0.013 | 0.000 | 0.000 | 0.001 |
| Firmicutes | Bacilli | Bacillales | Planococcaceae 2 | Incertae Sedis 3 | 0.001 | 0.000 | 0.026 | 0.004 | 0.000 | 0.000 |
| Firmicutes | Bacilli | Bacillales | Planococcaceae 2 | Incertae Sedis 4 | 0.000 | 0.000 | 0.000 | 0.004 | 0.000 | 0.000 |
| Firmicutes | Bacilli | Bacillales | Planococcaceae 2 | Incertae Sedis 5 | 0.000 | 0.000 | 0.000 | 0.007 | 0.000 | 0.000 |
| Firmicutes | Bacilli | Bacillales | Planococcaceae 2 | Incertae Sedis 6 | 0.000 | 0.000 | 0.013 | 0.000 | 0.000 | 0.000 |
| Firmicutes | Bacilli | Bacillales | Planococcaceae 2 | *Planococcus* 1 | 0.000 | 0.000 | 0.013 | 0.007 | 0.000 | 0.000 |
| Firmicutes | Bacilli | Bacillales | Planococcaceae 2 | Unclassified | 0.000 | 0.000 | 0.013 | 0.000 | 0.000 | 0.000 |
| Firmicutes | Bacilli | Bacillales | Staphylococcaceae | *Jeotgalicoccus* | 0.000 | 0.000 | 0.000 | 0.004 | 0.000 | 0.001 |
| Firmicutes | Bacilli | Bacillales | Staphylococcaceae | *Macrococcus* | 0.000 | 0.000 | 0.000 | 0.007 | 0.000 | 0.001 |
| Firmicutes | Bacilli | Bacillales | Staphylococcaceae | *Staphylococcus* | 0.000 | 0.000 | 0.039 | 0.021 | 0.000 | 0.002 |
| Firmicutes | Bacilli | Bacillales | Staphylococcaceae | *Staphylococcus* 1 | 0.000 | 0.000 | 0.077 | 0.000 | 0.000 | 0.002 |
| Firmicutes | Bacilli | Bacillales | Thermoactinomycetaceae | *Baia* | 0.000 | 0.000 | 0.000 | 0.004 | 0.000 | 0.000 |
| Firmicutes | Bacilli | Bacillales | Thermoactinomycetaceae | *Lihuaxuella* | 0.001 | 0.000 | 0.000 | 0.000 | 0.000 | 0.000 |
| Firmicutes | Bacilli | Bacillales | Thermoactinomycetaceae | *Marinithermofilum* | 0.000 | 0.000 | 0.000 | 0.000 | 0.004 | 0.000 |
| Firmicutes | Bacilli | Bacillales | Thermoactinomycetaceae | *Thermoactinomyces* | 0.000 | 0.000 | 0.000 | 0.000 | 0.000 | 0.002 |
| Firmicutes | Bacilli | Bacillales | Unclassified | *Geomicrobium* | 0.000 | 0.000 | 0.000 | 0.000 | 0.012 | 0.010 |
| Firmicutes | Bacilli | Bacillales | Unclassified | Unclassified | 0.000 | 0.000 | 0.000 | 0.011 | 0.004 | 0.001 |
| Firmicutes | Bacilli | Cluster 4-15 | Unclassified | Unclassified | 0.000 | 0.000 | 0.000 | 0.004 | 0.000 | 0.000 |
| Firmicutes | Bacilli | Lactobacillales | Aerococcaceae | *Aerococcus* | 0.004 | 0.000 | 0.013 | 0.004 | 0.000 | 0.001 |
| Firmicutes | Bacilli | Lactobacillales | Aerococcaceae | *Facklamia* | 0.000 | 0.000 | 0.013 | 0.004 | 0.000 | 0.000 |
| Firmicutes | Bacilli | Lactobacillales | Carnobacteriaceae | *Alkalibacterium* | 0.001 | 0.000 | 0.000 | 0.000 | 0.004 | 0.001 |
| Firmicutes | Bacilli | Lactobacillales | Carnobacteriaceae | *Alloiococcus* | 0.001 | 0.000 | 0.000 | 0.000 | 0.000 | 0.000 |
| Firmicutes | Bacilli | Lactobacillales | Carnobacteriaceae | *Atopobacter* | 0.000 | 0.000 | 0.000 | 0.000 | 0.004 | 0.000 |
| Firmicutes | Bacilli | Lactobacillales | Carnobacteriaceae | *Atopostipes* | 0.001 | 0.000 | 0.000 | 0.000 | 0.004 | 0.001 |
| Firmicutes | Bacilli | Lactobacillales | Carnobacteriaceae | *Carnobacterium* | 0.002 | 0.000 | 0.000 | 0.000 | 0.000 | 0.000 |
| Firmicutes | Bacilli | Lactobacillales | Carnobacteriaceae | *Desemzia* | 0.000 | 0.000 | 0.013 | 0.000 | 0.000 | 0.001 |

**Table S2 continued.**

| | | | | | | | | | | |
|---|---|---|---|---|---|---|---|---|---|---|
| Firmicutes | Bacilli | Lactobacillales | Carnobacteriaceae | *Marinilactibacillus* | 0.000 | 0.000 | 0.000 | 0.000 | 0.004 | 0.000 |
| Firmicutes | Bacilli | Lactobacillales | Carnobacteriaceae | *Trichococcus* | 0.002 | 0.000 | 0.064 | 0.000 | 0.000 | 0.002 |
| Firmicutes | Bacilli | Lactobacillales | Carnobacteriaceae | *Trichococcus* 1 | 0.001 | 0.000 | 0.000 | 0.000 | 0.000 | 0.000 |
| Firmicutes | Bacilli | Lactobacillales | Cockroach cluster | Unclassified | 0.002 | 0.000 | 0.039 | 0.000 | 0.000 | 0.000 |
| Firmicutes | Bacilli | Lactobacillales | Enterococcaceae | *Enterococcus* | 0.299 | 0.000 | 4.190 | 0.046 | 0.085 | 0.103 |
| Firmicutes | Bacilli | Lactobacillales | Enterococcaceae | *Enterococcus* 1 | 0.009 | 0.000 | 0.155 | 0.004 | 0.000 | 0.002 |
| Firmicutes | Bacilli | Lactobacillales | Enterococcaceae | *Enterococcus* 2 | 0.017 | 0.000 | 0.503 | 0.014 | 0.000 | 0.000 |
| Firmicutes | Bacilli | Lactobacillales | Enterococcaceae | *Enterococcus* 3 | 0.000 | 0.000 | 0.013 | 0.000 | 0.000 | 0.000 |
| Firmicutes | Bacilli | Lactobacillales | Enterococcaceae | *Melissococcus* | 0.000 | 0.000 | 0.052 | 0.000 | 0.000 | 0.001 |
| Firmicutes | Bacilli | Lactobacillales | Enterococcaceae | *Pilibacter* | 0.000 | 0.000 | 0.013 | 0.000 | 0.000 | 0.000 |
| Firmicutes | Bacilli | Lactobacillales | Enterococcaceae | Termite cluster 1 | 0.001 | 0.000 | 0.013 | 0.000 | 0.000 | 0.000 |
| Firmicutes | Bacilli | Lactobacillales | Enterococcaceae | *Vagococcus* | 0.001 | 0.000 | 0.077 | 0.008 | 0.020 | 0.022 |
| Firmicutes | Bacilli | Lactobacillales | Enterococcaceae 1 | Unclassified | 0.002 | 0.000 | 0.000 | 0.004 | 0.000 | 0.000 |
| Firmicutes | Bacilli | Lactobacillales | Enterococcaceae 1 | Unclassified | 0.000 | 0.000 | 0.000 | 0.000 | 0.000 | |
| Firmicutes | Bacilli | Lactobacillales | Enterococcaceae 2 | Unclassified | 0.000 | 0.000 | 0.219 | 0.004 | 0.000 | 0.002 |
| Firmicutes | Bacilli | Lactobacillales | Enterococcaceae 2 | Unclassified | 0.015 | 0.000 | 0.000 | 0.000 | 0.000 | 0.000 |
| Firmicutes | Bacilli | Lactobacillales | Enterococcaceaea | *Tetragenococcus* | 0.000 | 0.000 | 0.000 | 0.000 | 0.000 | 0.001 |
| Firmicutes | Bacilli | Lactobacillales | Insect cluster | Unclassified | 0.000 | 0.000 | 0.026 | 0.000 | 0.000 | 0.000 |
| Firmicutes | Bacilli | Lactobacillales | Lactobacillaceae | *Lactobacillus* | 0.002 | 0.000 | 0.013 | 0.014 | 0.000 | 0.000 |
| Firmicutes | Bacilli | Lactobacillales | Lactobacillaceae | *Lactobacillus* 1 | 0.001 | 0.000 | 0.000 | 0.007 | 0.004 | 0.000 |
| Firmicutes | Bacilli | Lactobacillales | Lactobacillaceae | *Lactobacillus* 4 | 0.000 | 0.000 | 0.013 | 0.000 | 0.000 | 0.000 |
| Firmicutes | Bacilli | Lactobacillales | Lactobacillaceae | *Lactobacillus* 8 | 0.000 | 0.000 | 0.039 | 0.000 | 0.000 | 0.000 |
| Firmicutes | Bacilli | Lactobacillales | Lactobacillaceae | Unclassified | 0.000 | 0.000 | 0.013 | 0.000 | 0.000 | 0.000 |
| Firmicutes | Bacilli | Lactobacillales | Leuconostocaceae | *Leuconostoc* | 0.000 | 0.000 | 0.000 | 0.004 | 0.000 | 0.000 |
| Firmicutes | Bacilli | Lactobacillales | Leuconostocaceae | *Weissella* | 0.000 | 0.000 | 0.000 | 0.004 | 0.000 | 0.000 |
| Firmicutes | Bacilli | Lactobacillales | q48f312-pp90 | Unclassified | 0.000 | 0.000 | 0.026 | 0.000 | 0.000 | 0.003 |
| Firmicutes | Bacilli | Lactobacillales | q48f312-pp90 | Unclassified | 0.001 | 0.000 | 0.000 | 0.000 | 0.000 | 0.000 |

**Table S2 continued.**

| | | | | | | | | | | |
|---|---|---|---|---|---|---|---|---|---|---|
| Firmicutes | Bacilli | Lactobacillales | Streptococcaceae | *Lactococcus* | 0.002 | 0.000 | 0.013 | 0.007 | 0.175 | 0.000 |
| Firmicutes | Bacilli | Lactobacillales | Streptococcaceae | *Lactococcus* 1 | 0.030 | 0.032 | 0.155 | 1.069 | 0.008 | 0.001 |
| Firmicutes | Bacilli | Lactobacillales | Streptococcaceae | *Lactococcus* 3 | 0.004 | 0.000 | 0.013 | 0.000 | 0.004 | 0.000 |
| Firmicutes | Bacilli | Lactobacillales | Streptococcaceae | *Lactovum* | 1.295 | 0.478 | 15.612 | 0.363 | 0.028 | 0.024 |
| Firmicutes | Bacilli | Lactobacillales | Streptococcaceae | *Streptococcus* | 0.032 | 0.064 | 0.374 | 0.118 | 0.008 | 0.001 |
| Firmicutes | Bacilli | Lactobacillales | Streptococcaceae | Unclassified | 0.000 | 0.000 | 0.000 | 0.004 | 0.000 | 0.000 |
| Firmicutes | Bacilli | Lactobacillales | Streptococcaceae | Uncultured 1 | 0.001 | 0.000 | 0.000 | 0.000 | 0.000 | 0.000 |
| Firmicutes | Bacilli | Lactobacillales | Streptococcaceae | Uncultured 1 | 0.000 | 0.000 | 0.000 | 0.000 | 0.008 | 0.000 |
| Firmicutes | Bacilli | Lactobacillales | Termite cluster 2 | Unclassified | 0.000 | 0.000 | 0.000 | 0.000 | 0.004 | 0.001 |
| Firmicutes | Bacilli | Lactobacillales | Termite cluster 2 | Unclassified | 0.001 | 0.000 | 0.000 | 0.000 | 0.000 | 0.000 |
| Firmicutes | Bacilli | Lactobacillales | Unclassified | Unclassified | 0.000 | 0.000 | 0.013 | 0.000 | 0.000 | 0.000 |
| Firmicutes | Bacilli | Lactobacillales | Unclassified | Unclassified | 0.001 | 0.000 | 0.000 | 0.000 | 0.000 | 0.000 |
| Firmicutes | Bacilli | Unclassified | Unclassified | Unclassified | 0.000 | 0.000 | 0.000 | 0.007 | 0.004 | 0.000 |
| Firmicutes | Bacilli | VAN12 | Unclassified | Unclassified | 0.000 | 0.000 | 0.013 | 0.007 | 0.000 | 0.000 |
| Firmicutes | CK-1C4-19 | Unclassified | Unclassified | Unclassified | 0.000 | 0.000 | 0.000 | 0.004 | 0.008 | 0.002 |
| Firmicutes | Clostridia | Clostridiales | Catabacteriaceae | *Catabacter* | 0.080 | 0.159 | 0.013 | 0.029 | 0.110 | 0.049 |
| Firmicutes | Clostridia | Clostridiales | Christensenellaceae | *Christensenella* | 0.010 | 0.032 | 0.026 | 0.004 | 0.065 | 0.030 |
| Firmicutes | Clostridia | Clostridiales | Clostridiaceae | *Alkaliphilus* | 0.000 | 0.000 | 0.000 | 0.000 | 0.000 | 0.001 |
| Firmicutes | Clostridia | Clostridiales | Clostridiaceae | *Anaerobacter* | 0.000 | 0.000 | 0.000 | 0.004 | 0.000 | 0.000 |
| Firmicutes | Clostridia | Clostridiales | Clostridiaceae | *Brassicibacter* | 0.000 | 0.000 | 0.000 | 0.000 | 0.000 | 0.001 |
| Firmicutes | Clostridia | Clostridiales | Clostridiaceae | *Butyricicoccus* | 0.002 | 0.000 | 0.000 | 0.004 | 0.000 | 0.000 |
| Firmicutes | Clostridia | Clostridiales | Clostridiaceae | *Caloramator* | 0.000 | 0.000 | 0.000 | 0.004 | 0.000 | 0.001 |
| Firmicutes | Clostridia | Clostridiales | Clostridiaceae | *Clostridium* | 0.173 | 0.159 | 5.324 | 1.058 | 0.094 | 0.049 |
| Firmicutes | Clostridia | Clostridiales | Clostridiaceae | *Crassaminicella* | 0.000 | 0.000 | 0.000 | 0.004 | 0.000 | 0.000 |
| Firmicutes | Clostridia | Clostridiales | Clostridiaceae | *Desnuesiella* | 0.000 | 0.000 | 0.000 | 0.014 | 0.000 | 0.000 |
| Firmicutes | Clostridia | Clostridiales | Clostridiaceae | *Falcatimonas* | 0.001 | 0.000 | 0.000 | 0.000 | 0.000 | 0.000 |
| Firmicutes | Clostridia | Clostridiales | Clostridiaceae | *Lactonifactor* | 0.017 | 0.000 | 0.374 | 0.064 | 0.000 | 0.003 |

**Table S2 continued.**

| | | | | | | | | | | |
|---|---|---|---|---|---|---|---|---|---|---|
| Firmicutes | Clostridia | Clostridiales | Clostridiaceae | *Lutispora* | 0.002 | 0.000 | 0.000 | 0.007 | 0.000 | 0.000 |
| Firmicutes | Clostridia | Clostridiales | Clostridiaceae | *Mordavella* | 0.001 | 0.000 | 0.000 | 0.000 | 0.000 | 0.000 |
| Firmicutes | Clostridia | Clostridiales | Clostridiaceae | *Natronincola* | 0.002 | 0.000 | 0.000 | 0.000 | 0.000 | 0.000 |
| Firmicutes | Clostridia | Clostridiales | Clostridiaceae | *Oceanirhabdus* | 0.001 | 0.000 | 0.000 | 0.000 | 0.000 | 0.000 |
| Firmicutes | Clostridia | Clostridiales | Clostridiaceae | *Oxobacter* | 0.002 | 0.000 | 0.000 | 0.000 | 0.000 | 0.001 |
| Firmicutes | Clostridia | Clostridiales | Clostridiaceae | *Thermobrachium* | 0.000 | 0.000 | 0.000 | 0.004 | 0.000 | 0.000 |
| Firmicutes | Clostridia | Clostridiales | Clostridiaceae | *Thermotalea* | 0.001 | 0.000 | 0.000 | 0.000 | 0.000 | 0.001 |
| Firmicutes | Clostridia | Clostridiales | Eubacteriaceae | *Eubacterium* | 0.014 | 0.000 | 0.077 | 0.004 | 0.028 | 0.006 |
| Firmicutes | Clostridia | Clostridiales | Eubacteriaceae | *Intestinibacillus* | 0.001 | 0.000 | 0.000 | 0.000 | 0.000 | 0.000 |
| Firmicutes | Clostridia | Clostridiales | Family XIII Incertae Sedis | *Emergencia* | 0.002 | 0.000 | 0.000 | 0.004 | 0.000 | 0.000 |
| Firmicutes | Clostridia | Clostridiales | Family XIII Incertae Sedis | *Anaerovorax* | 0.000 | 0.000 | 0.000 | 0.004 | 0.004 | 0.001 |
| Firmicutes | Clostridia | Clostridiales | Family XIII Incertae Sedis | *Casaltella* | 0.000 | 0.000 | 0.000 | 0.000 | 0.004 | 0.001 |
| Firmicutes | Clostridia | Clostridiales | Gracilibacteraceae | *Gracilibacter* | 0.017 | 0.032 | 0.000 | 0.007 | 0.024 | 0.010 |
| Firmicutes | Clostridia | Clostridiales | Heliobacteriaceae | *Heliobacterium* | 0.001 | 0.000 | 0.000 | 0.000 | 0.000 | 0.000 |
| Firmicutes | Clostridia | Clostridiales | Hungateiclostridiaceae | *Anaerobacterium* | 0.007 | 0.000 | 0.000 | 0.014 | 0.012 | 0.002 |
| Firmicutes | Clostridia | Clostridiales | Hungateiclostridiaceae | *Ercella* | 0.001 | 0.000 | 0.000 | 0.004 | 0.000 | 0.000 |
| Firmicutes | Clostridia | Clostridiales | Hungateiclostridiaceae | *Pseudobacteroides* | 0.001 | 0.000 | 0.000 | 0.000 | 0.004 | 0.001 |
| Firmicutes | Clostridia | Clostridiales | Hungateiclostridiaceae | *Ruminiclostridium* | 0.002 | 0.032 | 0.000 | 0.000 | 0.000 | 0.000 |
| Firmicutes | Clostridia | Clostridiales | Hungateiclostridiaceae | *Saccharofermentans* | 0.001 | 0.000 | 0.000 | 0.004 | 0.000 | 0.000 |
| Firmicutes | Clostridia | Clostridiales | Lachnospiraceae | *Anaerobium* | 0.001 | 0.000 | 0.000 | 0.014 | 0.000 | 0.000 |
| Firmicutes | Clostridia | Clostridiales | Lachnospiraceae | *Anaerocolumna* | 0.000 | 0.000 | 0.000 | 0.018 | 0.000 | 0.000 |
| Firmicutes | Clostridia | Clostridiales | Lachnospiraceae | *Anaerotaenia* | 0.005 | 0.000 | 0.000 | 0.029 | 0.000 | 0.000 |
| Firmicutes | Clostridia | Clostridiales | Lachnospiraceae | *Anaerotignum* | 0.001 | 0.032 | 0.000 | 0.000 | 0.000 | 0.001 |
| Firmicutes | Clostridia | Clostridiales | Lachnospiraceae | *Butyrivibrio* | 0.004 | 0.000 | 0.168 | 0.000 | 0.000 | 0.000 |
| Firmicutes | Clostridia | Clostridiales | Lachnospiraceae | *Cellulosilyticum* | 0.001 | 0.000 | 0.000 | 0.000 | 0.000 | 0.000 |
| Firmicutes | Clostridia | Clostridiales | Lachnospiraceae | *Coprococcus* | 0.002 | 0.000 | 0.000 | 0.004 | 0.004 | 0.000 |
| Firmicutes | Clostridia | Clostridiales | Lachnospiraceae | *Cuneatibacter* | 0.016 | 0.000 | 0.000 | 0.004 | 0.000 | 0.000 |

**Table S2 continued.**

| | | | | | | | | | | |
|---|---|---|---|---|---|---|---|---|---|---|
| Firmicutes | Clostridia | Clostridiales | Lachnospiraceae | *Eisenbergiella* | 0.001 | 0.000 | 0.013 | 0.000 | 0.000 | 0.000 |
| Firmicutes | Clostridia | Clostridiales | Lachnospiraceae | *Faecalicatena* | 0.000 | 0.000 | 0.000 | 0.004 | 0.000 | 0.000 |
| Firmicutes | Clostridia | Clostridiales | Lachnospiraceae | *Herbinix* | 0.001 | 0.000 | 0.000 | 0.000 | 0.000 | 0.000 |
| Firmicutes | Clostridia | Clostridiales | Lachnospiraceae | *Lachnoanaerobaculum* | 0.001 | 0.000 | 0.013 | 0.000 | 0.004 | 0.000 |
| Firmicutes | Clostridia | Clostridiales | Lachnospiraceae | *Lachnobacterium* | 0.001 | 0.000 | 0.000 | 0.000 | 0.000 | 0.000 |
| Firmicutes | Clostridia | Clostridiales | Lachnospiraceae | *Lachnoclostridium* | 0.001 | 0.000 | 0.013 | 0.000 | 0.000 | 0.000 |
| Firmicutes | Clostridia | Clostridiales | Lachnospiraceae | *Lachnotalea* | 0.000 | 0.000 | 0.026 | 0.004 | 0.000 | 0.000 |
| Firmicutes | Clostridia | Clostridiales | Lachnospiraceae | *Mobilisporobacter* | 0.000 | 0.000 | 0.000 | 0.004 | 0.000 | 0.000 |
| Firmicutes | Clostridia | Clostridiales | Lachnospiraceae | *Mobilitalea* | 0.000 | 0.000 | 0.026 | 0.000 | 0.000 | 0.000 |
| Firmicutes | Clostridia | Clostridiales | Lachnospiraceae | *Muricomes* | 0.001 | 0.000 | 0.000 | 0.000 | 0.000 | 0.000 |
| Firmicutes | Clostridia | Clostridiales | Lachnospiraceae | *Murimonas* | 0.000 | 0.000 | 0.013 | 0.000 | 0.000 | 0.000 |
| Firmicutes | Clostridia | Clostridiales | Lachnospiraceae | *Parasporobacterium* | 0.000 | 0.000 | 0.000 | 0.014 | 0.000 | 0.000 |
| Firmicutes | Clostridia | Clostridiales | Lachnospiraceae | *Pseudobutyrivibrio* | 0.002 | 0.000 | 0.000 | 0.000 | 0.000 | 0.000 |
| Firmicutes | Clostridia | Clostridiales | Lachnospiraceae | *Roseburia* | 0.001 | 0.000 | 0.000 | 0.000 | 0.000 | 0.000 |
| Firmicutes | Clostridia | Clostridiales | Lachnospiraceae | *Tyzzerella* | 0.004 | 0.000 | 0.026 | 0.004 | 0.000 | 0.000 |
| Firmicutes | Clostridia | Clostridiales | Oscillospiraceae | *Oscillibacter* | 0.021 | 0.032 | 0.013 | 0.007 | 0.020 | 0.005 |
| Firmicutes | Clostridia | Clostridiales | Peptococcaceae | *Dehalobacter* | 0.000 | 0.000 | 0.013 | 0.004 | 0.004 | 0.000 |
| Firmicutes | Clostridia | Clostridiales | Peptococcaceae | *Desulfitispora* | 0.000 | 0.032 | 0.000 | 0.000 | 0.000 | 0.000 |
| Firmicutes | Clostridia | Clostridiales | Peptococcaceae | *Desulfitobacterium* | 0.002 | 0.000 | 0.000 | 0.000 | 0.004 | 0.002 |
| Firmicutes | Clostridia | Clostridiales | Peptococcaceae | *Desulfosporosinus* | 0.002 | 0.000 | 0.000 | 0.000 | 0.004 | 0.002 |
| Firmicutes | Clostridia | Clostridiales | Peptococcaceae | *Desulfotomaculum* | 0.004 | 0.000 | 0.026 | 0.004 | 0.000 | 0.001 |
| Firmicutes | Clostridia | Clostridiales | Peptostreptococcaceae | *Acetoanaerobium* | 0.001 | 0.000 | 0.000 | 0.000 | 0.000 | 0.001 |
| Firmicutes | Clostridia | Clostridiales | Peptostreptococcaceae | *Filifactor* | 0.000 | 0.000 | 0.000 | 0.000 | 0.004 | 0.001 |
| Firmicutes | Clostridia | Clostridiales | Peptostreptococcaceae | *Intestinibacter* | 0.000 | 0.000 | 0.000 | 0.004 | 0.000 | 0.000 |
| Firmicutes | Clostridia | Clostridiales | Peptostreptococcaceae | *Peptoclostridium* | 0.000 | 0.000 | 0.000 | 0.004 | 0.000 | 0.001 |
| Firmicutes | Clostridia | Clostridiales | Peptostreptococcaceae | *Proteocatella* | 0.000 | 0.000 | 0.000 | 0.004 | 0.000 | 0.000 |
| Firmicutes | Clostridia | Clostridiales | Peptostreptococcaceae | *Sporacetigenium* | 0.000 | 0.000 | 0.000 | 0.011 | 0.000 | 0.000 |

**Table S2 continued.**

| | | | | | | | | | | |
|---|---|---|---|---|---|---|---|---|---|---|
| Firmicutes | Clostridia | Clostridiales | Peptostreptococcaceae | *Tepidibacter* | 0.000 | 0.000 | 0.000 | 0.007 | 0.000 | 0.000 |
| Firmicutes | Clostridia | Clostridiales | Ruminococcaceae | *Anaerofilum* | 0.000 | 0.000 | 0.013 | 0.004 | 0.000 | 0.000 |
| Firmicutes | Clostridia | Clostridiales | Ruminococcaceae | *Anaerotruncus* | 0.257 | 0.383 | 0.180 | 0.110 | 0.159 | 0.071 |
| Firmicutes | Clostridia | Clostridiales | Ruminococcaceae | *Ethanoligenens* | 0.001 | 0.000 | 0.000 | 0.000 | 0.000 | 0.000 |
| Firmicutes | Clostridia | Clostridiales | Ruminococcaceae | *Flavonifractor* | 0.001 | 0.000 | 0.000 | 0.000 | 0.000 | 0.000 |
| Firmicutes | Clostridia | Clostridiales | Ruminococcaceae | *Harryflintia* | 0.001 | 0.032 | 0.000 | 0.000 | 0.000 | 0.001 |
| Firmicutes | Clostridia | Clostridiales | Ruminococcaceae | *Herbivorax* | 0.001 | 0.000 | 0.000 | 0.000 | 0.000 | 0.000 |
| Firmicutes | Clostridia | Clostridiales | Ruminococcaceae | *Negativibacillus* | 0.001 | 0.000 | 0.000 | 0.000 | 0.000 | 0.000 |
| Firmicutes | Clostridia | Clostridiales | Ruminococcaceae | *Phocea* | 0.006 | 0.000 | 0.000 | 0.004 | 0.000 | 0.005 |
| Firmicutes | Clostridia | Clostridiales | Ruminococcaceae | *Pseudoflavonifractor* | 0.000 | 0.000 | 0.000 | 0.004 | 0.000 | 0.000 |
| Firmicutes | Clostridia | Clostridiales | Ruminococcaceae | *Ruminococcus* | 0.001 | 0.000 | 0.013 | 0.000 | 0.000 | 0.000 |
| Firmicutes | Clostridia | Clostridiales | Ruminococcaceae | *Ruthenibacterium* | 0.001 | 0.000 | 0.000 | 0.000 | 0.000 | 0.000 |
| Firmicutes | Clostridia | Clostridiales | Ruminococcaceae | *Sporobacter* | 0.010 | 0.000 | 0.000 | 0.004 | 0.000 | 0.001 |
| Firmicutes | Clostridia | Clostridiales | Syntrophomonadaceae | *Pelospora* | 0.009 | 0.000 | 0.000 | 0.000 | 0.000 | 0.000 |
| Firmicutes | Clostridia | Clostridiales | Unclassified | *Fenollaria* | 0.000 | 0.032 | 0.000 | 0.000 | 0.000 | 0.000 |
| Firmicutes | Clostridia | Clostridiales | Unclassified | *Intestinimonas* | 0.001 | 0.000 | 0.000 | 0.000 | 0.000 | 0.000 |
| Firmicutes | Clostridia | Clostridiales | Vallitaleaceae | *Vallitalea* | 0.001 | 0.000 | 0.000 | 0.004 | 0.000 | 0.000 |
| Firmicutes | Clostridia | Thermoanaerobacterales | Family III. Incertae Sedis | *Thermosediminibacter* | 0.002 | 0.000 | 0.000 | 0.000 | 0.000 | 0.000 |
| Firmicutes | Clostridia | Thermoanaerobacterales | Thermoanaerobacteraceae | *Fervidicola* | 0.002 | 0.000 | 0.000 | 0.000 | 0.000 | 0.000 |
| Firmicutes | Clostridia | Thermoanaerobacterales | Thermoanaerobacteraceae | *Thermoanaerobacter* | 0.001 | 0.000 | 0.000 | 0.000 | 0.000 | 0.001 |
| Firmicutes | Clostridia | Thermoanaerobacterales | Thermodesulfobiaceae | *Coprothermobacter* | 0.004 | 0.000 | 0.000 | 0.004 | 0.000 | 0.001 |
| Firmicutes | Clostridia 1 | Clostridiales | Clostridiaceae 1 | Arthropod cluster | 0.004 | 0.000 | 0.000 | 0.007 | 0.000 | 0.003 |
| Firmicutes | Clostridia 1 | Clostridiales | Clostridiaceae 1 | *Clostridium* 1 | 0.016 | 0.032 | 0.013 | 0.164 | 0.004 | 0.003 |
| Firmicutes | Clostridia 1 | Clostridiales | Clostridiaceae 1 | *Clostridium* 10 | 0.000 | 0.000 | 0.000 | 0.000 | 0.000 | 0.001 |
| Firmicutes | Clostridia 1 | Clostridiales | Clostridiaceae 1 | *Clostridium* 11 | 0.002 | 0.000 | 0.013 | 0.014 | 0.000 | 0.000 |
| Firmicutes | Clostridia 1 | Clostridiales | Clostridiaceae 1 | *Clostridium* 2 | 0.002 | 0.000 | 0.000 | 0.000 | 0.000 | 0.000 |

**Table S2 continued.**

| | | | | | | | | | | |
|---|---|---|---|---|---|---|---|---|---|---|
| Firmicutes | Clostridia 1 | Clostridiales | Clostridiaceae 1 | *Clostridium* 3 | 0.002 | 0.032 | 0.000 | 0.157 | 0.000 | 0.001 |
| Firmicutes | Clostridia 1 | Clostridiales | Clostridiaceae 1 | *Clostridium* 5 | 0.000 | 0.000 | 0.000 | 0.014 | 0.000 | 0.000 |
| Firmicutes | Clostridia 1 | Clostridiales | Clostridiaceae 1 | *Clostridium* 8 | 0.002 | 0.032 | 0.000 | 0.043 | 0.004 | 0.000 |
| Firmicutes | Clostridia 1 | Clostridiales | Clostridiaceae 1 | *Clostridium* 9 | 0.000 | 0.000 | 0.000 | 0.000 | 0.000 | 0.001 |
| Firmicutes | Clostridia 1 | Clostridiales | Clostridiaceae 1 | *Sarcina* | 0.001 | 0.000 | 0.000 | 0.011 | 0.004 | 0.002 |
| Firmicutes | Clostridia 1 | Clostridiales | Clostridiaceae 1 | Unclassified | 0.000 | 0.000 | 0.000 | 0.217 | 0.004 | 0.003 |
| Firmicutes | Clostridia 1 | Clostridiales | Clostridiaceae 1 | Unclassified | 0.002 | 0.000 | 0.000 | 0.000 | 0.000 | 0.000 |
| Firmicutes | Clostridia 1 | Clostridiales | Clostridiaceae 4 | *Caminicella* | 0.005 | 0.000 | 0.000 | 0.004 | 0.000 | 0.001 |
| Firmicutes | Clostridia 1 | Clostridiales | Eubacteriaceae 1 | *Acetobacterium* | 0.002 | 0.000 | 0.000 | 0.000 | 0.000 | 0.000 |
| Firmicutes | Clostridia 1 | Clostridiales | Eubacteriaceae 1 | *Anaerofustis* | 0.001 | 0.000 | 0.000 | 0.000 | 0.000 | 0.000 |
| Firmicutes | Clostridia 1 | Clostridiales | Eubacteriaceae 1 | *Eubacterium* 1 | 0.004 | 0.000 | 0.000 | 0.000 | 0.000 | 0.002 |
| Firmicutes | Clostridia 1 | Clostridiales | Eubacteriaceae 1 | *Eubacterium* 2 | 0.007 | 0.000 | 0.000 | 0.004 | 0.000 | 0.003 |
| Firmicutes | Clostridia 1 | Clostridiales | Eubacteriaceae 1 | *Pseudoramibacte*r | 0.002 | 0.000 | 0.000 | 0.000 | 0.000 | 0.000 |
| Firmicutes | Clostridia 1 | Clostridiales | Family XI Incertae Sedis | *Tissierella* 3 | 0.002 | 0.000 | 0.000 | 0.000 | 0.000 | 0.000 |
| Firmicutes | Clostridia 1 | Clostridiales | Family XII Incertae Sedis | *Acidaminobacter* | 0.001 | 0.000 | 0.000 | 0.000 | 0.000 | 0.000 |
| Firmicutes | Clostridia 1 | Clostridiales | Family XII Incertae Sedis | *Fusibacter* | 0.004 | 0.000 | 0.000 | 0.004 | 0.000 | 0.000 |
| Firmicutes | Clostridia 1 | Clostridiales | Family XIII Incertae Sedis | *Anaerovorax* 1 | 0.040 | 0.064 | 0.000 | 0.032 | 0.028 | 0.017 |
| Firmicutes | Clostridia 1 | Clostridiales | Family XIII Incertae Sedis | *Anaerovorax* 2 | 0.001 | 0.000 | 0.000 | 0.000 | 0.004 | 0.000 |
| Firmicutes | Clostridia 1 | Clostridiales | Family XIII Incertae Sedis | *Eubacterium* 3 | 0.031 | 0.000 | 0.116 | 0.068 | 0.134 | 0.041 |
| Firmicutes | Clostridia 1 | Clostridiales | Family XIII Incertae Sedis | Gut cluster 1 | 0.011 | 0.000 | 0.013 | 0.004 | 0.012 | 0.015 |
| Firmicutes | Clostridia 1 | Clostridiales | Family XIII Incertae Sedis | Gut cluster 2 | 0.001 | 0.000 | 0.000 | 0.004 | 0.000 | 0.000 |
| Firmicutes | Clostridia 1 | Clostridiales | Family XIII Incertae Sedis | Gut cluster 3 | 0.148 | 0.064 | 0.000 | 0.007 | 0.000 | 0.001 |
| Firmicutes | Clostridia 1 | Clostridiales | Family XIII Incertae Sedis | Gut cluster 4 | 0.000 | 0.000 | 0.000 | 0.000 | 0.069 | 0.044 |
| Firmicutes | Clostridia 1 | Clostridiales | Family XIII Incertae Sedis | *Mogibacterium* | 0.000 | 0.000 | 0.000 | 0.004 | 0.000 | 0.002 |
| Firmicutes | Clostridia 1 | Clostridiales | Family XIII Incertae Sedis | Termite cockroach cluster 1 | 0.548 | 0.447 | 0.026 | 0.207 | 0.293 | 0.226 |
| Firmicutes | Clostridia 1 | Clostridiales | Family XIII Incertae Sedis | Termite cockroach cluster 2 | 0.198 | 0.383 | 0.000 | 0.053 | 0.480 | 0.329 |
| Firmicutes | Clostridia 1 | Clostridiales | Family XIII Incertae Sedis | Unclassified | 0.000 | 0.000 | 0.000 | 0.004 | 0.000 | 0.000 |

**Table S2 continued.**

| | | | | | | | | | | |
|---|---|---|---|---|---|---|---|---|---|---|
| Firmicutes | Clostridia 1 | Clostridiales | Family XIII Incertae Sedis | Unclassified | 0.001 | 0.000 | 0.000 | 0.000 | 0.000 | 0.000 |
| Firmicutes | Clostridia 1 | Clostridiales | Gracilibacteraceae | Uncultured | 0.009 | 0.000 | 0.000 | 0.007 | 0.000 | 0.001 |
| Firmicutes | Clostridia 1 | Clostridiales | Lachnospiraceae | *Acetitomaculum* | 0.004 | 0.000 | 0.026 | 0.007 | 0.004 | 0.000 |
| Firmicutes | Clostridia 1 | Clostridiales | Lachnospiraceae | *Anaerostipes* | 0.000 | 0.032 | 0.000 | 0.011 | 0.000 | 0.000 |
| Firmicutes | Clostridia 1 | Clostridiales | Lachnospiraceae | *Blautia* | 0.046 | 0.000 | 0.387 | 0.025 | 0.008 | 0.007 |
| Firmicutes | Clostridia 1 | Clostridiales | Lachnospiraceae | *Butyrivibrio* 1 | 0.002 | 0.000 | 0.039 | 0.000 | 0.000 | 0.000 |
| Firmicutes | Clostridia 1 | Clostridiales | Lachnospiraceae | *Butyrivibrio* 2 | 0.009 | 0.000 | 0.039 | 0.000 | 0.000 | 0.000 |
| Firmicutes | Clostridia 1 | Clostridiales | Lachnospiraceae | *Butyrivibrio* 3 | 0.077 | 0.000 | 0.142 | 0.011 | 0.000 | 0.000 |
| Firmicutes | Clostridia 1 | Clostridiales | Lachnospiraceae | *Butyrivibrio* 4 | 0.048 | 0.000 | 0.000 | 0.000 | 0.000 | 0.000 |
| Firmicutes | Clostridia 1 | Clostridiales | Lachnospiraceae | *Butyrivibrio-Pseudobutyrivibrio* | 0.016 | 0.000 | 0.013 | 0.000 | 0.000 | 0.000 |
| Firmicutes | Clostridia 1 | Clostridiales | Lachnospiraceae | *Candidatus* Arthromitus | 4.450 | 3.923 | 0.026 | 1.564 | 0.216 | 0.076 |
| Firmicutes | Clostridia 1 | Clostridiales | Lachnospiraceae | *Catonella* | 0.001 | 0.000 | 0.000 | 0.000 | 0.000 | 0.000 |
| Firmicutes | Clostridia 1 | Clostridiales | Lachnospiraceae | *Clostridium piliforme* cluster | 0.054 | 0.032 | 0.000 | 0.011 | 0.004 | 0.000 |
| Firmicutes | Clostridia 1 | Clostridiales | Lachnospiraceae | *Coprococcus* 1 | 0.009 | 0.032 | 0.090 | 0.011 | 0.000 | 0.001 |
| Firmicutes | Clostridia 1 | Clostridiales | Lachnospiraceae | *Coprococcus* 2 | 0.005 | 0.032 | 0.000 | 0.000 | 0.000 | 0.000 |
| Firmicutes | Clostridia 1 | Clostridiales | Lachnospiraceae | *Dorea* | 0.005 | 0.000 | 0.168 | 0.018 | 0.000 | 0.001 |
| Firmicutes | Clostridia 1 | Clostridiales | Lachnospiraceae | Gut cluster 1 | 0.121 | 0.064 | 1.599 | 0.164 | 0.020 | 0.013 |
| Firmicutes | Clostridia 1 | Clostridiales | Lachnospiraceae | Gut cluster 10 | 0.010 | 0.000 | 0.322 | 0.000 | 0.000 | 0.001 |
| Firmicutes | Clostridia 1 | Clostridiales | Lachnospiraceae | Gut cluster 11 | 0.043 | 0.032 | 0.168 | 0.011 | 0.012 | 0.000 |
| Firmicutes | Clostridia 1 | Clostridiales | Lachnospiraceae | Gut cluster 13 | 1.882 | 2.105 | 0.026 | 0.684 | 0.980 | 0.342 |
| Firmicutes | Clostridia 1 | Clostridiales | Lachnospiraceae | Gut cluster 14 | 0.320 | 0.893 | 0.000 | 0.338 | 0.028 | 0.026 |
| Firmicutes | Clostridia 1 | Clostridiales | Lachnospiraceae | Gut cluster 15 | 0.174 | 0.223 | 0.000 | 0.192 | 0.179 | 0.045 |
| Firmicutes | Clostridia 1 | Clostridiales | Lachnospiraceae | Gut cluster 2 | 0.001 | 0.000 | 0.000 | 0.000 | 0.000 | 0.000 |
| Firmicutes | Clostridia 1 | Clostridiales | Lachnospiraceae | Gut cluster 3 | 0.064 | 0.032 | 0.064 | 0.011 | 0.008 | 0.010 |
| Firmicutes | Clostridia 1 | Clostridiales | Lachnospiraceae | Gut cluster 4 | 0.000 | 0.000 | 0.155 | 0.000 | 0.004 | 0.000 |
| Firmicutes | Clostridia 1 | Clostridiales | Lachnospiraceae | Gut cluster 5 | 0.017 | 0.064 | 0.129 | 0.153 | 0.000 | 0.001 |
| Firmicutes | Clostridia 1 | Clostridiales | Lachnospiraceae | Gut cluster 6 | 0.153 | 0.287 | 0.748 | 0.043 | 0.004 | 0.001 |

**Table S2 continued.**

| | | | | | | | | | | |
|---|---|---|---|---|---|---|---|---|---|---|
| Firmicutes | Clostridia 1 | Clostridiales | Lachnospiraceae | Gut cluster 7 | 0.004 | 0.000 | 0.013 | 0.004 | 0.000 | 0.003 |
| Firmicutes | Clostridia 1 | Clostridiales | Lachnospiraceae | Gut cluster 8 | 0.017 | 0.000 | 0.090 | 0.000 | 0.000 | 0.003 |
| Firmicutes | Clostridia 1 | Clostridiales | Lachnospiraceae | Gut cluster 9 | 0.020 | 0.000 | 0.387 | 0.007 | 0.000 | 0.000 |
| Firmicutes | Clostridia 1 | Clostridiales | Lachnospiraceae | *Hespellia* | 0.000 | 0.000 | 0.026 | 0.000 | 0.000 | 0.000 |
| Firmicutes | Clostridia 1 | Clostridiales | Lachnospiraceae | Incertae Sedis 1 | 0.014 | 0.000 | 0.077 | 0.011 | 0.004 | 0.001 |
| Firmicutes | Clostridia 1 | Clostridiales | Lachnospiraceae | Incertae Sedis 10 | 0.045 | 0.064 | 0.052 | 0.014 | 0.000 | 0.000 |
| Firmicutes | Clostridia 1 | Clostridiales | Lachnospiraceae | Incertae Sedis 11 | 0.083 | 0.032 | 1.689 | 0.061 | 0.020 | 0.002 |
| Firmicutes | Clostridia 1 | Clostridiales | Lachnospiraceae | Incertae Sedis 12 | 0.020 | 0.000 | 0.064 | 0.004 | 0.004 | 0.000 |
| Firmicutes | Clostridia 1 | Clostridiales | Lachnospiraceae | Incertae Sedis 13 | 0.054 | 0.000 | 0.013 | 0.000 | 0.000 | 0.001 |
| Firmicutes | Clostridia 1 | Clostridiales | Lachnospiraceae | Incertae Sedis 14 | 1.093 | 0.000 | 0.039 | 0.025 | 0.008 | 0.002 |
| Firmicutes | Clostridia 1 | Clostridiales | Lachnospiraceae | Incertae Sedis 15 | 0.030 | 0.032 | 0.013 | 0.004 | 0.000 | 0.000 |
| Firmicutes | Clostridia 1 | Clostridiales | Lachnospiraceae | Incertae Sedis 16 | 0.019 | 0.000 | 0.052 | 0.011 | 0.000 | 0.005 |
| Firmicutes | Clostridia 1 | Clostridiales | Lachnospiraceae | Incertae Sedis 17 | 0.011 | 0.032 | 0.052 | 0.000 | 0.000 | 0.005 |
| Firmicutes | Clostridia 1 | Clostridiales | Lachnospiraceae | Incertae Sedis 18 | 0.000 | 0.000 | 0.013 | 0.000 | 0.000 | 0.000 |
| Firmicutes | Clostridia 1 | Clostridiales | Lachnospiraceae | Incertae Sedis 19 | 0.056 | 0.096 | 0.232 | 0.011 | 0.004 | 0.000 |
| Firmicutes | Clostridia 1 | Clostridiales | Lachnospiraceae | Incertae Sedis 2 | 0.000 | 0.000 | 0.013 | 0.000 | 0.000 | 0.000 |
| Firmicutes | Clostridia 1 | Clostridiales | Lachnospiraceae | Incertae Sedis 20 | 0.002 | 0.000 | 0.013 | 0.000 | 0.000 | 0.000 |
| Firmicutes | Clostridia 1 | Clostridiales | Lachnospiraceae | Incertae Sedis 21 | 0.025 | 0.000 | 0.064 | 0.006 | 0.004 | 0.000 |
| Firmicutes | Clostridia 1 | Clostridiales | Lachnospiraceae | Incertae Sedis 22 | 0.012 | 0.000 | 0.000 | 0.000 | 0.000 | 0.000 |
| Firmicutes | Clostridia 1 | Clostridiales | Lachnospiraceae | Incertae Sedis 23 | 0.042 | 0.032 | 0.103 | 0.014 | 0.000 | 0.000 |
| Firmicutes | Clostridia 1 | Clostridiales | Lachnospiraceae | Incertae Sedis 25 | 0.020 | 0.032 | 0.413 | 0.014 | 0.000 | 0.003 |
| Firmicutes | Clostridia 1 | Clostridiales | Lachnospiraceae | Incertae Sedis 26 | 0.004 | 0.000 | 0.000 | 0.000 | 0.000 | 0.000 |
| Firmicutes | Clostridia 1 | Clostridiales | Lachnospiraceae | Incertae Sedis 27 | 0.007 | 0.000 | 0.013 | 0.000 | 0.000 | 0.000 |
| Firmicutes | Clostridia 1 | Clostridiales | Lachnospiraceae | Incertae Sedis 28 | 0.006 | 0.064 | 0.000 | 0.029 | 0.000 | 0.001 |
| Firmicutes | Clostridia 1 | Clostridiales | Lachnospiraceae | Incertae Sedis 3 | 0.001 | 0.000 | 0.013 | 0.004 | 0.000 | 0.001 |
| Firmicutes | Clostridia 1 | Clostridiales | Lachnospiraceae | Incertae Sedis 30 | 0.005 | 0.064 | 0.000 | 0.004 | 0.000 | 0.000 |
| Firmicutes | Clostridia 1 | Clostridiales | Lachnospiraceae | Incertae Sedis 31 | 0.000 | 0.000 | 0.193 | 0.036 | 0.000 | 0.000 |

**Table S2 continued.**

| | | | | | | | | | | |
|---|---|---|---|---|---|---|---|---|---|---|
| Firmicutes | Clostridia 1 | Clostridiales | Lachnospiraceae | Incertae Sedis 32 | 0.012 | 0.032 | 0.000 | 0.043 | 0.000 | 0.001 |
| Firmicutes | Clostridia 1 | Clostridiales | Lachnospiraceae | Incertae Sedis 33 | 0.001 | 0.000 | 0.000 | 0.004 | 0.000 | 0.000 |
| Firmicutes | Clostridia 1 | Clostridiales | Lachnospiraceae | Incertae Sedis 34 | 0.032 | 0.159 | 0.013 | 0.039 | 0.008 | 0.003 |
| Firmicutes | Clostridia 1 | Clostridiales | Lachnospiraceae | Incertae Sedis 35 | 0.001 | 0.032 | 0.026 | 0.000 | 0.000 | 0.000 |
| Firmicutes | Clostridia 1 | Clostridiales | Lachnospiraceae | Incertae Sedis 5 | 0.001 | 0.000 | 0.000 | 0.000 | 0.000 | 0.000 |
| Firmicutes | Clostridia 1 | Clostridiales | Lachnospiraceae | Incertae Sedis 6 | 0.000 | 0.000 | 0.013 | 0.000 | 0.000 | 0.000 |
| Firmicutes | Clostridia 1 | Clostridiales | Lachnospiraceae | Incertae Sedis 8 | 0.020 | 0.000 | 0.039 | 0.007 | 0.004 | 0.001 |
| Firmicutes | Clostridia 1 | Clostridiales | Lachnospiraceae | Incertae Sedis 9 | 0.047 | 0.000 | 0.413 | 0.018 | 0.000 | 0.000 |
| Firmicutes | Clostridia 1 | Clostridiales | Lachnospiraceae | *Johnsonella* | 0.005 | 0.000 | 0.000 | 0.000 | 0.000 | 0.000 |
| Firmicutes | Clostridia 1 | Clostridiales | Lachnospiraceae | *Lachnospira* | 0.005 | 0.000 | 0.000 | 0.007 | 0.004 | 0.000 |
| Firmicutes | Clostridia 1 | Clostridiales | Lachnospiraceae | *Marvinbryantia* | 0.019 | 0.000 | 0.090 | 0.004 | 0.000 | 0.000 |
| Firmicutes | Clostridia 1 | Clostridiales | Lachnospiraceae | Mixed gut cluster | 0.059 | 0.000 | 0.077 | 0.000 | 0.000 | 0.000 |
| Firmicutes | Clostridia 1 | Clostridiales | Lachnospiraceae | *Oribacterium* | 0.000 | 0.032 | 0.039 | 0.000 | 0.000 | 0.000 |
| Firmicutes | Clostridia 1 | Clostridiales | Lachnospiraceae | *Parasporobacterium-Sporobacterium* | 0.080 | 0.096 | 0.013 | 0.064 | 0.000 | 0.000 |
| Firmicutes | Clostridia 1 | Clostridiales | Lachnospiraceae | *Robinsoniella* Insects | 0.001 | 0.000 | 0.026 | 0.004 | 0.000 | 0.000 |
| Firmicutes | Clostridia 1 | Clostridiales | Lachnospiraceae | *Roseburia* 1 | 0.028 | 0.000 | 0.052 | 0.004 | 0.000 | 0.001 |
| Firmicutes | Clostridia 1 | Clostridiales | Lachnospiraceae | *Roseburia* 4 | 0.038 | 0.000 | 0.000 | 0.004 | 0.004 | 0.000 |
| Firmicutes | Clostridia 1 | Clostridiales | Lachnospiraceae | *Shuttleworthia* | 0.015 | 0.000 | 0.000 | 0.000 | 0.000 | 0.000 |
| Firmicutes | Clostridia 1 | Clostridiales | Lachnospiraceae | *Syntrophococcus* | 0.002 | 0.000 | 0.000 | 0.004 | 0.000 | 0.000 |
| Firmicutes | Clostridia 1 | Clostridiales | Lachnospiraceae | Termite cluster | 2.082 | 2.520 | 0.026 | 1.621 | 0.460 | 0.054 |
| Firmicutes | Clostridia 1 | Clostridiales | Lachnospiraceae | Termite cluster 2 | 0.515 | 0.351 | 0.219 | 0.096 | 0.004 | 0.005 |
| Firmicutes | Clostridia 1 | Clostridiales | Lachnospiraceae | Termite cockroach cluster | 0.584 | 0.032 | 0.000 | 0.360 | 0.004 | 0.002 |
| Firmicutes | Clostridia 1 | Clostridiales | Lachnospiraceae | Unclassified | 0.000 | 0.191 | 0.168 | 0.025 | 0.024 | 0.016 |
| Firmicutes | Clostridia 1 | Clostridiales | Lachnospiraceae | Uncultured 1 | 0.381 | 0.000 | 0.000 | 0.000 | 0.000 | 0.000 |
| Firmicutes | Clostridia 1 | Clostridiales | Lachnospiraceae | Uncultured 1 | 0.000 | 0.196 | 3.042 | 0.143 | 0.024 | 0.000 |
| Firmicutes | Clostridia 1 | Clostridiales | Lachnospiraceae | Uncultured 1 | 0.000 | 0.000 | 0.000 | 0.000 | 0.000 | 0.005 |
| Firmicutes | Clostridia 1 | Clostridiales | Lachnospiraceae | Uncultured 10 | 0.012 | 0.000 | 0.026 | 0.000 | 0.000 | 0.000 |

**Table S2 continued.**

| | | | | | | | | | | |
|---|---|---|---|---|---|---|---|---|---|---|
| Firmicutes | Clostridia 1 | Clostridiales | Lachnospiraceae | Uncultured 12 | 0.001 | 0.000 | 0.000 | 0.000 | 0.000 | 0.000 |
| Firmicutes | Clostridia 1 | Clostridiales | Lachnospiraceae | Uncultured 13 | 0.004 | 0.000 | 0.168 | 0.004 | 0.114 | 0.002 |
| Firmicutes | Clostridia 1 | Clostridiales | Lachnospiraceae | Uncultured 15 | 0.005 | 0.000 | 0.026 | 0.004 | 0.000 | 0.000 |
| Firmicutes | Clostridia 1 | Clostridiales | Lachnospiraceae | Uncultured 16 | 0.001 | 0.000 | 0.000 | 0.004 | 0.000 | 0.000 |
| Firmicutes | Clostridia 1 | Clostridiales | Lachnospiraceae | Uncultured 17 | 0.004 | 0.000 | 0.026 | 0.004 | 0.000 | 0.000 |
| Firmicutes | Clostridia 1 | Clostridiales | Lachnospiraceae | Uncultured 18 | 0.001 | 0.000 | 0.000 | 0.000 | 0.000 | 0.000 |
| Firmicutes | Clostridia 1 | Clostridiales | Lachnospiraceae | Uncultured 19 | 0.000 | 0.000 | 0.000 | 0.000 | 0.000 | 0.001 |
| Firmicutes | Clostridia 1 | Clostridiales | Lachnospiraceae | Uncultured 25 | 0.001 | 0.000 | 0.026 | 0.000 | 0.000 | 0.000 |
| Firmicutes | Clostridia 1 | Clostridiales | Lachnospiraceae | Uncultured 27 | 0.017 | 0.000 | 0.052 | 0.000 | 0.004 | 0.001 |
| Firmicutes | Clostridia 1 | Clostridiales | Lachnospiraceae | Uncultured 28 | 0.014 | 0.032 | 0.064 | 0.004 | 0.000 | 0.000 |
| Firmicutes | Clostridia 1 | Clostridiales | Lachnospiraceae | Uncultured 29 | 0.002 | 0.000 | 0.013 | 0.004 | 0.000 | 0.000 |
| Firmicutes | Clostridia 1 | Clostridiales | Lachnospiraceae | Uncultured 30 | 0.017 | 0.000 | 0.297 | 0.004 | 0.000 | 0.000 |
| Firmicutes | Clostridia 1 | Clostridiales | Lachnospiraceae | Uncultured 31 | 0.558 | 0.638 | 1.921 | 0.306 | 0.020 | 0.000 |
| Firmicutes | Clostridia 1 | Clostridiales | Lachnospiraceae | Uncultured 33 | 0.015 | 0.000 | 0.013 | 0.000 | 0.000 | 0.000 |
| Firmicutes | Clostridia 1 | Clostridiales | Lachnospiraceae | Uncultured 34 | 0.001 | 0.000 | 0.000 | 0.000 | 0.000 | 0.000 |
| Firmicutes | Clostridia 1 | Clostridiales | Lachnospiraceae | Uncultured 37 | 0.002 | 0.000 | 0.000 | 0.000 | 0.000 | 0.000 |
| Firmicutes | Clostridia 1 | Clostridiales | Lachnospiraceae | Uncultured 40 | 0.004 | 0.000 | 0.026 | 0.000 | 0.000 | 0.000 |
| Firmicutes | Clostridia 1 | Clostridiales | Lachnospiraceae | Uncultured 41 | 0.001 | 0.000 | 0.013 | 0.004 | 0.000 | 0.000 |
| Firmicutes | Clostridia 1 | Clostridiales | Lachnospiraceae | Uncultured 42 | 0.001 | 0.000 | 0.026 | 0.000 | 0.000 | 0.001 |
| Firmicutes | Clostridia 1 | Clostridiales | Lachnospiraceae | Uncultured 43 | 0.006 | 0.032 | 0.077 | 0.007 | 0.000 | 0.000 |
| Firmicutes | Clostridia 1 | Clostridiales | Lachnospiraceae | Uncultured 44 | 0.023 | 0.064 | 0.374 | 0.007 | 0.000 | 0.000 |
| Firmicutes | Clostridia 1 | Clostridiales | Lachnospiraceae | Uncultured 45 | 0.005 | 0.000 | 0.116 | 0.007 | 0.000 | 0.001 |
| Firmicutes | Clostridia 1 | Clostridiales | Lachnospiraceae | Uncultured 48 | 0.002 | 0.000 | 0.026 | 0.011 | 0.000 | 0.000 |
| Firmicutes | Clostridia 1 | Clostridiales | Lachnospiraceae | Uncultured 5 | 0.000 | 0.000 | 0.000 | 0.000 | 0.000 | 0.001 |
| Firmicutes | Clostridia 1 | Clostridiales | Lachnospiraceae | Uncultured 50 | 0.016 | 0.032 | 0.000 | 0.007 | 0.000 | 0.000 |
| Firmicutes | Clostridia 1 | Clostridiales | Lachnospiraceae | Uncultured 53 | 0.002 | 0.000 | 0.013 | 0.000 | 0.000 | 0.000 |
| Firmicutes | Clostridia 1 | Clostridiales | Lachnospiraceae | Uncultured 54 | 0.767 | 0.255 | 0.039 | 0.171 | 0.000 | 0.000 |

**Table S2 continued.**

| | | | | | | | | | | |
|---|---|---|---|---|---|---|---|---|---|---|
| Firmicutes | Clostridia 1 | Clostridiales | Lachnospiraceae | Uncultured 55 | 0.004 | 0.000 | 0.013 | 0.000 | 0.000 | 0.000 |
| Firmicutes | Clostridia 1 | Clostridiales | Lachnospiraceae | Uncultured 56 | 0.000 | 0.032 | 0.000 | 0.000 | 0.000 | 0.000 |
| Firmicutes | Clostridia 1 | Clostridiales | Lachnospiraceae | Uncultured 57 | 0.014 | 0.000 | 0.000 | 0.004 | 0.004 | 0.000 |
| Firmicutes | Clostridia 1 | Clostridiales | Lachnospiraceae | Uncultured 58 | 0.002 | 0.000 | 0.000 | 0.011 | 0.000 | 0.000 |
| Firmicutes | Clostridia 1 | Clostridiales | Lachnospiraceae | Uncultured 60 | 0.009 | 0.052 | 0.000 | 0.000 | 0.000 | 0.001 |
| Firmicutes | Clostridia 1 | Clostridiales | Lachnospiraceae | Uncultured 61 | 0.016 | 0.032 | 0.000 | 0.000 | 0.000 | 0.000 |
| Firmicutes | Clostridia 1 | Clostridiales | Lachnospiraceae | Uncultured 62 | 0.058 | 0.064 | 0.000 | 0.029 | 0.000 | 0.000 |
| Firmicutes | Clostridia 1 | Clostridiales | Lachnospiraceae | Uncultured 63 | 0.001 | 0.000 | 0.083 | 0.000 | 0.000 | 0.001 |
| Firmicutes | Clostridia 1 | Clostridiales | Lachnospiraceae | Uncultured 64 | 0.006 | 0.000 | 0.000 | 0.000 | 0.000 | 0.000 |
| Firmicutes | Clostridia 1 | Clostridiales | Lachnospiraceae | Uncultured 66 | 0.009 | 0.064 | 0.000 | 0.011 | 0.008 | 0.006 |
| Firmicutes | Clostridia 1 | Clostridiales | Lachnospiraceae | Unclassified | 0.087 | 0.000 | 0.000 | 0.000 | 0.000 | 0.000 |
| Firmicutes | Clostridia 1 | Clostridiales | Peptostreptococcaceae | Incertae Sedis 1 | 0.009 | 0.032 | 0.000 | 0.004 | 0.004 | 0.007 |
| Firmicutes | Clostridia 1 | Clostridiales | Peptostreptococcaceae | Incertae Sedis 2 | 0.001 | 0.000 | 0.000 | 0.000 | 0.004 | 0.001 |
| Firmicutes | Clostridia 1 | Clostridiales | Peptostreptococcaceae | Incertae Sedis 3 | 0.001 | 0.000 | 0.000 | 0.000 | 0.000 | 0.000 |
| Firmicutes | Clostridia 1 | Clostridiales | Peptostreptococcaceae | Incertae Sedis 5 | 0.001 | 0.000 | 0.000 | 0.000 | 0.000 | 0.000 |
| Firmicutes | Clostridia 1 | Clostridiales | Peptostreptococcaceae | Uncultured 3 | 0.007 | 0.000 | 0.000 | 0.004 | 0.000 | 0.001 |
| Firmicutes | Clostridia 1 | Clostridiales | Ruminococcaceae | *Acetanaerobacterium* | 0.019 | 0.032 | 0.000 | 0.004 | 0.004 | 0.003 |
| Firmicutes | Clostridia 1 | Clostridiales | Ruminococcaceae | *Acetivibrio* | 0.000 | 0.000 | 0.000 | 0.000 | 0.000 | 0.001 |
| Firmicutes | Clostridia 1 | Clostridiales | Ruminococcaceae | *Faecalibacterium* | 0.012 | 0.000 | 0.000 | 0.007 | 0.000 | 0.005 |
| Firmicutes | Clostridia 1 | Clostridiales | Ruminococcaceae | *Fastidiosipila* | 0.016 | 0.032 | 0.000 | 0.000 | 0.012 | 0.001 |
| Firmicutes | Clostridia 1 | Clostridiales | Ruminococcaceae | Gut cluster 1 | 0.108 | 0.064 | 0.013 | 0.050 | 0.061 | 0.070 |
| Firmicutes | Clostridia 1 | Clostridiales | Ruminococcaceae | Gut cluster 10 | 0.005 | 0.032 | 0.013 | 0.011 | 0.008 | 0.000 |
| Firmicutes | Clostridia 1 | Clostridiales | Ruminococcaceae | Gut cluster 3 | 0.142 | 0.478 | 0.026 | 0.053 | 0.110 | 0.036 |
| Firmicutes | Clostridia 1 | Clostridiales | Ruminococcaceae | Gut cluster 4 | 0.000 | 0.000 | 0.000 | 0.000 | 0.012 | 0.009 |
| Firmicutes | Clostridia 1 | Clostridiales | Ruminococcaceae | Gut cluster 5 | 0.716 | 0.734 | 0.000 | 0.011 | 0.525 | 0.436 |
| Firmicutes | Clostridia 1 | Clostridiales | Ruminococcaceae | Gut cluster 6 | 0.126 | 0.223 | 0.000 | 0.043 | 0.329 | 0.456 |
| Firmicutes | Clostridia 1 | Clostridiales | Ruminococcaceae | Gut cluster 7 | 0.778 | 1.946 | 0.013 | 0.784 | 0.020 | 0.005 |

**Table S2 continued.**

| | | | | | | | | | | |
|---|---|---|---|---|---|---|---|---|---|---|
| Firmicutes | Clostridia 1 | Clostridiales | Ruminococcaceae | Gut cluster 8 | 2.860 | 2.297 | 0.013 | 0.691 | 2.664 | 1.260 |
| Firmicutes | Clostridia 1 | Clostridiales | Ruminococcaceae | Gut cluster 9 | 0.113 | 0.255 | 0.000 | 0.039 | 0.061 | 0.024 |
| Firmicutes | Clostridia 1 | Clostridiales | Ruminococcaceae | *Hydrogenoanaerobacterium* | 0.173 | 0.191 | 0.000 | 0.053 | 0.077 | 0.028 |
| Firmicutes | Clostridia 1 | Clostridiales | Ruminococcaceae | Incertae Sedis 1 | 0.078 | 0.255 | 0.013 | 0.132 | 0.000 | 0.000 |
| Firmicutes | Clostridia 1 | Clostridiales | Ruminococcaceae | Incertae Sedis 10 | 0.001 | 0.000 | 0.000 | 0.000 | 0.000 | 0.000 |
| Firmicutes | Clostridia 1 | Clostridiales | Ruminococcaceae | Incertae Sedis 2 | 0.007 | 0.000 | 0.000 | 0.007 | 0.004 | 0.000 |
| Firmicutes | Clostridia 1 | Clostridiales | Ruminococcaceae | Incertae Sedis 3 | 0.006 | 0.000 | 0.052 | 0.000 | 0.000 | 0.005 |
| Firmicutes | Clostridia 1 | Clostridiales | Ruminococcaceae | Incertae Sedis 5 | 0.064 | 0.000 | 0.064 | 0.039 | 0.195 | 0.094 |
| Firmicutes | Clostridia 1 | Clostridiales | Ruminococcaceae | Incertae Sedis 6 | 0.057 | 0.064 | 0.000 | 0.021 | 0.049 | 0.025 |
| Firmicutes | Clostridia 1 | Clostridiales | Ruminococcaceae | Incertae Sedis 7 | 0.002 | 0.032 | 0.000 | 0.004 | 0.004 | 0.000 |
| Firmicutes | Clostridia 1 | Clostridiales | Ruminococcaceae | Insect cluster | 1.516 | 2.169 | 0.219 | 0.360 | 0.256 | 0.000 |
| Firmicutes | Clostridia 1 | Clostridiales | Ruminococcaceae | Insect cluster | 0.000 | 0.000 | 0.000 | 0.000 | 0.000 | 0.192 |
| Firmicutes | Clostridia 1 | Clostridiales | Ruminococcaceae | *Papillibacter* | 0.145 | 0.191 | 0.000 | 0.050 | 0.073 | 0.092 |
| Firmicutes | Clostridia 1 | Clostridiales | Ruminococcaceae | *Ruminococcus* 1 | 0.032 | 0.032 | 0.013 | 0.000 | 0.016 | 0.003 |
| Firmicutes | Clostridia 1 | Clostridiales | Ruminococcaceae | *Ruminococcus* 2 | 0.002 | 0.000 | 0.000 | 0.000 | 0.000 | 0.001 |
| Firmicutes | Clostridia 1 | Clostridiales | Ruminococcaceae | *Subdoligranulum* | 0.007 | 0.032 | 0.000 | 0.000 | 0.000 | 0.001 |
| Firmicutes | Clostridia 1 | Clostridiales | Ruminococcaceae | Termite cockroach cluster | 1.150 | 1.754 | 0.026 | 0.338 | 1.403 | 0.535 |
| Firmicutes | Clostridia 1 | Clostridiales | Ruminococcaceae | Termite group aaa | 0.136 | 0.191 | 0.000 | 0.050 | 0.016 | 0.029 |
| Firmicutes | Clostridia 1 | Clostridiales | Ruminococcaceae | Unclassified | 0.000 | 0.223 | 0.000 | 0.075 | 0.155 | 0.063 |
| Firmicutes | Clostridia 1 | Clostridiales | Ruminococcaceae | Uncultured 10 | 0.000 | 0.000 | 0.000 | 0.000 | 0.004 | 0.000 |
| Firmicutes | Clostridia 1 | Clostridiales | Ruminococcaceae | Uncultured 12 | 0.069 | 0.319 | 0.052 | 0.061 | 0.037 | 0.017 |
| Firmicutes | Clostridia 1 | Clostridiales | Ruminococcaceae | Uncultured 14 | 0.001 | 0.000 | 0.000 | 0.000 | 0.000 | 0.000 |
| Firmicutes | Clostridia 1 | Clostridiales | Ruminococcaceae | Uncultured 19 | 0.002 | 0.000 | 0.000 | 0.007 | 0.024 | 0.002 |
| Firmicutes | Clostridia 1 | Clostridiales | Ruminococcaceae | Uncultured 2 | 0.001 | 0.000 | 0.000 | 0.004 | 0.000 | 0.000 |
| Firmicutes | Clostridia 1 | Clostridiales | Ruminococcaceae | Uncultured 20 | 0.026 | 0.096 | 0.000 | 0.018 | 0.037 | 0.031 |
| Firmicutes | Clostridia 1 | Clostridiales | Ruminococcaceae | Uncultured 21 | 0.001 | 0.000 | 0.000 | 0.000 | 0.000 | 0.000 |
| Firmicutes | Clostridia 1 | Clostridiales | Ruminococcaceae | Uncultured 24 | 0.192 | 0.128 | 0.000 | 0.043 | 0.081 | 0.041 |

**Table S2 continued.**

| | | | | | | | | | | |
|---|---|---|---|---|---|---|---|---|---|---|
| Firmicutes | Clostridia 1 | Clostridiales | Ruminococcaceae | Uncultured 26 | 0.002 | 0.000 | 0.000 | 0.000 | 0.000 | 0.000 |
| Firmicutes | Clostridia 1 | Clostridiales | Ruminococcaceae | Uncultured 27 | 0.001 | 0.000 | 0.000 | 0.000 | 0.000 | 0.002 |
| Firmicutes | Clostridia 1 | Clostridiales | Ruminococcaceae | Uncultured 28 | 0.069 | 0.000 | 0.000 | 0.021 | 0.045 | 0.037 |
| Firmicutes | Clostridia 1 | Clostridiales | Ruminococcaceae | Uncultured 29 | 0.110 | 0.032 | 0.000 | 0.068 | 0.000 | 0.000 |
| Firmicutes | Clostridia 1 | Clostridiales | Ruminococcaceae | Uncultured 3 | 0.000 | 0.000 | 0.000 | 0.000 | 0.008 | 0.001 |
| Firmicutes | Clostridia 1 | Clostridiales | Ruminococcaceae | Uncultured 31 | 0.000 | 0.000 | 0.000 | 0.007 | 0.004 | 0.001 |
| Firmicutes | Clostridia 1 | Clostridiales | Ruminococcaceae | Uncultured 32 | 0.000 | 0.000 | 0.000 | 0.000 | 0.004 | 0.000 |
| Firmicutes | Clostridia 1 | Clostridiales | Ruminococcaceae | Uncultured 33 | 0.059 | 0.032 | 0.000 | 0.011 | 0.069 | 0.026 |
| Firmicutes | Clostridia 1 | Clostridiales | Ruminococcaceae | Uncultured 34 | 0.004 | 0.000 | 0.000 | 0.011 | 0.028 | 0.014 |
| Firmicutes | Clostridia 1 | Clostridiales | Ruminococcaceae | Uncultured 5 | 0.001 | 0.000 | 0.000 | 0.000 | 0.000 | 0.001 |
| Firmicutes | Clostridia 1 | Clostridiales | Ruminococcaceae | Uncultured 6 | 0.004 | 0.000 | 0.000 | 0.000 | 0.000 | 0.000 |
| Firmicutes | Clostridia 1 | Clostridiales | Ruminococcaceae | Uncultured 8 | 0.000 | 0.000 | 0.000 | 0.000 | 0.012 | 0.005 |
| Firmicutes | Clostridia 1 | Clostridiales | Ruminococcaceae | Uncultured 8 | 0.000 | 0.000 | 0.000 | 0.000 | 0.000 | 0.000 |
| Firmicutes | Clostridia 1 | Clostridiales | Ruminococcaceae | Uncultured 9 | 0.004 | 0.000 | 0.000 | 0.000 | 0.004 | 0.001 |
| Firmicutes | Clostridia 1 | Clostridiales | Ruminococcaceae | Unclassified | 0.108 | 0.000 | 0.000 | 0.000 | 0.000 | 0.000 |
| Firmicutes | Clostridia 1 | Clostridiales | Unclassified | Unclassified | 0.000 | 0.032 | 0.000 | 0.000 | 0.000 | 0.000 |
| Firmicutes | Clostridia 1 | Thermoanaerobacterales | Unclassified | Unclassified | 0.000 | 0.032 | 0.000 | 0.004 | 0.000 | 0.000 |
| Firmicutes | Clostridia 2 | Clostridiales | Family XVI Incertae Sedis | *Carboxydocella* | 0.000 | 0.032 | 0.000 | 0.000 | 0.000 | 0.000 |
| Firmicutes | Clostridia 2 | Clostridiales | Syntrophomonadaceae | *Syntrophomonas* 2 | 0.001 | 0.000 | 0.000 | 0.000 | 0.000 | 0.000 |
| Firmicutes | Clostridia 2 | Clostridiales | Syntrophomonadaceae | Unclassified | 0.001 | 0.000 | 0.000 | 0.000 | 0.000 | 0.000 |
| Firmicutes | Clostridia 2 | Clostridiales 1 | Heliobacteriaceae | Uncultured | 0.000 | 0.000 | 0.000 | 0.000 | 0.004 | 0.000 |
| Firmicutes | Clostridia 2 | Clostridiales 1 | Heliobacteriaceae | Unclassified | 0.001 | 0.000 | 0.000 | 0.000 | 0.000 | 0.000 |
| Firmicutes | Clostridia 2 | Clostridiales 1 | OPB54 | Unclassified | 0.000 | 0.032 | 0.000 | 0.004 | 0.004 | 0.000 |
| Firmicutes | Clostridia 2 | Clostridiales 1 | OPB54 | Unclassified | 0.002 | 0.000 | 0.000 | 0.000 | 0.000 | 0.000 |
| Firmicutes | Clostridia 2 | Clostridiales 1 | Peptococcaceae 1 | *Thermincola* | 0.001 | 0.000 | 0.000 | 0.000 | 0.000 | 0.000 |
| Firmicutes | Clostridia 2 | Clostridiales 1 | Peptococcaceae 1 | Unclassified | 0.000 | 0.000 | 0.000 | 0.000 | 0.000 | 0.002 |
| Firmicutes | Clostridia 2 | Clostridiales 1 | Peptococcaceae 1 | Uncultured 2 | 0.074 | 0.159 | 0.000 | 0.029 | 0.077 | 0.032 |

**Table S2 continued.**

| | | | | | | | | | | |
|---|---|---|---|---|---|---|---|---|---|---|
| Firmicutes | Clostridia 2 | Clostridiales 1 | Peptococcaceae 1 | Uncultured 4 | 0.002 | 0.000 | 0.000 | 0.004 | 0.012 | 0.002 |
| Firmicutes | Clostridia 2 | Clostridiales 1 | Peptococcaceae 1 | Uncultured 5 | 0.308 | 0.383 | 0.090 | 0.082 | 0.393 | 0.159 |
| Firmicutes | Clostridia 2 | Clostridiales 1 | Peptococcaceae 1 | Unclassified | 0.001 | 0.000 | 0.000 | 0.000 | 0.000 | 0.000 |
| Firmicutes | Clostridia 2 | Clostridiales 1 | Peptococcaceae 2 | Uncultured gut Group A | 0.002 | 0.000 | 0.000 | 0.000 | 0.000 | 0.001 |
| Firmicutes | Clostridia 2 | Clostridiales 1 | Peptococcaceae 3 | *Desulfotomaculum* 1 | 0.001 | 0.000 | 0.000 | 0.004 | 0.000 | 0.000 |
| Firmicutes | Clostridia 2 | Clostridiales 1 | Peptococcaceae 3 | *Pelotomaculum* 3 | 0.004 | 0.000 | 0.000 | 0.000 | 0.000 | 0.001 |
| Firmicutes | Clostridia 2 | Clostridiales 1 | Peptococcaceae 3 | Unclassified | 0.000 | 0.000 | 0.013 | 0.000 | 0.000 | 0.000 |
| Firmicutes | Clostridia 2 | Clostridiales 1 | Peptococcaceae 3 | Unclassified | 0.002 | 0.000 | 0.000 | 0.000 | 0.000 | 0.000 |
| Firmicutes | Clostridia 2 | Clostridiales 1 | Veillonellaceae | *Acetonema* a | 0.001 | 0.000 | 0.000 | 0.007 | 0.000 | 0.000 |
| Firmicutes | Clostridia 2 | Clostridiales 1 | Veillonellaceae | *Anaeroarcus-Anaeromusa* | 0.000 | 0.000 | 0.000 | 0.000 | 0.000 | 0.001 |
| Firmicutes | Clostridia 2 | Clostridiales 1 | Veillonellaceae | *Dendrosporobacter* | 0.054 | 0.032 | 0.026 | 0.039 | 0.016 | 0.005 |
| Firmicutes | Clostridia 2 | Clostridiales 1 | Veillonellaceae | *Mitsuokella* | 0.000 | 0.000 | 0.000 | 0.000 | 0.000 | 0.001 |
| Firmicutes | Clostridia 2 | Clostridiales 1 | Veillonellaceae | *Succiniclasticum* | 0.000 | 0.000 | 0.000 | 0.000 | 0.004 | 0.000 |
| Firmicutes | Clostridia 2 | Clostridiales 1 | Veillonellaceae | *Thermosinus* | 0.157 | 0.096 | 0.013 | 0.135 | 0.000 | 0.001 |
| Firmicutes | Clostridia 2 | Clostridiales 1 | Veillonellaceae | Unclassified | 0.000 | 0.000 | 0.013 | 0.000 | 0.000 | 0.000 |
| Firmicutes | Clostridia 2 | Clostridiales 1 | Veillonellaceae | Uncultured 3 | 0.001 | 0.000 | 0.000 | 0.000 | 0.000 | 0.000 |
| Firmicutes | Clostridia 2 | Clostridiales 1 | Veillonellaceae | Uncultured 6 | 0.001 | 0.000 | 0.013 | 0.000 | 0.000 | 0.000 |
| Firmicutes | Clostridia 2 | Clostridiales 1 | Veillonellaceae | Uncultured 7 | 0.095 | 0.096 | 0.000 | 0.070 | 0.065 | 0.026 |
| Firmicutes | Clostridia 2 | Clostridiales 1 | Veillonellaceae | Uncultured 8 | 0.014 | 0.000 | 0.000 | 0.000 | 0.000 | 0.000 |
| Firmicutes | Clostridia 2 | Clostridiales 1 | Veillonellaceae | Unclassified | 0.001 | 0.000 | 0.000 | 0.000 | 0.000 | 0.000 |
| Firmicutes | Clostridia 2 | Clostridiales 1 | Veillonellaceae | *Veillonella* | 0.001 | 0.000 | 0.000 | 0.000 | 0.000 | 0.000 |
| Firmicutes | Clostridia 2 | Clostridiales 2 | Unclassified | Unclassified | 0.002 | 0.000 | 0.000 | 0.000 | 0.000 | 0.000 |
| Firmicutes | Clostridia 2 | Thermoanerobacterales | Thermoanaerobacteraceae | *Gelria* | 0.002 | 0.000 | 0.000 | 0.000 | 0.004 | 0.000 |
| Firmicutes | Clostridia 2 | Unclassified | Unclassified | Unclassified | 0.000 | 0.000 | 0.013 | 0.000 | 0.000 | 0.000 |
| Firmicutes | Clostridia 3 | Thermoanaerobacterales 3 | Thermoanaerobacteraceae | *Thermanaeromonas* | 0.000 | 0.000 | 0.000 | 0.000 | 0.000 | 0.001 |
| Firmicutes | Clostridia 3 | Thermoanaerobacterales 3 | Thermoanaerobacteraceae | *Thermoanaerobacter* 1 | 0.000 | 0.000 | 0.000 | 0.000 | 0.000 | 0.001 |

**Table S2 continued.**

| | | | | | | | | | | |
|---|---|---|---|---|---|---|---|---|---|---|
| Firmicutes | Clostridia 4 | Thermoanaerobacterales 4 | Family III Incertae Sedis | *Caldicellulosiruptor* | 0.001 | 0.000 | 0.000 | 0.000 | 0.000 | 0.000 |
| Firmicutes | Clostridia 5 | Thermoanaerobacterales 5 | Family III Incertae Sedis | *Tepidanaerobacter* | 0.000 | 0.000 | 0.000 | 0.000 | 0.004 | 0.000 |
| Firmicutes | Clostridia 5 | Thermoanaerobacterales 5 | Family III Incertae Sedis | Unclassified | 0.004 | 0.000 | 0.000 | 0.000 | 0.000 | 0.000 |
| Firmicutes | Clsotridia | Clostridiales | Unclassified | *Natranaerovirga* | 0.000 | 0.000 | 0.000 | 0.000 | 0.000 | 0.001 |
| Firmicutes | Clsotridia | Clsotridiales | Peptostreptococcaceae | *Romboutsia* | 0.000 | 0.000 | 0.000 | 0.000 | 0.000 | 0.001 |
| Firmicutes | Erysipelotrichi | Erysipelotrichales | Erysipelotrichaceae | *Allobaculum* | 0.000 | 0.000 | 0.000 | 0.000 | 0.000 | 0.002 |
| Firmicutes | Erysipelotrichi | Erysipelotrichales | Erysipelotrichaceae | *Asteroleplasma* | 0.043 | 0.000 | 0.013 | 0.004 | 0.000 | 0.000 |
| Firmicutes | Erysipelotrichi | Erysipelotrichales | Erysipelotrichaceae | *Coprobacillus* | 0.001 | 0.000 | 0.000 | 0.000 | 0.000 | 0.000 |
| Firmicutes | Erysipelotrichi | Erysipelotrichales | Erysipelotrichaceae | Incertae Sedis 8 | 0.273 | 0.032 | 0.632 | 0.100 | 0.016 | 0.014 |
| Firmicutes | Erysipelotrichi | Erysipelotrichales | Erysipelotrichaceae | *Solobacterium* | 0.004 | 0.000 | 0.000 | 0.000 | 0.000 | 0.001 |
| Firmicutes | Erysipelotrichi | Erysipelotrichales | Erysipelotrichaceae | *Turicibacter* | 0.002 | 0.000 | 0.013 | 0.011 | 0.004 | 0.000 |
| Firmicutes | Erysipelotrichi | Erysipelotrichales | Erysipelotrichaceae | Unclassified | 0.000 | 0.000 | 0.116 | 0.011 | 0.110 | 0.048 |
| Firmicutes | Erysipelotrichi | Erysipelotrichales | Erysipelotrichaceae | Uncultured 10 | 0.000 | 0.000 | 0.000 | 0.000 | 0.000 | 0.001 |
| Firmicutes | Erysipelotrichi | Erysipelotrichales | Erysipelotrichaceae | Uncultured 11 | 0.001 | 0.000 | 0.000 | 0.000 | 0.004 | 0.001 |
| Firmicutes | Erysipelotrichi | Erysipelotrichales | Erysipelotrichaceae | Uncultured 12 | 0.000 | 0.000 | 0.013 | 0.004 | 0.024 | 0.007 |
| Firmicutes | Erysipelotrichi | Erysipelotrichales | Erysipelotrichaceae | Uncultured 3 | 0.001 | 0.032 | 0.348 | 0.007 | 0.130 | 0.555 |
| Firmicutes | Erysipelotrichi | Erysipelotrichales | Erysipelotrichaceae | Uncultured 4 | 0.002 | 0.000 | 0.013 | 0.000 | 0.004 | 0.002 |
| Firmicutes | Erysipelotrichi | Erysipelotrichales | Erysipelotrichaceae | Uncultured 5 | 0.000 | 0.000 | 0.000 | 0.000 | 0.000 | 0.001 |
| Firmicutes | Erysipelotrichi | Erysipelotrichales | Erysipelotrichaceae | Uncultured 6 | 0.001 | 0.000 | 0.000 | 0.000 | 0.000 | 0.000 |
| Firmicutes | Erysipelotrichi | Erysipelotrichales | Erysipelotrichaceae | Uncultured 7 | 0.001 | 0.000 | 0.000 | 0.000 | 0.000 | 0.000 |
| Firmicutes | Erysipelotrichi | Erysipelotrichales | Erysipelotrichaceae | Unclassified | 0.020 | 0.000 | 0.000 | 0.000 | 0.000 | 0.000 |
| Firmicutes | Erysipelotrichia | Erysipelotrichales | Erysipelotrichaceae | *Breznakia* | 0.020 | 0.032 | 3.764 | 0.068 | 0.146 | 0.076 |
| Firmicutes | Erysipelotrichia | Erysipelotrichales | Erysipelotrichaceae | *Catenibacterium* | 0.001 | 0.000 | 0.013 | 0.004 | 0.000 | 0.000 |
| Firmicutes | Erysipelotrichia | Erysipelotrichales | Erysipelotrichaceae | *Catenisphaera* | 0.001 | 0.000 | 0.000 | 0.000 | 0.000 | 0.000 |
| Firmicutes | Erysipelotrichia | Erysipelotrichales | Erysipelotrichaceae | *Dielma* | 0.000 | 0.000 | 0.000 | 0.000 | 0.004 | 0.000 |
| Firmicutes | Erysipelotrichia | Erysipelotrichales | Erysipelotrichaceae | *Erysipelothrix* | 0.035 | 0.032 | 0.193 | 0.014 | 0.460 | 0.203 |

**Table S2 continued.**

| | | | | | | | | | | |
|---|---|---|---|---|---|---|---|---|---|---|
| Firmicutes | Erysipelotrichia | Erysipelotrichales | Erysipelotrichaceae | *Holdemania* | 0.001 | 0.000 | 0.000 | 0.000 | 0.000 | 0.000 |
| Firmicutes | Erysipelotrichia | Erysipelotrichales | Erysipelotrichaceae | Incertae Sedis 4 | 0.001 | 0.032 | 0.013 | 0.004 | 0.008 | 0.000 |
| Firmicutes | Erysipelotrichia | Erysipelotrichales | Erysipelotrichaceae | Incertae Sedis 5 | 0.011 | 0.000 | 0.155 | 0.011 | 0.016 | 0.002 |
| Firmicutes | Negativicutes | Selenemonadales | Sporomusaceae | *Desulfosporomusa* | 0.000 | 0.000 | 0.077 | 0.000 | 0.000 | 0.000 |
| Firmicutes | Negativicutes | Selenomonadales | Selenomonadaceae | *Selenomonas* | 0.000 | 0.000 | 0.000 | 0.004 | 0.000 | 0.001 |
| Firmicutes | Negativicutes | Selenomonadales | Sporomusaceae | *Anaerospora* | 0.000 | 0.000 | 0.013 | 0.011 | 0.000 | 0.001 |
| Firmicutes | Negativicutes | Selenomonadales | Sporomusaceae | *Propionispora* | 0.000 | 0.000 | 0.013 | 0.000 | 0.000 | 0.000 |
| Firmicutes | Negativicutes | Selenomonadales | Sporomusaceae | *Sporomusa* | 0.001 | 0.000 | 0.541 | 0.014 | 0.000 | 0.005 |
| Firmicutes | Negativicutes | Veillonellales | Veillonellaceae | *Megasphaera* | 0.000 | 0.000 | 0.013 | 0.000 | 0.000 | 0.000 |
| Firmicutes | Negativicutes | Veillonellales | Veillonellaceae | *Negativicoccus* | 0.001 | 0.000 | 0.000 | 0.000 | 0.000 | 0.000 |
| Firmicutes | RF3 | Unclassified | Unclassified | Unclassified | 0.000 | 0.000 | 0.000 | 0.004 | 0.000 | 0.001 |
| Firmicutes | RF3 | Unclassified | Unclassified | Unclassified | 0.001 | 0.000 | 0.000 | 0.000 | 0.000 | 0.000 |
| Firmicutes | Tissierellia | Tissierellales | Gottschalkiaceae | *Andreesenia* | 0.001 | 0.000 | 0.000 | 0.000 | 0.000 | 0.000 |
| Firmicutes | Tissierellia | Tissierellales | Gottschalkiaceae | *Gottschalkia* | 0.000 | 0.000 | 0.000 | 0.000 | 0.000 | 0.001 |
| Firmicutes | Tissierellia | Tissierellales | Peptoniphilaceae | *Anaerococcus* | 0.000 | 0.000 | 0.000 | 0.007 | 0.000 | 0.000 |
| Firmicutes | Tissierellia | Tissierellales | Peptoniphilaceae | *Anaerosphaera* | 0.001 | 0.000 | 0.000 | 0.000 | 0.000 | 0.000 |
| Firmicutes | Tissierellia | Tissierellales | Peptoniphilaceae | *Helcococcus* | 0.000 | 0.000 | 0.000 | 0.000 | 0.000 | 0.001 |
| Firmicutes | Tissierellia | Tissierellales | Peptoniphilaceae | *Peptoniphilus* | 0.004 | 0.000 | 0.000 | 0.007 | 0.000 | 0.000 |
| Firmicutes | Tissierellia | Unclassified | Unclassified | *Sedimentibacter* | 0.004 | 0.000 | 0.000 | 0.000 | 0.000 | 0.002 |
| Fusobacteria | Fusobacteria | Fusobacteriales | boneC3G7 | Unclassified | 0.000 | 0.000 | 0.000 | 0.000 | 0.000 | 0.002 |
| Fusobacteria | Fusobacteria | Fusobacteriales | Fish gut cluster | Unclassified | 0.000 | 0.000 | 0.000 | 0.000 | 0.000 | 0.001 |
| Fusobacteria | Fusobacteria | Fusobacteriales | Fusobacteriaceae | *Cetobacterium* | 0.000 | 0.000 | 0.000 | 0.000 | 0.000 | 0.003 |
| Fusobacteria | Fusobacteria | Fusobacteriales | Leptotrichiaceae | *Leptotrichia* | 0.000 | 0.000 | 0.000 | 0.004 | 0.000 | 0.006 |
| Fusobacteria | Fusobacteria | Fusobacteriales | Leptotrichiaceae | Uncultured 1 | 0.000 | 0.000 | 0.000 | 0.000 | 0.000 | 0.003 |
| Fusobacteria | Fusobacteria | Fusobacteriales | Leptotrichiaceae | Uncultured 5 | 0.000 | 0.000 | 0.000 | 0.000 | 0.000 | 0.001 |
| Fusobacteria | Fusobacteria | KD3-66 | Unclassified | Unclassified | 0.000 | 0.000 | 0.000 | 0.004 | 0.000 | 0.000 |
| Fusobacteria | Fusobacteriia | Fusobacteriales | Fusobacteriaceae | *Fusobacterium* | 0.000 | 0.000 | 0.000 | 0.000 | 0.004 | 0.000 |

**Table S2 continued.**

| | | | | | | | | | | |
|---|---|---|---|---|---|---|---|---|---|---|
| Fusobacteria | Fusobacteriia | Fusobacteriales | Fusobacteriaceae | *Hypnocyclicus* | 0.000 | 0.000 | 0.000 | 0.000 | 0.000 | 0.001 |
| Fusobacteria | Fusobacteriia | Fusobacteriales | Leptotrichiaceae | *Oceanivirga* | 0.000 | 0.000 | 0.000 | 0.000 | 0.000 | 0.001 |
| Fusobacteria | Fusobacteriia | Fusobacteriales | Leptotrichiaceae | *Sebaldella* | 0.000 | 0.032 | 0.000 | 0.007 | 1.391 | 6.780 |
| Gemmatimonadetes | Gemmatimonadetes | AT425-EubC11 terrestrial group | Unclassified | Unclassified | 0.000 | 0.000 | 0.000 | 0.011 | 0.000 | 0.000 |
| Gemmatimonadetes | Gemmatimonadetes | AT425-EubC11 terrestrial group | Unclassified | Unclassified | 0.001 | 0.000 | 0.000 | 0.000 | 0.000 | 0.000 |
| Gemmatimonadetes | Gemmatimonadetes | BD2-11 terrestrial group | Unclassified | Unclassified | 0.000 | 0.000 | 0.000 | 0.025 | 0.000 | 0.000 |
| Gemmatimonadetes | Gemmatimonadetes | Gemmatimonadales | Gemmatimonadaceae | *Gemmatimonas* | 0.000 | 0.000 | 0.000 | 0.018 | 0.000 | 0.000 |
| Gemmatimonadetes | Gemmatimonadetes | Gemmatimonadales | Gemmatimonadaceae | Uncultured 1 | 0.000 | 0.000 | 0.000 | 0.082 | 0.000 | 0.000 |
| Gemmatimonadetes | Gemmatimonadetes | Gemmatimonadales | Gemmatimonadaceae | Uncultured 10 | 0.000 | 0.000 | 0.000 | 0.014 | 0.000 | 0.000 |
| Gemmatimonadetes | Gemmatimonadetes | Gemmatimonadales | Gemmatimonadaceae | Uncultured 2 | 0.000 | 0.000 | 0.000 | 0.004 | 0.000 | 0.000 |
| Gemmatimonadetes | Gemmatimonadetes | Gemmatimonadales | Gemmatimonadaceae | Uncultured 4 | 0.000 | 0.000 | 0.000 | 0.007 | 0.000 | 0.000 |
| Gemmatimonadetes | Gemmatimonadetes | Gemmatimonadales | Gemmatimonadaceae | Uncultured 5 | 0.000 | 0.000 | 0.000 | 0.007 | 0.000 | 0.000 |
| Gemmatimonadetes | Gemmatimonadetes | Gemmatimonadales | Gemmatimonadaceae | Uncultured 7 | 0.001 | 0.000 | 0.000 | 0.053 | 0.000 | 0.001 |
| Gemmatimonadetes | Gemmatimonadetes | Gemmatimonadales | Gemmatimonadaceae | Uncultured 8 | 0.000 | 0.000 | 0.000 | 0.039 | 0.000 | 0.000 |
| Gemmatimonadetes | Gemmatimonadetes | Gemmatimonadales | Gemmatimonadaceae | Uncultured 9 | 0.000 | 0.000 | 0.000 | 0.007 | 0.000 | 0.000 |
| Gemmatimonadetes | Gemmatimonadetes | Kazan-1B-37 | Unclassified | Unclassified | 0.001 | 0.000 | 0.000 | 0.000 | 0.000 | 0.000 |
| Gemmatimonadetes | Gemmatimonadetes | PAUC43f marine benthic group | Unclassified | Unclassified | 0.000 | 0.000 | 0.000 | 0.000 | 0.000 | 0.001 |
| Gemmatimonadetes | Gemmatimonadetes | S0134 terrestrial group | Unclassified | Unclassified | 0.000 | 0.000 | 0.000 | 0.089 | 0.000 | 0.000 |
| Gemmatimonadetes | Gemmatimonadetes | S0134 terrestrial group | Unclassified | Unclassified | 0.001 | 0.000 | 0.000 | 0.000 | 0.000 | 0.000 |
| Gemmatimonadetes | Longimicrobia | Longimicrobiales | Longimicrobiaceae | *Longimicrobium* | 0.000 | 0.000 | 0.000 | 0.004 | 0.000 | 0.000 |
| Gemmatimonadetes | Unclassified | Unclassified | Unclassified | Unclassified | 0.000 | 0.000 | 0.000 | 0.029 | 0.000 | 0.000 |
| Hyd24-12 | Unclassified | Unclassified | Unclassified | Unclassified | 0.001 | 0.000 | 0.000 | 0.000 | 0.000 | 0.000 |
| JL-ETNP-Z39 | Unclassified | Unclassified | Unclassified | Unclassified | 0.000 | 0.000 | 0.000 | 0.007 | 0.000 | 0.000 |
| Lentisphaerae | Lentisphaeria | BS5 | Unclassified | Unclassified | 0.000 | 0.191 | 0.000 | 0.000 | 0.000 | 0.000 |
| Lentisphaerae | Lentisphaeria | LD1-PA34 | Unclassified | Unclassified | 0.000 | 0.032 | 0.000 | 0.000 | 0.000 | 0.000 |
| Lentisphaerae | Lentisphaeria | RFP12 gut group | Unclassified | Unclassified | 0.000 | 0.000 | 0.000 | 0.000 | 0.004 | 0.003 |

**Table S2 continued.**

| | | | | | | | | | | |
|---|---|---|---|---|---|---|---|---|---|---|
| Lentisphaerae | Lentisphaeria | RFP12 gut group | Unclassified | Unclassified | 0.006 | 0.000 | 0.000 | 0.000 | 0.000 | 0.000 |
| Lentisphaerae | Lentisphaeria | Victivallales | Victivallaceae | Uncultured | 0.001 | 0.000 | 0.000 | 0.000 | 0.000 | 0.000 |
| Lentisphaerae | Lentisphaeria | WCHB1-41 | Unclassified | Unclassified | 0.000 | 0.000 | 0.000 | 0.000 | 0.000 | 0.001 |
| Nitrospirae | Nitrospira | Nitrospirales | Cluster 319-6A21 | Unclassified | 0.000 | 0.000 | 0.000 | 0.029 | 0.000 | 0.001 |
| Nitrospirae | Nitrospira | Nitrospirales | Cluster 4-29 | Unclassified | 0.000 | 0.000 | 0.013 | 0.000 | 0.004 | 0.000 |
| Nitrospirae | Nitrospira | Nitrospirales | Nitrospiraceae | *Candidatus* Magnetobacterium | 0.001 | 0.000 | 0.000 | 0.000 | 0.000 | 0.000 |
| Nitrospirae | Nitrospira | Nitrospirales | Nitrospiraceae | *Nitrospira* | 0.000 | 0.000 | 0.013 | 0.110 | 0.000 | 0.003 |
| Nitrospirae | Nitrospira | Nitrospirales | Nitrospiraceae | *Thermodesulfovibrio* | 0.001 | 0.000 | 0.000 | 0.000 | 0.000 | 0.000 |
| Nitrospirae | Nitrospira | Nitrospirales | Nitrospiraceae | Uncultured 1 | 0.005 | 0.000 | 0.000 | 0.000 | 0.020 | 0.000 |
| Nitrospirae | Nitrospira | Nitrospirales | Nitrospiraceae | Uncultured 1 | 0.000 | 0.000 | 0.000 | 0.000 | 0.000 | 0.002 |
| Nitrospirae | Nitrospira | Nitrospirales | Nitrospiraceae | Uncultured 3 | 0.001 | 0.000 | 0.000 | 0.000 | 0.000 | 0.000 |
| NPL-UPA2 | Unclassified | Unclassified | Unclassified | Unclassified | 0.001 | 0.000 | 0.000 | 0.000 | 0.000 | 0.000 |
| Planctomycetes | H05-P-BN-P5 | Unclassified | Unclassified | Unclassified | 0.000 | 0.000 | 0.000 | 0.004 | 0.000 | 0.000 |
| Planctomycetes | OM190 | Unclassified | Unclassified | Unclassified | 0.000 | 0.000 | 0.000 | 0.021 | 0.000 | 0.000 |
| Planctomycetes | Phycisphaerae | CPla-3 termite group | Unclassified | Unclassified | 0.000 | 0.000 | 0.000 | 0.004 | 0.000 | 0.000 |
| Planctomycetes | Phycisphaerae | Phycisphaerales | Phycisphaeraceae | AKYG587 | 0.000 | 0.000 | 0.000 | 0.007 | 0.000 | 0.000 |
| Planctomycetes | Phycisphaerae | Phycisphaerales | Phycisphaeraceae | SM1A02 | 0.000 | 0.000 | 0.000 | 0.004 | 0.000 | 0.000 |
| Planctomycetes | Phycisphaerae | S-70 | Unclassified | Unclassified | 0.009 | 0.000 | 0.000 | 0.000 | 0.000 | 0.000 |
| Planctomycetes | Phycisphaerae | WD2101 soil group | Unclassified | Unclassified | 0.000 | 0.000 | 0.000 | 0.007 | 0.000 | 0.000 |
| Planctomycetes | Pla4 lineage | Unclassified | Unclassified | Unclassified | 0.000 | 0.000 | 0.000 | 0.007 | 0.000 | 0.000 |
| Planctomycetes | Planctomycetacia | Planctomycetales | Planctomycetaceae | Gemmata | 0.000 | 0.000 | 0.000 | 0.007 | 0.000 | 0.000 |
| Planctomycetes | Planctomycetacia | Planctomycetales | Planctomycetaceae | Gut cluster 2 | 0.017 | 0.032 | 0.000 | 0.011 | 0.012 | 0.002 |
| Planctomycetes | Planctomycetacia | Planctomycetales | Planctomycetaceae | *Isosphaera* | 0.000 | 0.000 | 0.000 | 0.007 | 0.000 | 0.000 |
| Planctomycetes | Planctomycetacia | Planctomycetales | Planctomycetaceae | Pir4 lineage | 0.000 | 0.000 | 0.000 | 0.007 | 0.000 | 0.000 |
| Planctomycetes | Planctomycetacia | Planctomycetales | Planctomycetaceae | *Pirellula* | 0.000 | 0.000 | 0.000 | 0.007 | 0.000 | 0.000 |
| Planctomycetes | Planctomycetacia | Planctomycetales | Planctomycetaceae | *Rhodopirellula* | 0.000 | 0.000 | 0.000 | 0.004 | 0.000 | 0.000 |
| Planctomycetes | Planctomycetacia | Planctomycetales | Planctomycetaceae | *Singulisphaera* | 0.000 | 0.000 | 0.000 | 0.004 | 0.000 | 0.000 |

**Table S2 continued.**

| | | | | | | | | | | |
|---|---|---|---|---|---|---|---|---|---|---|
| Planctomycetes | Planctomycetacia | Planctomycetales | Planctomycetaceae | Termite cockroach cluster 2 | 0.177 | 0.128 | 0.000 | 0.064 | 0.020 | 0.010 |
| Planctomycetes | Planctomycetacia | Planctomycetales | Planctomycetaceae | Unclassified | 0.000 | 0.032 | 0.000 | 0.000 | 0.000 | 0.000 |
| Planctomycetes | Planctomycetacia | Planctomycetales | Planctomycetaceae | Uncultured 11 | 0.000 | 0.000 | 0.000 | 0.004 | 0.000 | 0.000 |
| Planctomycetes | Planctomycetacia | Planctomycetales | Planctomycetaceae | Uncultured 23 | 0.000 | 0.000 | 0.000 | 0.004 | 0.000 | 0.000 |
| Planctomycetes | Planctomycetacia | Planctomycetales | Planctomycetaceae | Uncultured 7 | 0.000 | 0.032 | 0.000 | 0.000 | 0.000 | 0.000 |
| Planctomycetes | vadinHA49 | Insect cluster | Unclassified | Unclassified | 0.000 | 1.148 | 0.000 | 0.787 | 0.403 | 0.278 |
| Planctomycetes | vadinHA49 | Insect cluster | Unclassified | Unclassified | 1.214 | 0.000 | 0.000 | 0.000 | 0.000 | 0.000 |
| Planctomycetes | vadinHA49 | Unclassified | Unclassified | Unclassified | 0.000 | 0.096 | 0.013 | 0.021 | 0.041 | 0.014 |
| Planctomycetes | vadinHA49 | Unclassified | Unclassified | Unclassified | 0.059 | 0.000 | 0.000 | 0.000 | 0.000 | 0.000 |
| Proteobacteria | Alphaproteobacteria | Caulobacterales | Caulobacteraceae | *Asticcacaulis* | 0.001 | 0.000 | 0.000 | 0.011 | 0.000 | 0.000 |
| Proteobacteria | Alphaproteobacteria | Caulobacterales | Caulobacteraceae | *Brevundimonas* | 0.002 | 0.000 | 0.000 | 0.143 | 0.000 | 0.002 |
| Proteobacteria | Alphaproteobacteria | Caulobacterales | Caulobacteraceae | *Caulobacter* | 0.001 | 0.000 | 0.000 | 0.110 | 0.000 | 0.000 |
| Proteobacteria | Alphaproteobacteria | Caulobacterales | Caulobacteraceae | *Phenylobacterium* | 0.000 | 0.000 | 0.000 | 0.078 | 0.000 | 0.000 |
| Proteobacteria | Alphaproteobacteria | Caulobacterales | Caulobacteraceae | Unclassified | 0.000 | 0.000 | 0.000 | 0.025 | 0.000 | 0.000 |
| Proteobacteria | Alphaproteobacteria | Caulobacterales | Caulobacteraceae | Uncultured 3 | 0.002 | 0.000 | 0.013 | 0.110 | 0.000 | 0.000 |
| Proteobacteria | Alphaproteobacteria | Caulobacterales | Caulobacteraceae | Uncultured 6 | 0.000 | 0.000 | 0.000 | 0.000 | 0.000 | 0.001 |
| Proteobacteria | Alphaproteobacteria | Caulobacterales | Hyphomonadaceae | Uncultured 1 | 0.001 | 0.000 | 0.000 | 0.000 | 0.000 | 0.000 |
| Proteobacteria | Alphaproteobacteria | Caulobacterales | Hyphomonadaceae | Uncultured 2 | 0.000 | 0.000 | 0.000 | 0.021 | 0.000 | 0.000 |
| Proteobacteria | Alphaproteobacteria | DB1-14 | Unclassified | Unclassified | 0.000 | 0.000 | 0.000 | 0.025 | 0.000 | 0.000 |
| Proteobacteria | Alphaproteobacteria | DB1-14 | Unclassified | Unclassified | 0.010 | 0.000 | 0.000 | 0.000 | 0.000 | 0.000 |
| Proteobacteria | Alphaproteobacteria | Kiloneillales | Kiloniellaceae | *Kiloniella* | 0.014 | 0.032 | 0.000 | 0.014 | 0.008 | 0.000 |
| Proteobacteria | Alphaproteobacteria | Kordiimonadales | Kordiimonadaceae | Kordiimonas | 0.000 | 0.000 | 0.000 | 0.004 | 0.000 | 0.000 |
| Proteobacteria | Alphaproteobacteria | MNG3 | Unclassified | Unclassified | 0.000 | 0.000 | 0.000 | 0.004 | 0.000 | 0.000 |
| Proteobacteria | Alphaproteobacteria | MNG3 | Unclassified | Unclassified | 0.001 | 0.000 | 0.000 | 0.000 | 0.000 | 0.000 |
| Proteobacteria | Alphaproteobacteria | Parvularculales | Parvularculaceae | *Parvularcula* | 0.033 | 0.032 | 0.000 | 0.018 | 0.024 | 0.014 |
| Proteobacteria | Alphaproteobacteria | Rhizobiales | Aurantimonadaceae | *Aurantimonas* | 0.000 | 0.000 | 0.000 | 0.004 | 0.000 | 0.000 |
| Proteobacteria | Alphaproteobacteria | Rhizobiales | Aurantimonadaceae | *Aureimonas* | 0.000 | 0.000 | 0.000 | 0.039 | 0.000 | 0.000 |

**Table S2 continued.**

| | | | | | | | | | | |
|---|---|---|---|---|---|---|---|---|---|---|
| Proteobacteria | Alphaproteobacteria | Rhizobiales | Aurantimonadaceae | *Fulvimarina* | 0.000 | 0.000 | 0.000 | 0.004 | 0.000 | 0.000 |
| Proteobacteria | Alphaproteobacteria | Rhizobiales | Bartonellaceae | *Bartonella* | 0.007 | 0.032 | 0.000 | 0.032 | 0.004 | 0.000 |
| Proteobacteria | Alphaproteobacteria | Rhizobiales | Beijerinckiaceae | *Beijerinckia* | 0.002 | 0.032 | 0.000 | 0.004 | 0.000 | 0.001 |
| Proteobacteria | Alphaproteobacteria | Rhizobiales | Beijerinkiaceae | *Methylovirgula* | 0.000 | 0.000 | 0.000 | 0.004 | 0.000 | 0.000 |
| Proteobacteria | Alphaproteobacteria | Rhizobiales | Beijernckiaceae | *Methylocapsa* | 0.001 | 0.000 | 0.000 | 0.000 | 0.000 | 0.000 |
| Proteobacteria | Alphaproteobacteria | Rhizobiales | Beijernckiaceae | *Methylocella* | 0.001 | 0.000 | 0.000 | 0.004 | 0.000 | 0.000 |
| Proteobacteria | Alphaproteobacteria | Rhizobiales | Bradyrhizobiaceae | *Afipia* | 0.006 | 0.000 | 0.000 | 0.232 | 0.000 | 0.000 |
| Proteobacteria | Alphaproteobacteria | Rhizobiales | Bradyrhizobiaceae | *Bradyrhizobium* | 0.006 | 0.000 | 0.039 | 0.428 | 0.004 | 0.000 |
| Proteobacteria | Alphaproteobacteria | Rhizobiales | Bradyrhizobiaceae | *Nitrobacter* | 0.001 | 0.032 | 0.013 | 0.053 | 0.004 | 0.000 |
| Proteobacteria | Alphaproteobacteria | Rhizobiales | Bradyrhizobiaceae | *Rhodopseudomonas* | 0.004 | 0.000 | 0.000 | 0.014 | 0.000 | 0.000 |
| Proteobacteria | Alphaproteobacteria | Rhizobiales | Bradyrhizobiaceae | *Tardiphaga* | 0.001 | 0.000 | 0.000 | 0.011 | 0.000 | 0.000 |
| Proteobacteria | Alphaproteobacteria | Rhizobiales | Brucellaceae | *Brucella* | 0.000 | 0.000 | 0.000 | 0.004 | 0.000 | 0.000 |
| Proteobacteria | Alphaproteobacteria | Rhizobiales | Brucellaceae | *Mycoplana* | 0.000 | 0.000 | 0.000 | 0.011 | 0.000 | 0.000 |
| Proteobacteria | Alphaproteobacteria | Rhizobiales | Brucellaceae | *Ochrobactrum* | 0.027 | 0.000 | 0.000 | 0.036 | 0.000 | 0.001 |
| Proteobacteria | Alphaproteobacteria | Rhizobiales | Brucellaceae | *Paenochrobactrum* | 0.000 | 0.000 | 0.000 | 0.000 | 0.000 | 0.001 |
| Proteobacteria | Alphaproteobacteria | Rhizobiales | Brucellaceae | *Pseudochrobactrum* | 0.001 | 0.000 | 0.000 | 0.014 | 0.000 | 0.000 |
| Proteobacteria | Alphaproteobacteria | Rhizobiales | Chelatococcaceae | *Chelatococcus* | 0.000 | 0.000 | 0.000 | 0.004 | 0.000 | 0.001 |
| Proteobacteria | Alphaproteobacteria | Rhizobiales | Cohaesibacteraceae | *Breoghania* | 0.001 | 0.000 | 0.000 | 0.000 | 0.000 | 0.000 |
| Proteobacteria | Alphaproteobacteria | Rhizobiales | Cohaesibacteraceae | *Cohaesibacter* | 0.009 | 0.000 | 0.013 | 0.014 | 0.000 | 0.000 |
| Proteobacteria | Alphaproteobacteria | Rhizobiales | Hyphomicrobiaceae | *Arsenicitalea* | 0.000 | 0.000 | 0.000 | 0.004 | 0.000 | 0.000 |
| Proteobacteria | Alphaproteobacteria | Rhizobiales | Hyphomicrobiaceae | *Blastochloris* | 0.001 | 0.000 | 0.000 | 0.000 | 0.000 | 0.000 |
| Proteobacteria | Alphaproteobacteria | Rhizobiales | Hyphomicrobiaceae | *Devosia* | 0.007 | 0.032 | 0.000 | 0.121 | 0.000 | 0.001 |
| Proteobacteria | Alphaproteobacteria | Rhizobiales | Hyphomicrobiaceae | *Dichotomicrobium* | 0.000 | 0.000 | 0.000 | 0.004 | 0.000 | 0.000 |
| Proteobacteria | Alphaproteobacteria | Rhizobiales | Hyphomicrobiaceae | *Hyphomicrobium* | 0.001 | 0.032 | 0.000 | 0.014 | 0.000 | 0.001 |
| Proteobacteria | Alphaproteobacteria | Rhizobiales | Hyphomicrobiaceae | *Methylorhabdus* | 0.000 | 0.000 | 0.000 | 0.004 | 0.000 | 0.000 |
| Proteobacteria | Alphaproteobacteria | Rhizobiales | Hyphomicrobiaceae | *Methyloterrigena* | 0.000 | 0.000 | 0.000 | 0.004 | 0.000 | 0.000 |
| Proteobacteria | Alphaproteobacteria | Rhizobiales | Hyphomicrobiaceae | *Pedomicrobium* | 0.001 | 0.000 | 0.000 | 0.029 | 0.000 | 0.002 |

**Table S2 continued.**

| | | | | | | | | | | |
|---|---|---|---|---|---|---|---|---|---|---|
| Proteobacteria | Alphaproteobacteria | Rhizobiales | Hyphomicrobiaceae | *Rhodoplanes* | 0.000 | 0.000 | 0.000 | 0.011 | 0.000 | 0.000 |
| Proteobacteria | Alphaproteobacteria | Rhizobiales | Hypomicrobiaceae | *Prosthecomicrobium* | 0.000 | 0.000 | 0.000 | 0.011 | 0.000 | 0.000 |
| Proteobacteria | Alphaproteobacteria | Rhizobiales | Methylobacteriaceae | *Methanobacterium* | 0.002 | 0.000 | 0.000 | 0.000 | 0.000 | 0.000 |
| Proteobacteria | Alphaproteobacteria | Rhizobiales | Methylobacteriaceae | *Methylobacterium* | 0.005 | 0.000 | 0.013 | 0.374 | 0.012 | 0.005 |
| Proteobacteria | Alphaproteobacteria | Rhizobiales | Methylobacteriaceae | *Microvirga* | 0.000 | 0.000 | 0.000 | 0.096 | 0.000 | 0.001 |
| Proteobacteria | Alphaproteobacteria | Rhizobiales | Methylocystaceae | *Chenggangzhangella* | 0.001 | 0.000 | 0.000 | 0.004 | 0.000 | 0.001 |
| Proteobacteria | Alphaproteobacteria | Rhizobiales | Methylocystaceae | *Hansschlegelia* | 0.001 | 0.000 | 0.013 | 0.011 | 0.004 | 0.000 |
| Proteobacteria | Alphaproteobacteria | Rhizobiales | Methylocystaceae | *Pleomorphomonas* | 0.002 | 0.000 | 0.000 | 0.025 | 0.000 | 0.001 |
| Proteobacteria | Alphaproteobacteria | Rhizobiales | Methylocystaceae | Uncultured 1 | 0.010 | 0.000 | 0.013 | 0.039 | 0.016 | 0.002 |
| Proteobacteria | Alphaproteobacteria | Rhizobiales | Methylocystaceae | Uncultured 2 | 0.000 | 0.000 | 0.000 | 0.011 | 0.000 | 0.000 |
| Proteobacteria | Alphaproteobacteria | Rhizobiales | Phyllobacteriaceae | *Aliihoeflea* | 0.000 | 0.000 | 0.000 | 0.011 | 0.000 | 0.000 |
| Proteobacteria | Alphaproteobacteria | Rhizobiales | Phyllobacteriaceae | *Corticibacterium* | 0.000 | 0.000 | 0.000 | 0.007 | 0.000 | 0.000 |
| Proteobacteria | Alphaproteobacteria | Rhizobiales | Phyllobacteriaceae | *Hoeflea* | 0.001 | 0.000 | 0.000 | 0.004 | 0.000 | 0.000 |
| Proteobacteria | Alphaproteobacteria | Rhizobiales | Phyllobacteriaceae | *Mesorhizobium* | 0.001 | 0.000 | 0.000 | 0.025 | 0.000 | 0.000 |
| Proteobacteria | Alphaproteobacteria | Rhizobiales | Phyllobacteriaceae | *Nitratireductor* | 0.002 | 0.000 | 0.000 | 0.004 | 0.000 | 0.000 |
| Proteobacteria | Alphaproteobacteria | Rhizobiales | Phyllobacteriaceae | *Phyllobacterium* | 0.004 | 0.032 | 0.000 | 0.004 | 0.000 | 0.000 |
| Proteobacteria | Alphaproteobacteria | Rhizobiales | Phyllobacteriaceae | *Pseudaminobacter* | 0.000 | 0.000 | 0.000 | 0.004 | 0.000 | 0.000 |
| Proteobacteria | Alphaproteobacteria | Rhizobiales | Phyllobacteriaceae | *Tianweitania* | 0.000 | 0.000 | 0.000 | 0.007 | 0.000 | 0.000 |
| Proteobacteria | Alphaproteobacteria | Rhizobiales | Rhizobiaceae | *Agrobacterium* | 0.000 | 0.000 | 0.000 | 0.025 | 0.000 | 0.000 |
| Proteobacteria | Alphaproteobacteria | Rhizobiales | Rhizobiaceae | *Ciceribacter* | 0.004 | 0.000 | 0.000 | 0.011 | 0.000 | 0.000 |
| Proteobacteria | Alphaproteobacteria | Rhizobiales | Rhizobiaceae | *Ensifer* | 0.000 | 0.000 | 0.000 | 0.014 | 0.000 | 0.000 |
| Proteobacteria | Alphaproteobacteria | Rhizobiales | Rhizobiaceae | *Kaistia* | 0.000 | 0.000 | 0.000 | 0.018 | 0.000 | 0.000 |
| Proteobacteria | Alphaproteobacteria | Rhizobiales | Rhizobiaceae | *Rhizobium* | 0.033 | 0.096 | 0.000 | 0.267 | 0.000 | 0.007 |
| Proteobacteria | Alphaproteobacteria | Rhizobiales | Rhizobiaceae | *Rhodobium* | 0.002 | 0.000 | 0.000 | 0.004 | 0.000 | 0.000 |
| Proteobacteria | Alphaproteobacteria | Rhizobiales | Rhizobiaceae | *Shinella* | 0.002 | 0.000 | 0.000 | 0.046 | 0.000 | 0.000 |
| Proteobacteria | Alphaproteobacteria | Rhizobiales | Rhizobiaceae | *Sinorhizobium* | 0.001 | 0.000 | 0.000 | 0.007 | 0.000 | 0.000 |
| Proteobacteria | Alphaproteobacteria | Rhizobiales | Rhodobiaceae | *Amorphus* | 0.000 | 0.000 | 0.000 | 0.004 | 0.000 | 0.000 |

**Table S2 continued.**

| | | | | | | | | | | |
|---|---|---|---|---|---|---|---|---|---|---|
| Proteobacteria | Alphaproteobacteria | Rhizobiales | Rhodobiaceae | *Parvibaculum* | 0.001 | 0.000 | 0.000 | 0.000 | 0.000 | 0.000 |
| Proteobacteria | Alphaproteobacteria | Rhizobiales | Rhodobiaceae | *Rhodobium* 2 | 0.001 | 0.000 | 0.000 | 0.021 | 0.008 | 0.001 |
| Proteobacteria | Alphaproteobacteria | Rhizobiales | Rhodobiaceae | *Rhodoligotrophos* | 0.000 | 0.000 | 0.000 | 0.004 | 0.000 | 0.000 |
| Proteobacteria | Alphaproteobacteria | Rhizobiales | Rhodobiaceae | *Roseospirillum* | 0.002 | 0.000 | 0.000 | 0.000 | 0.000 | 0.000 |
| Proteobacteria | Alphaproteobacteria | Rhizobiales | Rhodobiaceae | *Tepidicaulis* | 0.000 | 0.000 | 0.000 | 0.004 | 0.000 | 0.000 |
| Proteobacteria | Alphaproteobacteria | Rhizobiales | Unclassified | *Alsobacter* | 0.000 | 0.000 | 0.000 | 0.004 | 0.000 | 0.000 |
| Proteobacteria | Alphaproteobacteria | Rhizobiales | Unclassified | *Bauldia* | 0.000 | 0.000 | 0.000 | 0.004 | 0.000 | 0.000 |
| Proteobacteria | Alphaproteobacteria | Rhizobiales | Unclassified | *Nordella* | 0.000 | 0.000 | 0.000 | 0.007 | 0.000 | 0.000 |
| Proteobacteria | Alphaproteobacteria | Rhizobiales | Unclassified | *Pseudorhodoplanes* | 0.000 | 0.000 | 0.000 | 0.004 | 0.000 | 0.000 |
| Proteobacteria | Alphaproteobacteria | Rhizobiales | Xanthobacteraceae | *Ancylobacter* | 0.000 | 0.000 | 0.000 | 0.011 | 0.000 | 0.000 |
| Proteobacteria | Alphaproteobacteria | Rhizobiales | Xanthobacteraceae | *Labrys* | 0.004 | 0.000 | 0.000 | 0.000 | 0.000 | 0.000 |
| Proteobacteria | Alphaproteobacteria | Rhizobiales | Xanthobacteraceae | *Pseudoxanthobacter* | 0.000 | 0.032 | 0.000 | 0.011 | 0.000 | 0.000 |
| Proteobacteria | Alphaproteobacteria | Rhizobiales | Xanthobacteraceae | *Xanthobacter* | 0.000 | 0.000 | 0.000 | 0.004 | 0.000 | 0.000 |
| Proteobacteria | Alphaproteobacteria | Rhizobiales 1 | Aurantimonadaceae | *Aurantimonas* 1 | 0.000 | 0.000 | 0.000 | 0.004 | 0.000 | 0.000 |
| Proteobacteria | Alphaproteobacteria | Rhizobiales 1 | Aurantimonadaceae | *Aurantimonas* 3 | 0.000 | 0.000 | 0.000 | 0.004 | 0.000 | 0.000 |
| Proteobacteria | Alphaproteobacteria | Rhizobiales 1 | Aurantimonadaceae | Unclassified | 0.000 | 0.000 | 0.000 | 0.011 | 0.000 | 0.000 |
| Proteobacteria | Alphaproteobacteria | Rhizobiales 1 | Aurantimonadaceae | Uncultured | 0.000 | 0.000 | 0.000 | 0.004 | 0.000 | 0.000 |
| Proteobacteria | Alphaproteobacteria | Rhizobiales 1 | Brucellaceae | *Ochrobactrum* 1 | 0.000 | 0.000 | 0.000 | 0.004 | 0.000 | 0.000 |
| Proteobacteria | Alphaproteobacteria | Rhizobiales 1 | Brucellaceae | *Ochrobactrum* 2 | 0.012 | 0.000 | 0.000 | 0.000 | 0.000 | 0.000 |
| Proteobacteria | Alphaproteobacteria | Rhizobiales 1 | Brucellaceae | *Ochrobactrum* 3 | 0.000 | 0.000 | 0.000 | 0.011 | 0.000 | 0.000 |
| Proteobacteria | Alphaproteobacteria | Rhizobiales 1 | Brucellaceae | *Ochrobactrum* 4 | 0.006 | 0.032 | 0.000 | 0.007 | 0.000 | 0.000 |
| Proteobacteria | Alphaproteobacteria | Rhizobiales 1 | Brucellaceae | Unclassified | 0.001 | 0.000 | 0.000 | 0.000 | 0.000 | 0.000 |
| Proteobacteria | Alphaproteobacteria | Rhizobiales 1 | Hyphomicrobiaceae | *Devosia-Prosthecomicrobium* | 0.010 | 0.032 | 0.013 | 0.082 | 0.000 | 0.000 |
| Proteobacteria | Alphaproteobacteria | Rhizobiales 1 | Hyphomicrobiaceae | Unclassified | 0.000 | 0.000 | 0.000 | 0.007 | 0.000 | 0.000 |
| Proteobacteria | Alphaproteobacteria | Rhizobiales 1 | Hyphomicrobiaceae | Uncultured 2 | 0.001 | 0.000 | 0.000 | 0.014 | 0.000 | 0.000 |
| Proteobacteria | Alphaproteobacteria | Rhizobiales 1 | Hyphomicrobiaceae | Unclassified | 0.001 | 0.000 | 0.000 | 0.000 | 0.000 | 0.000 |
| Proteobacteria | Alphaproteobacteria | Rhizobiales 1 | Phyllobacteriaceae | *Ahrensia* | 0.005 | 0.000 | 0.000 | 0.004 | 0.000 | 0.001 |

**Table S2 continued.**

| | | | | | | | | | | |
|---|---|---|---|---|---|---|---|---|---|---|
| Proteobacteria | Alphaproteobacteria | Rhizobiales 1 | Phyllobacteriaceae | *Aminobacter* 1 | 0.004 | 0.032 | 0.000 | 0.011 | 0.000 | 0.000 |
| Proteobacteria | Alphaproteobacteria | Rhizobiales 1 | Phyllobacteriaceae | *Mesorhizobium* 11 | 0.000 | 0.000 | 0.000 | 0.004 | 0.000 | 0.000 |
| Proteobacteria | Alphaproteobacteria | Rhizobiales 1 | Phyllobacteriaceae | *Mesorhizobium* 12 | 0.000 | 0.000 | 0.000 | 0.011 | 0.000 | 0.000 |
| Proteobacteria | Alphaproteobacteria | Rhizobiales 1 | Phyllobacteriaceae | *Pseudoaminobacter* | 0.000 | 0.032 | 0.000 | 0.000 | 0.000 | 0.000 |
| Proteobacteria | Alphaproteobacteria | Rhizobiales 1 | Phyllobacteriaceae | Unclassified | 0.000 | 0.000 | 0.000 | 0.021 | 0.000 | 0.001 |
| Proteobacteria | Alphaproteobacteria | Rhizobiales 1 | Phyllobacteriaceae | Uncultured 3 | 0.000 | 0.000 | 0.000 | 0.004 | 0.000 | 0.000 |
| Proteobacteria | Alphaproteobacteria | Rhizobiales 1 | Phyllobacteriaceae | Uncultured 4 | 0.000 | 0.000 | 0.000 | 0.007 | 0.000 | 0.000 |
| Proteobacteria | Alphaproteobacteria | Rhizobiales 1 | Phyllobacteriaceae | Uncultured 5 | 0.004 | 0.000 | 0.000 | 0.000 | 0.000 | 0.001 |
| Proteobacteria | Alphaproteobacteria | Rhizobiales 1 | Rhizobiaceae | *Rhizobium-Agrobacterium* | 0.028 | 0.096 | 0.039 | 0.264 | 0.004 | 0.003 |
| Proteobacteria | Alphaproteobacteria | Rhizobiales 1 | Rhizobiaceae | *Sinorhizobium-Ensifer* | 0.000 | 0.000 | 0.000 | 0.025 | 0.004 | 0.001 |
| Proteobacteria | Alphaproteobacteria | Rhizobiales 1 | Shinella genera incertae sedis | Shinella genera incertae sedis | 0.000 | 0.000 | 0.000 | 0.000 | 0.004 | 0.000 |
| Proteobacteria | Alphaproteobacteria | Rhizobiales 1 | Unclassified | Unclassified | 0.000 | 0.000 | 0.000 | 0.004 | 0.000 | 0.002 |
| Proteobacteria | Alphaproteobacteria | Rhizobiales 2 | alphaI cluster 2 | Unclassified | 0.000 | 0.000 | 0.000 | 0.004 | 0.000 | 0.000 |
| Proteobacteria | Alphaproteobacteria | Rhizobiales 2 | BCf3-20 | Unclassified | 0.000 | 0.000 | 0.000 | 0.004 | 0.000 | 0.000 |
| Proteobacteria | Alphaproteobacteria | Rhizobiales 2 | BCf3-20 | Unclassified | 0.001 | 0.000 | 0.000 | 0.000 | 0.000 | 0.000 |
| Proteobacteria | Alphaproteobacteria | Rhizobiales 2 | Beijerinckiaceae | Uncultured 2 | 0.001 | 0.000 | 0.000 | 0.014 | 0.000 | 0.000 |
| Proteobacteria | Alphaproteobacteria | Rhizobiales 2 | Beijerinckiaceae | Unclassified | 0.002 | 0.000 | 0.000 | 0.000 | 0.000 | 0.000 |
| Proteobacteria | Alphaproteobacteria | Rhizobiales 2 | Bradyrhizobiaceae | *Balneimonas* | 0.001 | 0.000 | 0.000 | 0.036 | 0.000 | 0.000 |
| Proteobacteria | Alphaproteobacteria | Rhizobiales 2 | Bradyrhizobiaceae | *Blastobacter* | 0.000 | 0.000 | 0.000 | 0.007 | 0.000 | 0.000 |
| Proteobacteria | Alphaproteobacteria | Rhizobiales 2 | Bradyrhizobiaceae | *Bosea* | 0.000 | 0.000 | 0.000 | 0.075 | 0.004 | 0.006 |
| Proteobacteria | Alphaproteobacteria | Rhizobiales 2 | Bradyrhizobiaceae | *Bradyrhizobium* 1 | 0.000 | 0.000 | 0.000 | 0.004 | 0.000 | 0.000 |
| Proteobacteria | Alphaproteobacteria | Rhizobiales 2 | Bradyrhizobiaceae | *Bradyrhizobium* 10 | 0.000 | 0.000 | 0.000 | 0.125 | 0.000 | 0.000 |
| Proteobacteria | Alphaproteobacteria | Rhizobiales 2 | Bradyrhizobiaceae | *Bradyrhizobium* 12 | 0.000 | 0.000 | 0.000 | 0.103 | 0.000 | 0.000 |
| Proteobacteria | Alphaproteobacteria | Rhizobiales 2 | Bradyrhizobiaceae | *Bradyrhizobium* 13 | 0.000 | 0.000 | 0.000 | 0.036 | 0.000 | 0.000 |
| Proteobacteria | Alphaproteobacteria | Rhizobiales 2 | Bradyrhizobiaceae | *Bradyrhizobium* 6 | 0.000 | 0.000 | 0.000 | 0.025 | 0.000 | 0.000 |
| Proteobacteria | Alphaproteobacteria | Rhizobiales 2 | Bradyrhizobiaceae | Unclassified | 0.000 | 0.000 | 0.000 | 0.082 | 0.000 | 0.001 |
| Proteobacteria | Alphaproteobacteria | Rhizobiales 2 | Bradyrhizobiaceae | Uncultured 1 | 0.001 | 0.000 | 0.000 | 0.004 | 0.000 | 0.000 |

**Table S2 continued.**

| | | | | | | | | | | |
|---|---|---|---|---|---|---|---|---|---|---|
| Proteobacteria | Alphaproteobacteria | Rhizobiales 2 | Bradyrhizobiaceae | Uncultured 2 | 0.001 | 0.000 | 0.000 | 0.004 | 0.000 | 0.000 |
| Proteobacteria | Alphaproteobacteria | Rhizobiales 2 | Bradyrhizobiaceae | Uncultured 3 | 0.000 | 0.000 | 0.000 | 0.018 | 0.000 | 0.000 |
| Proteobacteria | Alphaproteobacteria | Rhizobiales 2 | Bradyrhizobiaceae | Uncultured 4 | 0.001 | 0.000 | 0.000 | 0.004 | 0.000 | 0.000 |
| Proteobacteria | Alphaproteobacteria | Rhizobiales 2 | Bradyrhizobiaceae | Unclassified | 0.001 | 0.000 | 0.000 | 0.000 | 0.000 | 0.000 |
| Proteobacteria | Alphaproteobacteria | Rhizobiales 2 | CCSD-DF730-B17 | Unclassified | 0.000 | 0.000 | 0.000 | 0.007 | 0.000 | 0.000 |
| Proteobacteria | Alphaproteobacteria | Rhizobiales 2 | Hyphomicrobiaceae | *Rhodomicrobium* | 0.002 | 0.000 | 0.000 | 0.000 | 0.000 | 0.000 |
| Proteobacteria | Alphaproteobacteria | Rhizobiales 2 | Hyphomicrobiaceae 1 | *Ancalomicrobium* | 0.001 | 0.000 | 0.000 | 0.043 | 0.000 | 0.000 |
| Proteobacteria | Alphaproteobacteria | Rhizobiales 2 | Hyphomicrobiaceae 1 | Unclassified | 0.000 | 0.000 | 0.000 | 0.014 | 0.000 | 0.001 |
| Proteobacteria | Alphaproteobacteria | Rhizobiales 2 | Hyphomicrobiaceae 1 | Uncultured | 0.000 | 0.000 | 0.000 | 0.004 | 0.000 | 0.000 |
| Proteobacteria | Alphaproteobacteria | Rhizobiales 2 | Hyphomicrobiaceae 1 | Unclassified | 0.001 | 0.000 | 0.000 | 0.000 | 0.000 | 0.000 |
| Proteobacteria | Alphaproteobacteria | Rhizobiales 2 | Methylocystaceae | *Methylopila* | 0.000 | 0.000 | 0.000 | 0.000 | 0.004 | 0.000 |
| Proteobacteria | Alphaproteobacteria | Rhizobiales 2 | Methylocystaceae 1 | *Methylosinus* | 0.000 | 0.000 | 0.000 | 0.004 | 0.000 | 0.000 |
| Proteobacteria | Alphaproteobacteria | Rhizobiales 2 | Unclassified | Unclassified | 0.000 | 0.000 | 0.000 | 0.014 | 0.008 | 0.000 |
| Proteobacteria | Alphaproteobacteria | Rhizobiales 2 | Xanthobacteraceae | *Pseudolabrys* | 0.001 | 0.000 | 0.000 | 0.014 | 0.000 | 0.000 |
| Proteobacteria | Alphaproteobacteria | Rhizobiales 2 | Xanthobacteraceae | Unclassified | 0.000 | 0.000 | 0.000 | 0.004 | 0.000 | 0.000 |
| Proteobacteria | Alphaproteobacteria | Rhizobiales 2 | Xanthobacteraceae | Uncultured 1 | 0.002 | 0.000 | 0.000 | 0.064 | 0.000 | 0.001 |
| Proteobacteria | Alphaproteobacteria | Rhizobiales 2 | Xanthobacteraceae | Uncultured 3 | 0.000 | 0.000 | 0.000 | 0.011 | 0.000 | 0.000 |
| Proteobacteria | Alphaproteobacteria | Rhizobiales 3 | Candidatus Liberibacter | *Candidatus* Liberibacter | 0.000 | 0.000 | 0.000 | 0.007 | 0.000 | 0.001 |
| Proteobacteria | Alphaproteobacteria | Rhizobiales 3 | Hyphomicrobiaceae | *Filomicrobium* 1 | 0.000 | 0.000 | 0.000 | 0.004 | 0.000 | 0.001 |
| Proteobacteria | Alphaproteobacteria | Rhizobiales 3 | Hyphomicrobiaceae | *Hyphomicrobium* 1 | 0.001 | 0.000 | 0.000 | 0.007 | 0.000 | 0.000 |
| Proteobacteria | Alphaproteobacteria | Rhizobiales 3 | Hyphomicrobiaceae | *Hyphomicrobium* 2 | 0.004 | 0.064 | 0.000 | 0.025 | 0.000 | 0.001 |
| Proteobacteria | Alphaproteobacteria | Rhizobiales 3 | Hyphomicrobiaceae | Unclassified | 0.000 | 0.000 | 0.000 | 0.007 | 0.000 | 0.000 |
| Proteobacteria | Alphaproteobacteria | Rhizobiales 3 | Hyphomicrobiaceae | Uncultured | 0.000 | 0.000 | 0.000 | 0.007 | 0.004 | 0.000 |
| Proteobacteria | Alphaproteobacteria | Rhizobiales 3 | Methylobacteriaceae | *Meganema* | 0.001 | 0.000 | 0.000 | 0.000 | 0.000 | 0.000 |
| Proteobacteria | Alphaproteobacteria | Rhizobiales 3 | Methylobacteriaceae | Uncultured | 0.001 | 0.000 | 0.000 | 0.007 | 0.000 | 0.000 |
| Proteobacteria | Alphaproteobacteria | Rhizobiales 3 | Nordella Nordella | Unclassified | 0.000 | 0.032 | 0.000 | 0.029 | 0.000 | 0.001 |
| Proteobacteria | Alphaproteobacteria | Rhizobiales 3 | Nordella Nordella | Unclassified | 0.001 | 0.000 | 0.000 | 0.000 | 0.000 | 0.000 |

**Table S2 continued.**

| | | | | | | | | | | |
|---|---|---|---|---|---|---|---|---|---|---|
| Proteobacteria | Alphaproteobacteria | Rhizobiales 3 | Unclassified | Unclassified | 0.000 | 0.000 | 0.000 | 0.004 | 0.000 | 0.000 |
| Proteobacteria | Alphaproteobacteria | Rhodobacterales | Hyphomonadaceae | *Hirschia* | 0.002 | 0.000 | 0.000 | 0.029 | 0.000 | 0.000 |
| Proteobacteria | Alphaproteobacteria | Rhodobacterales | Hyphomonadaceae | *Hyphomonas* | 0.000 | 0.000 | 0.000 | 0.007 | 0.000 | 0.000 |
| Proteobacteria | Alphaproteobacteria | Rhodobacterales | Rhodobacteraceae | *Agaricicola* | 0.000 | 0.000 | 0.000 | 0.007 | 0.000 | 0.000 |
| Proteobacteria | Alphaproteobacteria | Rhodobacterales | Rhodobacteraceae | *Gemmobacter* | 0.000 | 0.000 | 0.000 | 0.004 | 0.000 | 0.000 |
| Proteobacteria | Alphaproteobacteria | Rhodobacterales | Rhodobacteraceae | *Labrenzia* | 0.000 | 0.000 | 0.000 | 0.004 | 0.000 | 0.000 |
| Proteobacteria | Alphaproteobacteria | Rhodobacterales | Rhodobacteraceae | *Nesiotobacter* | 0.001 | 0.000 | 0.000 | 0.000 | 0.000 | 0.000 |
| Proteobacteria | Alphaproteobacteria | Rhodobacterales | Rhodobacteraceae | *Oceanicola* | 0.001 | 0.000 | 0.000 | 0.000 | 0.000 | 0.000 |
| Proteobacteria | Alphaproteobacteria | Rhodobacterales | Rhodobacteraceae | *Paracoccus* | 0.001 | 0.000 | 0.000 | 0.036 | 0.004 | 0.000 |
| Proteobacteria | Alphaproteobacteria | Rhodobacterales | Rhodobacteraceae | *Pseudovibrio* | 0.001 | 0.000 | 0.000 | 0.000 | 0.000 | 0.000 |
| Proteobacteria | Alphaproteobacteria | Rhodobacterales | Rhodobacteraceae | *Rhodobacter* | 0.000 | 0.000 | 0.000 | 0.004 | 0.000 | 0.000 |
| Proteobacteria | Alphaproteobacteria | Rhodobacterales | Rhodobacteraceae | *Rhodovulum* | 0.002 | 0.000 | 0.000 | 0.004 | 0.000 | 0.001 |
| Proteobacteria | Alphaproteobacteria | Rhodobacterales | Rhodobacteraceae | *Roseibium* | 0.000 | 0.000 | 0.000 | 0.000 | 0.000 | 0.001 |
| Proteobacteria | Alphaproteobacteria | Rhodobacterales | Rhodobacteraceae | *Rubellimicrobium* | 0.000 | 0.000 | 0.000 | 0.043 | 0.000 | 0.000 |
| Proteobacteria | Alphaproteobacteria | Rhodobacterales | Rhodobacteraceae 1 | *Albimonas* | 0.000 | 0.000 | 0.000 | 0.004 | 0.004 | 0.000 |
| Proteobacteria | Alphaproteobacteria | Rhodobacterales | Rhodobacteraceae 1 | *Amaricoccus* | 0.000 | 0.000 | 0.000 | 0.000 | 0.004 | 0.000 |
| Proteobacteria | Alphaproteobacteria | Rhodobacterales | Rhodobacteraceae 1 | *Paracoccus* 1 | 0.000 | 0.000 | 0.000 | 0.004 | 0.000 | 0.000 |
| Proteobacteria | Alphaproteobacteria | Rhodobacterales | Rhodobacteraceae 1 | *Rhodobacter* 3 | 0.000 | 0.000 | 0.000 | 0.000 | 0.000 | 0.003 |
| Proteobacteria | Alphaproteobacteria | Rhodobacterales | Rhodobacteraceae 1 | *Rhodobacter* 5 | 0.002 | 0.000 | 0.000 | 0.000 | 0.000 | 0.000 |
| Proteobacteria | Alphaproteobacteria | Rhodobacterales | Rhodobacteraceae 1 | *Roseobacter* clade | 0.000 | 0.000 | 0.000 | 0.004 | 0.000 | 0.001 |
| Proteobacteria | Alphaproteobacteria | Rhodobacterales | Rhodobacteraceae 1 | *Shimia* | 0.000 | 0.000 | 0.000 | 0.004 | 0.000 | 0.000 |
| Proteobacteria | Alphaproteobacteria | Rhodobacterales | Rhodobacteraceae 1 | *Tropicimonas* | 0.002 | 0.000 | 0.000 | 0.000 | 0.000 | 0.001 |
| Proteobacteria | Alphaproteobacteria | Rhodobacterales | Rhodobacteraceae 1 | Unclassified | 0.000 | 0.000 | 0.013 | 0.000 | 0.000 | 0.000 |
| Proteobacteria | Alphaproteobacteria | Rhodobacterales | Rhodobacteraceae 1 | Uncultured 21 | 0.001 | 0.000 | 0.000 | 0.000 | 0.000 | 0.000 |
| Proteobacteria | Alphaproteobacteria | Rhodobacterales | Rhodobacteraceae 1 | Uncultured 23 | 0.000 | 0.000 | 0.000 | 0.004 | 0.000 | 0.000 |
| Proteobacteria | Alphaproteobacteria | Rhodobacterales | Rhodobacteraceae 1 | Uncultured 5 | 0.002 | 0.000 | 0.000 | 0.004 | 0.000 | 0.000 |
| Proteobacteria | Alphaproteobacteria | Rhodobacterales | Rhodobacteraceae 1 | Uncultured 7 | 0.004 | 0.000 | 0.000 | 0.004 | 0.000 | 0.000 |

**Table S2 continued.**

| | | | | | | | | | | |
|---|---|---|---|---|---|---|---|---|---|---|
| Proteobacteria | Alphaproteobacteria | Rhodobacterales | Rhodobacteraceae 3 | *Pannonibacter* | 0.000 | 0.000 | 0.000 | 0.007 | 0.000 | 0.000 |
| Proteobacteria | Alphaproteobacteria | Rhodobacterales | Rhodobacteraceae 3 | *Stappia* 2 | 0.001 | 0.000 | 0.000 | 0.000 | 0.000 | 0.000 |
| Proteobacteria | Alphaproteobacteria | Rhodobacterales | Rhodobacteraceae 3 | Unclassified | 0.000 | 0.000 | 0.000 | 0.000 | 0.000 | 0.001 |
| Proteobacteria | Alphaproteobacteria | Rhododbacterales | Rhodobacteraceae | *Stappia* | 0.001 | 0.000 | 0.000 | 0.004 | 0.000 | 0.000 |
| Proteobacteria | Alphaproteobacteria | Rhodospirillales | Acetobacteraceae | *Acetobacter* | 0.000 | 0.000 | 0.000 | 0.004 | 0.000 | 0.000 |
| Proteobacteria | Alphaproteobacteria | Rhodospirillales | Acetobacteraceae | *Acidisoma* | 0.000 | 0.000 | 0.000 | 0.007 | 0.000 | 0.000 |
| Proteobacteria | Alphaproteobacteria | Rhodospirillales | Acetobacteraceae | *Belnapia* | 0.000 | 0.000 | 0.000 | 0.007 | 0.000 | 0.000 |
| Proteobacteria | Alphaproteobacteria | Rhodospirillales | Acetobacteraceae | *Crenalkalicoccus* | 0.000 | 0.000 | 0.000 | 0.007 | 0.000 | 0.000 |
| Proteobacteria | Alphaproteobacteria | Rhodospirillales | Acetobacteraceae | *Komagataeibacter* | 0.001 | 0.000 | 0.000 | 0.000 | 0.000 | 0.000 |
| Proteobacteria | Alphaproteobacteria | Rhodospirillales | Acetobacteraceae | *Neokomagataea* | 0.002 | 0.000 | 0.000 | 0.004 | 0.000 | 0.000 |
| Proteobacteria | Alphaproteobacteria | Rhodospirillales | Acetobacteraceae | *Paracraurococcus* | 0.000 | 0.000 | 0.000 | 0.004 | 0.000 | 0.000 |
| Proteobacteria | Alphaproteobacteria | Rhodospirillales | Acetobacteraceae | *Rhodovastum* | 0.001 | 0.000 | 0.000 | 0.004 | 0.000 | 0.000 |
| Proteobacteria | Alphaproteobacteria | Rhodospirillales | Acetobacteraceae | *Roseomonas* | 0.001 | 0.000 | 0.000 | 0.021 | 0.000 | 0.000 |
| Proteobacteria | Alphaproteobacteria | Rhodospirillales | Acetobacteraceae | *Rubritepida* | 0.000 | 0.000 | 0.000 | 0.007 | 0.000 | 0.000 |
| Proteobacteria | Alphaproteobacteria | Rhodospirillales | Geminicoccaceae | *Geminicoccus* | 0.000 | 0.000 | 0.000 | 0.004 | 0.000 | 0.000 |
| Proteobacteria | Alphaproteobacteria | Rhodospirillales | Rhodospirillaceae | *Aestuariispira* | 0.000 | 0.000 | 0.000 | 0.000 | 0.004 | 0.000 |
| Proteobacteria | Alphaproteobacteria | Rhodospirillales | Rhodospirillaceae | *Azospirillum* | 0.000 | 0.000 | 0.000 | 0.171 | 0.000 | 0.002 |
| Proteobacteria | Alphaproteobacteria | Rhodospirillales | Rhodospirillaceae | *Defluviicoccus* | 0.002 | 0.032 | 0.013 | 0.007 | 0.000 | 0.000 |
| Proteobacteria | Alphaproteobacteria | Rhodospirillales | Rhodospirillaceae | *Dongia* | 0.004 | 0.000 | 0.000 | 0.018 | 0.000 | 0.003 |
| Proteobacteria | Alphaproteobacteria | Rhodospirillales | Rhodospirillaceae | *Elstera* | 0.001 | 0.000 | 0.000 | 0.000 | 0.004 | 0.000 |
| Proteobacteria | Alphaproteobacteria | Rhodospirillales | Rhodospirillaceae | *Fodinicurvata* | 0.001 | 0.000 | 0.000 | 0.000 | 0.004 | 0.000 |
| Proteobacteria | Alphaproteobacteria | Rhodospirillales | Rhodospirillaceae | *Haematospirillum* | 0.009 | 0.000 | 0.000 | 0.004 | 0.012 | 0.002 |
| Proteobacteria | Alphaproteobacteria | Rhodospirillales | Rhodospirillaceae | *Inquilinus* | 0.002 | 0.000 | 0.000 | 0.007 | 0.000 | 0.000 |
| Proteobacteria | Alphaproteobacteria | Rhodospirillales | Rhodospirillaceae | *Limibacillus* | 0.001 | 0.000 | 0.000 | 0.004 | 0.012 | 0.000 |
| Proteobacteria | Alphaproteobacteria | Rhodospirillales | Rhodospirillaceae | *Magnetospira* | 0.000 | 0.000 | 0.000 | 0.007 | 0.000 | 0.000 |
| Proteobacteria | Alphaproteobacteria | Rhodospirillales | Rhodospirillaceae | *Magnetospirillum* | 0.001 | 0.000 | 0.000 | 0.000 | 0.000 | 0.000 |
| Proteobacteria | Alphaproteobacteria | Rhodospirillales | Rhodospirillaceae | *Magnetovibrio* | 0.012 | 0.032 | 0.000 | 0.018 | 0.000 | 0.000 |

**Table S2 continued.**

| | | | | | | | | | | |
|---|---|---|---|---|---|---|---|---|---|---|
| Proteobacteria | Alphaproteobacteria | Rhodospirillales | Rhodospirillaceae | *Marispirillum* | 0.002 | 0.000 | 0.000 | 0.007 | 0.008 | 0.003 |
| Proteobacteria | Alphaproteobacteria | Rhodospirillales | Rhodospirillaceae | *Nisaea* | 0.001 | 0.000 | 0.000 | 0.000 | 0.000 | 0.000 |
| Proteobacteria | Alphaproteobacteria | Rhodospirillales | Rhodospirillaceae | *Novispirillum* | 0.038 | 0.096 | 0.000 | 0.032 | 0.008 | 0.010 |
| Proteobacteria | Alphaproteobacteria | Rhodospirillales | Rhodospirillaceae | *Oceanibaculum* | 0.001 | 0.000 | 0.000 | 0.000 | 0.000 | 0.000 |
| Proteobacteria | Alphaproteobacteria | Rhodospirillales | Rhodospirillaceae | *Pelagibius* | 0.030 | 0.032 | 0.000 | 0.000 | 0.004 | 0.001 |
| Proteobacteria | Alphaproteobacteria | Rhodospirillales | Rhodospirillaceae | *Rhodospirillum* | 0.001 | 0.032 | 0.000 | 0.000 | 0.000 | 0.000 |
| Proteobacteria | Alphaproteobacteria | Rhodospirillales | Rhodospirillaceae | *Rhodovibrio* | 0.004 | 0.000 | 0.000 | 0.004 | 0.012 | 0.000 |
| Proteobacteria | Alphaproteobacteria | Rhodospirillales | Rhodospirillaceae | *Skermanella* | 0.002 | 0.000 | 0.000 | 0.014 | 0.008 | 0.000 |
| Proteobacteria | Alphaproteobacteria | Rhodospirillales | Rhodospirillaceae | *Tagaea* | 0.000 | 0.000 | 0.000 | 0.000 | 0.004 | 0.000 |
| Proteobacteria | Alphaproteobacteria | Rhodospirillales | Rhodospirillaceae | *Taonella* | 0.000 | 0.000 | 0.000 | 0.007 | 0.000 | 0.000 |
| Proteobacteria | Alphaproteobacteria | Rhodospirillales | Rhodospirillaceae | *Thalassospira* | 0.657 | 0.000 | 0.013 | 1.055 | 1.619 | 0.071 |
| Proteobacteria | Alphaproteobacteria | Rhodospirillales | Rhodospirillaceae | *Tistrella* | 0.001 | 0.000 | 0.000 | 0.000 | 0.000 | 0.000 |
| Proteobacteria | Alphaproteobacteria | Rhodospirillales | Rhodospirillaceae | Uncultured 1 | 0.001 | 0.000 | 0.000 | 0.004 | 0.004 | 0.001 |
| Proteobacteria | Alphaproteobacteria | Rhodospirillales | Unclassified | *Enhydrobacter* | 0.001 | 0.000 | 0.000 | 0.000 | 0.004 | 0.001 |
| Proteobacteria | Alphaproteobacteria | Rhodospirillales | Unclassified | *Reyranella* | 0.000 | 0.000 | 0.000 | 0.007 | 0.000 | 0.000 |
| Proteobacteria | Alphaproteobacteria | Rhodospirillales 1 | Acetobacteraceae | *Acidiphilium* | 0.001 | 0.000 | 0.000 | 0.007 | 0.000 | 0.000 |
| Proteobacteria | Alphaproteobacteria | Rhodospirillales 1 | Acetobacteraceae | *Acidocella* | 0.001 | 0.000 | 0.000 | 0.004 | 0.000 | 0.000 |
| Proteobacteria | Alphaproteobacteria | Rhodospirillales 1 | Acetobacteraceae | *Craurococcus* | 0.000 | 0.000 | 0.000 | 0.004 | 0.000 | 0.000 |
| Proteobacteria | Alphaproteobacteria | Rhodospirillales 1 | Acetobacteraceae | *Gluconobacter* | 0.000 | 0.000 | 0.000 | 0.000 | 0.000 | 0.001 |
| Proteobacteria | Alphaproteobacteria | Rhodospirillales 1 | Acetobacteraceae | Unclassified | 0.000 | 0.000 | 0.000 | 0.021 | 0.000 | 0.000 |
| Proteobacteria | Alphaproteobacteria | Rhodospirillales 1 | Acetobacteraceae | Uncultured 1 | 0.000 | 0.000 | 0.000 | 0.000 | 0.000 | 0.001 |
| Proteobacteria | Alphaproteobacteria | Rhodospirillales 1 | Rhodospirillaceae | marine group AEGEAN-169 | 0.001 | 0.000 | 0.000 | 0.000 | 0.000 | 0.000 |
| Proteobacteria | Alphaproteobacteria | Rhodospirillales 1 | Rhodospirillaceae | Unclassified | 0.000 | 0.000 | 0.000 | 0.000 | 0.000 | 0.002 |
| Proteobacteria | Alphaproteobacteria | Rhodospirillales 1 | Rhodospirillaceae | Uncultured 1 | 0.001 | 0.000 | 0.013 | 0.000 | 0.000 | 0.001 |
| Proteobacteria | Alphaproteobacteria | Rhodospirillales 1 | Rhodospirillaceae | Uncultured 2 | 0.453 | 0.702 | 0.000 | 0.460 | 0.077 | 0.142 |
| Proteobacteria | Alphaproteobacteria | Rhodospirillales 1 | Rhodospirillaceae | Uncultured a | 0.015 | 0.032 | 0.000 | 0.014 | 0.000 | 0.000 |
| Proteobacteria | Alphaproteobacteria | Rhodospirillales 1 | Rhodospirillaceae | Unclassified | 0.002 | 0.000 | 0.000 | 0.000 | 0.000 | 0.000 |

**Table S2 continued.**

| | | | | | | | | | | |
|---|---|---|---|---|---|---|---|---|---|---|
| Proteobacteria | Alphaproteobacteria | Rhodospirillales 1 | Unclassified | Unclassified | 0.000 | 0.000 | 0.000 | 0.007 | 0.000 | 0.000 |
| Proteobacteria | Alphaproteobacteria | Rhodospirillales 2 | Candidatus Alysiosphaera | *Candidatus* Alysiosphaera | 0.001 | 0.000 | 0.000 | 0.004 | 0.000 | 0.000 |
| Proteobacteria | Alphaproteobacteria | Rhodospirillales 2 | DA111 | Unclassified | 0.000 | 0.000 | 0.000 | 0.018 | 0.000 | 0.001 |
| Proteobacteria | Alphaproteobacteria | Rhodospirillales 2 | DA111 | Unclassified | 0.002 | 0.000 | 0.000 | 0.000 | 0.000 | 0.000 |
| Proteobacteria | Alphaproteobacteria | Rhodospirillales 2 | I-10 | Unclassified | 0.000 | 0.000 | 0.000 | 0.018 | 0.000 | 0.000 |
| Proteobacteria | Alphaproteobacteria | Rhodospirillales 2 | JG37-AG-20 | Unclassified | 0.000 | 0.032 | 0.000 | 0.004 | 0.000 | 0.000 |
| Proteobacteria | Alphaproteobacteria | Rhodospirillales 2 | JG37-AG-20 | Unclassified | 0.004 | 0.000 | 0.000 | 0.000 | 0.000 | 0.000 |
| Proteobacteria | Alphaproteobacteria | Rhodospirillales 2 | KCM-B-15 | Unclassified | 0.000 | 0.000 | 0.000 | 0.014 | 0.000 | 0.000 |
| Proteobacteria | Alphaproteobacteria | Rhodospirillales 2 | MNH4 | Unclassified | 0.000 | 0.000 | 0.000 | 0.057 | 0.004 | 0.001 |
| Proteobacteria | Alphaproteobacteria | Rhodospirillales 2 | MNH4 | Unclassified | 0.009 | 0.000 | 0.000 | 0.000 | 0.000 | 0.000 |
| Proteobacteria | Alphaproteobacteria | Rhodospirillales 2 | Rhodospirillaceae 2 | *Azospirillum* 1 | 0.000 | 0.000 | 0.000 | 0.007 | 0.000 | 0.000 |
| Proteobacteria | Alphaproteobacteria | Rhodospirillales 2 | Rhodospirillaceae 2 | Unclassified | 0.000 | 0.000 | 0.000 | 0.036 | 0.000 | 0.000 |
| Proteobacteria | Alphaproteobacteria | Rhodospirillales 2 | Unclassified | Unclassified | 0.000 | 0.000 | 0.000 | 0.032 | 0.000 | 0.000 |
| Proteobacteria | Alphaproteobacteria | Rhodospirillales 2 | Unclassified | Unclassified | 0.001 | 0.000 | 0.000 | 0.000 | 0.000 | 0.000 |
| Proteobacteria | Alphaproteobacteria | Rhodospirillales 2 | wr0007 | Unclassified | 0.000 | 0.000 | 0.000 | 0.043 | 0.000 | 0.000 |
| Proteobacteria | Alphaproteobacteria | Rhodospirillales 2 | wr0007 | Unclassified | 0.005 | 0.000 | 0.000 | 0.000 | 0.000 | 0.000 |
| Proteobacteria | Alphaproteobacteria | Rickettsiales | Anaplasmataceae | *Anaplasma* | 0.000 | 0.000 | 0.000 | 0.000 | 0.000 | 0.001 |
| Proteobacteria | Alphaproteobacteria | Rickettsiales | Anaplasmataceae | *Candidatus* Xenohaliotis | 0.001 | 0.000 | 0.000 | 0.000 | 0.000 | 0.000 |
| Proteobacteria | Alphaproteobacteria | Rickettsiales | Anaplasmataceae | *Ehrlichia* | 0.002 | 0.000 | 0.039 | 0.000 | 0.012 | 0.008 |
| Proteobacteria | Alphaproteobacteria | Rickettsiales | Anaplasmataceae | *Neorickettsia* | 0.002 | 0.000 | 0.000 | 0.004 | 0.004 | 0.001 |
| Proteobacteria | Alphaproteobacteria | Rickettsiales | Anaplasmataceae | *Unclassified* | 0.000 | 0.000 | 0.000 | 0.000 | 0.000 | 0.003 |
| Proteobacteria | Alphaproteobacteria | Rickettsiales | Anaplasmataceae | *Wolbachia* | 2.188 | 8.112 | 44.321 | 2.234 | 15.303 | 31.272 |
| Proteobacteria | Alphaproteobacteria | Rickettsiales | Candidatus Captivus | *Candidatus* Captivus | 0.019 | 0.000 | 0.000 | 0.014 | 0.000 | 0.003 |
| Proteobacteria | Alphaproteobacteria | Rickettsiales | Candidatus Hepatincola | *Candidatus* Hepatincola | 0.004 | 0.000 | 0.000 | 0.007 | 0.000 | 0.000 |
| Proteobacteria | Alphaproteobacteria | Rickettsiales | Candidatus Midichloria | *Candidatus* Midichloria | 0.000 | 0.000 | 0.000 | 0.004 | 0.000 | 0.000 |
| Proteobacteria | Alphaproteobacteria | Rickettsiales | Candidatus Odyssella | *Candidatus* Odyssella | 0.002 | 0.000 | 0.000 | 0.004 | 0.000 | 0.000 |
| Proteobacteria | Alphaproteobacteria | Rickettsiales | LWSR-14 | Unclassified | 0.000 | 0.000 | 0.026 | 0.000 | 0.004 | 0.003 |

**Table S2 continued.**

| | | | | | | | | | | |
|---|---|---|---|---|---|---|---|---|---|---|
| Proteobacteria | Alphaproteobacteria | Rickettsiales | Mitochondria | Mitochondria | 0.017 | 0.000 | 0.000 | 0.061 | 0.004 | 0.003 |
| Proteobacteria | Alphaproteobacteria | Rickettsiales | Rickettsiaceae | *Orientia* | 0.000 | 0.000 | 0.000 | 0.000 | 0.004 | 0.000 |
| Proteobacteria | Alphaproteobacteria | Rickettsiales | Rickettsiaceae | *Rickettsia* | 0.002 | 0.000 | 0.000 | 0.014 | 0.004 | 0.002 |
| Proteobacteria | Alphaproteobacteria | Rickettsiales | Rickettsiaceae | Unclassified | 0.000 | 0.000 | 0.000 | 0.004 | 0.000 | 0.002 |
| Proteobacteria | Alphaproteobacteria | Rickettsiales | SAR116 | Unclassified | 0.000 | 0.000 | 0.000 | 0.000 | 0.000 | 0.001 |
| Proteobacteria | Alphaproteobacteria | Rickettsiales | SAR116 | Unclassified | 0.001 | 0.000 | 0.000 | 0.000 | 0.000 | 0.000 |
| Proteobacteria | Alphaproteobacteria | Rickettsiales | SM2D12 | Unclassified | 0.000 | 0.000 | 0.000 | 0.007 | 0.000 | 0.000 |
| Proteobacteria | Alphaproteobacteria | Rickettsiales | SM2D12 | Unclassified | 0.002 | 0.000 | 0.000 | 0.000 | 0.000 | 0.000 |
| Proteobacteria | Alphaproteobacteria | Rickettsiales | Unclassified | Unclassified | 0.000 | 0.032 | 0.000 | 0.014 | 0.000 | 0.001 |
| Proteobacteria | Alphaproteobacteria | SAR11 clade | Surface 1 | Unclassified | 0.001 | 0.000 | 0.000 | 0.000 | 0.000 | 0.000 |
| Proteobacteria | Alphaproteobacteria | SAR11 clade | Unclassified | Unclassified | 0.002 | 0.000 | 0.000 | 0.000 | 0.000 | 0.000 |
| Proteobacteria | Alphaproteobacteria | Sneathiellales | Sneathiellaceae | *Sneathiella* | 0.000 | 0.000 | 0.000 | 0.004 | 0.000 | 0.000 |
| Proteobacteria | Alphaproteobacteria | Sphingomonadales | Ellin6055 | Unclassified | 0.000 | 0.000 | 0.000 | 0.021 | 0.000 | 0.001 |
| Proteobacteria | Alphaproteobacteria | Sphingomonadales | Erythrobacteraceae | *Altererythrobacter* | 0.004 | 0.000 | 0.000 | 0.029 | 0.000 | 0.000 |
| Proteobacteria | Alphaproteobacteria | Sphingomonadales | Erythrobacteraceae | *Erythrobacter* | 0.004 | 0.000 | 0.000 | 0.071 | 0.000 | 0.000 |
| Proteobacteria | Alphaproteobacteria | Sphingomonadales | Erythrobacteraceae | *Porphyrobacter* | 0.001 | 0.000 | 0.000 | 0.007 | 0.000 | 0.000 |
| Proteobacteria | Alphaproteobacteria | Sphingomonadales | Erythrobacteraceae | Unclassified | 0.000 | 0.000 | 0.000 | 0.004 | 0.000 | 0.000 |
| Proteobacteria | Alphaproteobacteria | Sphingomonadales | Erythrobacteraceae | Uncultured 2 | 0.000 | 0.000 | 0.000 | 0.004 | 0.000 | 0.000 |
| Proteobacteria | Alphaproteobacteria | Sphingomonadales | Erythrobacteraceae | Uncultured 3 | 0.000 | 0.000 | 0.000 | 0.018 | 0.000 | 0.000 |
| Proteobacteria | Alphaproteobacteria | Sphingomonadales | JG34-KF-161 | Unclassified | 0.000 | 0.000 | 0.000 | 0.018 | 0.000 | 0.000 |
| Proteobacteria | Alphaproteobacteria | Sphingomonadales | Sphingomonadaceae | *Hephaestia* | 0.000 | 0.000 | 0.000 | 0.004 | 0.000 | 0.000 |
| Proteobacteria | Alphaproteobacteria | Sphingomonadales | Sphingomonadaceae | *Novosphingobium* | 0.001 | 0.000 | 0.000 | 0.053 | 0.008 | 0.000 |
| Proteobacteria | Alphaproteobacteria | Sphingomonadales | Sphingomonadaceae | *Sandaracinobacter* | 0.001 | 0.000 | 0.000 | 0.004 | 0.000 | 0.000 |
| Proteobacteria | Alphaproteobacteria | Sphingomonadales | Sphingomonadaceae | *Sandarakinorhabdus* | 0.000 | 0.000 | 0.000 | 0.000 | 0.000 | 0.001 |
| Proteobacteria | Alphaproteobacteria | Sphingomonadales | Sphingomonadaceae | *Sphingobium* | 0.001 | 0.000 | 0.000 | 0.082 | 0.000 | 0.000 |
| Proteobacteria | Alphaproteobacteria | Sphingomonadales | Sphingomonadaceae | *Sphingobium* 1 | 0.000 | 0.000 | 0.000 | 0.046 | 0.000 | 0.000 |
| Proteobacteria | Alphaproteobacteria | Sphingomonadales | Sphingomonadaceae | *Sphingomicrobium* | 0.001 | 0.000 | 0.000 | 0.000 | 0.000 | 0.000 |

**Table S2 continued.**

| | | | | | | | | | | |
|---|---|---|---|---|---|---|---|---|---|---|
| Proteobacteria | Alphaproteobacteria | Sphingomonadales | Sphingomonadaceae | *Sphingomonas* | 0.006 | 0.000 | 0.000 | 0.645 | 0.004 | 0.001 |
| Proteobacteria | Alphaproteobacteria | Sphingomonadales | Sphingomonadaceae | *Sphingomonas* 1 | 0.000 | 0.000 | 0.000 | 0.004 | 0.000 | 0.000 |
| Proteobacteria | Alphaproteobacteria | Sphingomonadales | Sphingomonadaceae | *Sphingomonas* 2 | 0.002 | 0.000 | 0.013 | 0.289 | 0.000 | 0.001 |
| Proteobacteria | Alphaproteobacteria | Sphingomonadales | Sphingomonadaceae | *Sphingomonas* 3 | 0.004 | 0.000 | 0.000 | 0.242 | 0.000 | 0.001 |
| Proteobacteria | Alphaproteobacteria | Sphingomonadales | Sphingomonadaceae | *Sphingopyxis* | 0.000 | 0.000 | 0.000 | 0.011 | 0.000 | 0.000 |
| Proteobacteria | Alphaproteobacteria | Sphingomonadales | Sphingomonadaceae | *Sphingopyxis* 1 | 0.000 | 0.000 | 0.000 | 0.018 | 0.000 | 0.005 |
| Proteobacteria | Alphaproteobacteria | Sphingomonadales | Sphingomonadaceae | *Sphingosinicella* | 0.000 | 0.000 | 0.000 | 0.025 | 0.000 | 0.000 |
| Proteobacteria | Alphaproteobacteria | Sphingomonadales | Unclassified | Unclassified | 0.000 | 0.000 | 0.000 | 0.089 | 0.000 | 0.000 |
| Proteobacteria | Alphaproteobacteria | Sphinomonadales | Sphinomonadaceae | *Blastomonas* | 0.000 | 0.000 | 0.000 | 0.021 | 0.000 | 0.000 |
| Proteobacteria | Alphaproteobacteria | Unclassified | Unclassified | Unclassified | 0.000 | 0.000 | 0.000 | 0.004 | 0.004 | 0.000 |
| Proteobacteria | Betaproteobacteria | B1-7BS | Unclassified | Unclassified | 0.000 | 0.000 | 0.000 | 0.000 | 0.008 | 0.000 |
| Proteobacteria | Betaproteobacteria | Burkholderiales | Alcaligenaceae | *Advenella* | 0.000 | 0.000 | 0.000 | 0.007 | 0.000 | 0.000 |
| Proteobacteria | Betaproteobacteria | Burkholderiales | Alcaligenaceae | *Azohydromonas* | 0.000 | 0.000 | 0.000 | 0.039 | 0.000 | 0.000 |
| Proteobacteria | Betaproteobacteria | Burkholderiales | Alcaligenaceae | *Candidimonas* | 0.000 | 0.000 | 0.000 | 0.014 | 0.000 | 0.000 |
| Proteobacteria | Betaproteobacteria | Burkholderiales | Alcaligenaceae | *Castellaniella* | 0.000 | 0.000 | 0.000 | 0.018 | 0.000 | 0.000 |
| Proteobacteria | Betaproteobacteria | Burkholderiales | Alcaligenaceae | *Derxia* | 0.000 | 0.000 | 0.013 | 0.004 | 0.000 | 0.000 |
| Proteobacteria | Betaproteobacteria | Burkholderiales | Alcaligenaceae | *Pelistega* | 0.000 | 0.000 | 0.000 | 0.004 | 0.000 | 0.000 |
| Proteobacteria | Betaproteobacteria | Burkholderiales | Alcaligenaceae | *Pigmentiphaga* | 0.000 | 0.000 | 0.000 | 0.014 | 0.000 | 0.000 |
| Proteobacteria | Betaproteobacteria | Burkholderiales | Alcaligenaceae | *Pusillimonas* | 0.000 | 0.000 | 0.000 | 0.004 | 0.000 | 0.000 |
| Proteobacteria | Betaproteobacteria | Burkholderiales | Alcaligenaceae 1 | *Achromobacter* 1 | 0.000 | 0.000 | 0.000 | 0.004 | 0.000 | 0.000 |
| Proteobacteria | Betaproteobacteria | Burkholderiales | Alcaligenaceae 1 | *Achromobacter* 2 | 0.000 | 0.000 | 0.000 | 0.086 | 0.000 | 0.000 |
| Proteobacteria | Betaproteobacteria | Burkholderiales | Alcaligenaceae 1 | *Achromobacter* 3 | 0.000 | 0.000 | 0.013 | 0.036 | 0.000 | 0.000 |
| Proteobacteria | Betaproteobacteria | Burkholderiales | Alcaligenaceae 1 | *Bordetella* 2 | 0.000 | 0.000 | 0.000 | 0.029 | 0.000 | 0.000 |
| Proteobacteria | Betaproteobacteria | Burkholderiales | Alcaligenaceae 1 | GKS98 freshwater group | 0.000 | 0.000 | 0.000 | 0.014 | 0.000 | 0.000 |
| Proteobacteria | Betaproteobacteria | Burkholderiales | Alcaligenaceae 1 | *Kinetoplastibacterium* | 0.000 | 0.000 | 0.000 | 0.004 | 0.000 | 0.000 |
| Proteobacteria | Betaproteobacteria | Burkholderiales | Alcaligenaceae 1 | *Tetrathiobacter* | 0.000 | 0.032 | 0.000 | 0.021 | 0.000 | 0.000 |
| Proteobacteria | Betaproteobacteria | Burkholderiales | Alcaligenaceae 1 | Unclassified | 0.000 | 0.032 | 0.026 | 0.538 | 0.000 | 0.000 |

**Table S2 continued.**

| | | | | | | | | | | |
|---|---|---|---|---|---|---|---|---|---|---|
| Proteobacteria | Betaproteobacteria | Burkholderiales | Alcaligenaceae 1 | Uncultured 2 | 0.000 | 0.000 | 0.000 | 0.004 | 0.000 | 0.000 |
| Proteobacteria | Betaproteobacteria | Burkholderiales | Alcaligenaceae 1 | Uncultured 3 | 0.000 | 0.000 | 0.000 | 0.004 | 0.000 | 0.000 |
| Proteobacteria | Betaproteobacteria | Burkholderiales | Alcanigenaceae | *Ampullimonas* | 0.000 | 0.000 | 0.000 | 0.007 | 0.000 | 0.000 |
| Proteobacteria | Betaproteobacteria | Burkholderiales | Alcanigenaceae | *Bordetella* | 0.001 | 0.064 | 0.064 | 1.639 | 0.000 | 0.000 |
| Proteobacteria | Betaproteobacteria | Burkholderiales | Alkaligenaceae | *Achromobacter* | 0.000 | 0.000 | 0.000 | 0.021 | 0.000 | 0.000 |
| Proteobacteria | Betaproteobacteria | Burkholderiales | Burkholderiaceae | *Burkholderia* | 0.000 | 0.032 | 0.064 | 0.510 | 0.000 | 0.001 |
| Proteobacteria | Betaproteobacteria | Burkholderiales | Burkholderiaceae | *Cupriavidus* | 0.009 | 0.000 | 0.052 | 0.734 | 0.000 | 0.005 |
| Proteobacteria | Betaproteobacteria | Burkholderiales | Burkholderiaceae | *Pandoraea* | 0.005 | 0.000 | 0.052 | 1.090 | 0.000 | 0.000 |
| Proteobacteria | Betaproteobacteria | Burkholderiales | Burkholderiaceae | *Paraburkholderia* | 0.000 | 0.000 | 0.000 | 0.036 | 0.000 | 0.000 |
| Proteobacteria | Betaproteobacteria | Burkholderiales | Burkholderiaceae | *Polynucleobacter* | 0.002 | 0.000 | 0.000 | 0.004 | 0.000 | 0.000 |
| Proteobacteria | Betaproteobacteria | Burkholderiales | Burkholderiaceae | *Ralstonia* | 0.033 | 0.000 | 0.142 | 1.753 | 0.012 | 0.017 |
| Proteobacteria | Betaproteobacteria | Burkholderiales | Burkholderiaceae 1 | *Burkholderia* 1 | 0.006 | 0.000 | 0.013 | 0.139 | 0.008 | 0.002 |
| Proteobacteria | Betaproteobacteria | Burkholderiales | Burkholderiaceae 1 | *Burkholderia* 10 | 0.000 | 0.000 | 0.000 | 0.007 | 0.000 | 0.000 |
| Proteobacteria | Betaproteobacteria | Burkholderiales | Burkholderiaceae 1 | *Burkholderia* 15 | 0.000 | 0.000 | 0.000 | 0.011 | 0.000 | 0.000 |
| Proteobacteria | Betaproteobacteria | Burkholderiales | Burkholderiaceae 1 | *Limnobacter* | 0.001 | 0.000 | 0.000 | 0.007 | 0.000 | 0.000 |
| Proteobacteria | Betaproteobacteria | Burkholderiales | Burkholderiaceae 1 | Unclassified | 0.000 | 0.000 | 0.013 | 0.014 | 0.000 | 0.000 |
| Proteobacteria | Betaproteobacteria | Burkholderiales | Burkholderiaceae 1 | Unclassified | 0.002 | 0.000 | 0.000 | 0.000 | 0.000 | 0.000 |
| Proteobacteria | Betaproteobacteria | Burkholderiales | Comamonadaceae | *Acidovorax* | 0.000 | 0.000 | 0.000 | 0.025 | 0.000 | 0.000 |
| Proteobacteria | Betaproteobacteria | Burkholderiales | Comamonadaceae | *Acidovorax-Hylemonella* | 0.001 | 0.000 | 0.000 | 0.036 | 0.000 | 0.000 |
| Proteobacteria | Betaproteobacteria | Burkholderiales | Comamonadaceae | *Acidovorax-Verminephrobacter* | 0.001 | 0.000 | 0.000 | 0.114 | 0.000 | 0.001 |
| Proteobacteria | Betaproteobacteria | Burkholderiales | Comamonadaceae | *Alicycliphilus* | 0.000 | 0.000 | 0.000 | 0.011 | 0.000 | 0.000 |
| Proteobacteria | Betaproteobacteria | Burkholderiales | Comamonadaceae | *Azohydromonas* 2 | 0.000 | 0.000 | 0.000 | 0.011 | 0.000 | 0.000 |
| Proteobacteria | Betaproteobacteria | Burkholderiales | Comamonadaceae | *Brachymonas* 1 | 0.000 | 0.000 | 0.000 | 0.004 | 0.000 | 0.000 |
| Proteobacteria | Betaproteobacteria | Burkholderiales | Comamonadaceae | *Caenimonas* | 0.000 | 0.000 | 0.000 | 0.007 | 0.000 | 0.000 |
| Proteobacteria | Betaproteobacteria | Burkholderiales | Comamonadaceae | *Comamonas* | 0.000 | 0.000 | 0.000 | 0.043 | 0.000 | 0.000 |
| Proteobacteria | Betaproteobacteria | Burkholderiales | Comamonadaceae | *Comamonas* 1 | 0.000 | 0.000 | 0.000 | 0.007 | 0.000 | 0.000 |
| Proteobacteria | Betaproteobacteria | Burkholderiales | Comamonadaceae | *Comamonas* 5 | 0.004 | 0.000 | 0.000 | 0.253 | 0.000 | 0.000 |

**Table S2 continued.**

| | | | | | | | | | | |
|---|---|---|---|---|---|---|---|---|---|---|
| Proteobacteria | Betaproteobacteria | Burkholderiales | Comamonadaceae | *Comamonas* 6 | 0.002 | 0.000 | 0.000 | 0.139 | 0.000 | 0.000 |
| Proteobacteria | Betaproteobacteria | Burkholderiales | Comamonadaceae | *Comamonas* 8 | 0.000 | 0.000 | 0.000 | 0.007 | 0.004 | 0.000 |
| Proteobacteria | Betaproteobacteria | Burkholderiales | Comamonadaceae | *Delftia* | 0.000 | 0.000 | 0.000 | 0.004 | 0.000 | 0.000 |
| Proteobacteria | Betaproteobacteria | Burkholderiales | Comamonadaceae | *Diaphorobacter* | 0.000 | 0.000 | 0.000 | 0.007 | 0.000 | 0.000 |
| Proteobacteria | Betaproteobacteria | Burkholderiales | Comamonadaceae | *Giesbergeria* | 0.000 | 0.000 | 0.000 | 0.004 | 0.000 | 0.000 |
| Proteobacteria | Betaproteobacteria | Burkholderiales | Comamonadaceae | *Hydrogenophaga* | 0.001 | 0.000 | 0.000 | 0.011 | 0.008 | 0.000 |
| Proteobacteria | Betaproteobacteria | Burkholderiales | Comamonadaceae | *Hydrogenophaga* 1 | 0.000 | 0.000 | 0.000 | 0.000 | 0.004 | 0.000 |
| Proteobacteria | Betaproteobacteria | Burkholderiales | Comamonadaceae | *Hydrogenophaga* 2 | 0.000 | 0.000 | 0.000 | 0.000 | 0.004 | 0.000 |
| Proteobacteria | Betaproteobacteria | Burkholderiales | Comamonadaceae | *Hydrogenophaga* 3 | 0.000 | 0.000 | 0.000 | 0.004 | 0.000 | 0.000 |
| Proteobacteria | Betaproteobacteria | Burkholderiales | Comamonadaceae | *Ideonella* 1 | 0.000 | 0.000 | 0.000 | 0.004 | 0.000 | 0.000 |
| Proteobacteria | Betaproteobacteria | Burkholderiales | Comamonadaceae | *Ideonella* 2 | 0.000 | 0.000 | 0.000 | 0.004 | 0.000 | 0.000 |
| Proteobacteria | Betaproteobacteria | Burkholderiales | Comamonadaceae | *Kinneretia* | 0.000 | 0.000 | 0.000 | 0.004 | 0.000 | 0.000 |
| Proteobacteria | Betaproteobacteria | Burkholderiales | Comamonadaceae | *Lampropedia* | 0.000 | 0.000 | 0.000 | 0.004 | 0.000 | 0.000 |
| Proteobacteria | Betaproteobacteria | Burkholderiales | Comamonadaceae | *Leptothrix* 2 | 0.002 | 0.000 | 0.000 | 0.007 | 0.000 | 0.000 |
| Proteobacteria | Betaproteobacteria | Burkholderiales | Comamonadaceae | *Methylibium* 1 | 0.009 | 0.000 | 0.000 | 0.082 | 0.016 | 0.001 |
| Proteobacteria | Betaproteobacteria | Burkholderiales | Comamonadaceae | *Methylibium* 2 | 0.000 | 0.000 | 0.000 | 0.007 | 0.000 | 0.000 |
| Proteobacteria | Betaproteobacteria | Burkholderiales | Comamonadaceae | *Ottowia* | 0.000 | 0.000 | 0.000 | 0.021 | 0.000 | 0.000 |
| Proteobacteria | Betaproteobacteria | Burkholderiales | Comamonadaceae | *Pelomonas* | 0.000 | 0.000 | 0.000 | 0.046 | 0.000 | 0.000 |
| Proteobacteria | Betaproteobacteria | Burkholderiales | Comamonadaceae | *Pseudorhodoferax* | 0.000 | 0.000 | 0.000 | 0.018 | 0.000 | 0.000 |
| Proteobacteria | Betaproteobacteria | Burkholderiales | Comamonadaceae | *Ramlibacter* | 0.000 | 0.000 | 0.000 | 0.029 | 0.000 | 0.000 |
| Proteobacteria | Betaproteobacteria | Burkholderiales | Comamonadaceae | *Rhodoferax* | 0.000 | 0.000 | 0.000 | 0.018 | 0.000 | 0.000 |
| Proteobacteria | Betaproteobacteria | Burkholderiales | Comamonadaceae | *Simplicispira* | 0.000 | 0.000 | 0.000 | 0.004 | 0.000 | 0.000 |
| Proteobacteria | Betaproteobacteria | Burkholderiales | Comamonadaceae | *Simplicispira* 2 | 0.004 | 0.032 | 0.000 | 0.007 | 0.000 | 0.001 |
| Proteobacteria | Betaproteobacteria | Burkholderiales | Comamonadaceae | *Sphaerotilus* | 0.000 | 0.000 | 0.000 | 0.004 | 0.000 | 0.000 |
| Proteobacteria | Betaproteobacteria | Burkholderiales | Comamonadaceae | Unclassified | 0.000 | 0.000 | 0.013 | 0.025 | 0.004 | 0.000 |
| Proteobacteria | Betaproteobacteria | Burkholderiales | Comamonadaceae | Uncultured 11 | 0.000 | 0.000 | 0.000 | 0.011 | 0.000 | 0.000 |
| Proteobacteria | Betaproteobacteria | Burkholderiales | Comamonadaceae | Uncultured 12 | 0.000 | 0.000 | 0.000 | 0.007 | 0.000 | 0.000 |

**Table S2 continued.**

| | | | | | | | | | | |
|---|---|---|---|---|---|---|---|---|---|---|
| Proteobacteria | Betaproteobacteria | Burkholderiales | Comamonadaceae | Uncultured 13 | 0.000 | 0.000 | 0.000 | 0.004 | 0.000 | 0.000 |
| Proteobacteria | Betaproteobacteria | Burkholderiales | Comamonadaceae | Uncultured 14 | 0.002 | 0.000 | 0.000 | 0.000 | 0.000 | 0.000 |
| Proteobacteria | Betaproteobacteria | Burkholderiales | Comamonadaceae | Uncultured 15 | 0.000 | 0.000 | 0.000 | 0.007 | 0.000 | 0.000 |
| Proteobacteria | Betaproteobacteria | Burkholderiales | Comamonadaceae | Uncultured 17 | 0.000 | 0.000 | 0.000 | 0.007 | 0.000 | 0.000 |
| Proteobacteria | Betaproteobacteria | Burkholderiales | Comamonadaceae | Uncultured 18 | 0.000 | 0.000 | 0.000 | 0.004 | 0.004 | 0.000 |
| Proteobacteria | Betaproteobacteria | Burkholderiales | Comamonadaceae | Uncultured 2 | 0.000 | 0.000 | 0.000 | 0.004 | 0.000 | 0.000 |
| Proteobacteria | Betaproteobacteria | Burkholderiales | Comamonadaceae | Uncultured 20 | 0.000 | 0.000 | 0.000 | 0.004 | 0.000 | 0.000 |
| Proteobacteria | Betaproteobacteria | Burkholderiales | Comamonadaceae | Uncultured 21 | 0.000 | 0.000 | 0.000 | 0.011 | 0.000 | 0.000 |
| Proteobacteria | Betaproteobacteria | Burkholderiales | Comamonadaceae | Uncultured 22 | 0.000 | 0.000 | 0.000 | 0.004 | 0.000 | 0.000 |
| Proteobacteria | Betaproteobacteria | Burkholderiales | Comamonadaceae | Uncultured 24 | 0.000 | 0.000 | 0.000 | 0.004 | 0.000 | 0.000 |
| Proteobacteria | Betaproteobacteria | Burkholderiales | Comamonadaceae | Uncultured 25 | 0.011 | 0.000 | 0.000 | 0.000 | 0.004 | 0.000 |
| Proteobacteria | Betaproteobacteria | Burkholderiales | Comamonadaceae | Uncultured 26 | 0.001 | 0.000 | 0.000 | 0.000 | 0.000 | 0.000 |
| Proteobacteria | Betaproteobacteria | Burkholderiales | Comamonadaceae | Uncultured 4 | 0.001 | 0.000 | 0.000 | 0.000 | 0.004 | 0.000 |
| Proteobacteria | Betaproteobacteria | Burkholderiales | Comamonadaceae | *Variovorax* | 0.000 | 0.000 | 0.000 | 0.007 | 0.000 | 0.000 |
| Proteobacteria | Betaproteobacteria | Burkholderiales | Comamonadaceae | *Variovorax* 1 | 0.002 | 0.000 | 0.000 | 0.060 | 0.000 | 0.000 |
| Proteobacteria | Betaproteobacteria | Burkholderiales | Comamonadaceae | *Variovorax* 2 | 0.000 | 0.000 | 0.000 | 0.018 | 0.000 | 0.001 |
| Proteobacteria | Betaproteobacteria | Burkholderiales | Comamonadaceae | *Variovorax* a | 0.000 | 0.000 | 0.000 | 0.004 | 0.004 | 0.000 |
| Proteobacteria | Betaproteobacteria | Burkholderiales | Comamonadaceae | *Variovorax* b | 0.000 | 0.000 | 0.000 | 0.004 | 0.000 | 0.000 |
| Proteobacteria | Betaproteobacteria | Burkholderiales | Comamonadaceae | *Verminephrobacter* | 0.000 | 0.000 | 0.000 | 0.004 | 0.000 | 0.000 |
| Proteobacteria | Betaproteobacteria | Burkholderiales | Comamonadaceae | *Xenophilus* | 0.000 | 0.000 | 0.000 | 0.014 | 0.000 | 0.000 |
| Proteobacteria | Betaproteobacteria | Burkholderiales | Oxalobacteraceae | *Collimonas* | 0.023 | 0.032 | 0.000 | 0.018 | 0.000 | 0.003 |
| Proteobacteria | Betaproteobacteria | Burkholderiales | Oxalobacteraceae | *Herbaspirillum* | 0.001 | 0.000 | 0.000 | 0.018 | 0.000 | 0.000 |
| Proteobacteria | Betaproteobacteria | Burkholderiales | Oxalobacteraceae | *Herbaspirillum* 1 | 0.007 | 0.000 | 0.000 | 0.004 | 0.000 | 0.000 |
| Proteobacteria | Betaproteobacteria | Burkholderiales | Oxalobacteraceae | *Herbaspirillum* 2 | 0.001 | 0.032 | 0.000 | 0.000 | 0.000 | 0.000 |
| Proteobacteria | Betaproteobacteria | Burkholderiales | Oxalobacteraceae | *Herbaspirillum* 3 | 0.001 | 0.000 | 0.000 | 0.000 | 0.004 | 0.001 |
| Proteobacteria | Betaproteobacteria | Burkholderiales | Oxalobacteraceae | *Herminiimonas* 1 | 0.002 | 0.000 | 0.000 | 0.004 | 0.000 | 0.001 |
| Proteobacteria | Betaproteobacteria | Burkholderiales | Oxalobacteraceae | *Janthinobacterium* | 0.000 | 0.000 | 0.000 | 0.004 | 0.000 | 0.000 |

**Table S2 continued.**

| | | | | | | | | | | |
|---|---|---|---|---|---|---|---|---|---|---|
| Proteobacteria | Betaproteobacteria | Burkholderiales | Oxalobacteraceae | *Janthinobacterium* 1 | 0.000 | 0.000 | 0.000 | 0.004 | 0.004 | 0.000 |
| Proteobacteria | Betaproteobacteria | Burkholderiales | Oxalobacteraceae | *Janthinobacterium* 2 | 0.001 | 0.000 | 0.000 | 0.004 | 0.000 | 0.000 |
| Proteobacteria | Betaproteobacteria | Burkholderiales | Oxalobacteraceae | *Massilia* | 0.000 | 0.000 | 0.000 | 0.021 | 0.004 | 0.002 |
| Proteobacteria | Betaproteobacteria | Burkholderiales | Oxalobacteraceae | *Massilia* 1 | 0.000 | 0.000 | 0.000 | 0.004 | 0.000 | 0.000 |
| Proteobacteria | Betaproteobacteria | Burkholderiales | Oxalobacteraceae | *Massilia* 10 | 0.001 | 0.000 | 0.000 | 0.011 | 0.000 | 0.000 |
| Proteobacteria | Betaproteobacteria | Burkholderiales | Oxalobacteraceae | *Massilia* 9 | 0.000 | 0.000 | 0.000 | 0.025 | 0.000 | 0.000 |
| Proteobacteria | Betaproteobacteria | Burkholderiales | Oxalobacteraceae | *Massilia* sp a | 0.000 | 0.000 | 0.000 | 0.004 | 0.000 | 0.000 |
| Proteobacteria | Betaproteobacteria | Burkholderiales | Oxalobacteraceae | *Naxibacter* | 0.009 | 0.000 | 0.026 | 0.196 | 0.008 | 0.002 |
| Proteobacteria | Betaproteobacteria | Burkholderiales | Oxalobacteraceae | *Noviherbaspirillum* | 0.000 | 0.000 | 0.000 | 0.007 | 0.000 | 0.001 |
| Proteobacteria | Betaproteobacteria | Burkholderiales | Oxalobacteraceae | *Oxalicibacterium* | 0.000 | 0.032 | 0.000 | 0.000 | 0.000 | 0.000 |
| Proteobacteria | Betaproteobacteria | Burkholderiales | Oxalobacteraceae | *Oxalobacter* | 0.061 | 0.191 | 0.000 | 0.018 | 0.041 | 0.020 |
| Proteobacteria | Betaproteobacteria | Burkholderiales | Oxalobacteraceae | *Paucimonas* | 0.001 | 0.032 | 0.000 | 0.011 | 0.000 | 0.000 |
| Proteobacteria | Betaproteobacteria | Burkholderiales | Oxalobacteraceae | Unclassified | 0.000 | 0.032 | 0.000 | 0.011 | 0.000 | 0.001 |
| Proteobacteria | Betaproteobacteria | Burkholderiales | Oxalobacteraceae | Uncultured 10 | 0.001 | 0.000 | 0.000 | 0.000 | 0.000 | 0.000 |
| Proteobacteria | Betaproteobacteria | Burkholderiales | Oxalobacteraceae | Uncultured 9 | 0.001 | 0.032 | 0.000 | 0.000 | 0.000 | 0.001 |
| Proteobacteria | Betaproteobacteria | Burkholderiales | Oxalobacteraceae | Uncultured a | 0.000 | 0.000 | 0.000 | 0.007 | 0.000 | 0.000 |
| Proteobacteria | Betaproteobacteria | Burkholderiales | Oxalobacteraceae | Undibacterium | 0.130 | 0.159 | 0.000 | 0.029 | 0.045 | 0.017 |
| Proteobacteria | Betaproteobacteria | Burkholderiales | Oxalobacteraceae | Unclassified | 0.005 | 0.000 | 0.000 | 0.000 | 0.000 | 0.000 |
| Proteobacteria | Betaproteobacteria | Burkholderiales | Unclassified | *Aquabacterium* | 0.007 | 0.000 | 0.026 | 0.007 | 0.004 | 0.005 |
| Proteobacteria | Betaproteobacteria | Burkholderiales | Unclassified | *Aquincola* | 0.000 | 0.000 | 0.000 | 0.004 | 0.000 | 0.000 |
| Proteobacteria | Betaproteobacteria | Burkholderiales | Unclassified | *Inhella* | 0.000 | 0.000 | 0.000 | 0.014 | 0.014 | 0.000 |
| Proteobacteria | Betaproteobacteria | Burkholderiales | Unclassified | *Leptothrix* | 0.000 | 0.000 | 0.000 | 0.007 | 0.000 | 0.000 |
| Proteobacteria | Betaproteobacteria | Burkholderiales | Unclassified | *Roseateles* | 0.000 | 0.000 | 0.000 | 0.007 | 0.000 | 0.000 |
| Proteobacteria | Betaproteobacteria | Burkholderiales | Unclassified | *Thiobacter* | 0.000 | 0.000 | 0.000 | 0.000 | 0.016 | 0.000 |
| Proteobacteria | Betaproteobacteria | Burkholderiales | Unclassified | Unclassified | 0.000 | 0.000 | 0.000 | 0.004 | 0.000 | 0.000 |
| Proteobacteria | Betaproteobacteria | Hydrogenophilales | Hydrogenophilaceae | *Thiobacillus* 1 | 0.005 | 0.000 | 0.000 | 0.004 | 0.012 | 0.001 |
| Proteobacteria | Betaproteobacteria | Hydrogenophilales | Hydrogenophilaceae | Unclassified | 0.001 | 0.000 | 0.000 | 0.000 | 0.000 | 0.000 |

**Table S2 continued.**

| | | | | | | | | | | |
|---|---|---|---|---|---|---|---|---|---|---|
| Proteobacteria | Betaproteobacteria | Methylophilales | Methylophilaceae | Unclassified | 0.000 | 0.000 | 0.000 | 0.200 | 0.000 | 0.000 |
| Proteobacteria | Betaproteobacteria | Methylophilales | Methylophilaceae | Uncultured | 0.001 | 0.000 | 0.000 | 0.228 | 0.000 | 0.001 |
| Proteobacteria | Betaproteobacteria | Methylophilales | Methylophilaceae | Unclassified | 0.001 | 0.000 | 0.000 | 0.000 | 0.000 | 0.000 |
| Proteobacteria | Betaproteobacteria | Neisseriales | Chromobacteriaceae | *Chitinilyticum* | 0.001 | 0.000 | 0.000 | 0.000 | 0.000 | 0.000 |
| Proteobacteria | Betaproteobacteria | Neisseriales | Chromobacteriaceae | *Chromobacterium* | 0.000 | 0.000 | 0.000 | 0.004 | 0.000 | 0.000 |
| Proteobacteria | Betaproteobacteria | Neisseriales | Neisseriaceae 1 | *Neisseria* 1 | 0.000 | 0.000 | 0.000 | 0.004 | 0.000 | 0.000 |
| Proteobacteria | Betaproteobacteria | Neisseriales | Neisseriaceae 1 | Uncultured 1 | 0.000 | 0.000 | 0.000 | 0.007 | 0.000 | 0.002 |
| Proteobacteria | Betaproteobacteria | Neisseriales | Neisseriaceae 1 | Uncultured 4 | 0.000 | 0.000 | 0.000 | 0.011 | 0.000 | 0.000 |
| Proteobacteria | Betaproteobacteria | Neisseriales | Neisseriaceae 1 | *Vogesella* | 0.000 | 0.000 | 0.000 | 0.004 | 0.000 | 0.000 |
| Proteobacteria | Betaproteobacteria | Nitrosomonadales | Gallionellaceae | *Candidatus* Nitrotoga | 0.000 | 0.000 | 0.000 | 0.004 | 0.000 | 0.000 |
| Proteobacteria | Betaproteobacteria | Nitrosomonadales | Gallionellaceae | Unclassified | 0.000 | 0.000 | 0.000 | 0.004 | 0.004 | 0.000 |
| Proteobacteria | Betaproteobacteria | Nitrosomonadales | Gallionellaceae | Uncultured 1 | 0.000 | 0.000 | 0.000 | 0.011 | 0.000 | 0.000 |
| Proteobacteria | Betaproteobacteria | Nitrosomonadales | Gallionellaceae | Uncultured 2 | 0.005 | 0.000 | 0.000 | 0.004 | 0.016 | 0.001 |
| Proteobacteria | Betaproteobacteria | Nitrosomonadales | Gallionellaceae | Uncultured 3 | 0.001 | 0.000 | 0.000 | 0.000 | 0.000 | 0.000 |
| Proteobacteria | Betaproteobacteria | Nitrosomonadales | Gallionellaceae | Unclassified | 0.005 | 0.000 | 0.000 | 0.000 | 0.000 | 0.000 |
| Proteobacteria | Betaproteobacteria | Nitrosomonadales | Methylophilaceae | *Methylophilus* | 0.004 | 0.000 | 0.000 | 0.150 | 0.000 | 0.000 |
| Proteobacteria | Betaproteobacteria | Nitrosomonadales | Methylophilaceae | *Methylotenera* | 0.000 | 0.000 | 0.000 | 0.007 | 0.000 | 0.000 |
| Proteobacteria | Betaproteobacteria | Nitrosomonadales | Methylophilaceae | *Methylovorus* | 0.000 | 0.000 | 0.000 | 0.007 | 0.000 | 0.000 |
| Proteobacteria | Betaproteobacteria | Nitrosomonadales | Nitrosomonadaceae | *Nitrosomonas* | 0.006 | 0.000 | 0.000 | 0.004 | 0.004 | 0.007 |
| Proteobacteria | Betaproteobacteria | Nitrosomonadales | Nitrosomonadaceae | *Nitrosospira* | 0.000 | 0.000 | 0.000 | 0.004 | 0.000 | 0.000 |
| Proteobacteria | Betaproteobacteria | Nitrosomonadales | Nitrosomonadaceae | Unclassified | 0.000 | 0.000 | 0.000 | 0.004 | 0.000 | 0.000 |
| Proteobacteria | Betaproteobacteria | Nitrosomonadales | Nitrosomonadaceae | Uncultured 1 | 0.006 | 0.000 | 0.000 | 0.214 | 0.012 | 0.001 |
| Proteobacteria | Betaproteobacteria | Nitrosomonadales | Nitrosomonadaceae | Uncultured 2 | 0.001 | 0.000 | 0.000 | 0.000 | 0.004 | 0.000 |
| Proteobacteria | Betaproteobacteria | Nitrosomonadales | Sterolibacteriaceae | *Methyloversatilis* | 0.006 | 0.000 | 0.000 | 0.000 | 0.020 | 0.000 |
| Proteobacteria | Betaproteobacteria | Rhodcyclales | Zoogloeaceae | *Thauera* | 0.017 | 0.000 | 0.000 | 0.000 | 0.037 | 0.003 |
| Proteobacteria | Betaproteobacteria | Rhodocyclales | Azonexaceae | *Azonexus* | 0.000 | 0.000 | 0.000 | 0.000 | 0.004 | 0.000 |
| Proteobacteria | Betaproteobacteria | Rhodocyclales | Rhodocyclaceae | *Azospira* | 0.000 | 0.000 | 0.000 | 0.004 | 0.004 | 0.000 |

**Table S2 continued.**

| | | | | | | | | | | |
|---|---|---|---|---|---|---|---|---|---|---|
| Proteobacteria | Betaproteobacteria | Rhodocyclales | Rhodocyclaceae | *Dechloromonas* | 0.000 | 0.000 | 0.000 | 0.004 | 0.000 | 0.000 |
| Proteobacteria | Betaproteobacteria | Rhodocyclales | Rhodocyclaceae | *Propionivibrio* | 0.236 | 0.128 | 0.000 | 0.036 | 0.033 | 0.047 |
| Proteobacteria | Betaproteobacteria | Rhodocyclales | Rhodocyclaceae | *Rhodocyclus* | 0.001 | 0.000 | 0.000 | 0.000 | 0.000 | 0.000 |
| Proteobacteria | Betaproteobacteria | Rhodocyclales | Rhodocyclaceae 1 | Uncultured a | 0.000 | 0.000 | 0.000 | 0.004 | 0.000 | 0.000 |
| Proteobacteria | Betaproteobacteria | Rhodocyclales | Rhodocyclaceae 2 | *Azoarcus* 2 | 0.005 | 0.000 | 0.000 | 0.004 | 0.008 | 0.001 |
| Proteobacteria | Betaproteobacteria | Rhodocyclales | Rhodocyclaceae 2 | Insect cluster | 0.156 | 0.000 | 0.000 | 0.031 | 0.464 | 0.000 |
| Proteobacteria | Betaproteobacteria | Rhodocyclales | Rhodocyclaceae 2 | Insect cluster | 0.000 | 0.000 | 0.000 | 0.000 | 0.000 | 0.018 |
| Proteobacteria | Betaproteobacteria | Rhodocyclales | Rhodocyclaceae 2 | Unclassified | 0.000 | 0.000 | 0.000 | 0.004 | 0.008 | 0.000 |
| Proteobacteria | Betaproteobacteria | Rhodocyclales | Rhodocyclaceae 2 | Unclassified | 0.002 | 0.000 | 0.000 | 0.000 | 0.000 | 0.000 |
| Proteobacteria | Betaproteobacteria | Rhodocyclales | Rhodocyclaceae 3 | *Azonexus* 2 | 0.000 | 0.000 | 0.000 | 0.000 | 0.008 | 0.000 |
| Proteobacteria | Betaproteobacteria | Rhodocyclales | Rhodocyclaceae 3 | *Dechloromonas* 1 | 0.011 | 0.000 | 0.000 | 0.000 | 0.037 | 0.001 |
| Proteobacteria | Betaproteobacteria | Rhodocyclales | Rhodocyclaceae 3 | Termite cluster | 0.021 | 0.000 | 0.000 | 0.004 | 0.081 | 0.006 |
| Proteobacteria | Betaproteobacteria | Rhodocyclales | Rhodocyclaceae 3 | Unclassified | 0.000 | 0.000 | 0.000 | 0.000 | 0.004 | 0.000 |
| Proteobacteria | Betaproteobacteria | Rhodocyclales | Rhodocyclaceae 3 | Uncultured 1 | 0.000 | 0.000 | 0.000 | 0.000 | 0.000 | 0.001 |
| Proteobacteria | Betaproteobacteria | Rhodocyclales | Rhodocyclaceae 3 | Uncultured 2 | 0.001 | 0.000 | 0.000 | 0.000 | 0.000 | 0.001 |
| Proteobacteria | Betaproteobacteria | Rhodocyclales | Rhodocyclaceae 3 | Uncultured 5 | 0.002 | 0.000 | 0.000 | 0.000 | 0.008 | 0.001 |
| Proteobacteria | Betaproteobacteria | Rhodocyclales | Zoogloeaceae | *Azoarcus* | 0.004 | 0.000 | 0.000 | 0.007 | 0.012 | 0.000 |
| Proteobacteria | Betaproteobacteria | Rhodocyclales | Zoogloeaceae | *Uliginosibacterium* | 0.002 | 0.000 | 0.000 | 0.000 | 0.000 | 0.000 |
| Proteobacteria | Betaproteobacteria | Rhodocyclales | Zoogloeaceae | *Zoogloea* | 0.014 | 0.000 | 0.000 | 0.007 | 0.049 | 0.000 |
| Proteobacteria | Betaproteobacteria | SC-I-84 | Insect cluster II | Unclassified | 0.000 | 0.000 | 0.000 | 0.004 | 0.000 | 0.002 |
| Proteobacteria | Betaproteobacteria | SC-I-84 | Unclassified | Unclassified | 0.000 | 0.032 | 0.000 | 0.025 | 0.000 | 0.000 |
| Proteobacteria | Betaproteobacteria | TRA3-20 | Unclassified | Unclassified | 0.000 | 0.000 | 0.000 | 0.071 | 0.000 | 0.000 |
| Proteobacteria | Betaproteobacteria | Unclassified | Unclassified | *Paucibacter* | 0.000 | 0.000 | 0.000 | 0.000 | 0.000 | 0.001 |
| Proteobacteria | Betaproteobacteria | Unclassified | Unclassified | Unclassified | 0.000 | 0.000 | 0.000 | 0.000 | 0.004 | 0.000 |
| Proteobacteria | Betaproteobacteria | Unclassified | Unclassified | Unclassified | 0.001 | 0.000 | 0.000 | 0.000 | 0.000 | 0.000 |
| Proteobacteria | Deltaproteobacteria | Bdellovibrionales | Bacteriovoracaceae | *Bacteriovorax* 1 | 0.017 | 0.000 | 0.000 | 0.000 | 0.000 | 0.000 |
| Proteobacteria | Deltaproteobacteria | Bdellovibrionales | Bacteriovoracaceae | *Bacteriovorax* 2 | 0.001 | 0.000 | 0.000 | 0.011 | 0.000 | 0.000 |

**Table S2 continued.**

| | | | | | | | | | | |
|---|---|---|---|---|---|---|---|---|---|---|
| Proteobacteria | Deltaproteobacteria | Bdellovibrionales | Bacteriovoraceae | Uncultured 1 | 0.001 | 0.000 | 0.000 | 0.000 | 0.000 | 0.000 |
| Proteobacteria | Deltaproteobacteria | Bdellovibrionales | Bacteriovoraceae | Unclassified | 0.030 | 0.000 | 0.000 | 0.000 | 0.000 | 0.000 |
| Proteobacteria | Deltaproteobacteria | Bdellovibrionales | Bdellovibrionaceae | *Bdellovibrio* | 0.000 | 0.000 | 0.013 | 0.021 | 0.000 | 0.000 |
| Proteobacteria | Deltaproteobacteria | Bdellovibrionales | Bdellovibrionaceae | OM27 clade | 0.000 | 0.000 | 0.000 | 0.007 | 0.000 | 0.000 |
| Proteobacteria | Deltaproteobacteria | Desulfarculales | Desulfarculaceae | *Desulfarculus* | 0.049 | 0.064 | 0.000 | 0.021 | 0.053 | 0.015 |
| Proteobacteria | Deltaproteobacteria | Desulfarculales | Desulfarculaceae | *Dethiosulfatarculus* | 0.001 | 0.000 | 0.000 | 0.000 | 0.000 | 0.001 |
| Proteobacteria | Deltaproteobacteria | Desulfarculales | Desulfarculaceae | Uncultured | 0.010 | 0.033 | 0.000 | 0.014 | 0.000 | 0.010 |
| Proteobacteria | Deltaproteobacteria | Desulfarculales | Desulfarculaceae | Unclassified | 0.004 | 0.000 | 0.000 | 0.000 | 0.000 | 0.000 |
| Proteobacteria | Deltaproteobacteria | Desulfobacterales | Desulfobacteraceae | *Algorimarina* | 0.000 | 0.000 | 0.000 | 0.004 | 0.000 | 0.000 |
| Proteobacteria | Deltaproteobacteria | Desulfobacterales | Desulfobacteraceae | *Desulfatiferula* | 0.001 | 0.000 | 0.000 | 0.004 | 0.004 | 0.000 |
| Proteobacteria | Deltaproteobacteria | Desulfobacterales | Desulfobacteraceae | *Desulfatitalea* | 0.001 | 0.000 | 0.000 | 0.000 | 0.000 | 0.000 |
| Proteobacteria | Deltaproteobacteria | Desulfobacterales | Desulfobacteraceae | *Desulfonema* | 0.001 | 0.000 | 0.000 | 0.000 | 0.000 | 0.000 |
| Proteobacteria | Deltaproteobacteria | Desulfobacterales | Desulfobacteraceae | *Desulfosarcina* | 0.001 | 0.000 | 0.000 | 0.000 | 0.000 | 0.000 |
| Proteobacteria | Deltaproteobacteria | Desulfobacterales | Desulfobacteraceae | endosymbionts | 0.001 | 0.000 | 0.000 | 0.000 | 0.004 | 0.001 |
| Proteobacteria | Deltaproteobacteria | Desulfobacterales | Desulfobacteraceae | Unclassified | 0.000 | 0.032 | 0.000 | 0.000 | 0.000 | 0.000 |
| Proteobacteria | Deltaproteobacteria | Desulfobacterales | Desulfobacteraceae | Unclassified | 0.001 | 0.000 | 0.000 | 0.000 | 0.000 | 0.000 |
| Proteobacteria | Deltaproteobacteria | Desulfobacterales | Desulfobulbaceae | *Desulfobacterium* | 0.000 | 0.000 | 0.000 | 0.000 | 0.000 | 0.002 |
| Proteobacteria | Deltaproteobacteria | Desulfobacterales | Desulfobulbaceae | *Desulfobulbus* | 0.000 | 0.032 | 0.000 | 0.004 | 0.000 | 0.001 |
| Proteobacteria | Deltaproteobacteria | Desulfobacterales | Desulfobulbaceae | *Desulfocapsa* 1 | 0.001 | 0.000 | 0.000 | 0.004 | 0.000 | 0.000 |
| Proteobacteria | Deltaproteobacteria | Desulfobacterales | Desulfobulbaceae | *Desulfofustis* | 0.001 | 0.000 | 0.000 | 0.000 | 0.004 | 0.000 |
| Proteobacteria | Deltaproteobacteria | Desulfobacterales | Desulfobulbaceae | Unclassified | 0.000 | 0.000 | 0.000 | 0.000 | 0.004 | 0.000 |
| Proteobacteria | Deltaproteobacteria | Desulfobacterales | Nitrospinaceae | *Nitrospina* | 0.002 | 0.000 | 0.000 | 0.000 | 0.004 | 0.000 |
| Proteobacteria | Deltaproteobacteria | Desulfovibrionales | Desulfonatronaceae | *Desulfonatronum* | 0.000 | 0.000 | 0.000 | 0.000 | 0.004 | 0.002 |
| Proteobacteria | Deltaproteobacteria | Desulfovibrionales | Desulfovibrionaceae | *Bilophila* | 0.001 | 0.000 | 0.000 | 0.000 | 0.012 | 0.003 |
| Proteobacteria | Deltaproteobacteria | Desulfovibrionales | Desulfovibrionaceae | *Desulfocurvus* | 0.001 | 0.000 | 0.000 | 0.000 | 0.000 | 0.001 |
| Proteobacteria | Deltaproteobacteria | Desulfovibrionales | Desulfovibrionaceae | *Desulfohalophilus* | 0.001 | 0.000 | 0.000 | 0.000 | 0.004 | 0.000 |
| Proteobacteria | Deltaproteobacteria | Desulfovibrionales | Desulfovibrionaceae | *Desulfomicrobium* | 0.001 | 0.000 | 0.000 | 0.004 | 0.033 | 0.013 |

**Table S2 continued.**

| | | | | | | | | | | |
|---|---|---|---|---|---|---|---|---|---|---|
| Proteobacteria | Deltaproteobacteria | Desulfovibrionales | Desulfovibrionaceae | *Desulfovibrio* | 0.084 | 0.064 | 0.000 | 0.011 | 1.273 | 0.737 |
| Proteobacteria | Deltaproteobacteria | Desulfovibrionales | Desulfovibrionaceae | *Desulfovibrio* 2 | 0.019 | 0.000 | 0.000 | 0.000 | 0.020 | 0.016 |
| Proteobacteria | Deltaproteobacteria | Desulfovibrionales | Desulfovibrionaceae | *Desulfovibrio* 4 | 0.001 | 0.000 | 0.000 | 0.000 | 0.004 | 0.000 |
| Proteobacteria | Deltaproteobacteria | Desulfovibrionales | Desulfovibrionaceae | *Desulfovibrio* 5 | 0.001 | 0.000 | 0.000 | 0.000 | 0.000 | 0.005 |
| Proteobacteria | Deltaproteobacteria | Desulfovibrionales | Desulfovibrionaceae | Gut cluster 1 | 0.058 | 0.032 | 0.000 | 0.025 | 3.050 | 1.536 |
| Proteobacteria | Deltaproteobacteria | Desulfovibrionales | Desulfovibrionaceae | Gut cluster 2 | 0.727 | 0.702 | 0.000 | 0.289 | 0.704 | 0.660 |
| Proteobacteria | Deltaproteobacteria | Desulfovibrionales | Desulfovibrionaceae | Gut cluster 3 | 1.644 | 3.126 | 0.013 | 0.673 | 4.681 | 2.782 |
| Proteobacteria | Deltaproteobacteria | Desulfovibrionales | Desulfovibrionaceae | *Lawsonia* | 0.005 | 0.032 | 0.000 | 0.004 | 0.024 | 0.003 |
| Proteobacteria | Deltaproteobacteria | Desulfovibrionales | Desulfovibrionaceae | *Mailhella* | 0.000 | 0.000 | 0.000 | 0.000 | 0.004 | 0.000 |
| Proteobacteria | Deltaproteobacteria | Desulfovibrionales | Desulfovibrionaceae | *Pseudodesulfovibrio* | 0.001 | 0.000 | 0.000 | 0.000 | 0.000 | 0.000 |
| Proteobacteria | Deltaproteobacteria | Desulfovibrionales | Desulfovibrionaceae | Termite cluster II | 1.123 | 0.734 | 0.000 | 0.353 | 1.257 | 2.079 |
| Proteobacteria | Deltaproteobacteria | Desulfovibrionales | Desulfovibrionaceae | Termite cluster III | 0.087 | 0.128 | 0.000 | 0.021 | 0.065 | 0.017 |
| Proteobacteria | Deltaproteobacteria | Desulfurellales | Desulfurellaceae | Uncultured | 0.000 | 0.000 | 0.000 | 0.007 | 0.000 | 0.000 |
| Proteobacteria | Deltaproteobacteria | Desulfuromonadales | BVA18 | Unclassified | 0.000 | 0.000 | 0.000 | 0.004 | 0.000 | 0.000 |
| Proteobacteria | Deltaproteobacteria | Desulfuromonadales | Desulfuromonadaceae | *Desulfuromonas* | 0.000 | 0.000 | 0.000 | 0.004 | 0.000 | 0.000 |
| Proteobacteria | Deltaproteobacteria | Desulfuromonadales | Desulfuromonadaceae | *Desulfuromonas* 3 | 0.002 | 0.032 | 0.000 | 0.000 | 0.000 | 0.000 |
| Proteobacteria | Deltaproteobacteria | Desulfuromonadales | Desulfuromonadaceae | *Desulfuromusa* | 0.001 | 0.000 | 0.000 | 0.000 | 0.000 | 0.000 |
| Proteobacteria | Deltaproteobacteria | Desulfuromonadales | Desulfuromonadaceae | *Pelobacter* | 0.001 | 0.000 | 0.000 | 0.000 | 0.000 | 0.000 |
| Proteobacteria | Deltaproteobacteria | Desulfuromonadales | Geobacteraceae | *Geobacter* | 0.001 | 0.000 | 0.000 | 0.011 | 0.004 | 0.000 |
| Proteobacteria | Deltaproteobacteria | Desulfuromonadales | Geobacteraceae | *Geothermobacter* | 0.001 | 0.000 | 0.000 | 0.000 | 0.000 | 0.000 |
| Proteobacteria | Deltaproteobacteria | Desulfuromonadales | GR-WP33-58 | Unclassified | 0.000 | 0.000 | 0.000 | 0.007 | 0.008 | 0.000 |
| Proteobacteria | Deltaproteobacteria | Desulfuromonadales | GR-WP33-58 | Unclassified | 0.002 | 0.000 | 0.000 | 0.000 | 0.000 | 0.000 |
| Proteobacteria | Deltaproteobacteria | Desulfuromonadales | Unclassified | Unclassified | 0.001 | 0.000 | 0.000 | 0.000 | 0.000 | 0.000 |
| Proteobacteria | Deltaproteobacteria | F-1404R | Unclassified | Unclassified | 0.001 | 0.000 | 0.000 | 0.000 | 0.000 | 0.000 |
| Proteobacteria | Deltaproteobacteria | GR-WP33-30 | Unclassified | Unclassified | 0.000 | 0.000 | 0.000 | 0.014 | 0.000 | 0.000 |
| Proteobacteria | Deltaproteobacteria | GR-WP33-30 | Uncultured 1 | Unclassified | 0.000 | 0.000 | 0.000 | 0.139 | 0.000 | 0.000 |
| Proteobacteria | Deltaproteobacteria | GR-WP33-30 | Uncultured 1 | Unclassified | 0.001 | 0.000 | 0.000 | 0.000 | 0.000 | 0.000 |

**Table S2 continued.**

| | | | | | | | | | | |
|---|---|---|---|---|---|---|---|---|---|---|
| Proteobacteria | Deltaproteobacteria | GR-WP33-30 | Uncultured 2 | Unclassified | 0.000 | 0.000 | 0.000 | 0.021 | 0.000 | 0.001 |
| Proteobacteria | Deltaproteobacteria | GR-WP33-30 | Unclassified | Unclassified | 0.001 | 0.000 | 0.000 | 0.000 | 0.000 | 0.000 |
| Proteobacteria | Deltaproteobacteria | Myxococcales | Archangiaceae | *Angiococcus* | 0.000 | 0.000 | 0.000 | 0.007 | 0.000 | 0.000 |
| Proteobacteria | Deltaproteobacteria | Myxococcales | Archangiaceae | *Cystobacter* | 0.001 | 0.000 | 0.000 | 0.011 | 0.000 | 0.000 |
| Proteobacteria | Deltaproteobacteria | Myxococcales | Archangiaceae | *Stigmatella* | 0.001 | 0.000 | 0.000 | 0.000 | 0.000 | 0.000 |
| Proteobacteria | Deltaproteobacteria | Myxococcales | Cluster 319-6G20 | Unclassified | 0.000 | 0.000 | 0.000 | 0.075 | 0.000 | 0.008 |
| Proteobacteria | Deltaproteobacteria | Myxococcales | Cluster 319-6G20 | Unclassified | 0.001 | 0.000 | 0.000 | 0.000 | 0.000 | 0.000 |
| Proteobacteria | Deltaproteobacteria | Myxococcales | Cystobacteraceae | *Anaeromyxobacter* | 0.001 | 0.000 | 0.000 | 0.029 | 0.000 | 0.000 |
| Proteobacteria | Deltaproteobacteria | Myxococcales | Cystobacteraceae | Unclassified | 0.000 | 0.000 | 0.000 | 0.025 | 0.000 | 0.000 |
| Proteobacteria | Deltaproteobacteria | Myxococcales | Haliangiaceae | *Haliangium* | 0.000 | 0.000 | 0.000 | 0.128 | 0.004 | 0.000 |
| Proteobacteria | Deltaproteobacteria | Myxococcales | JG37-AG-15 | Unclassified | 0.000 | 0.000 | 0.000 | 0.071 | 0.000 | 0.000 |
| Proteobacteria | Deltaproteobacteria | Myxococcales | JG37-AG-15 | Unclassified | 0.001 | 0.000 | 0.000 | 0.000 | 0.000 | 0.000 |
| Proteobacteria | Deltaproteobacteria | Myxococcales | Kofleriaceae | *Kofleria* | 0.000 | 0.000 | 0.000 | 0.014 | 0.000 | 0.000 |
| Proteobacteria | Deltaproteobacteria | Myxococcales | mle1-27 | Unclassified | 0.000 | 0.000 | 0.000 | 0.007 | 0.000 | 0.000 |
| Proteobacteria | Deltaproteobacteria | Myxococcales | mle1-27 | Unclassified | 0.001 | 0.000 | 0.000 | 0.000 | 0.000 | 0.000 |
| Proteobacteria | Deltaproteobacteria | Myxococcales | Myxococcaceae | *Aggregicoccus* | 0.000 | 0.000 | 0.000 | 0.004 | 0.000 | 0.000 |
| Proteobacteria | Deltaproteobacteria | Myxococcales | Myxococcaceae | *Corallococcus* | 0.000 | 0.000 | 0.000 | 0.011 | 0.000 | 0.000 |
| Proteobacteria | Deltaproteobacteria | Myxococcales | Myxococcaceae | *Myxococcus* | 0.000 | 0.000 | 0.000 | 0.004 | 0.000 | 0.000 |
| Proteobacteria | Deltaproteobacteria | Myxococcales | Myxococcaceae | Unclassified | 0.000 | 0.000 | 0.000 | 0.004 | 0.000 | 0.000 |
| Proteobacteria | Deltaproteobacteria | Myxococcales | Nannocystaceae | *Nannocystis* | 0.000 | 0.000 | 0.000 | 0.011 | 0.000 | 0.000 |
| Proteobacteria | Deltaproteobacteria | Myxococcales | Nannocystaceae | *Pseudenhygromyxa* | 0.000 | 0.000 | 0.000 | 0.004 | 0.000 | 0.000 |
| Proteobacteria | Deltaproteobacteria | Myxococcales | nocystaceae | Unclassified | 0.000 | 0.000 | 0.000 | 0.025 | 0.000 | 0.000 |
| Proteobacteria | Deltaproteobacteria | Myxococcales | Phaselicystidaceae | *Phaselicystis* | 0.000 | 0.000 | 0.000 | 0.014 | 0.004 | 0.000 |
| Proteobacteria | Deltaproteobacteria | Myxococcales | Polyangiaceae | *Aetherobacter* | 0.000 | 0.000 | 0.000 | 0.000 | 0.004 | 0.000 |
| Proteobacteria | Deltaproteobacteria | Myxococcales | Polyangiaceae | *Byssovorax* | 0.000 | 0.000 | 0.000 | 0.004 | 0.000 | 0.000 |
| Proteobacteria | Deltaproteobacteria | Myxococcales | Polyangiaceae | *Chondromyces* | 0.000 | 0.000 | 0.000 | 0.004 | 0.000 | 0.000 |
| Proteobacteria | Deltaproteobacteria | Myxococcales | Polyangiaceae | *Racemicystis* | 0.000 | 0.000 | 0.000 | 0.004 | 0.000 | 0.000 |

**Table S2 continued.**

| | | | | | | | | | | |
|---|---|---|---|---|---|---|---|---|---|---|
| Proteobacteria | Deltaproteobacteria | Myxococcales | Polyangiaceae | *Sorangium* | 0.000 | 0.000 | 0.000 | 0.032 | 0.000 | 0.000 |
| Proteobacteria | Deltaproteobacteria | Myxococcales | Polyangiaceae | Unclassified | 0.000 | 0.000 | 0.000 | 0.007 | 0.000 | 0.000 |
| Proteobacteria | Deltaproteobacteria | Myxococcales | Sandaracinaceae | *Sandaracinus* | 0.000 | 0.000 | 0.000 | 0.011 | 0.000 | 0.000 |
| Proteobacteria | Deltaproteobacteria | Myxococcales | Unclassified | *Enhygromyxa* | 0.001 | 0.000 | 0.000 | 0.000 | 0.000 | 0.000 |
| Proteobacteria | Deltaproteobacteria | Myxococcales | Unclassified | *Minicystis* | 0.000 | 0.000 | 0.000 | 0.007 | 0.000 | 0.000 |
| Proteobacteria | Deltaproteobacteria | Myxococcales | Unclassified | Unclassified | 0.000 | 0.000 | 0.000 | 0.032 | 0.000 | 0.000 |
| Proteobacteria | Deltaproteobacteria | Myxococcales | Unclassified | Unclassified | 0.000 | 0.000 | 0.000 | 0.000 | 0.000 | 0.002 |
| Proteobacteria | Deltaproteobacteria | Myxococcales | Uncultured 2 | Unclassified | 0.000 | 0.000 | 0.000 | 0.004 | 0.000 | 0.000 |
| Proteobacteria | Deltaproteobacteria | Myxococcales | Uncultured 3 | Unclassified | 0.000 | 0.000 | 0.000 | 0.039 | 0.000 | 0.001 |
| Proteobacteria | Deltaproteobacteria | Myxococcales | Uncultured 3 | Unclassified | 0.001 | 0.000 | 0.000 | 0.000 | 0.000 | 0.000 |
| Proteobacteria | Deltaproteobacteria | Myxococcales | Uncultured 4 | Unclassified | 0.000 | 0.000 | 0.000 | 0.029 | 0.000 | 0.000 |
| Proteobacteria | Deltaproteobacteria | Myxococcales | Uncultured 5 | Unclassified | 0.000 | 0.000 | 0.000 | 0.007 | 0.000 | 0.000 |
| Proteobacteria | Deltaproteobacteria | Myxococcales | Vulgatibacteraceae | *Vulgatibacter* | 0.000 | 0.000 | 0.000 | 0.007 | 0.000 | 0.000 |
| Proteobacteria | Deltaproteobacteria | Rs-K70 | Insect cluster | Unclassified | 0.000 | 0.096 | 0.000 | 0.043 | 0.049 | 0.045 |
| Proteobacteria | Deltaproteobacteria | Rs-K70 | Insect cluster | Unclassified | 0.114 | 0.000 | 0.000 | 0.000 | 0.000 | 0.000 |
| Proteobacteria | Deltaproteobacteria | Rs-K70 | Termite cluster I | Unclassified | 0.000 | 0.638 | 0.000 | 0.157 | 0.077 | 0.065 |
| Proteobacteria | Deltaproteobacteria | Rs-K70 | Termite cluster I | Unclassified | 0.590 | 0.000 | 0.000 | 0.000 | 0.000 | 0.000 |
| Proteobacteria | Deltaproteobacteria | Rs-K70 | Termite cluster III | Unclassified | 0.000 | 0.000 | 0.000 | 0.021 | 0.008 | 0.011 |
| Proteobacteria | Deltaproteobacteria | Rs-K70 | Termite cluster III | Unclassified | 0.069 | 0.000 | 0.000 | 0.000 | 0.000 | 0.000 |
| Proteobacteria | Deltaproteobacteria | Rs-K70 | Termite cluster IV | Unclassified | 0.000 | 0.096 | 0.000 | 0.068 | 0.069 | 0.059 |
| Proteobacteria | Deltaproteobacteria | Rs-K70 | Termite cluster IV | Unclassified | 0.117 | 0.000 | 0.000 | 0.000 | 0.000 | 0.000 |
| Proteobacteria | Deltaproteobacteria | Rs-K70 | Termite cluster V | Unclassified | 0.001 | 0.000 | 0.000 | 0.000 | 0.000 | 0.000 |
| Proteobacteria | Deltaproteobacteria | SAR324 clade(Marine group B) | Unclassified | Unclassified | 0.001 | 0.000 | 0.000 | 0.000 | 0.000 | 0.000 |
| Proteobacteria | Deltaproteobacteria | Sh765B-TzT-29 | Unclassified | Unclassified | 0.000 | 0.000 | 0.000 | 0.032 | 0.004 | 0.001 |
| Proteobacteria | Deltaproteobacteria | Syntrophobacterales | Syntrophaceae | *Syntrophus* | 0.035 | 0.000 | 0.000 | 0.000 | 0.016 | 0.003 |
| Proteobacteria | Deltaproteobacteria | Syntrophobacterales | Syntrophaceae | Unclassified | 0.001 | 0.000 | 0.000 | 0.000 | 0.000 | 0.000 |
| Proteobacteria | Deltaproteobacteria | Syntrophobacterales | Syntrophobacteraceae | *Desulfoglaeba* | 0.001 | 0.000 | 0.000 | 0.000 | 0.000 | 0.000 |

**Table S2 continued.**

| | | | | | | | | | | |
|---|---|---|---|---|---|---|---|---|---|---|
| Proteobacteria | Deltaproteobacteria | Syntrophobacterales | Syntrophobacteraceae | *Desulfosoma* | 0.000 | 0.000 | 0.000 | 0.004 | 0.000 | 0.000 |
| Proteobacteria | Deltaproteobacteria | Syntrophobacterales | Syntrophobacteraceae | Unclassified | 0.000 | 0.000 | 0.000 | 0.004 | 0.000 | 0.001 |
| Proteobacteria | Deltaproteobacteria | Unclassified | Unclassified | Unclassified | 0.001 | 0.000 | 0.000 | 0.000 | 0.000 | 0.000 |
| Proteobacteria | Epsilonproteobacteria | Campylobacterales | Campylobacteraceae | *Arcobacter* | 0.147 | 0.032 | 0.000 | 0.342 | 0.525 | 3.010 |
| Proteobacteria | Epsilonproteobacteria | Campylobacterales | Campylobacteraceae | *Campylobacter* | 0.004 | 0.000 | 0.000 | 0.000 | 0.016 | 0.002 |
| Proteobacteria | Epsilonproteobacteria | Campylobacterales | Campylobacteraceae | *Sulfurospirillum* | 2.197 | 1.053 | 0.000 | 0.449 | 1.793 | 0.356 |
| Proteobacteria | Epsilonproteobacteria | Campylobacterales | Helicobacteraceae | *Helicobacter* | 0.001 | 0.000 | 0.000 | 0.000 | 0.000 | 0.000 |
| Proteobacteria | Epsilonproteobacteria | Campylobacterales | Helicobacteraceae | Rs-M59 termite group | 0.004 | 0.000 | 0.000 | 0.000 | 0.008 | 0.002 |
| Proteobacteria | Epsilonproteobacteria | Campylobacterales | Helicobacteraceae | *Sulfuricurvum* | 0.004 | 0.000 | 0.000 | 0.000 | 0.000 | 0.000 |
| Proteobacteria | Epsilonproteobacteria | Campylobacterales | Helicobacteraceae | *Sulfurimonas* | 0.001 | 0.000 | 0.000 | 0.000 | 0.000 | 0.000 |
| Proteobacteria | Epsilonproteobacteria | Campylobacterales | Helicobacteraceae | *Sulfurovum* | 0.001 | 0.000 | 0.000 | 0.000 | 0.012 | 0.003 |
| Proteobacteria | Epsilonproteobacteria | Campylobacterales | Helicobacteraceae | Unclassified | 0.000 | 0.000 | 0.000 | 0.000 | 0.000 | 0.001 |
| Proteobacteria | Epsilonproteobacteria | Nautiliales | Nautiliaceae | *Nitratifractor* | 0.001 | 0.000 | 0.000 | 0.000 | 0.000 | 0.000 |
| Proteobacteria | Epsilonproteobacteria | Nautiliales | Nautiliaceae | *Nitratiruptor* | 0.001 | 0.000 | 0.000 | 0.000 | 0.000 | 0.000 |
| Proteobacteria | Epsilonproteobacteria | Unclassified | Unclassified | Unclassified | 0.001 | 0.000 | 0.000 | 0.000 | 0.000 | 0.000 |
| Proteobacteria | Gammaproteobacteria | Aeromonadales | Aeromonadaceae | *Aeromonas* | 0.000 | 0.000 | 0.039 | 0.032 | 0.000 | 0.001 |
| Proteobacteria | Gammaproteobacteria | Aeromonadales | Aeromonadaceae | *Oceanisphaera* | 0.000 | 0.000 | 0.000 | 0.004 | 0.000 | 0.000 |
| Proteobacteria | Gammaproteobacteria | Alteromonadales | Alteromonadaceae | *Alishewanella* | 0.000 | 0.000 | 0.013 | 0.011 | 0.000 | 0.000 |
| Proteobacteria | Gammaproteobacteria | Alteromonadales | Colwelliaceae | *Colwellia* | 0.000 | 0.000 | 0.000 | 0.000 | 0.000 | 0.001 |
| Proteobacteria | Gammaproteobacteria | Alteromonadales | Pseudoalteromonadaceae | *Pseudoalteromonas* | 0.000 | 0.032 | 0.000 | 0.000 | 0.000 | 0.000 |
| Proteobacteria | Gammaproteobacteria | Alteromonadales | Shewanellaceae | *Shewanella* | 0.000 | 0.000 | 0.000 | 0.007 | 0.004 | 0.001 |
| Proteobacteria | Gammaproteobacteria | Cardiobacteriales | Cardiobacteriaceae | Uncultured | 0.001 | 0.000 | 0.000 | 0.004 | 0.000 | 0.000 |
| Proteobacteria | Gammaproteobacteria | Cellvibrionales | Cellvibrionaceae | *Cellvibrio* | 0.000 | 0.000 | 0.000 | 0.039 | 0.000 | 0.000 |
| Proteobacteria | Gammaproteobacteria | Cellvibrionales | Porticoccaceae | *Porticoccus* | 0.000 | 0.000 | 0.000 | 0.004 | 0.000 | 0.000 |
| Proteobacteria | Gammaproteobacteria | Cellvibrionales | Spongiibacteraceae | *Melitea* | 0.001 | 0.000 | 0.000 | 0.000 | 0.000 | 0.000 |
| Proteobacteria | Gammaproteobacteria | Chromatiales | Chromatiaceae | *Rheinheimera* | 0.000 | 0.000 | 0.000 | 0.242 | 0.000 | 0.000 |
| Proteobacteria | Gammaproteobacteria | Chromatiales | Ectothiorhodospiraceae | *Ectothiorhodospira* | 0.000 | 0.000 | 0.000 | 0.000 | 0.004 | 0.000 |

**Table S2 continued.**

| | | | | | | | | | | |
|---|---|---|---|---|---|---|---|---|---|---|
| Proteobacteria | Gammaproteobacteria | Chromatiales | Ectothiorhodospiraceae | *Thioalkalivibrio* | 0.000 | 0.000 | 0.000 | 0.004 | 0.000 | 0.000 |
| Proteobacteria | Gammaproteobacteria | Chromatiales | Halothiobacillaceae | *Halothiobacillus* | 0.000 | 0.000 | 0.000 | 0.004 | 0.000 | 0.000 |
| Proteobacteria | Gammaproteobacteria | Chromatiales | Woeseiaceae | *Woeseia* | 0.000 | 0.000 | 0.000 | 0.004 | 0.000 | 0.000 |
| Proteobacteria | Gammaproteobacteria | Enterobacterales | Enterobacteriaceae | *Buttiauxella* | 0.004 | 0.000 | 0.013 | 0.007 | 0.004 | 0.002 |
| Proteobacteria | Gammaproteobacteria | Enterobacterales | Enterobacteriaceae | *Citrobacter* | 0.000 | 0.000 | 0.013 | 0.011 | 0.004 | 0.000 |
| Proteobacteria | Gammaproteobacteria | Enterobacterales | Enterobacteriaceae | *Cronobacter* | 0.002 | 0.000 | 0.000 | 0.007 | 0.004 | 0.000 |
| Proteobacteria | Gammaproteobacteria | Enterobacterales | Enterobacteriaceae | *Enterobacter* | 0.023 | 0.000 | 0.039 | 0.043 | 0.008 | 0.005 |
| Proteobacteria | Gammaproteobacteria | Enterobacterales | Enterobacteriaceae | *Escherichia* | 0.001 | 0.000 | 0.000 | 0.050 | 0.000 | 0.000 |
| Proteobacteria | Gammaproteobacteria | Enterobacterales | Enterobacteriaceae | *Klebsiella* | 0.001 | 0.000 | 0.013 | 0.014 | 0.000 | 0.000 |
| Proteobacteria | Gammaproteobacteria | Enterobacterales | Enterobacteriaceae | *Kluyvera* | 0.005 | 0.032 | 0.013 | 0.014 | 0.000 | 0.003 |
| Proteobacteria | Gammaproteobacteria | Enterobacterales | Enterobacteriaceae | *Kosakonia* | 0.001 | 0.000 | 0.000 | 0.032 | 0.000 | 0.000 |
| Proteobacteria | Gammaproteobacteria | Enterobacterales | Enterobacteriaceae | *Leclercia* | 0.001 | 0.000 | 0.000 | 0.004 | 0.000 | 0.005 |
| Proteobacteria | Gammaproteobacteria | Enterobacterales | Enterobacteriaceae | *Pseudocitrobacter* | 0.001 | 0.000 | 0.000 | 0.000 | 0.000 | 0.000 |
| Proteobacteria | Gammaproteobacteria | Enterobacterales | Enterobacteriaceae | *Raoultella* | 0.000 | 0.000 | 0.000 | 0.004 | 0.000 | 0.000 |
| Proteobacteria | Gammaproteobacteria | Enterobacterales | Enterobacteriaceae | *Salmonella* | 0.074 | 0.032 | 0.000 | 0.082 | 0.049 | 0.009 |
| Proteobacteria | Gammaproteobacteria | Enterobacterales | Enterobacteriaceae | *Shigella* | 0.000 | 0.000 | 0.000 | 0.043 | 0.000 | 0.001 |
| Proteobacteria | Gammaproteobacteria | Enterobacterales | Enterobacteriaceae | *Shimwellia* | 0.000 | 0.000 | 0.000 | 0.004 | 0.000 | 0.000 |
| Proteobacteria | Gammaproteobacteria | Enterobacterales | Enterobacteriaceae | *Trabulsiella* | 0.183 | 0.191 | 0.026 | 0.253 | 0.061 | 0.094 |
| Proteobacteria | Gammaproteobacteria | Enterobacterales | Erwiniaceae | *Erwinia* | 0.002 | 0.000 | 0.013 | 0.007 | 0.000 | 0.001 |
| Proteobacteria | Gammaproteobacteria | Enterobacterales | Erwiniaceae | *Pantoea* | 0.005 | 0.000 | 0.026 | 0.057 | 0.008 | 0.001 |
| Proteobacteria | Gammaproteobacteria | Enterobacterales | Morganellaceae | *Morganella* | 0.000 | 0.000 | 0.000 | 0.000 | 0.000 | 0.001 |
| Proteobacteria | Gammaproteobacteria | Enterobacterales | Morganellaceae | *Providencia* | 0.001 | 0.000 | 0.000 | 0.014 | 0.000 | 0.006 |
| Proteobacteria | Gammaproteobacteria | Enterobacterales | Pectobacteriaceae | *Pectobacterium* | 0.001 | 0.000 | 0.000 | 0.032 | 0.000 | 0.001 |
| Proteobacteria | Gammaproteobacteria | Enterobacterales | Yersiniaceae | *Serratia* | 0.000 | 0.000 | 0.000 | 0.036 | 0.000 | 0.000 |
| Proteobacteria | Gammaproteobacteria | Enterobacterales | Yersiniaceae | *Yersinia* | 0.000 | 0.000 | 0.000 | 0.007 | 0.000 | 0.001 |
| Proteobacteria | Gammaproteobacteria | Legionellales | Coxiellaceae | *Diplorickettsia* | 0.000 | 0.000 | 0.000 | 0.004 | 0.000 | 0.000 |
| Proteobacteria | Gammaproteobacteria | Legionellales | Legionellaceae | *Fluoribacter* | 0.000 | 0.000 | 0.000 | 0.011 | 0.000 | 0.000 |

**Table S2 continued.**

| | | | | | | | | | | |
|---|---|---|---|---|---|---|---|---|---|---|
| Proteobacteria | Gammaproteobacteria | Legionellales | Legionellaceae | *Legionella* | 0.001 | 0.000 | 0.000 | 0.093 | 0.000 | 0.000 |
| Proteobacteria | Gammaproteobacteria | Legionellales | Legionellaceae | *Tatlockia* | 0.000 | 0.000 | 0.000 | 0.007 | 0.000 | 0.000 |
| Proteobacteria | Gammaproteobacteria | Methylococcales | Methylococcaceae | *Methylocaldum* | 0.000 | 0.000 | 0.000 | 0.004 | 0.000 | 0.000 |
| Proteobacteria | Gammaproteobacteria | Methylococcales | Methylothermaceae | *Methylohalobius* | 0.000 | 0.000 | 0.000 | 0.004 | 0.000 | 0.000 |
| Proteobacteria | Gammaproteobacteria | Nevskiales | Algiphilaceae | *Algiphilus* | 0.000 | 0.000 | 0.000 | 0.004 | 0.000 | 0.000 |
| Proteobacteria | Gammaproteobacteria | Nevskiales | Sinobacteraceae | *Fontimonas* | 0.000 | 0.000 | 0.000 | 0.011 | 0.000 | 0.000 |
| Proteobacteria | Gammaproteobacteria | Nevskiales | Sinobacteraceae | *Hydrocarboniphaga* | 0.000 | 0.000 | 0.000 | 0.021 | 0.000 | 0.000 |
| Proteobacteria | Gammaproteobacteria | Nevskiales | Sinobacteraceae | *Nevskia* | 0.000 | 0.000 | 0.000 | 0.021 | 0.000 | 0.000 |
| Proteobacteria | Gammaproteobacteria | Nevskiales | Sinobacteraceae | *Panacagrimonas* | 0.000 | 0.000 | 0.000 | 0.071 | 0.000 | 0.000 |
| Proteobacteria | Gammaproteobacteria | Nevskiales | Sinobacteraceae | *Solimonas* | 0.000 | 0.000 | 0.000 | 0.014 | 0.000 | 0.000 |
| Proteobacteria | Gammaproteobacteria | Nevskiales | Steroidobacteraceae | *Povalibacter* | 0.000 | 0.000 | 0.000 | 0.018 | 0.000 | 0.000 |
| Proteobacteria | Gammaproteobacteria | Nevskiales | Steroidobacteraceae | *Steroidobacter* | 0.000 | 0.000 | 0.000 | 0.107 | 0.000 | 0.000 |
| Proteobacteria | Gammaproteobacteria | Oceanospirillales | Alcanivoracaceae | *Alcanivorax* | 0.000 | 0.000 | 0.000 | 0.004 | 0.000 | 0.000 |
| Proteobacteria | Gammaproteobacteria | Oceanospirillales | Halomonadaceae | *Aidingimonas* | 0.000 | 0.000 | 0.000 | 0.004 | 0.000 | 0.000 |
| Proteobacteria | Gammaproteobacteria | Oceanospirillales | Oceanospirillaceae | *Marinobacterium* | 0.000 | 0.000 | 0.000 | 0.004 | 0.000 | 0.000 |
| Proteobacteria | Gammaproteobacteria | Oceanospirillales | Oceanospirillaceae | *Marinomonas* | 0.000 | 0.000 | 0.000 | 0.004 | 0.000 | 0.000 |
| Proteobacteria | Gammaproteobacteria | Pasteurellales | Pasteurellaceae | *Haemophilus* | 0.000 | 0.000 | 0.000 | 0.011 | 0.000 | 0.000 |
| Proteobacteria | Gammaproteobacteria | Pseudomonadales | Moraxellaceae | *Acinetobacter* | 0.009 | 0.032 | 0.142 | 2.722 | 0.008 | 0.015 |
| Proteobacteria | Gammaproteobacteria | Pseudomonadales | Moraxellaceae | *Alkanindiges* | 0.000 | 0.000 | 0.000 | 0.021 | 0.000 | 0.000 |
| Proteobacteria | Gammaproteobacteria | Pseudomonadales | Moraxellaceae | *Moraxella* | 0.000 | 0.000 | 0.013 | 0.014 | 0.016 | 0.006 |
| Proteobacteria | Gammaproteobacteria | Pseudomonadales | Moraxellaceae | *Perlucidibaca* | 0.000 | 0.000 | 0.013 | 0.018 | 0.000 | 0.000 |
| Proteobacteria | Gammaproteobacteria | Pseudomonadales | Moraxellaceae | *Psychrobacter* | 0.000 | 0.000 | 0.000 | 0.004 | 0.000 | 0.001 |
| Proteobacteria | Gammaproteobacteria | Pseudomonadales | Pseudomonadaceae | *Pseudomonas* | 0.002 | 0.000 | 0.077 | 0.210 | 0.033 | 0.016 |
| Proteobacteria | Gammaproteobacteria | Thiotrichales | Piscirickettsiaceae | endosymbionts 2 | 0.000 | 0.000 | 0.000 | 0.004 | 0.000 | 0.000 |
| Proteobacteria | Gammaproteobacteria | Thiotrichales | Piscirickettsiaceae | Unclassified | 0.000 | 0.000 | 0.000 | 0.000 | 0.000 | 0.001 |
| Proteobacteria | Gammaproteobacteria | Thiotrichales | Piscirickettsiaceae | Unclassified | 0.001 | 0.000 | 0.000 | 0.000 | 0.000 | 0.000 |
| Proteobacteria | Gammaproteobacteria | Thiotrichales | Thiotrichaceae | *Thiothrix* | 0.000 | 0.000 | 0.000 | 0.004 | 0.000 | 0.000 |

**Table S2 continued.**

| | | | | | | | | | | |
|---|---|---|---|---|---|---|---|---|---|---|
| Proteobacteria | Gammaproteobacteria | Unclassified | Unclassified | *Ignatzschineria* | 0.000 | 0.000 | 0.000 | 0.004 | 0.000 | 0.000 |
| Proteobacteria | Gammaproteobacteria | Unclassified | Unclassified | *Methylohalomonas* | 0.000 | 0.000 | 0.000 | 0.004 | 0.000 | 0.000 |
| Proteobacteria | Gammaproteobacteria | Vibrionales | Vibrionaceae | *Aliivibrio* | 0.000 | 0.000 | 0.000 | 0.000 | 0.004 | 0.000 |
| Proteobacteria | Gammaproteobacteria | Xanthomonadales | Rhodanobacteraceae | *Aquimonas* | 0.000 | 0.000 | 0.000 | 0.004 | 0.000 | 0.000 |
| Proteobacteria | Gammaproteobacteria | Xanthomonadales | Rhodanobacteraceae | *Dyella* | 0.000 | 0.000 | 0.000 | 0.021 | 0.000 | 0.000 |
| Proteobacteria | Gammaproteobacteria | Xanthomonadales | Rhodanobacteraceae | *Frateuria* | 0.000 | 0.000 | 0.000 | 0.004 | 0.000 | 0.000 |
| Proteobacteria | Gammaproteobacteria | Xanthomonadales | Rhodanobacteraceae | *Fulvimonas* | 0.000 | 0.000 | 0.000 | 0.004 | 0.000 | 0.000 |
| Proteobacteria | Gammaproteobacteria | Xanthomonadales | Rhodanobacteraceae | *Luteibacter* | 0.000 | 0.032 | 0.013 | 0.071 | 0.000 | 0.000 |
| Proteobacteria | Gammaproteobacteria | Xanthomonadales | Xanthomonadaceae | *Arenimonas* | 0.000 | 0.000 | 0.000 | 0.021 | 0.000 | 0.000 |
| Proteobacteria | Gammaproteobacteria | Xanthomonadales | Xanthomonadaceae | *Dokdonella* | 0.000 | 0.000 | 0.000 | 0.004 | 0.000 | 0.000 |
| Proteobacteria | Gammaproteobacteria | Xanthomonadales | Xanthomonadaceae | *Luteimonas* | 0.000 | 0.000 | 0.000 | 0.043 | 0.000 | 0.000 |
| Proteobacteria | Gammaproteobacteria | Xanthomonadales | Xanthomonadaceae | *Luteimonas* 3 | 0.000 | 0.000 | 0.000 | 0.004 | 0.000 | 0.001 |
| Proteobacteria | Gammaproteobacteria | Xanthomonadales | Xanthomonadaceae | *Lysobacter* | 0.000 | 0.032 | 0.000 | 0.068 | 0.000 | 0.000 |
| Proteobacteria | Gammaproteobacteria | Xanthomonadales | Xanthomonadaceae | *Lysobacter* 1 | 0.000 | 0.000 | 0.000 | 0.004 | 0.000 | 0.000 |
| Proteobacteria | Gammaproteobacteria | Xanthomonadales | Xanthomonadaceae | *Lysobacter* 2 | 0.000 | 0.000 | 0.000 | 0.021 | 0.000 | 0.000 |
| Proteobacteria | Gammaproteobacteria | Xanthomonadales | Xanthomonadaceae | *Lysobacter* 3 | 0.000 | 0.000 | 0.000 | 0.011 | 0.000 | 0.000 |
| Proteobacteria | Gammaproteobacteria | Xanthomonadales | Xanthomonadaceae | *Pseudoxanthomonas* | 0.004 | 0.000 | 0.013 | 0.164 | 0.004 | 0.002 |
| Proteobacteria | Gammaproteobacteria | Xanthomonadales | Xanthomonadaceae | *Rhodanobacter* 1 | 0.000 | 0.000 | 0.000 | 0.004 | 0.000 | 0.000 |
| Proteobacteria | Gammaproteobacteria | Xanthomonadales | Xanthomonadaceae | *Silanimonas* | 0.000 | 0.000 | 0.013 | 0.004 | 0.000 | 0.000 |
| Proteobacteria | Gammaproteobacteria | Xanthomonadales | Xanthomonadaceae | *Stenotrophomonas* | 0.001 | 0.000 | 0.000 | 0.114 | 0.000 | 0.001 |
| Proteobacteria | Gammaproteobacteria | Xanthomonadales | Xanthomonadaceae | *Thermomonas* | 0.000 | 0.000 | 0.000 | 0.004 | 0.000 | 0.000 |
| Proteobacteria | Gammaproteobacteria | Xanthomonadales | Xanthomonadaceae | *Thermomonas* 1 | 0.000 | 0.000 | 0.000 | 0.011 | 0.000 | 0.000 |
| Proteobacteria | Gammaproteobacteria | Xanthomonadales | Xanthomonadaceae | *Thermomonas* 2 | 0.001 | 0.000 | 0.000 | 0.007 | 0.000 | 0.000 |
| Proteobacteria | Gammaproteobacteria | Xanthomonadales | Xanthomonadaceae | Unclassified | 0.000 | 0.000 | 0.000 | 0.014 | 0.004 | 0.000 |
| Proteobacteria | Gammaproteobacteria | Xanthomonadales | Xanthomonadaceae | Uncultured 5 | 0.000 | 0.000 | 0.000 | 0.061 | 0.000 | 0.000 |
| Proteobacteria | Gammaproteobacteria | Xanthomonadales | Xanthomonadaceae | *Xanthomonas* | 0.000 | 0.000 | 0.000 | 0.011 | 0.012 | 0.000 |
| Proteobacteria | Gammaproteobacteria 1 | Alteromonadales | Alteromonadaceae | *Marinobacter* | 0.000 | 0.000 | 0.000 | 0.000 | 0.000 | 0.001 |

**Table S2 continued.**

| | | | | | | | | | | |
|---|---|---|---|---|---|---|---|---|---|---|
| Proteobacteria | Gammaproteobacteria 1 | Alteromonadales | Alteromonadaceae | OM60 NOR5 clade | 0.000 | 0.000 | 0.000 | 0.004 | 0.000 | 0.001 |
| Proteobacteria | Gammaproteobacteria 1 | Alteromonadales 1 | Alteromonadaceae 1 | *Glaciecola* | 0.000 | 0.000 | 0.000 | 0.004 | 0.000 | 0.000 |
| Proteobacteria | Gammaproteobacteria 1 | Alteromonadales 1 | Alteromonadaceae 1 | Unclassified | 0.001 | 0.000 | 0.000 | 0.000 | 0.000 | 0.000 |
| Proteobacteria | Gammaproteobacteria 1 | Alteromonadales 1 | Psychromonadaceae | *Psychromonas* | 0.000 | 0.000 | 0.000 | 0.000 | 0.004 | 0.000 |
| Proteobacteria | Gammaproteobacteria 1 | B38 | Unclassified | Unclassified | 0.000 | 0.000 | 0.013 | 0.018 | 0.000 | 0.000 |
| Proteobacteria | Gammaproteobacteria 1 | B38 | Unclassified | Unclassified | 0.005 | 0.000 | 0.000 | 0.000 | 0.000 | 0.000 |
| Proteobacteria | Gammaproteobacteria 1 | Chromatiales | Chromatiaceae | *Rheinheimera* 1 | 0.000 | 0.000 | 0.000 | 0.064 | 0.000 | 0.000 |
| Proteobacteria | Gammaproteobacteria 1 | Chromatiales | Chromatiaceae | *Rheinheimera* 3 | 0.000 | 0.000 | 0.000 | 0.011 | 0.000 | 0.000 |
| Proteobacteria | Gammaproteobacteria 1 | Chromatiales | Chromatiaceae | *Rheinheimera* 4 | 0.000 | 0.000 | 0.000 | 0.007 | 0.000 | 0.000 |
| Proteobacteria | Gammaproteobacteria 1 | Chromatiales | Chromatiaceae | Unclassified | 0.000 | 0.000 | 0.000 | 0.004 | 0.000 | 0.000 |
| Proteobacteria | Gammaproteobacteria 1 | Chromatiales | Chromatiaceae | Uncultured | 0.000 | 0.000 | 0.000 | 0.011 | 0.000 | 0.000 |
| Proteobacteria | Gammaproteobacteria 1 | Enterobacteriales | Enterobacteriaceae | *Brenneria*-Erwinia-Pectobacterium | 0.001 | 0.000 | 0.013 | 0.061 | 0.000 | 0.003 |
| Proteobacteria | Gammaproteobacteria 1 | Enterobacteriales | Enterobacteriaceae | *Citrobacter* 1 | 0.016 | 0.000 | 0.000 | 0.032 | 0.016 | 0.009 |
| Proteobacteria | Gammaproteobacteria 1 | Enterobacteriales | Enterobacteriaceae | *Citrobacter* 2 | 0.012 | 0.000 | 0.000 | 0.032 | 0.073 | 0.007 |
| Proteobacteria | Gammaproteobacteria 1 | Enterobacteriales | Enterobacteriaceae | *Enterobacter* 1 | 0.048 | 0.032 | 0.077 | 0.132 | 0.041 | 0.024 |
| Proteobacteria | Gammaproteobacteria 1 | Enterobacteriales | Enterobacteriaceae | *Enterobacter* 2 | 0.002 | 0.000 | 0.000 | 0.000 | 0.000 | 0.000 |
| Proteobacteria | Gammaproteobacteria 1 | Enterobacteriales | Enterobacteriaceae | *Enterobacter* 4 | 0.015 | 0.032 | 0.026 | 0.043 | 0.016 | 0.008 |
| Proteobacteria | Gammaproteobacteria 1 | Enterobacteriales | Enterobacteriaceae | *Escherichia blattae* cluster | 0.000 | 0.000 | 0.000 | 0.007 | 0.000 | 0.001 |
| Proteobacteria | Gammaproteobacteria 1 | Enterobacteriales | Enterobacteriaceae | *Escherichia-Shigella* | 0.026 | 0.000 | 0.064 | 0.449 | 0.000 | 0.005 |
| Proteobacteria | Gammaproteobacteria 1 | Enterobacteriales | Enterobacteriaceae | Gut cluster | 0.001 | 0.000 | 0.000 | 0.031 | 0.000 | 0.000 |
| Proteobacteria | Gammaproteobacteria 1 | Enterobacteriales | Enterobacteriaceae | *Klebsiella* 1 | 0.005 | 0.032 | 0.077 | 0.046 | 0.000 | 0.001 |
| Proteobacteria | Gammaproteobacteria 1 | Enterobacteriales | Enterobacteriaceae | *Klebsiella* 2 | 0.004 | 0.000 | 0.000 | 0.007 | 0.004 | 0.001 |
| Proteobacteria | Gammaproteobacteria 1 | Enterobacteriales | Enterobacteriaceae | *Phytobacter* | 0.001 | 0.000 | 0.000 | 0.007 | 0.000 | 0.000 |
| Proteobacteria | Gammaproteobacteria 1 | Enterobacteriales | Enterobacteriaceae | *Plesiomonas* | 0.001 | 0.000 | 0.000 | 0.000 | 0.000 | 0.000 |
| Proteobacteria | Gammaproteobacteria 1 | Enterobacteriales | Enterobacteriaceae | *Rahnella* | 0.000 | 0.000 | 0.000 | 0.004 | 0.000 | 0.000 |
| Proteobacteria | Gammaproteobacteria 1 | Enterobacteriales | Enterobacteriaceae | *Salmonella* | 0.000 | 0.000 | 0.000 | 0.000 | 0.000 | 0.000 |
| Proteobacteria | Gammaproteobacteria 1 | Enterobacteriales | Enterobacteriaceae | *Serratia* 1 | 0.000 | 0.000 | 0.000 | 0.018 | 0.000 | 0.002 |

**Table S2 continued.**

| | | | | | | | | | | |
|---|---|---|---|---|---|---|---|---|---|---|
| Proteobacteria | Gammaproteobacteria 1 | Enterobacteriales | Enterobacteriaceae | *Tatumella* | 0.001 | 0.000 | 0.013 | 0.014 | 0.000 | 0.000 |
| Proteobacteria | Gammaproteobacteria 1 | Enterobacteriales | Enterobacteriaceae | *Yokenella* | 0.000 | 0.000 | 0.000 | 0.000 | 0.000 | 0.001 |
| Proteobacteria | Gammaproteobacteria 1 | Enterobacteriales | Enterobacteriaceae 1 | *Arsenophonus* | 0.000 | 0.000 | 0.000 | 0.004 | 0.000 | 0.000 |
| Proteobacteria | Gammaproteobacteria 1 | Enterobacteriales | Enterobacteriaceae 2 | *Candidatus* Hamiltonella | 0.000 | 0.000 | 0.013 | 0.000 | 0.000 | 0.000 |
| Proteobacteria | Gammaproteobacteria 1 | Enterobacteriales | Enterobacteriaceae 3 | Unclassified | 0.000 | 0.000 | 0.000 | 0.000 | 0.004 | 0.000 |
| Proteobacteria | Gammaproteobacteria 1 | Fish gut cluster | Unclassified | Unclassified | 0.000 | 0.000 | 0.013 | 0.000 | 0.000 | 0.000 |
| Proteobacteria | Gammaproteobacteria 1 | Oceanospirillales | Oceanospirillaceae | *Pseudospirillum* | 0.000 | 0.000 | 0.000 | 0.018 | 0.000 | 0.000 |
| Proteobacteria | Gammaproteobacteria 1 | Oceanospirillales | Oceanospirillaceae 1 | *Amphritea* | 0.000 | 0.000 | 0.000 | 0.004 | 0.000 | 0.000 |
| Proteobacteria | Gammaproteobacteria 1 | Oceanospirillales | Oceanospirillaceae 1 | *Nitrincola* | 0.000 | 0.000 | 0.013 | 0.000 | 0.000 | 0.000 |
| Proteobacteria | Gammaproteobacteria 1 | Oceanospirillales 1 | Halomonadaceae | *Halomonas* 1 | 0.001 | 0.000 | 0.000 | 0.004 | 0.000 | 0.000 |
| Proteobacteria | Gammaproteobacteria 1 | Oceanospirillales 2 | Oceanospirillaceae | *Reinekea* | 0.000 | 0.000 | 0.000 | 0.000 | 0.004 | 0.000 |
| Proteobacteria | Gammaproteobacteria 1 | Oceanospirillales 3 | Oceanospirillaceae | *Balneatrix* | 0.001 | 0.000 | 0.000 | 0.000 | 0.000 | 0.000 |
| Proteobacteria | Gammaproteobacteria 1 | Oceanospirillales 3 | Unclassified | Unclassified | 0.000 | 0.000 | 0.000 | 0.004 | 0.000 | 0.000 |
| Proteobacteria | Gammaproteobacteria 1 | Pasteurellales | Pasteurellaceae | *Haemophilus* 2 | 0.000 | 0.000 | 0.000 | 0.004 | 0.000 | 0.000 |
| Proteobacteria | Gammaproteobacteria 1 | Pasteurellales | Pasteurellaceae | *Haemophilus* 7 | 0.001 | 0.000 | 0.000 | 0.000 | 0.000 | 0.000 |
| Proteobacteria | Gammaproteobacteria 1 | Pseudomonadales | Moraxellaceae | Unclassified | 0.000 | 0.000 | 0.000 | 0.004 | 0.000 | 0.000 |
| Proteobacteria | Gammaproteobacteria 1 | Pseudomonadales | Pseudomonadaceae | *Azotobacter-Azomonas* | 0.000 | 0.000 | 0.000 | 0.011 | 0.000 | 0.000 |
| Proteobacteria | Gammaproteobacteria 1 | Pseudomonadales | Pseudomonadaceae | *Pseudomonas* 1 | 0.004 | 0.000 | 0.052 | 0.310 | 0.073 | 0.049 |
| Proteobacteria | Gammaproteobacteria 1 | Pseudomonadales | Pseudomonadaceae | *Pseudomonas* 2 | 0.001 | 0.000 | 0.013 | 0.100 | 0.004 | 0.005 |
| Proteobacteria | Gammaproteobacteria 1 | Pseudomonadales | Pseudomonadaceae | *Pseudomonas* 3 | 0.000 | 0.000 | 0.000 | 0.014 | 0.000 | 0.000 |
| Proteobacteria | Gammaproteobacteria 1 | Pseudomonadales | Pseudomonadaceae | *Pseudomonas* 4 | 0.000 | 0.000 | 0.000 | 0.025 | 0.000 | 0.000 |
| Proteobacteria | Gammaproteobacteria 1 | Pseudomonadales | Pseudomonadaceae | *Pseudomonas* 6 | 0.001 | 0.000 | 0.000 | 0.000 | 0.000 | 0.000 |
| Proteobacteria | Gammaproteobacteria 1 | Pseudomonadales | Pseudomonadaceae | *Pseudomonas* 7 | 0.000 | 0.000 | 0.000 | 0.004 | 0.000 | 0.000 |
| Proteobacteria | Gammaproteobacteria 1 | Vibrionales | Vibrionaceae | *Vibrio-Alivibrio* | 0.001 | 0.033 | 0.013 | 0.004 | 0.000 | 0.001 |
| Proteobacteria | Gammaproteobacteria 2 | Chromatiales | Chromatiaceae | *Nitrosococcus* | 0.000 | 0.000 | 0.000 | 0.018 | 0.000 | 0.000 |
| Proteobacteria | Gammaproteobacteria 2 | Chromatiales | Ectothiorhodospiraceae 1 | *Thioalkalivibrio* 1 | 0.000 | 0.000 | 0.000 | 0.004 | 0.000 | 0.000 |
| Proteobacteria | Gammaproteobacteria 2 | Chromatiales | Ectothiorhodospiraceae 1 | *Thiorhodospira* | 0.000 | 0.000 | 0.000 | 0.000 | 0.000 | 0.001 |

**Table S2 continued.**

| | | | | | | | | | | |
|---|---|---|---|---|---|---|---|---|---|---|
| Proteobacteria | Gammaproteobacteria 2 | Chromatiales | Ectothiorhodospiraceae 2 | Uncultured | 0.000 | 0.000 | 0.000 | 0.004 | 0.000 | 0.000 |
| Proteobacteria | Gammaproteobacteria 2 | Chromatiales | Ectothiorhodospiraceae 3 | Unclassified | 0.000 | 0.000 | 0.000 | 0.004 | 0.000 | 0.000 |
| Proteobacteria | Gammaproteobacteria 2 | Legionellales | Coxiellaceae | *Aquicella* | 0.000 | 0.000 | 0.000 | 0.232 | 0.000 | 0.000 |
| Proteobacteria | Gammaproteobacteria 2 | Legionellales | Coxiellaceae | *Coxiella* | 0.002 | 0.032 | 0.000 | 0.064 | 0.000 | 0.000 |
| Proteobacteria | Gammaproteobacteria 2 | Legionellales | Coxiellaceae | *Rickettsiella* | 0.001 | 0.000 | 0.026 | 0.135 | 0.000 | 0.000 |
| Proteobacteria | Gammaproteobacteria 2 | Legionellales | Coxiellaceae | Uncultured | 0.000 | 0.000 | 0.000 | 0.004 | 0.000 | 0.000 |
| Proteobacteria | Gammaproteobacteria 2 | Legionellales | Legionellaceae | *Legionella* 1 | 0.000 | 0.000 | 0.000 | 0.021 | 0.000 | 0.000 |
| Proteobacteria | Gammaproteobacteria 2 | Legionellales | Legionellaceae | *Legionella* 5 | 0.000 | 0.000 | 0.000 | 0.004 | 0.000 | 0.000 |
| Proteobacteria | Gammaproteobacteria 2 | Legionellales | Legionellaceae | *Legionella* 9 | 0.000 | 0.000 | 0.000 | 0.004 | 0.000 | 0.000 |
| Proteobacteria | Gammaproteobacteria 2 | Legionellales | Legionellaceae | Unclassified | 0.000 | 0.000 | 0.000 | 0.004 | 0.000 | 0.000 |
| Proteobacteria | Gammaproteobacteria 2 | Legionellales | Legionellaceae | Uncultured | 0.000 | 0.000 | 0.000 | 0.004 | 0.000 | 0.000 |
| Proteobacteria | Gammaproteobacteria 2 | marine group E01-9C-26 | Unclassified | Unclassified | 0.001 | 0.000 | 0.000 | 0.000 | 0.000 | 0.000 |
| Proteobacteria | Gammaproteobacteria 2 | Methylonatrum | Methylonatrum | *Methylonatrum* | 0.000 | 0.000 | 0.000 | 0.004 | 0.000 | 0.000 |
| Proteobacteria | Gammaproteobacteria 2 | NKB5 | Unclassified | Unclassified | 0.000 | 0.000 | 0.000 | 0.018 | 0.000 | 0.000 |
| Proteobacteria | Gammaproteobacteria 2 | Thiotrichales | EV818SWSAP88 | Unclassified | 0.000 | 0.000 | 0.000 | 0.004 | 0.000 | 0.000 |
| Proteobacteria | Gammaproteobacteria 2 | Xanthomonadales | Sinobacteraceae | marine benthic group | 0.000 | 0.000 | 0.000 | 0.007 | 0.000 | 0.000 |
| Proteobacteria | Gammaproteobacteria 2 | Xanthomonadales | Sinobacteraceae | Unclassified | 0.000 | 0.000 | 0.000 | 0.011 | 0.000 | 0.000 |
| Proteobacteria | Gammaproteobacteria 2 | Xanthomonadales | Sinobacteraceae | Uncultured 2 | 0.000 | 0.000 | 0.000 | 0.021 | 0.000 | 0.000 |
| Proteobacteria | Gammaproteobacteria 2 | Xanthomonadales | Sinobacteraceae | Uncultured 3 | 0.000 | 0.000 | 0.000 | 0.018 | 0.000 | 0.000 |
| Proteobacteria | Gammaproteobacteria 2 | Xanthomonadales | Sinobacteraceae | Uncultured 4 | 0.000 | 0.000 | 0.000 | 0.011 | 0.000 | 0.000 |
| Proteobacteria | Gammaproteobacteria 2 | Xanthomonadales | Sinobacteraceae | Uncultured 5 | 0.001 | 0.000 | 0.000 | 0.021 | 0.000 | 0.000 |
| Proteobacteria | Gammaproteobacteria 2 | Xanthomonadales | Sinobacteraceae | Uncultured 6 | 0.001 | 0.000 | 0.000 | 0.061 | 0.000 | 0.000 |
| Proteobacteria | Gammaproteobacteria 2 | Xanthomonadales | Sinobacteraceae | Uncultured 8 | 0.001 | 0.000 | 0.000 | 0.060 | 0.000 | 0.000 |
| Proteobacteria | Gammaproteobacteria 3 | JTB148 | Unclassified | Unclassified | 0.000 | 0.000 | 0.000 | 0.004 | 0.000 | 0.000 |
| Proteobacteria | Gammaproteobacteria 4 | Methylococcales | Hyd24-01 | Unclassified | 0.000 | 0.000 | 0.000 | 0.000 | 0.000 | 0.001 |
| Proteobacteria | Gammaproteobacteria 4 | Unclassified | Unclassified | Unclassified | 0.000 | 0.000 | 0.000 | 0.004 | 0.000 | 0.000 |
| Proteobacteria | Hydrogenophilalia | Hydrogenophilales | Hydrogenophilaceae | *Hydrogenophilus* | 0.005 | 0.000 | 0.000 | 0.000 | 0.016 | 0.002 |

**Table S2 continued.**

| | | | | | | | | | | |
|---|---|---|---|---|---|---|---|---|---|---|
| Proteobacteria | Hydrogenophilalia | Hydrogenophilales | Hydrogenophilaceae | *Tepidiphilus* | 0.007 | 0.000 | 0.000 | 0.011 | 0.020 | 0.001 |
| Proteobacteria | JTB23 | Unclassified | Unclassified | Unclassified | 0.000 | 0.000 | 0.000 | 0.004 | 0.000 | 0.000 |
| Proteobacteria | Oligoflexia | Bacteriovoracales | Bacteriovoracaceae | *Bacteriovorax* | 0.016 | 0.000 | 0.000 | 0.004 | 0.000 | 0.000 |
| Proteobacteria | Oligoflexia | Bacteriovoracales | Bacteriovoracaceae | *Peredibacter* | 0.052 | 0.000 | 0.000 | 0.011 | 0.000 | 0.000 |
| Proteobacteria | TA18 | Unclassified | Unclassified | Unclassified | 0.139 | 0.255 | 0.000 | 0.053 | 0.037 | 0.011 |
| Proteobacteria | Unclassified | Unclassified | Unclassified | Unclassified | 0.000 | 0.000 | 0.000 | 0.004 | 0.004 | 0.000 |
| Proteobacteria | Unclassified | Unclassified | Unclassified | Unclassified | 0.000 | 0.000 | 0.000 | 0.000 | 0.000 | 0.001 |
| Proteobacteria | Unclassified | Unclassified | Unclassified | Unclassified | 0.004 | 0.000 | 0.000 | 0.000 | 0.000 | 0.000 |
| Proteobacteria 1 | Deltaproteobacteria | Desulfobacterales | Nitrospinaceae | *Candidatus* Entotheonella | 0.000 | 0.000 | 0.000 | 0.011 | 0.000 | 0.000 |
| Proteobacteria 1 | Deltaproteobacteria | Desulfobacterales | Nitrospinaceae | Uncultured | 0.000 | 0.000 | 0.000 | 0.011 | 0.000 | 0.001 |
| Spirochaetes | Spirochaetes | Kazan-3B-09 | Unclassified | Unclassified | 0.000 | 0.032 | 0.000 | 0.000 | 0.000 | 0.000 |
| Spirochaetes | Spirochaetes | Spirochaetales | Leptospiraceae | *Leptospira* | 0.000 | 0.000 | 0.000 | 0.000 | 0.000 | 0.001 |
| Spirochaetes | Spirochaetes | Spirochaetales | Leptospiraceae | Rs-H88 termite group | 0.025 | 0.000 | 0.000 | 0.000 | 0.016 | 0.011 |
| Spirochaetes | Spirochaetes | Spirochaetales | Leptospiraceae | Uncultured 3 | 0.001 | 0.000 | 0.000 | 0.000 | 0.004 | 0.000 |
| Spirochaetes | Spirochaetes | Spirochaetales | Spirochaetaceae | *Spirochaeta* | 0.501 | 0.191 | 0.000 | 0.061 | 0.236 | 0.205 |
| Spirochaetes | Spirochaetes | Spirochaetales | Spirochaetaceae 10 | *Treponema caldaria* cluster | 0.032 | 0.096 | 0.000 | 0.025 | 0.008 | 0.007 |
| Spirochaetes | Spirochaetes | Spirochaetales | Spirochaetaceae 11 | Digestor cluster | 0.011 | 0.000 | 0.000 | 0.004 | 0.004 | 0.003 |
| Spirochaetes | Spirochaetes | Spirochaetales | Spirochaetaceae 5 | M2PT2-76 termite group | 0.016 | 0.000 | 0.000 | 0.004 | 0.000 | 0.002 |
| Spirochaetes | Spirochaetes | Spirochaetales | Spirochaetaceae Treponema | Animal cluster 1 | 0.001 | 0.032 | 0.000 | 0.000 | 0.000 | 0.000 |
| Spirochaetes | Spirochaetes | Spirochaetales | Spirochaetaceae Treponema | Animal cluster 2 | 0.006 | 0.000 | 0.000 | 0.000 | 0.012 | 0.000 |
| Spirochaetes | Spirochaetes | Spirochaetales | Spirochaetaceae Treponema | *Treponema* II | 0.006 | 0.000 | 0.000 | 0.004 | 0.000 | 0.000 |
| Spirochaetes | Spirochaetes | Spirochaetales | Spirochaetaceae Treponema | *Treponema* III | 0.000 | 1.308 | 0.000 | 0.096 | 0.004 | 0.023 |
| Spirochaetes | Spirochaetes | Spirochaetales | Spirochaetaceae Treponema I | *Treponema* Ia | 12.943 | 6.093 | 0.039 | 1.963 | 6.450 | 6.296 |
| Spirochaetes | Spirochaetes | Spirochaetales | Spirochaetaceae Treponema I | *Treponema* Ib | 0.134 | 0.064 | 0.026 | 0.053 | 0.037 | 0.040 |
| Spirochaetes | Spirochaetes | Spirochaetales | Spirochaetaceae Treponema I | *Treponema* Ic | 0.403 | 0.191 | 0.013 | 0.096 | 0.085 | 0.084 |
| Spirochaetes | Spirochaetes | Spirochaetales | Spirochaetaceae Treponema I | *Treponema* Id | 0.010 | 0.000 | 0.000 | 0.007 | 0.004 | 0.001 |
| Spirochaetes | Spirochaetes | Spirochaetales | Spirochaetaceae Treponema I | *Treponema* Ie | 1.784 | 0.893 | 0.000 | 0.353 | 0.805 | 0.959 |

**Table S2 continued.**

| | | | | | | | | | | |
|---|---|---|---|---|---|---|---|---|---|---|
| Spirochaetes | Spirochaetes | Spirochaetales | Spirochaetaceae Treponema I | *Treponema* If | 1.697 | 0.574 | 0.013 | 0.281 | 0.272 | 0.083 |
| Spirochaetes | Spirochaetes | Spirochaetales | Spirochaetaceae Treponema I | *Treponema* Ig | 0.114 | 0.064 | 0.000 | 0.021 | 0.016 | 0.028 |
| Spirochaetes | Spirochaetes | Spirochaetales | Spirochaetaceae Treponema I | *Treponema* Ih | 1.656 | 0.797 | 0.000 | 0.200 | 0.634 | 0.617 |
| Spirochaetes | Spirochaetia | Spirochaetales | Leptospiraceae | Uncultured 1 | 0.000 | 0.000 | 0.013 | 0.000 | 0.000 | 0.000 |
| Spirochaetes | Spirochaetia | Spirochaetales | Spirochaetaceae | *Treponema* | 0.038 | 0.000 | 0.000 | 0.000 | 0.004 | 0.002 |
| Synergistetes | Synergistia | Synergistales | Synergistaceae | *Acetomicrobium* | 0.000 | 0.032 | 0.000 | 0.000 | 0.004 | 0.000 |
| Synergistetes | Synergistia | Synergistales | Synergistaceae | *Aminomonas* | 0.001 | 0.000 | 0.000 | 0.000 | 0.000 | 0.000 |
| Synergistetes | Synergistia | Synergistales | Synergistaceae | *Candidatus* Tammella | 0.526 | 0.383 | 0.013 | 0.053 | 0.659 | 0.740 |
| Synergistetes | Synergistia | Synergistales | Synergistaceae | *Cloacibacillus* | 0.000 | 0.000 | 0.000 | 0.000 | 0.000 | 0.001 |
| Synergistetes | Synergistia | Synergistales | Synergistaceae | *Dethiosulfovibrio* | 0.002 | 0.000 | 0.000 | 0.000 | 0.000 | 0.000 |
| Synergistetes | Synergistia | Synergistales | Synergistaceae | Digester cluster 2 | 0.006 | 0.000 | 0.000 | 0.000 | 0.004 | 0.005 |
| Synergistetes | Synergistia | Synergistales | Synergistaceae | Digester cluster 3 | 0.002 | 0.000 | 0.000 | 0.000 | 0.008 | 0.002 |
| Synergistetes | Synergistia | Synergistales | Synergistaceae | Digester cluster 4 | 0.001 | 0.000 | 0.000 | 0.000 | 0.000 | 0.000 |
| Synergistetes | Synergistia | Synergistales | Synergistaceae | Digester cluster 5 | 0.010 | 0.000 | 0.000 | 0.000 | 0.000 | 0.000 |
| Synergistetes | Synergistia | Synergistales | Synergistaceae | Gut digester cluster | 0.000 | 0.000 | 0.000 | 0.000 | 0.000 | 0.001 |
| Synergistetes | Synergistia | Synergistales | Synergistaceae | *Lactivibrio* | 0.001 | 0.000 | 0.000 | 0.004 | 0.000 | 0.000 |
| Synergistetes | Synergistia | Synergistales | Synergistaceae | Oral cluster | 0.000 | 0.000 | 0.000 | 0.004 | 0.000 | 0.000 |
| Synergistetes | Synergistia | Synergistales | Synergistaceae | Termite cockroach cluster | 3.727 | 3.349 | 0.013 | 0.755 | 3.827 | 2.628 |
| Synergistetes | Synergistia | Synergistales | Synergistaceae | *Thermanaerovibrio* | 0.001 | 0.000 | 0.000 | 0.000 | 0.000 | 0.000 |
| Synergistetes | Synergistia | Synergistales | Synergistaceae | *Thermovirga* | 0.002 | 0.000 | 0.000 | 0.000 | 0.000 | 0.003 |
| TA06 | Unclassified | Unclassified | Unclassified | Unclassified | 0.001 | 0.000 | 0.000 | 0.000 | 0.000 | 0.000 |
| Tenericutes | Mollicutes | Acholeplasmatales | Acholeplasmataceae | *Acholeplasma* | 0.000 | 0.000 | 0.000 | 0.007 | 0.000 | 0.000 |
| Tenericutes | Mollicutes | Acholeplasmatales | Acholeplasmataceae | *Acholeplasma* 2 | 0.000 | 0.000 | 0.000 | 0.004 | 0.000 | 0.000 |
| Tenericutes | Mollicutes | Anaeroplasmatales | Anaeroplasmataceae | *Anaeroplasma* | 0.002 | 0.000 | 0.000 | 0.185 | 0.000 | 0.000 |
| Tenericutes | Mollicutes | Candidatus Phytoplasma | Candidatus Phytoplasma | *Candidatus* Phytoplasma | 0.000 | 0.000 | 0.000 | 0.014 | 0.000 | 0.000 |
| Tenericutes | Mollicutes | Entomoplasmatales | Spiroplasmataceae | *Spiroplasma* | 0.009 | 0.000 | 0.039 | 0.004 | 0.012 | 0.015 |
| Tenericutes | Mollicutes | Mycoplasmatales | Mycoplasmataceae | *Candidatus* Bacilloplasma | 0.000 | 0.000 | 0.000 | 0.000 | 0.000 | 0.001 |

**Table S2 continued.**

| | | | | | | | | | | |
|---|---|---|---|---|---|---|---|---|---|---|
| Tenericutes | Mollicutes | Mycoplasmatales | Mycoplasmataceae | *Mycoplasma* | 0.004 | 0.000 | 0.284 | 0.007 | 0.045 | 0.030 |
| Tenericutes | Mollicutes | Mycoplasmatales | Mycoplasmataceae | *Mycoplasma* 2 | 0.000 | 0.000 | 0.064 | 0.036 | 0.000 | 0.000 |
| Tenericutes | Mollicutes | RF9 | Unclassified | Unclassified | 0.000 | 0.319 | 0.013 | 0.289 | 0.008 | 0.005 |
| Tenericutes | Mollicutes | RF9 | Unclassified | Unclassified | 0.291 | 0.000 | 0.000 | 0.000 | 0.000 | 0.000 |
| Tenericutes | Mollicutes | Unclassified | Unclassified | Unclassified | 0.000 | 0.255 | 0.000 | 0.007 | 0.000 | 0.000 |
| Thaumarchaeota | Nitrososphaeria | Nitrososphaerales | Nitrososphaeraceae | *Nitrososphaera* | 0.000 | 0.000 | 0.000 | 0.011 | 0.000 | 0.000 |
| Thermodesulfobacteria | Thermodesulfobacteria | Thermodesulfobacteriales | Thermodesulfobacteriaceae | *Thermodesulfatator* | 0.001 | 0.000 | 0.000 | 0.000 | 0.000 | 0.000 |
| Thermotogae | Thermotogae | Thermotogales | Thermotogaceae | Uncultured 1 | 0.002 | 0.000 | 0.000 | 0.000 | 0.000 | 0.000 |
| Unclassified | Unclassified | Unclassified | Unclassified | *Bactoderma* | 0.000 | 0.000 | 0.000 | 0.004 | 0.000 | 0.000 |
| Unclassified | Unclassified | Unclassified | Unclassified | *Caldisphaera* | 0.000 | 0.032 | 0.000 | 0.000 | 0.000 | 0.000 |
| Unclassified | Unclassified | Unclassified | Unclassified | *Duganella* | 0.000 | 0.000 | 0.000 | 0.000 | 0.000 | 0.001 |
| Unclassified | Unclassified | Unclassified | Unclassified | *Elusimicrobium* | 0.000 | 0.000 | 0.000 | 0.000 | 0.000 | 0.001 |
| Unclassified | Unclassified | Unclassified | Unclassified | *Gordonia* | 0.000 | 0.032 | 0.000 | 0.004 | 0.000 | 0.000 |
| Unclassified | Unclassified | Unclassified | Unclassified | *Halarchaeum* | 0.000 | 0.000 | 0.000 | 0.004 | 0.000 | 0.000 |
| Unclassified | Unclassified | Unclassified | Unclassified | *Haloplasma* | 0.000 | 0.000 | 0.000 | 0.000 | 0.000 | 0.001 |
| Unclassified | Unclassified | Unclassified | Unclassified | *Halorhodospira* | 0.000 | 0.000 | 0.000 | 0.000 | 0.000 | 0.001 |
| Unclassified | Unclassified | Unclassified | Unclassified | *Humibacter* | 0.000 | 0.000 | 0.000 | 0.000 | 0.000 | 0.001 |
| Unclassified | Unclassified | Unclassified | Unclassified | *Ideonella* | 0.000 | 0.000 | 0.000 | 0.000 | 0.000 | 0.001 |
| Unclassified | Unclassified | Unclassified | Unclassified | *Ketogulonicigenium* | 0.000 | 0.000 | 0.000 | 0.000 | 0.000 | 0.001 |
| Unclassified | Unclassified | Unclassified | Unclassified | *Methanimicrococcus* | 0.000 | 1.085 | 0.000 | 0.135 | 0.000 | 0.000 |
| Unclassified | Unclassified | Unclassified | Unclassified | *Methanobrevibacter* | 0.000 | 0.032 | 0.000 | 0.007 | 0.000 | 0.001 |
| Unclassified | Unclassified | Unclassified | Unclassified | *Methanocella* | 0.000 | 0.000 | 0.000 | 0.007 | 0.000 | 0.000 |
| Unclassified | Unclassified | Unclassified | Unclassified | *Methanohalophilus* | 0.000 | 0.032 | 0.000 | 0.007 | 0.000 | 0.000 |
| Unclassified | Unclassified | Unclassified | Unclassified | *Methanomassiliicoccus* | 0.000 | 0.606 | 0.000 | 0.150 | 0.163 | 0.000 |
| Unclassified | Unclassified | Unclassified | Unclassified | *Pyrococcus* | 0.000 | 0.032 | 0.000 | 0.000 | 0.000 | 0.000 |
| Unclassified | Unclassified | Unclassified | Unclassified | *Thermogymnomonas* | 0.000 | 0.000 | 0.000 | 0.004 | 0.000 | 0.000 |
| Unclassified | Unclassified | Unclassified | Unclassified | Unclassified | 0.000 | 0.000 | 0.000 | 0.000 | 0.000 | 0.001 |

**Table S2 continued.**

| | | | | | | | | | | |
|---|---|---|---|---|---|---|---|---|---|---|
| Unidentified | Unidentified | Unidentified | Unidentified | Unidentified | 0.708 | 0.000 | 0.000 | 0.000 | 0.000 | 0.086 |
| Verrucomicrobia | Acidimethylosilex | Unclassified | Unclassified | Unclassified | 0.000 | 0.000 | 0.000 | 0.004 | 0.004 | 0.000 |
| Verrucomicrobia | OPB35 soil group | Unclassified | Unclassified | Unclassified | 0.000 | 0.000 | 0.000 | 0.110 | 0.000 | 0.000 |
| Verrucomicrobia | Opitutae | Opitutales | Opitutaceae | *Lacunisphaera* | 0.000 | 0.000 | 0.000 | 0.004 | 0.000 | 0.000 |
| Verrucomicrobia | Opitutae | Opitutales | Opitutaceae | *Opitutus* | 0.056 | 0.000 | 0.000 | 0.025 | 0.004 | 0.001 |
| Verrucomicrobia | Opitutae | Unclassified | Unclassified | Unclassified | 0.001 | 0.000 | 0.000 | 0.000 | 0.000 | 0.000 |
| Verrucomicrobia | Spartobacteria | Chthoniobacter | Unclassified | Unclassified | 0.000 | 0.000 | 0.000 | 0.039 | 0.000 | 0.000 |
| Verrucomicrobia | Spartobacteria | Chthoniobacterales | Xiphinematobacteraceae | *Candidatus* Xiphinematobacter | 0.000 | 0.000 | 0.000 | 0.036 | 0.000 | 0.000 |
| Verrucomicrobia | Spartobacteria | DA101 soil group | Unclassified | Unclassified | 0.000 | 0.000 | 0.000 | 0.011 | 0.000 | 0.000 |
| Verrucomicrobia | Spartobacteria | FukuN18 freshwater group | Unclassified | Unclassified | 0.000 | 0.000 | 0.000 | 0.082 | 0.000 | 0.000 |
| Verrucomicrobia | Spartobacteria | Unclassified | Unclassified | *Terrimicrobium* | 0.000 | 0.000 | 0.000 | 0.007 | 0.000 | 0.000 |
| Verrucomicrobia | Verrucomicrobiae | Verrucomicrobiales | DEV007 | Unclassified | 0.001 | 0.000 | 0.000 | 0.000 | 0.000 | 0.000 |
| Verrucomicrobia | Verrucomicrobiae | Verrucomicrobiales | Verrucomicrobiaceae | *Akkermansia* | 0.000 | 0.000 | 0.013 | 0.021 | 0.000 | 0.000 |
| Verrucomicrobia | Verrucomicrobiae | Verrucomicrobiales | Verrucomicrobiaceae 1 | *Haloferula* | 0.000 | 0.000 | 0.000 | 0.004 | 0.000 | 0.000 |
| Verrucomicrobia | Verrucomicrobiae | Verrucomicrobiales | Verrucomicrobiaceae 1 | Uncultured 3 | 0.000 | 0.000 | 0.000 | 0.004 | 0.000 | 0.000 |
| Verrucomicrobia | Verrucomicrobiae | Verrucomicrobiales | Verrucomicrobiaceae 2 | *Prosthecobacter* | 0.000 | 0.000 | 0.000 | 0.011 | 0.000 | 0.000 |
| Verrucomicrobia | Verrucomicrobiae | Verrucomicrobiales | Verrucomicrobiaceae 2 | *Verrucomicrobium* | 0.000 | 0.000 | 0.000 | 0.004 | 0.000 | 0.000 |
| WCHB1-60 | Unclassified | Unclassified | Unclassified | Unclassified | 0.000 | 0.000 | 0.000 | 0.007 | 0.000 | 0.000 |
